# ANDES: a novel best-match approach for enhancing gene set analysis in embedding spaces

**DOI:** 10.1101/2023.11.21.568145

**Authors:** Lechuan Li, Ruth Dannenfelser, Charlie Cruz, Vicky Yao

**Affiliations:** Department of Computer Science, Rice University

## Abstract

Embedding methods have emerged as a valuable class of approaches for distilling essential information from complex high-dimensional data into more accessible lower-dimensional spaces. Applications of embedding methods to biological data have demonstrated that gene embeddings can effectively capture physical, structural, and functional relationships between genes. However, this utility has been primarily realized by using gene embeddings for downstream machine learning tasks. Much less has been done to examine the embeddings directly, especially analyses of gene sets in embedding spaces. Here, we propose ANDES, a novel best-match approach that can be used with existing gene embeddings to compare gene sets while reconciling gene set diversity. This intuitive method has important downstream implications for improving the utility of embedding spaces for various tasks. Specifically, we show how ANDES, when applied to different gene embeddings encoding protein-protein interactions, can be used as a novel overrepresentation-based and rank-based gene set enrichment analysis method that achieves state-of-the-art performance. Additionally, ANDES can use multi-organism joint gene embeddings to facilitate functional knowledge transfer across organisms, allowing for phenotype mapping across model systems. Our flexible, straightforward best-match methodology can be extended to other embedding spaces with diverse community structures between set elements.

## Introduction

Methods to build embedding representations have become ubiquitous in diverse fields spanning text-based [1, 2], image-based [3, 4], and domain-specific [5–10] applications. In addition to the computational benefits that lower-dimension embedding representations provide, there is an implicit assumption that a quality embedding amplifies the important signal in the data while reducing noise. In the biomedical realm, gene embeddings are gaining traction as a valuable approach for predicting function [11–13], disease associations [14, 15], expanding gene set representations [16, 17], among other applications [18–21].

Given the utility of gene embeddings for downstream machine learning tasks, it is intuitive that gene embeddings capture important gene-gene relationship information (Figure 1A). However, little attention is paid to exploring the resulting embedding spaces, especially for the analyses of gene sets. In standard genomics analyses, gene sets (e.g., a set of differentially expressed genes, reported GWAS genes, or even a group of genes annotated to a particular pathway) are often a fundamental “functional unit.” Comparisons between sets are very routine, including gene set enrichment analysis [22, 23], disease-gene associations [24, 25], and drug repurposing [26, 27]. Yet, there appears to be limited to no research considering gene set comparisons in the context of embedding spaces.

**Figure 1.**
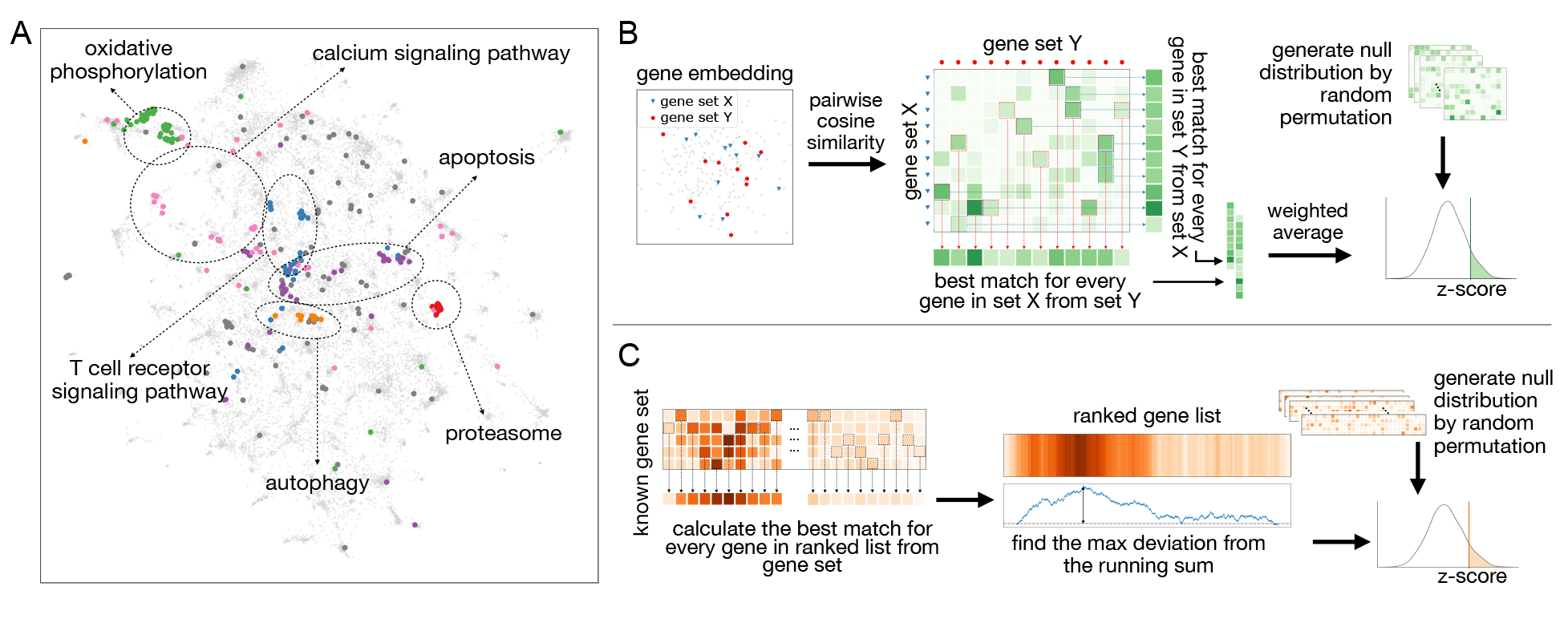
Overview of ANDES. **(A)** A UMAP plot of the node2vec gene embedding for a human PPI network with a set of Alzheimer’s disease genes (hsa05010) highlighted. Within this set of disease genes, several sub-clustered biological processes representing diverse biological functions are scattered across the embedding space. ANDES is capable of considering this functional diversity when matching gene sets. **(B)** Overview of the ANDES set similarity framework. Given two gene sets, ANDES first calculates the pairwise cosine similarity between every gene in each of the two sets. Based on the underlying similarity matrix, ANDES finds the best match for every gene (in both directions), then calculates the weighted average (taking into account gene set size) to yield a single score. Statistical significance is estimated using a cardinality-aware null distribution. **(C)** Overview of the ANDES rank-based gene set enrichment method. Given a ranked gene list based on an experimental result and a known gene set, ANDES calculates the best-match similarity for every gene in the ranked list. Walking down the ranked list, ANDES finds the max deviation from the running sum. The final enrichment score is also estimated using a cardinality-aware null distribution.

Here, we present an Algorithm for Network Data Embedding and Similarity analysis (ANDES) (Figure 1). The goal of ANDES is to calculate an interpretable measure of gene set similarity that accounts for the presence of functional diversity. Towards this end, ANDES identifies best-matching (most similar) genes between two sets reciprocally and calculates a score based on the embedding distances between these best-match similarities (Figure 1B). This best-match concept has parallels to an early method proposed for biological text mining [28], but functional analyses in embedding spaces require adjustments for biases due to gene set cardinalities. ANDES thus further incorporates a statistical significance estimation procedure that estimates the null distribution through Monte Carlo sampling to ensure comparable similarity estimations across different pairs of sets.

Outside of biological use cases, previous methods to summarize set relationships in embedding spaces typically consider variations of averaging embedding information across set members, ultimately ignoring the potential diversity within the set. One such averaging approach simply uses the centroid of all considered entities in the embedding space [29, 30]. However, gene sets, especially ones of interest, often contain a mixture of signals (e.g., a disease-associated gene set may include dysregulated genes from several pathways). The similarity calculated from two gene set centroids would fail to consider different subfunctional groups in the gene set and instead obscure this signal, especially when the subgroups are distinct. Though network-based gene set comparison methods have typically not been applied to embedding spaces, they can sometimes transfer to this domain. One such network-based method formulates the gene set comparison problem as a *t*-test between the two gene sets, with a permutation-based background correction for the size of each gene set [31]. However, while the corrected *t*-test method does take into account the variability of the gene set, it is ultimately still anchored in a comparison of means.

Gene set enrichment is a natural extension of the set-matching abilities of ANDES. Gene set enrichment methods fall into two main classes: overrepresentation-based approaches [22] and ranked-based approaches [23, 32], of which hypergeometric test and Gene Set Enrichment Analysis (GSEA) are respectively representative methods [23]. One of the fundamental limitations of both categories of methods is a complete reliance on gene annotations (e.g., functional annotations in the Gene Ontology); if a gene has no annotations, it cannot contribute to the enrichment analysis. Considering genes in functionally meaningful embedding spaces is one way to circumvent this limitation. ANDES can be used directly as an overrepresentation-based approach, and we also extend ANDES’ best-match approach for rank-based gene set enrichment (Figure 1C).

Other recent methods that attempt to lessen the dependency of gene set enrichment on existing annotations include Network-Based Gene Set Enrichment Analysis (NGSEA), which reranks the input gene list by incorporating the mean of its network neighbors [33], and Gene Set Proximity Analysis (GSPA), which allows users to supplement gene set annotations using a radius in an embedding space [34]. Unfortunately, we are unable to systematically evaluate NGSEA because it is only available as a web portal, and we cannot change the underlying gene sets used.

Through a series of evaluations, we demonstrate that ANDES can better estimate gene set functional similarity in gene embedding spaces compared to existing average-based methods. Furthermore, ANDES outperforms previous methods for gene set enrichment, and we also showcase its ability to prioritize new candidates for drug repurposing. Overall, we find that leveraging the intuition of best-match comparisons is an effective, generalizable approach that has important implications for interpretable analyses of embedding spaces.

## Results

### ANDES outperforms other set comparison approaches by effectively capturing substructure within gene sets

We first explore the extent to which ANDES and other set comparison methods can recover “functionally matched” gene sets in embedding spaces. “Matched” gene sets describe the same biological processes across different databases, such as KEGG pathways and Gene Ontology (GO) biological processes. When examining these gene sets in an embedding space that captures gene relationships, we expect a good set comparison method to identify these matches. We note that the same process described in KEGG and GO can naturally have overlapping genes, which would alter this set matching problem into an easier one primarily driven by set overlap. To prevent this, we only keep overlapping genes in the KEGG gene sets and remove them from GO gene sets. We compare ANDES against the mean embedding, mean score, and corrected t-score methods. Mean embedding and mean score are two intuitive approaches for set comparisons in embedding spaces, and the corrected t-score method has been used for set comparisons in functional network representations [31]. To the best of our knowledge, these represent the current scope of embedding set comparison methods, highlighting the lack of method development for this problem.

All set comparison approaches are agnostic to the type of embedding method used. To see if the type of embedding impacts performance, we use three different methods to generate gene embeddings from a protein-protein interaction (PPI) network: node2vec [35], NetMF [36], and a structure-preserving autoencoder method based on the architecture in [10]. ANDES consistently outperforms other methods and successfully identifies gene sets with similar functional roles, regardless of the underlying gene embedding method (Figure 2A). More specifically, ANDES significantly outperforms the mean score (node2vec: p=1.84 × 10^−6^, NetMF: 2.75 × 10^−3^, NN: p=3.14 × 10^−5^, Wilcoxon signed-rank test) and corrected t-score method (node2vec: p=7.56 × 10^−6^, NetMF: p=2.06 × 10^−4^, NN: p=2.23 × 10^−7^, Wilcoxon signed-rank test). ANDES also significantly outperforms the mean embedding method for the node2vec and autoencoder embedding methods (node2vec: p=0.039, NetMF: p=0.117, NN: p=0.015, Wilcoxon signed-rank test), and in general, ANDES is more stable. We note that if we do not remove the overlapping genes between matching KEGG and GO terms, ANDES shows an even larger performance advantage compared to existing methods (Figure S1).

**Figure 2.**
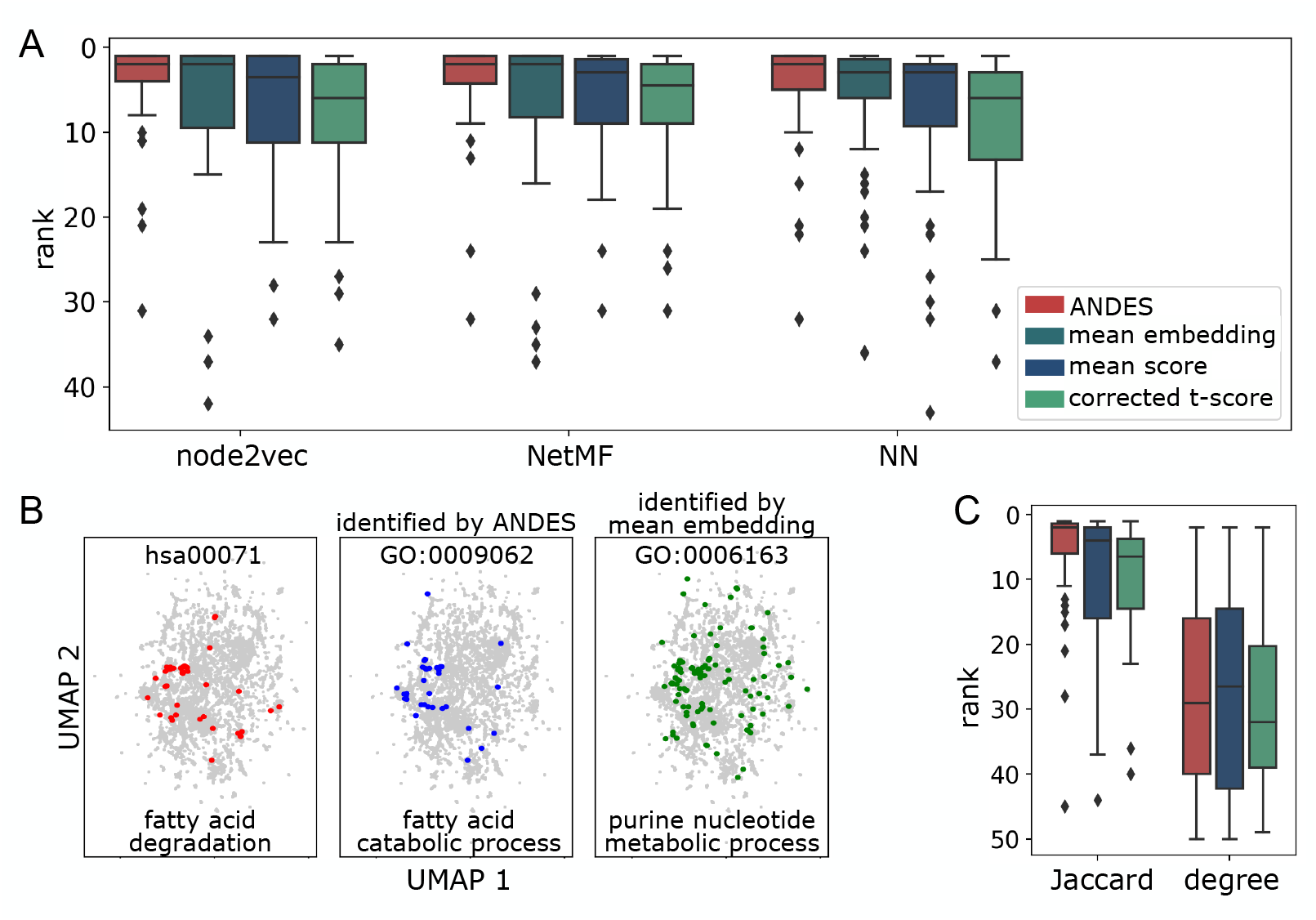
ANDES better matches gene sets that describe the same biological processes regardless of the underlying embedding or network structure. **(A)** Boxplots of the ranking of the correct matching GO term for 50 KEGG terms demonstrate that ANDES outperforms the mean embedding, corrected t-score, and mean score methods across three network embedding approaches (node2vec, NetMF, Neural Network (NN)). NN is a structure-aware autoencoder method (Methods). We also note that the handful of KEGG-GO pairs where ANDES performs poorly have consistently poor performance across methods (e.g., none of the 5 ANDES outlier KEGG terms in node2vec achieve a better ranking in any other methods). **(B)** UMAP of the node2vec PPI network embedding of genes in the KEGG fatty acid degradation gene set highlights a failure of the mean embedding method to capture meaningful substructure. Inspection of the embedding space reveals a similar substructure between the correct KEGG-GO term match prioritized by ANDES that is not seen in the top match for the mean embedding method. **(C)** Baseline approach for gene set matching in PPI networks. Matched KEGG-GO terms are ranked using pairwise similarity based on gene neighbor Jaccard similarity (Jaccard), or more naively, by the sum of node degrees (degree). Since these pairwise similarity matrices are directly calculated from network properties without using embeddings, we cannot calculate the mean embedding method and instead compare ANDES to only the corrected t-score and mean score.

Practically, we observe that the mean embedding method can have more extreme failure cases compared to all other methods, likely due to the inherent limitation of collapsing information from all genes in the gene set into a single mean embedding before subsequent comparisons. As an example, we show one specific failure of mean embedding that occurs with a matched pair of KEGG and GO terms (KEGG: hsa00071-fatty acid degradation, GO: GO:0009062-fatty acid catabolic process) (Figure 2B). A direct inspection of the distribution of each gene set’s genes in the embedding space (Figure 2B) quickly reveals that the correct KEGG-GO match has distinctly more similar embeddings, which ANDES can correctly identify and mean embedding cannot. Both the mean score and corrected t-score methods also rank the correct term higher than the mean embedding method.

Since ANDES’ best-match framework is not limited to using only embeddings as input, we also examine the extent to which matching gene sets can be identified using naive non-embedding network-only approaches, such as shared neighbor profiles, graph diffusion, and node degrees, using the original PPI network. Because these approaches do not use embeddings, it is impossible to calculate mean embeddings for the following comparisons. Using the Jaccard index of shared neighbors between two genes also captures sufficient functional signal to identify several correct KEGG-GO matches. In this setup, ANDES still significantly outperforms the mean score (p=1.22 × 10^−5^, Wilcoxon signed-rank test) and corrected t-score (p=2.19 × 10^−4^, Wilcoxon signed-rank test) methods (Figure 2C). We observe similar performance trends using a heat diffusion kernel on the PPI network, though the difference between ANDES and mean score is not significant (mean score: p=0.158, corrected t-score: p=4.12 × 10^−5^, Figure S2). Although both shared neighbor profiles and heat diffusion clearly capture functional signal, we note that using embedding approaches such as node2vec as input still leads to better performance (Jaccard: p=6.52 × 10^−3^, heat diffusion: p=7.82 × 10^−3^, Wilcoxon signed-rank test). Using a more naive exponential diffusion kernel and the simple sum of node degrees in the PPI network to measure gene similarity results in nearly random performance for all 3 methods explored in this comparison. In general, we observe that while ANDES can be successfully applied to these non-embedding approaches, the sparsity in the resultant PPI similarity matrix may be a limiting factor. Unlike gene similarity matrices based on embedding spaces, gene pairs with weak relationships have scores of zero using non-embedding approaches, which decreases the stability of the results. Altogether, this set of analyses demonstrates that the application of ANDES is not limited to only embedding spaces, but ANDES’ performance improves with the informativeness of the similarity matrix that is used as input.

### ANDES as a novel overrepresentation-based and rank-based gene set enrichment method

Since ANDES is a general framework to compare sets, comparing a single gene set of interest against a collection of annotated gene sets (e.g., KEGG) is directly comparable to existing overrepresentation-based approaches. We also further extend ANDES to handle comparisons of a ranked list of genes to gene sets, which allows ANDES to be used as a rank-based gene set enrichment approach (Figure 1C).

Here, our evaluations use a well-established gene set enrichment benchmark (GEO2KEGG [37]). GEO2KEGG annotates differential expression results (including FDR and log-fold-change per gene) to related pathways for 42 microarray studies, covering over 200 KEGG pathways. The rich data in GEO2KEGG enables us to test the recovery of corresponding KEGG pathways for each dataset through overrepresentation-based enrichment (using significance or fold change cutoffs) as well as rank-based gene set enrichment (by ranking all genes based on their fold changes). For the overrepresentation case, we compare ANDES performance with the hypergeometric test [22], considering gene sets of interest as differentially expressed genes per dataset (FDR≤0.05, keeping gene sets that are larger than 10 genes as is standard practice for overrepresentation analysis). In 22 out of 31 cases (71% datasets), ANDES set enrichment outperforms the hypergeometric test for KEGG pathway identification (p=6.78 × 10^−4^, Wilcoxon signed-rank test, Figure 3A). For the rank-based version, we compare ANDES with GSEA [23] and the embedding-based gene set enrichment method, GSPA [34]. Aggregating performance across all 42 datasets in the benchmark, ANDES’ rank-based gene set enrichment significantly improves over both GSEA (p=0.041; Wilcoxon signed-rank test) and GSPA (p=0.028; Wilcoxon signed-rank test) (Figure 3B). In general, we observe that ANDES consistently performs better than other methods, but especially when the original dataset has more samples (Figure S3). We thus also wanted to more carefully perform the enrichment analysis while ameliorating potential biases due to differing differential expression signals in the original GEO2KEGG datasets. Thus, for datasets with sufficient samples, we also further compute empirical p-values for enrichment scores by comparing against an estimated null distribution of the same dataset generated using 100 different condition label permutations. ANDES still outperforms GSEA (p=0.060; Wilcoxon signed-rank test) and GSPA (p=0.041; Wilcoxon signed-rank test) (Figure 3C), demonstrating ANDES’ ability of better prioritizing relevant functions based on true differential signals.

**Figure 3.**
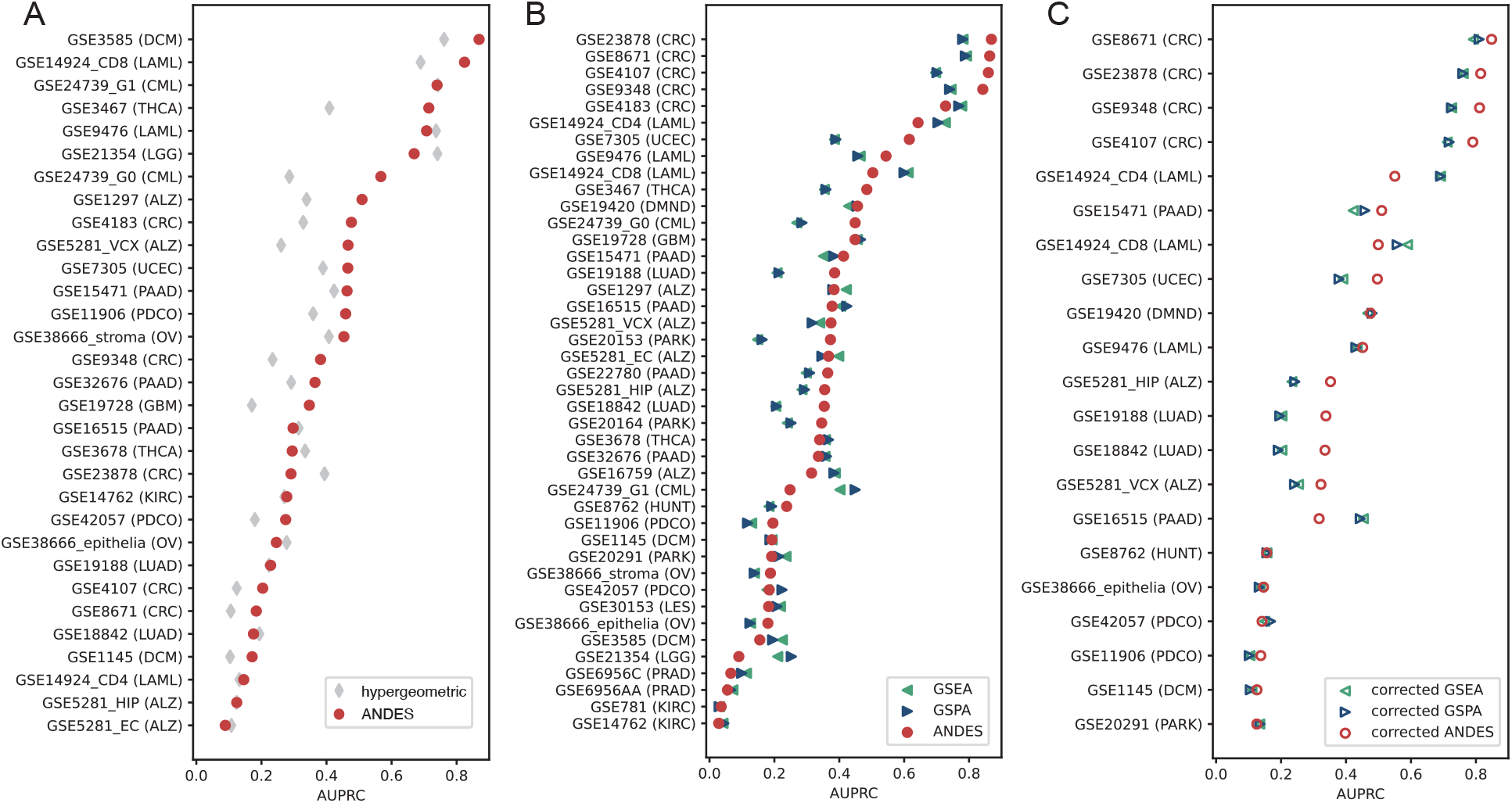
ANDES achieves state-of-the-art performance in overrepresentation-based and rank-based gene set enrichment for the GEO2KEGG [37] gene set enrichment benchmark. **(A)** Performance comparison between ANDES and hypergeometric test in retrieving annotated KEGG terms using genes that have FDR ≤ 0.05 in each dataset (where there are at least 10 genes that are significantly differentially expressed). **(B)** Performance comparison between ANDES, GSEA [23], and GSPA [34] in retrieving annotated KEGG terms using the full list of genes (no FDR cutoff), ranked by log_2_(fold change). In both cases, ANDES statistically outperforms other methods, demonstrating the advantage of incorporating gene embedding information using best-match into the gene set enrichment setting. **C**. Performance comparisons between ANDES, GSEA, and GSPA with empirically estimated p-values in retrieving annotated KEGG terms. Only expression datasets with at least 10 samples in both normal and diseased conditions (21 datasets) are included for sufficient variability in label permutations. corrected-ANDES still outperforms corrected-GSEA and corrected-GSPA

Together, these results highlight ANDES’ utility as a novel state-of-the-art method for overrepresentation-based and rank-based gene set enrichment. Furthermore, since ANDES utilizes gene embeddings, enrichment analyses can be performed in cases where existing annotations have low or even no overlap with the genes of interest, making it particularly valuable for the overrepresentation case.

### ANDES can be used to recapitulate known relationships between drugs and prioritize new candidates for drug repurposing

To highlight how ANDES can be used to discover new biology, we use ANDES to match disease gene sets from OMIM [38] with drug target genes from DrugBank [39]. This is a use case of practical importance, since the drug designing process for new compounds is quite laborious, involving many layers of development to ensure compound safety, delivery, efficacy, and stability. Thus, a computational effort that can potentially be used to repurpose a vetted compound can greatly help accelerate the development of new therapies. We first calculate a disease similarity profile for drugs using their ANDES scores, then apply dimensionality reduction using principal component analysis (PCA) on these profiles (Figure 4A, Figure S5). We see that, there are several “nervous system” drugs that are more distinct than others, as well as a subgroup that has overlap with other classes, including drugs that are “antineoplastic and immunomodulating agents.”

**Figure 4.**
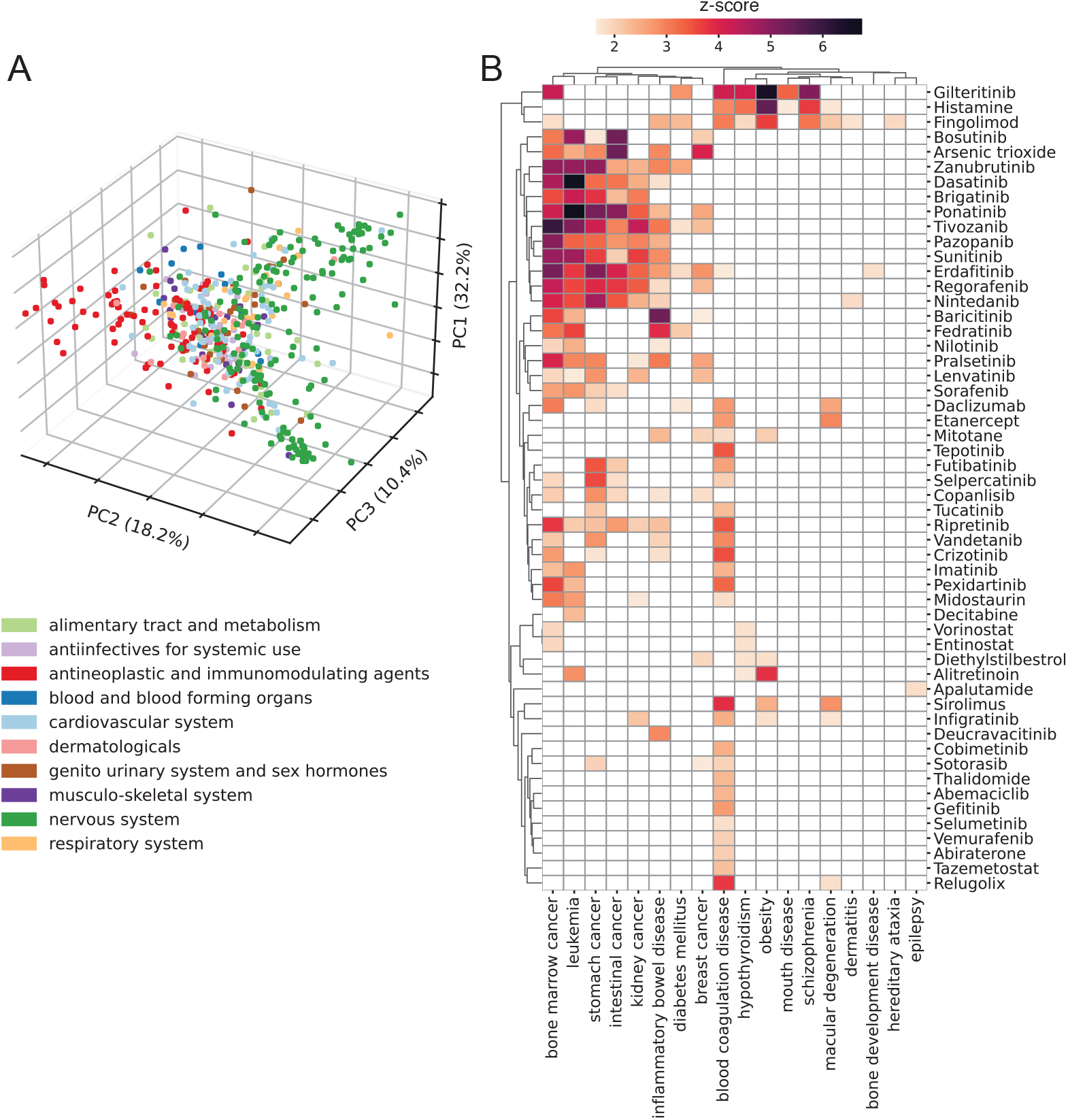
Analysis of drug-disease relationships using ANDES. **(A)** PCA plot of the first three PCs showing the relationship between drugs based on their association with diseases. Colors are based on the 1st level of ATC groups. The first three PCs explain 32.2%, 18.2%, and 10.4% of the total variance respectively. **(B)** Heatmap of ANDES gene set similarity *z*-scores (darker color indicates higher *z*-score) between diseases and drugs in the “antineoplastic and immunomodulating agents” therapeutic class. Diseases and drugs that have at least one association (*z*-score>1.64) are retained, yielding a heatmap of 18 diseases and 54 drugs.

As a proof of concept for highlighting avenues for drug repurposing, we take a closer look at the “antineoplastic and immunomodulating agents,” examining the ANDES gene set similarity scores for all drugs in this class (Figure 4B). Results for the other drug classes can be found in the supplement (Figures S6-17). Only diseases and drugs with at least one significant match (*z* > 1.64) are kept. Besides the most apparent cluster of cancers, ANDES also captures potentially novel indications or potential side effects. For example, ANDES predicts a strong association between obesity and two drugs, Histamine and Gilteritinib, which are well-documented associations. Specifically, Histamine can decrease hunger by affecting the appetite control center in the brain [40], and weight gain is a listed side effect of Gilteritinib [41]. While these are all known relationships, ANDES also predicts less-documented, potentially novel drug-disease relationships, such as a link between Fingolimod and schizophrenia. As recently as 2023, Fingolimod has been examined in rats for its potential to reverse schizophrenia phenotypes [42]. We observe a similar association between Sirolimus and macular degeneration. Sirolimus is an immunosuppressive agent used to prevent transplant rejection, but as of 2021, it has been found to have emerging promise as a therapeutic for age-related macular degeneration [43]. These two seemingly disparate indications likely share an inflammatory pathological pathway, which can be picked up with ANDES.

### ANDES can be effectively used with other types of gene embeddings, such as cross-organism embeddings for increasingly complex biological insights

We have already shown that ANDES performance is agnostic to the underlying embedding used, making it a modular framework. To make it even more powerful, we swap the human PPI-based gene embeddings for joint cross-organism gene embeddings using our previously published network embedding alignment method, ETNA [10]. This analysis highlights two general principles: (1) ANDES can still prioritize relevant signal when the underlying gene embedding is structurally more complex and (2) the scope of new biological insights can be expanded with the use of different embeddings.

Beyond showcasing the power of ANDES, being able to successfully map coordinated gene sets, such as pathways and processes between model organisms and humans, is an important problem. Model organisms are critical for studying aspects of human biology that are technically infeasible or unethical to study directly. Thus, improving functional knowledge transfer increases the potential impact of model system study.

To determine if ANDES can aid in functional knowledge transfer, we use ETNA to build three pairwise joint gene embeddings between humans and three model organisms: *M. musculus, D. melanogaster*, and *C. elegans*. ETNA’s joint embedding space enables the calculation of a similarity matrix for genes across species. Since genes across organisms can be annotated to the same GO terms, we can also evaluate to what extent the same GO term is prioritized in human when using the species-specific annotations in model organisms. For all three model organisms, ANDES consistently outperforms the mean embedding (*M. musculus*: p=4.30 × 10^−10^, *D. melanogaster*: p=0.147, *C. elegans*: p=2.28 × 10^−3^, Wilcoxon signed-rank test), mean score (*M. musculus*: p=1.01 × 10^−25^, *D. melanogaster*: p=1.19 × 10^−3^, *C. elegans*: p=9.88 × 10^−4^, Wilcoxon signed-rank test), and corrected t-score (*M. musculus*: p=3.66 × 10^−25^, *D. melanogaster*: p=3.62 × 10^−3^, *C. elegans*: p=2.27 × 10^−4^, Wilcoxon signed-rank test) (Figure 5A).

**Figure 5.**
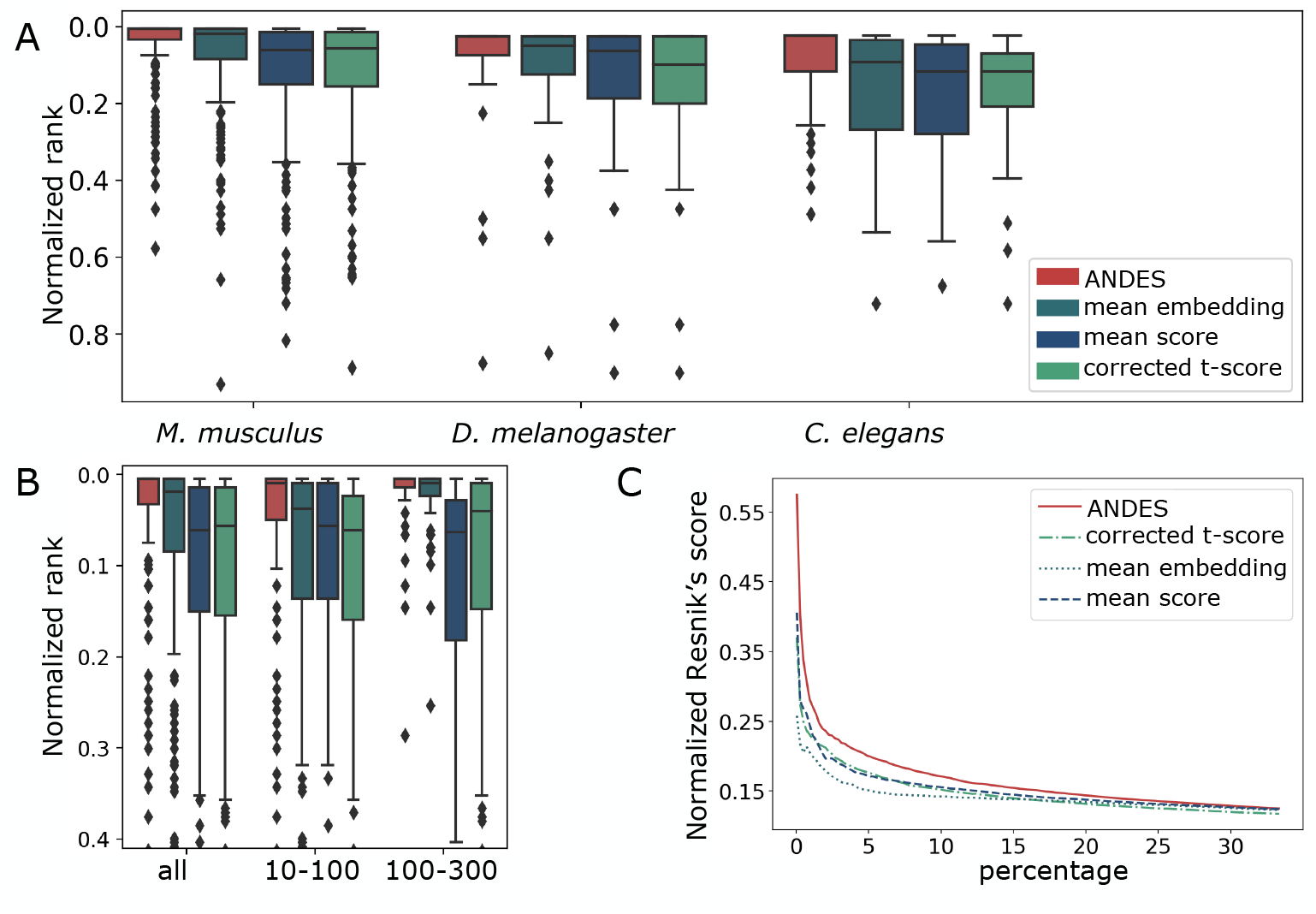
ANDES estimates gene set functional similarity across organisms better than existing methods. **(A)** Boxplot of the ranking of matched GO terms between human and three model organisms: *M. musculus, D. melanogaster*, and *C. elegans*, with 213, 40, and 43 shared GO slim terms, respectively. To facilitate comparison between organism pairs, the ranking is normalized by the number of shared GO terms. For each of the three organisms, ANDES consistently outperforms the mean embedding, corrected t-score, and mean score methods. **(B)** Boxplot of the ranking of matched GO terms for *H. sapiens* and *M. musculus*. Gene set pairs are grouped into two categories according to the sum of the number of genes in both gene sets (small [10-100] and large [101-300]). ANDES again consistently outperforms other methods regardless of gene set size. **(C)** Comparison of the cumulative average of the Resnik score walking down the ranked list for *H. sapiens* and *M. musculus*. ANDES consistently outperforms other methods until the score converges at ∼30% of all pairs.

When we group the GO terms shared between human and *M. musculus* by size, ANDES shows better performance, both when the gene set is large (mean embedding: p=0.023, mean score: 5.39 × 10^−11^, corrected t-score: 1.96 × 10^−12^, Wilcoxon signed-rank test) and small (mean embedding: p=5.91 × 10^−9^, mean score: 8.18 × 10^−17^, corrected t-score: 1.55 × 10^−14^, Wilcoxon signed-rank test). We notice that larger gene sets are easier to match across organisms (Figure 5B). We speculate that having more genes can result in more distinct patterns in the embedding space, leading to better mappings, especially for ANDES and the mean embedding method. Meanwhile, the mean score and corrected t-score methods are not able to take advantage of the additional information in larger gene sets and perform similarly. The mean embedding method performs poorly when the gene sets are small; this is likely because “outlier” genes can more easily skew the mean embedding, especially when there are distinct subprocesses within a gene set. Overall, ANDES is the only method that is a strong performer, consistently robust to gene set size.

So far, we have simplified the relationship between a pair of gene sets to simply “matched” or “unmatched,” but we can also evaluate unmatched terms based on how close they are to the correct target term in the ontology tree. To this end, we use Resnik measure [44], a semantic similarity measure leveraging the GO hierarchy, to quantify the similarity between two gene sets. Two gene sets that are close in the Gene Ontology are more likely to describe functionally similar biological processes and, therefore, have a higher Resnik score. Comparing predicted similarities for all gene sets between human and mouse, we calculate the cumulative average Resnik score of gene set pairs ranked by their similarity score in each set comparison method. Across all methods, the Resnik score is higher for the top-ranked pairs and gradually converges to randomness at around 30% of the ranked list of all pairs (Figure 5C). The trend also holds for *D. melanogaster* and *C. elegans* (Figure S18). Overall, ANDES consistently has the highest Resnik scores of all methods, demonstrating that it both identifies the exact match as well as other functionally related gene sets.

### Prioritizing mouse phenotypes for modeling human diseases with ANDES

After verifying that ANDES can recover conserved cross-organism functional signal (Figure 5), we further explore the potential of ANDES for cross-organism knowledge transfer. Phenotype prioritization is a vital aspect of effective knowledge transfer as some small phenotypic changes in the model organism (e.g., “decreased cervical vertebrae”) can be an important marker of the presence or extent of a human disease phenotype. Good matches here can be potential candidates for phenotypic screens. Towards this end, we systematically test associations between human disease gene sets from OMIM [38] and mouse phenotype gene sets from MGI [45]. We identify a range of significantly related disease-phenotype pairs, many of which merit further exploration (Figures S19-41). As a proof of concept, we show a smaller slim set of 13 human diseases that span a wide range of organ systems and pathological mechanisms, along with the top 5 associated mouse phenotypes (Figure 6).

**Figure 6.**
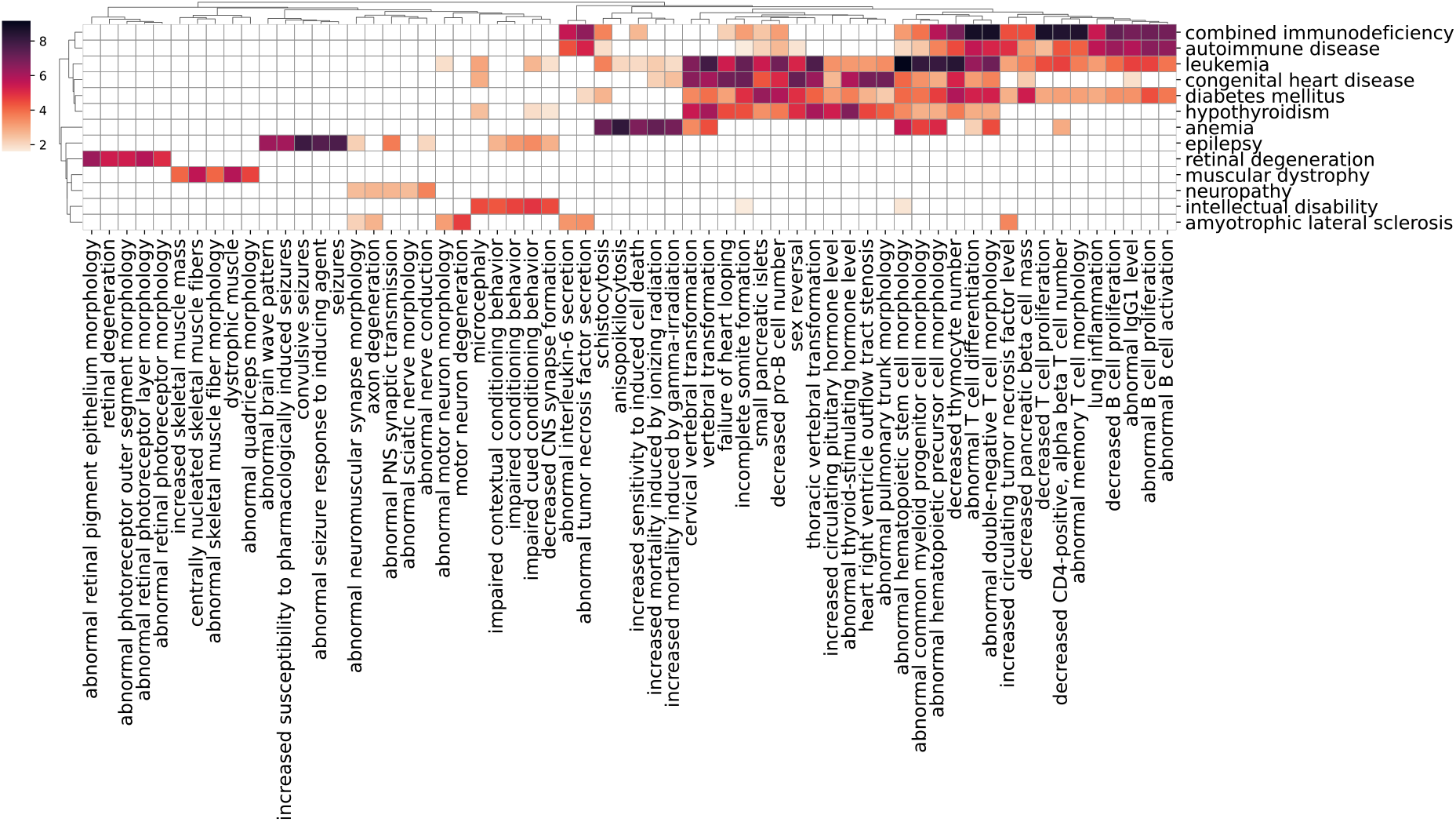
Heatmap of ANDES similarity scores for human disease and mouse phenotypes gene sets. Gene set similarity *z*-scores generated by ANDES are shown for 13 selected human diseases across various organ systems and pathological pathways. For each human disease, the top 5 mouse phenotypes predicted to be functionally similar are selected (62 mouse phenotypes). The intensity of color indicates the extent to which disease-phenotype associations exceed *z*-score=1.64 (corresponding to p-value=0.05).

While we do not have clear gold standards to evaluate mouse phenotype-human disease predictions, many of the disease-phenotype pairs we find make intuitive sense. Specifically, we find that diseases tend to cluster with ones that involve similar organ systems (e.g., combined immunodeficiency and autoimmune disease, Figure 6). Furthermore, phenotypes related to lymphocytes are associated with both immune diseases and leukemia, while “abnormal neuromuscular synapse morphology” is shared between neuropathy, epilepsy, and amyotrophic lateral sclerosis. Moreover, phenotype associations can also reflect differences between diseases related to the same organ system. Both epilepsy and intellectual disability’s top related phenotypes, “impaired conditioning behavior” [46], but seizure phenotypes are specific to epilepsy.

We also find that ANDES can capture both direct and secondary associations between human diseases and mouse phenotypes. For example, diabetes mellitus is related to the mouse phenotype “small pancreatic islets,” capturing the fact that these cells produce insulin (Figure 6). Furthermore, anemia, a disease with many variants that affect red blood cells, is enriched for the mouse phenotype, “anisopoikilocytosis,” a disorder where red blood cells have irregular sizes and shapes. ANDES can also identify secondary disease phenotypes not directly caused by the disease itself. For example, hypothyroidism is enriched in mouse phenotypes related to the brain and nervous system, which is a phenotype known to be associated with thyroid disease in humans [47]. Together, these results highlight the exciting potential of ANDES to not only model existing human-mouse phenotype mappings, but also identify new translational opportunities to aid in developing new model organism screens for specific human disease phenotypes.

## Discussion

Here, we introduce ANDES as a general-purpose method for comparing sets by considering best-match elements. In exploring ANDES’ application to gene embeddings, we have demonstrated how it can be used to prioritize functionally similar gene sets within a single organism or across organisms using more sophisticated joint embeddings. ANDES can leverage functional information from gene embeddings to avoid a complete reliance on gene annotations in both overrepresentation- and rank-based gene set enrichment analysis, and by doing so, achieves state-of-the-art performance. Unlike current embedding set similarity methods that rely on averaging, ANDES identifies the best matches between individual elements in a set, thus considering the diversity within a set to better capture inherent substructure that may be otherwise lost.

One limitation of our best-match approach is that it could potentially be sensitive to outliers. ANDES currently addresses this by combining information from all best-matching pairs and estimating statistical significance using cardinality-aware null distributions. But beyond these strategies, we note that we can also expand the ANDES framework to consider the top *k* matches per gene instead. The argument for using more matching genes would be to diminish the effect of outliers in driving the assessment of set similarity. This framing would place the best-match and mean score approaches at two ends of the spectrum with respect to choosing *k*, as *k* = 1 (a single element) is by definition the best-match approach, and *k* = 100% (all elements in both gene sets) is equivalent to the mean score approach. A cursory exploration of the effect of varying *k* on ANDES’ performance shows that the ability to identify functionally similar GO and KEGG gene sets using node2vec-embedded PPI networks decreases as *k* increases, eventually converging to the significantly lower mean score performance (Figure S42). Thus, at least with PPI embeddings, we find that using the best-match approach (i.e., *k* = 1) can avoid the introduction of an additional hyperparameter and performs well in practice. In other situations where outliers are of particular concern, it may be more worthwhile to examine the effect of varying *k*.

Another key aspect of ANDES is the similarity metric used to determine best matches. ANDES currently uses cosine similarity, but there are of course several possible alternatives, two natural ones being Euclidean distance and Pearson’s correlation. We find that using (inverse) Euclidean distance as the similarity metric results in the largest performance drop (Figure S43), likely due to the sensitivity of Euclidean distance to the magnitudes of the gene embeddings. Overall, the performance is similar when using the two scale-invariant methods (cosine similarity and Pearson’s correlation), though cosine similarity performs slightly better and would be our generally recommended metric.

Our novel ANDES framework has a myriad of downstream applications, especially when paired with different embeddings. Here, we only scratch the surface by showing the potential of ANDES for function prediction and drug repurposing when paired with a human gene embedding space, as well as cross-organism functional knowledge translation tasks when paired with a joint gene embedding. By matching phenotypes across human and mouse, we provide additional insight into opportunities for improved translational studies. We anticipate more interesting use cases with different gene embeddings or even embeddings of an entirely different modality. Furthermore, while we have analyzed several methods for generating PPI-based gene embeddings, integrating more gene information beyond PPIs may yield further improved gene set matching.

For example, genetic interaction ([48]), coexpression ([49]), and functional networks ([24]) can capture additional, complementary information with respect to gene functional similarity. Furthermore, since ubiquitously expressed genes can give rise to different disease-susceptibility and phenotypes in different tissues [50], using tissue-specific network models as input to ANDES could bring additional depth to the analyses and lead to new biological insights. We explore the possibility of extending ANDES’ enrichment analyses with tissue-specific functional networks [31], considering datasets from GEO2KEGG from five diseases: PDCO (Chronic Obstructive Pulmonary Disease), KIRC (Kidney Renal Clear Cell Carcinoma), PAAD (Pancreatic Adenocarcinoma), ALZ (Alzheimer’s Disease), and LAML (Acute Myeloid Leukemia), with their corresponding tissue-specific networks (lung, kidney, pancreas, brain, and blood, respectively). We find that using the tissue-specific networks consistently improves the enrichment for PDCO, KIRC, and PAAD. However, used directly, they do not result in much performance improvement on the ALZ and LAML datasets (Figure S44). For the LAML datasets, the CD8 samples in the GSE14924 dataset seem to benefit slightly from using the blood tissue network while the CD4 samples from the same dataset performs worse. These results suggest that the general blood tissue network may better capture and highlight CD8-related T cell signal rather than CD4 signal. In general, we reason that brain and blood may exhibit more regional and cell specificity with differing implications for disease, and thus using the general brain or blood tissue networks may not be specific enough to highlight the signal for ALZ and LAML. In these cases, more fine-grained tissue networks could be more beneficial. For example, Alzheimer’s disease modeling could benefit from using an entorhinal cortex network [51] and likewise for leukemia.

Beyond enrichment and set matching tasks, ANDES can also potentially be used to evaluate the quality of different embeddings when some set relationships are known *a priori*. In the gene embedding case, embedding spaces constructed in different ways might highlight different gene attributes. Analyzing known similarities (e.g., pathways, gene function, etc.) and how gene set matching changes might yield more unique insights into the information encoded in the latent space. A related extension for ANDES would be to further identify the best-match scores that drive different set matching results, thus providing gene-level insights for downstream analyses, similar to how network representations are used to provide functional insights for individual genes in [52].

Though we focus primarily on the utility of ANDES for gene embeddings, conceptually, ANDES’ best-match approach can be applied to any set comparisons as long as a corresponding similarity matrix for set elements exists, regardless of whether the input is an embedding representation. For PPI networks, we have found that embedding representations are better able to prioritize functionally similar gene sets than using the non-embedded network or network properties as input (Figure 2, Figure S2), but there could be alternative scenarios where the end-user is specifically interested in exploring set similarity using the non-embedded representation. In addition, by expanding the types of entities ANDES can be applied to, we foresee exciting applications for improved interpretability in other domains, such as protein language models ([53–55]) and single cell embeddings ([56–58]).

In conclusion, here we have presented a novel algorithm for set comparisons, ANDES, that can improve the utility and interpretability of analyses using embedding spaces. We hope that our best-match framework, paired with various embeddings, will be widely adopted and further adapted for additional novel use cases.

## Methods

As depicted in Figure 1A, a single gene set can comprise a mixture of different biological processes scattered throughout the embedding space. ANDES considers similarity while reconciling gene set diversity when measuring the relationship between two gene sets. Specifically, for each gene in a gene set, the method focuses on the “best-matching” gene (i.e., closest gene) in the other set, allowing ANDES to quantify the presence of functionally similar genes between the sets. The mean of these best-match scores represents the similarity between two gene sets. Finally, to enable systematic comparisons across different gene set pairs, background correction with an estimated null distribution is applied to standardize the scores.

### Calculation of the ANDES gene set similarity score

Given a low dimensional gene representation *E* ∈ ℝ^*n*×*d*^, where n is the number of genes and d is the embedding dimension, ANDES computes a pairwise similarity score between every pair of genes in the embedding. Here, we use cosine similarity, as it measures the angle between two vectors rather than only the distance and is scale-invariant, which means that it is not affected by the magnitude of the initial vector. This magnitude-invariance is an attractive feature because we find that the magnitude of embedding vectors is negatively correlated with the degree of the corresponding vertex in all three embedding methods used in our comparisons (Figure S45). Though degree has been shown to be a meaningful network property that relates to gene function, it can also capture study bias and is thus not preferable. Formally, *S* = cos(*E, E*) ∈ ℝ^*n*×*n*^, where each entry *S*_*i,j*_ of the matrix represents the similarity between two genes *i* and j. For two gene sets *X* = *x*_1_, *x*_2_, *x*_*m*_ and *Y* = *y*_1_, *y*_2_, *y*_*k*_, where |*X*| = *m* and |*Y* | = *k*, we define a similarity matrix for *X* and *Y* as *A* ∈ ℝ^*m*×*k*^, the sub-matrix of *S* with the corresponding entries matching genes from *X* and *Y* as rows and columns, respectively. The gene set similarity score (G*S*) is defined as:

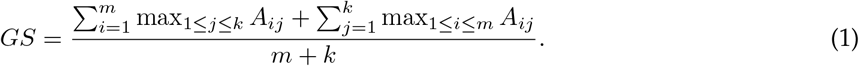

ANDES thus finds the best match for every gene in set *X* from set *Y* and vice versa. A large *GS* means most of the genes from *X* and *Y* can find similar genes to each other in the embedding space and are more likely to involve similar processes.

### Estimation of null distribution and statistical significance

ANDES uses an asymptotic approximation of Monte Carlo sampling to calculate a statistical significance score for every pair of gene sets. This procedure facilitates the comparison between different gene set pairs with varying numbers of genes (cardinalities). Since ANDES uses the max operator, this step is particularly crucial. In contrast to the mean operation, which has the same expected value for different random samples drawn from a Gaussian distribution, the expected maximum value will increase as set cardinality grows. Therefore, an appropriate cardinality-aware null distribution is essential for ANDES to eliminate bias resulting from varied gene set sizes.

The null distribution of the ANDES score between a pair of gene sets is approximated by 1,000 Monte Carlo samples, where each of the 2 sets has the same cardinality as the original pair. A restricted background gene list helps prevent the statistical significance of gene set similarities from becoming artificially inflated. While in most cases, the background can be all genes in the embedding, *E*, for systematic comparisons with a target annotation database (e.g., Gene Ontology (GO)), we use a more conservative background gene list that includes only genes that are in both the embedding and the target annotation database (e.g., genes with at least one GO annotation). To balance power and computational cost, we use a normal asymptotic approximation to estimate a *z*-score: 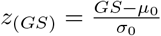, where *μ*_0_ and *σ*_0_ are the mean and standard deviation of the Monte Carlo approximations of the null distribution.

Since *G S*and *z*_(*GS*)_ can be used directly to quantify embedding-aware gene set similarities, they can be applied directly to gene set enrichment via overrepresentation analyses. In comparison with the standard Fisher’s exact test that is typically used for such comparisons, the immediate advantage of ANDES is that the overrepresentation analyses can identify significantly “related” gene sets even in the scenario where two gene sets of interest have completely no overlap.

### ANDES as a rank-based gene set enrichment method

In addition to overrepresentation analyses, we apply the best-match concept to develop a novel rank-based gene set enrichment method that considers distances between sets in gene embedding spaces. Given a ranked gene list *L* = *g*_1_, *g*_2_, … *g*_*l*_ and a gene set of interest *Y* = *y*_1_, *y*_2_, … *y*_*k*_, we define the similarity matrix for *L* and *Y* as *A* ∈ ℝ^*l*×*k*^, a sub-matrix of similarity matrix *S* with the corresponding entries matching genes from *L* and *Y* as rows and columns, respectively.

The best-match gene set enrichment score is:

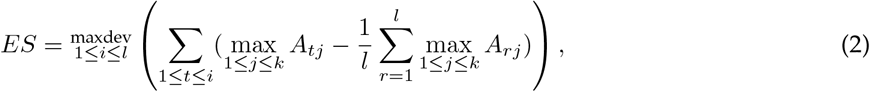

where maxdev is the maximum deviation from 0 and 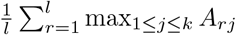 is the mean of all best-match scores from *L* to *Y* At each position *i* in *L*, ANDES is thus calculating the cumulative sum of mean-corrected best-match scores from genes *g*_1_, *g*_2_, … *g*_*i*_ to genes in *Y* As such, *ES* reflects the extent to which genes in *Y* are close (in embedding space) near the extremes (top or bottom) of the ranked gene list *L*. In this way, there are some similarities to the running-sum calculation used in the GSEA method [23], which updates an enrichment score based on the fraction of gene “hits” and “misses” as *L* is traversed. Similar to the limitation with Fisher’s exact test, GSEA only considers genes in *L* that have direct annotations in *Y* The best-match approach that ANDES takes is able to consider the extent to which each gene in *L* is close to genes in *Y* even if it does not have a direct annotation.

To assess the significance of *ES* for a given gene set *Y* ANDES uses an approach similar to that described above to calculate a normalized enrichment score (*NES*) through Monte Carlo sampling and asymptotic approximation, ensuring that the random gene sets have the same cardinality as *Y* Systematic comparisons against a target annotation database, such as GO, also use the more conservative background gene list (e.g., genes with at least one GO annotation).

### Gene embeddings using a protein-protein interaction network

While our proposed framework is agnostic to embedding method and data type, here we focus on gene embeddings generated from protein-protein interactions (PPIs). We use our previously assembled consensus PPI network [59], which considers physical interaction information from 8 different data sources, resulting in an unweighted and undirected network of 20,363 genes (vertices) and 822,311 interactions (edges). We compare three different embedding approaches: node2vec [35], NetMF [36], and a structure-preserving autoencoder method based on the architecture in [10], which we abbreviate as the neural network (NN) approach. Each method takes a different approach to embed the gene relationships characterized by the PPI network. Node2vec employs random walks on the graph, followed by the skip-gram model to embed node relationships. NetMF generates latent representations by solving a closed-form matrix factorization problem. Lastly, NN uses an autoencoder backbone with the following objective function:

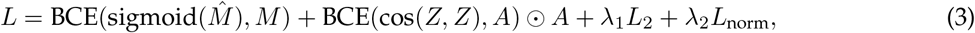

where *A* is adjacency matrix, *Z* is gene embedding matrix, *M* is the NetMF matrix, and *BCE* is binary cross entropy. The *L*_2_ norm on the autoencoder parameters is included to avoid overfitting and 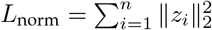 to avoid exploding norms. By optimizing this objective function, the neural network preserves the global and local structural information simultaneously. We fixed the embedding dimension sizes from all 3 methods to 128.

### Gene embeddings using tissue-specific functional networks

We use five human tissue-specific functional networks (lung, kidney, pancreas, brain, and blood) from GIANT [31], downloaded from humanbase.*i*o. Specifically, we use the “top edges” (edges with evidence supporting a tissue-specific functional interaction) as input to build the embeddings. In order to better utilize the weighted edge information in tissue specific networks, we use PecanPy’s [60] efficient node2vec+ [61] implementation. Node2vec+ is an extension of the node2vec method that demonstrats consistently strong performance in our benchmarks; the node2vec+ extension considers input edge weights during the random walk sampling process and thus is able to more fully take advantage of the functional networks.

### Gene set processing

For benchmarking, we use curated gene sets describing pathways, function, tissues, diseases, phenotypes, and drugs. We describe the gene sets and processing in the sections below.

#### Gene Ontology (GO)

To assess gene function, we use the biological process (BP) annotations from GO [62] (16 July 2020). To ensure high-quality annotations, we only keep terms with low-throughput experimental evidence codes (EXP, IDA, IMP, IGI, IEP). Furthermore, to avoid any circularity with the underlying PPIs used to construct embeddings, we exclude terms with evidence code IPI (Inferred from Physical Interaction). We further restrict the total number of GO terms using an expert-curated set of slim terms designed to emphasize key biological processes [31]. Leveraging the directed acyclic graph structure of the ontology, we propagate gene annotations from child terms to parent terms based on annotated “is a” and “part of” relationships; parent terms thus also contain genes that participate in more specific (child term) processes. After propagation, we apply a final filter to preserve terms with more than 10 and fewer than 300 annotated genes.

#### Kyoto Encyclopedia of Genes and Genomes (KEGG)

We obtain pathway gene sets from KEGG [63] using ConsensusPathDB [64]. In total, we obtain 333 unique human pathway gene sets.

#### Online Mendelian Inheritance in Man (OMIM)

We collect disease gene sets from OMIM (October 2023) [38]. These gene sets are then mapped to Disease Ontology [65] and propagated through the ontology structure, resulting in 284 unique disease gene sets.

#### Mouse Genome Informatics Phenotypes (MGI)

We assemble mouse phenotype gene sets from MGI (March 2022) [45]. After propagating genes from children to parents, we obtain 3,738 mouse phenotype gene sets.

#### DrugBank

Drug target information for 725 drugs, as well as other descriptions, such as ATC codes, were parsed from the academic licensed version of DrugBank [39] (Jan 2023).

### Benchmarking gene set similarity metrics in embedding spaces

The most straightforward way to compare sets in embedding space is to summarize their similarity through averaging. We compare ANDES with two variants of averaging (mean score and mean embedding) as a benchmark along with a corrected t-score approach [31].

Given two gene set embeddings *X* ∈ ℝ^*m*×*d*^ and *Y* ∈ ℝ^*k*×*d*^, where *m* and *k* are the number of genes in the set and d is the embedding dimension, the mean score method first calculates the pairwise cosine similarity *S* where 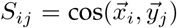 where 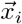 is the *i*-th row vector of *X* and 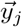 is the j-th row vector of *Y* The mean score method then simply computes the mean of these similarities,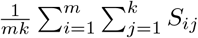.

The mean embedding approach instead first takes the average within a gene set, resulting in two gene set-level pooled embeddings 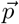 and 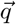 where

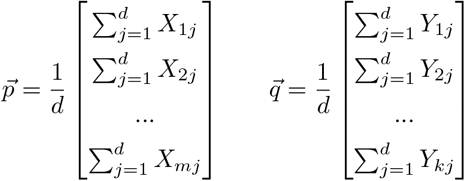

The final mean embedding score is the cosine similarity between the gene set-level pooled embeddings, cos 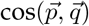.

The corrected t-score method [31] calculates an unequal variance *t*-test on two score distributions: the pairwise similarity score between two gene sets (between) and the scores associated with cardinality-matched gene sets across the genome (background). The final score is determined by comparing the between scores against a null distribution of the background scores.

We explore two gene set matching evaluations: (1) matching paired functional annotation datasets (KEGG and GO) with each other, and (2) matching GO gene sets across different model organisms. For the matched KEGG and GO comparisons, annotations for 50 KEGG pathways to corresponding GO biological processes are obtained based on the external database annotations from the KEGG web portal. To assess the ability to capture functional similarity beyond overlapping genes, we remove all overlapping genes between the KEGG- and GO-matched gene sets from GO gene sets when evaluating each method. In addition to comparing the different embedding methods using this evaluation paradigm, we also evaluate several baseline approaches that capture gene similarity based directly on the original PPI network, including shared neighbor profiles, graph diffusion (heat diffusion as well as exponential diffusion), and node degree. For the shared neighbor profile baseline, we use the Jaccard index of shared neighbors between a pair of genes as a measure of functional similarity. The heat diffusion analysis uses a diffusion step of 0.1 as recommended by [66]. Exponential diffusion and node degree constitute the most naive similarity approaches, where node degree is simply using the sum of a pair of genes’ degrees in the PPI as a measure of similarity.

For the cross-species evaluation, we look for exact matches between the same GO slim term in humans versus three model organisms: *M. musculus, S. cerevisiae, D. melanogaster*. While cross-species GO annotations can capture conserved biological processes, we need updated embedding spaces that jointly model genes from both species. To that end, we use our previously developed method, ETNA [10], to construct pairwise gene embedding spaces for human and each of the three model organisms. ETNA uses an autoencoder approach to generate within-species network embeddings based on PPI networks, then uses a cross-training approach with known orthologous genes as anchors to align the two embeddings into a joint embedding. This joint embedding enables cross-species comparisons of all genes represented in each PPI network.

### Evaluating gene set enrichment methods

To evaluate ANDES’ ability to identify functionally relevant pathways in an enrichment-analysis setting, we use a gold standard compendium of pathway-annotated gene expression data, GEO2KEGG, that has been routinely used for benchmarking gene set enrichment analyses [34, 37, 67]. GEO2KEGG consists of 42 human microarray profiles matched to various diseases, each of which has a set of curated KEGG pathway annotations. Using the corresponding annotated KEGG pathways as a gold standard, we can then calculate AUPRC for each of the 42 datasets. We compare the results of ANDES against three existing gene set analysis methods: hypergeometric test [22], GSEA [23], and GSPA [34]. We are unable to use NGSEA [33] and EnrichNet [68] for this benchmarking analysis because only web portals are available, with no ability to change the underlying gene sets for a fair comparison across methods. We note that GSPA does report that they outperform both NGSEA and EnrichNet. The comparison with hypergeometric test uses ANDES’ gene set similarity score and statistical significance calculation, taking genes with FDR≤0.05. Comparisons with rank-based gene set enrichment methods use ANDES’ best-match-based enrichment method, where the input gene list is ranked using log_2_(fold change).

To calculate empirical p-values to correct for potential biases in the amount of differential expression signal in the original dataset, we generate null distributions for each dataset with at least 10 samples in both normal and diseased conditions (21 datasets total) by permuting the sample labels 100 times. We include only datasets with at least 10 normal and 10 diseased samples to ensure more consistency across permutations. Using ANDES scores calculated on the datasets with permuted labels, we can then compute empirical p-values for each KEGG term and expression dataset pair by comparing the ANDES score based on the true dataset.

## Software Availability

Our implementation of ANDES and code for the analysis described herein is available on GitHub (https://github.com/ylaboratory/ANDES), released under the BSD 3-clause license for open source use, and also included as Supplemental Code.

## Competing Interest Statement

The authors declare no competing interests.

## Acknowledgements

The authors would like to thank members of the ylaboratory for helpful discussions.

## Funding

This work was supported by the Cancer Prevention & Research Institute of Texas (CPRIT RR190065) and the National Science Foundation (NSF DBI-2144534). VY is a CPRIT Scholar in Cancer Research.

## Author Contributions

LL and VY developed the method; LL implemented the method, with assistance in computational efficiency improvements from CC; LL performed methods evaluations; LL and RD applied the method to different case studies; VY supervised the study; LL, RD, and VY wrote the manuscript. All authors read and approved the final manuscript.

## Notes

https://github.com/ylaboratory/ANDES

### Competing Interest Statement

The authors have declared no competing interest.

### Summary of Updates

Expanded Discussion; Figure 3 expanded; updated all paired hypothesis tests to consistently use wilcoxon signed-rank test; updated supplemental files

## References

1. Church, K. W. Word2Vec. Natural Language Engineering 23, 155–162 (2017).

2. Devlin, J., Chang, M.-W., Lee, K. & Toutanova, K. Bert: Pre-training of deep bidirectional transformers for language understanding. arXiv preprint 1810.04805 (2018).

3. Khrulkov, V., Mirvakhabova, L., Ustinova, E., Oseledets, I. & Lempitsky, V. Hyperbolic image embeddings in Proceedings of the IEEE/CVF Conference on Computer Vision and Pattern Recognition (2020), 6418–6428.

4. Dosovitskiy, A. et al. An image is worth 16×16 words: Transformers for image recognition at scale. arXiv preprint 2010.11929 (2020).

5. Zhang, F., Yuan, N. J., Lian, D., Xie, X. & Ma, W.-Y. Collaborative knowledge base embedding for recommender systems in Proceedings of the 22nd ACM SIGKDD international conference on knowledge discovery and data mining (2016), 353–362.

6. Stanojevic, S., Li, Y., Ristivojevic, A. & Garmire, L. X. Computational methods for single-cell multi-omics integration and alignment. Genomics, Proteomics & Bioinformatics 20, 836–849 (2022).

7. Ma, A., McDermaid, A., Xu, J., Chang, Y. & Ma, Q. Integrative methods and practical challenges for single-cell multi-omics. Trends in biotechnology 38, 1007–1022 (2020).

8. Theodoris, C. V. et al. Transfer learning enables predictions in network biology. Nature, 1–9 (2023).

9. Du, J. et al. Gene2vec: distributed representation of genes based on co-expression. BMC genomics 20, 7–15 (2019).

10. Li, L. et al. Joint embedding of biological networks for cross-species functional alignment. Bioinformatics 39, btad529 (2023).

11. Kulmanov, M., Khan, M. A. & Hoehndorf, R. DeepGO: predicting protein functions from sequence and interactions using a deep ontology-aware classifier. Bioinformatics 34, 660–668 (2018).

12. Kulmanov, M. & Hoehndorf, R. DeepGOPlus: improved protein function prediction from sequence. Bioinformatics 36, 422–429 (2020).

13. Gligorijević, V. et al. Structure-based protein function prediction using graph convolutional networks. Nature communications 12, 3168 (2021).

14. Xiong, Y. et al. Heterogeneous network embedding enabling accurate disease association predictions. BMC medical genomics 12, 1–17 (2019).

15. Yu, Z., Huang, F., Zhao, X., Xiao, W. & Zhang, W. Predicting drug–disease associations through layer attention graph convolutional network. Briefings in bioinformatics 22, bbaa243 (2021).

16. Wang, S., Flynn, E. R. & Altman, R. B. Gaussian embedding for large-scale gene set analysis. Nature machine intelligence 2, 387–395 (2020).

17. Chen, H.-I. H. et al. GSAE: an autoencoder with embedded gene-set nodes for genomics functional characterization. BMC systems biology 12, 45–57 (2018).

18. Mostavi, M., Chiu, Y.-C., Huang, Y. & Chen, Y. Convolutional neural network models for cancer type prediction based on gene expression. BMC medical genomics 13, 1–13 (2020).

19. Kim, S., Lee, H., Kim, K. & Kang, J. Mut2Vec: distributed representation of cancerous mutations. BMC medical genomics 11, 57–69 (2018).

20. Bryant, P., Pozzati, G. & Elofsson, A. Improved prediction of protein-protein interactions using AlphaFold2. Nature communications 13, 1265 (2022).

21. Gao, K. Y. et al. Interpretable drug target prediction using deep neural representation. in IJCAI 2018 (2018), 3371–3377.

22. Hahne, F. et al. Hypergeometric testing used for gene set enrichment analysis. Bioconductor case studies, 207–220 (2008).

23. Subramanian, A. et al. Gene set enrichment analysis: a knowledge-based approach for interpreting genome-wide expression profiles. Proceedings of the National Academy of Sciences 102, 15545–15550 (2005).

24. Yao, V., Wong, A. K. & Troyanskaya, O. G. Enabling precision medicine through integrative network models. Journal of molecular biology 430, 2913–2923 (2018).

25. Wang, L., Jia, P., Wolfinger, R. D., Chen, X. & Zhao, Z. Gene set analysis of genome-wide association studies: methodological issues and perspectives. Genomics 98, 1–8 (2011).

26. Peyvandipour, A., Saberian, N., Shafi, A., Donato, M. & Draghici, S. A novel computational approach for drug repurposing using systems biology. Bioinformatics 34, 2817–2825 (2018).

27. Reay, W. R. & Cairns, M. J. Advancing the use of genome-wide association studies for drug repurposing. Nature Reviews Genetics 22, 658–671 (2021).

28. Azuaje, F., Wang, H. & Bodenreider, O. Ontology-driven similarity approaches to supporting gene functional assessment in Proceedings of the ISMB’2005 SIG meeting on Bio-ontologies 2005 (2005), 9–10.

29. Wieting, J., Bansal, M., Gimpel, K. & Livescu, K. Towards universal paraphrastic sentence embeddings. arXiv preprint 1511.08198 (2015).

30. Lin, Z. et al. Evolutionary-scale prediction of atomic-level protein structure with a language model. Science 379, 1123–1130 (2023).

31. Greene, C. S. et al. Understanding multicellular function and disease with human tissue-specific networks. Nature genetics 47, 569–576 (2015).

32. Kim, S.-Y. & Volsky, D. J. PAGE: parametric analysis of gene set enrichment. BMC bioinformatics 6, 1–12 (2005).

33. Han, H., Lee, S. & Lee, I. NGSEA: network-based gene set enrichment analysis for interpreting gene expression phenotypes with functional gene sets. Molecules and cells 42, 579 (2019).

34. Cousins, H. et al. Gene set proximity analysis: expanding gene set enrichment analysis through learned geometric embeddings, with drug-repurposing applications in COVID-19. Bioinformatics 39, btac735 (2023).

35. Grover, A. & Leskovec, J. node2vec: Scalable feature learning for networks in Proceedings of the 22nd ACM SIGKDD international conference on Knowledge discovery and data mining (2016), 855–864.

36. Qiu, J. et al. Network embedding as matrix factorization: Unifying deepwalk, line, pte, and node2vec in Proceedings of the eleventh ACM international conference on web search and data mining (2018), 459–467.

37. Tarca, A. L., Bhatti, G. & Romero, R. A comparison of gene set analysis methods in terms of sensitivity, prioritization and specificity. PloS one 8, e79217 (2013).

38. Hamosh, A., Scott, A. F., Amberger, J. S., Bocchini, C. A. & McKusick, V. A. Online Mendelian Inheritance in Man (OMIM), a knowledgebase of human genes and genetic disorders. Nucleic acids research 33, D514–D517 (2005).

39. Wishart, D. S. et al. DrugBank 5.0: a major update to the DrugBank database for 2018. Nucleic acids research 46, D1074–D1082 (2018).

40. Jørgensen, E. A., Knigge, U., Warberg, J. & Kjær, A. Histamine and the regulation of body weight. Neuroendocrinology 86, 210–214 (2007).

41. Perl, A. E. et al. Follow-up of patients with R/R FLT3-mutation–positive AML treated with gilteritinib in the phase 3 ADMIRAL trial. Blood, The Journal of the American Society of Hematology 139, 3366–3375 (2022).

42. Yu, X. et al. Fingolimod ameliorates schizophrenia-like cognitive impairments induced by phencyclidine in male rats. British Journal of Pharmacology 180, 161–173 (2023).

43. Suri, R. et al. Sirolimus loaded chitosan functionalized poly (lactic-co-glycolic acid)(PLGA) nanoparticles for potential treatment of age-related macular degeneration. International journal of biological macromolecules 191, 548–559 (2021).

44. Resnik, P. Semantic similarity in a taxonomy: An information-based measure and its application to problems of ambiguity in natural language. Journal of artificial intelligence research 11, 95–130 (1999).

45. Eppig, J. T. et al. Mouse Genome Informatics (MGI): resources for mining mouse genetic, genomic, and biological data in support of primary and translational research. Systems Genetics: Methods and Protocols, 47–73 (2017).

46. Holley, A. J. & Lugo, J. N. Effects of an acute seizure on associative learning and memory. Epilepsy & Behavior 54, 51–57 (2016).

47. Khaleghzadeh-Ahangar, H., Talebi, A. & Mohseni-Moghaddam, P. Thyroid disorders and development of cognitive impairment: A review study. Neuroendocrinology 112, 835–844 (2022).

48. Dixon, S. J., Costanzo, M., Baryshnikova, A., Andrews, B. & Boone, C. Systematic mapping of genetic interaction networks. Annual review of genetics 43, 601–625 (2009).

49. Obayashi, T., Kodate, S., Hibara, H., Kagaya, Y. & Kinoshita, K. COXPRESdb v8: an animal gene coexpression database navigating from a global view to detailed investigations. Nucleic Acids Research 51, D80–D87 (2023).

50. Hekselman, I. & Yeger-Lotem, E. Mechanisms of tissue and cell-type specificity in heritable traits and diseases. Nature Reviews Genetics 21, 137–150 (2020).

51. Roussarie, J.-P. et al. Selective neuronal vulnerability in Alzheimer’s disease: a network-based analysis. Neuron 107, 821–835 (2020).

52. Ietswaart, R., Gyori, B. M., Bachman, J. A., Sorger, P. K. & Churchman, L. S. GeneWalk identifies relevant gene functions for a biological context using network representation learning. Genome biology 22, 1–35 (2021).

53. Rives, A. et al. Biological structure and function emerge from scaling unsupervised learning to 250 million protein sequences. Proceedings of the National Academy of Sciences 118, e2016239118 (2021).

54. Weissenow, K., Heinzinger, M. & Rost, B. Protein language-model embeddings for fast, accurate, and alignment-free protein structure prediction. Structure 30, 1169–1177 (2022).

55. Villegas-Morcillo, A., Gomez, A. M. & Sanchez, V. An analysis of protein language model embeddings for fold prediction. Briefings in Bioinformatics 23, bbac142 (2022).

56. Liu, J., Huang, Y., Singh, R.Vert, J.-P. & Noble, W. S. Jointly embedding multiple single-cell omics measurements in Algorithms in bioinformatics:… International Workshop, WABI…, proceedings. WABI (Workshop) 143 (2019).

57. Zhao, Y., Cai, H., Zhang, Z., Tang, J. & Li, Y. Learning interpretable cellular and gene signature embeddings from single-cell transcriptomic data. Nature communications 12, 5261 (2021).

58. Chen, H., Ryu, J., Vinyard, M. E., Lerer, A. & Pinello, L. SIMBA: single-cell embedding along with features. Nature Methods 21, 1003–1013 (2024).

59. Dannenfelser, R. & Yao, V. Splitpea: quantifying protein interaction network rewiring changes due to alternative splicing in cancer. bioRxiv, 2023–09 (2023).

60. Liu, R. & Krishnan, A. PecanPy: a fast, efficient and parallelized Python implementation of node2vec. Bioinformatics 37, 3377–3379 (2021).

61. Liu, R., Hirn, M. & Krishnan, A. Accurately modeling biased random walks on weighted networks using node2vec+. Bioinformatics 39, btad047 (2023).

62. Consortium, G. O. The Gene Ontology (GO) database and informatics resource. Nucleic acids research 32, D258–D261 (2004).

63. Kanehisa, M. & Goto, S. KEGG: kyoto encyclopedia of genes and genomes. Nucleic acids research 28, 27–30 (2000).

64. Kamburov, A., Stelzl, U., Lehrach, H. & Herwig, R. The ConsensusPathDB interaction database: 2013 update. Nucleic acids research 41, D793–D800 (2013).

65. Schriml, L. M. et al. Disease Ontology: a backbone for disease semantic integration. Nucleic acids research 40, D940–D946 (2012).

66. Vandin, F., Clay, P., Upfal, E. & Raphael, B. J. in Biocomputing 2012 55–66 (World Scientific, 2012).

67. Tarca, A. L., Draghici, S., Bhatti, G. & Romero, R. Down-weighting overlapping genes improves gene set analysis. BMC bioinformatics 13, 1–14 (2012).

68. Glaab, E., Baudot, A., Krasnogor, N., Schneider, R. & Valencia, A. EnrichNet: network-based gene set enrichment analysis. Bioinformatics 28, i451–i457 (2012).

