## Supplemental Figures for "ANDES: a novel best-match approach for enhancing gene set analysis in embedding spaces"

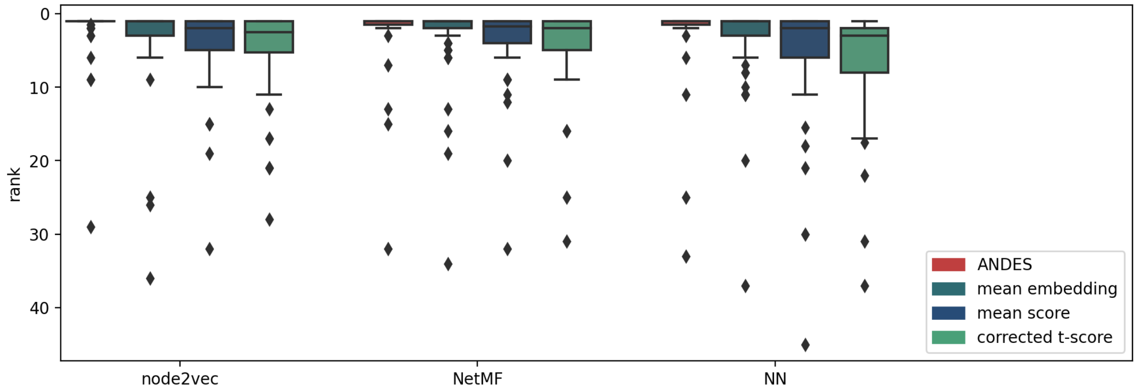


**Figure S1. Performance comparison of KEGG-GO gene set matching when including overlapping genes.** Boxplots of the ranking of the correct matching GO term for 50 KEGG terms when the original gene sets are used directly demonstrate that ANDES outperforms the mean embedding method (node2vec: p=0.022, NetMF: p=5.44 $\times 10^{-3}$, NN: p=5.23 $\times 10^{-3}$, Wilcoxon signed-rank test), corrected t-score (node2vec: p=3.84 $\times 10^{-6}$, NetMF: p=2.04 $\times 10^{-5}$, NN: p=4.45 $\times 10^{-7}$, Wilcoxon signed-rank test), and mean score (node2vec: p=2.54 $\times 10^{-6}$, NetMF: p=4.05 $\times 10^{-5}$, NN: p=4.37 $\times 10^{-5}$, Wilcoxon signed-rank test) methods across three network embedding approaches.

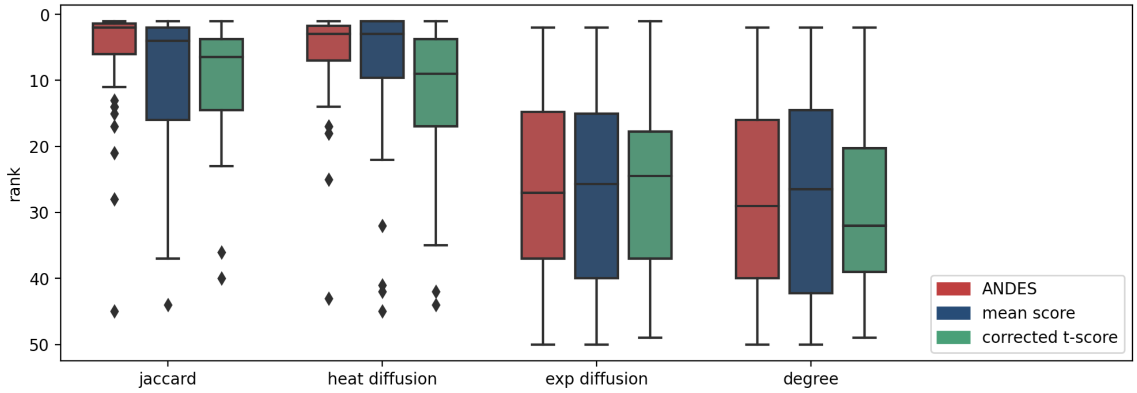


**Figure S2. Extending baseline comparisons to include diffusion methods on the original PPI network.** Matched KEGG-GO terms are ranked using pairwise similarity based on transformations and/or calculations based on the original PPI network, including gene neighbor Jaccard similarity (Jaccard), heat diffusion (heat diffusion), exponential diffusion (exp diffusion) or more naively, by the sum of node degrees (degree). Boxplots show performance comparisons for ANDES, mean score, and corrected t-score. Because pairwise similarity scores are directly calculated from network properties, the mean embedding method could not be included in the comparison.

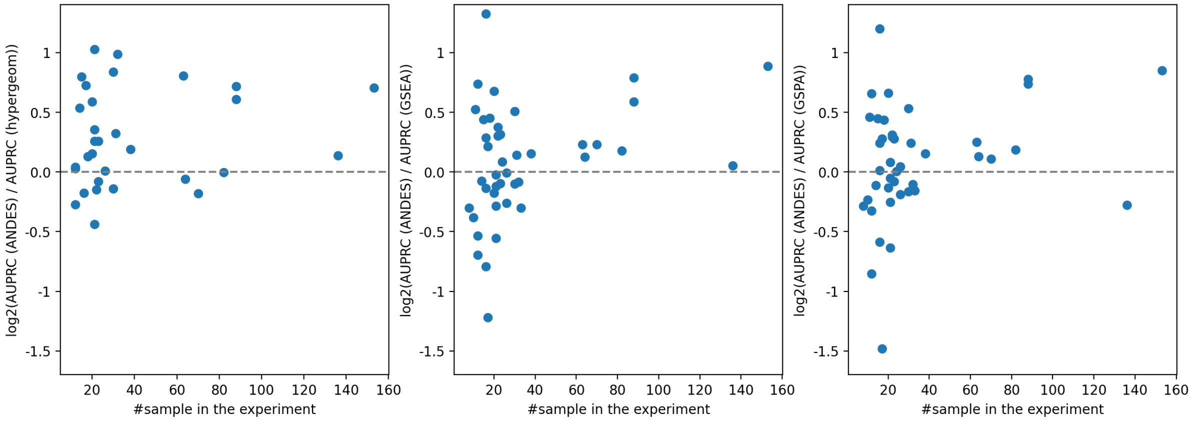


**Figure S3. ANDES has better performance in enrichment analyses when expression datasets have more samples.** Scatterplots show the number of samples in the original GEO2KEGG dataset (x-axis) versus the improvement of AUPRC for ANDES over existing methods (hypergeometric, GSEA, GSPA) as the log2 fold change of AUPRC (y-axis). ANDES has stably better performance for experiment with more samples.

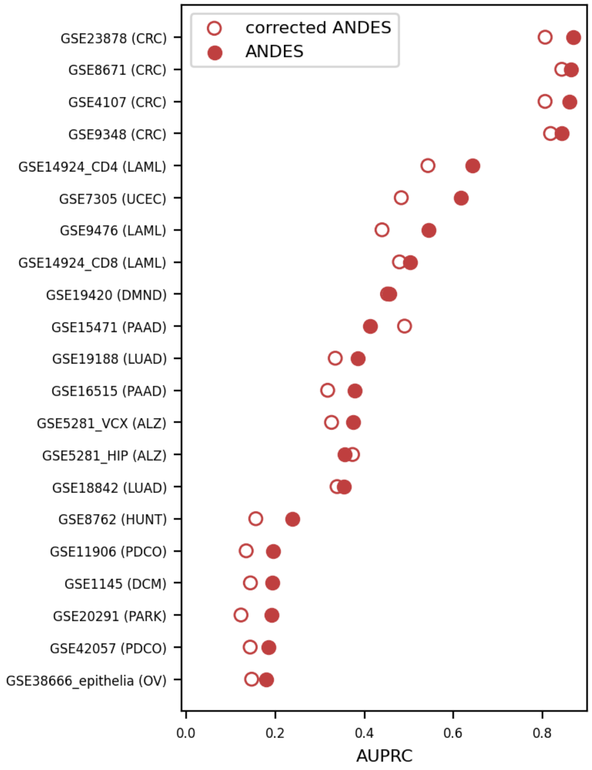


**Figure S4. Gene set enrichment analysis using empirically estimated p-values.** Empirical p-values for KEGG term enrichment scores are calculated based on a null distribution generated from 100 randomly permuted labels for expression datasets. Performance comparison in retrieving annotated KEGG terms using the GEO2KEGG benchmark between original ANDES and ANDES with empirically estimated p-values (corrected-ANDES) to correct for potential biases in the expression data. There is a slight performance drop using corrected-ANDES.

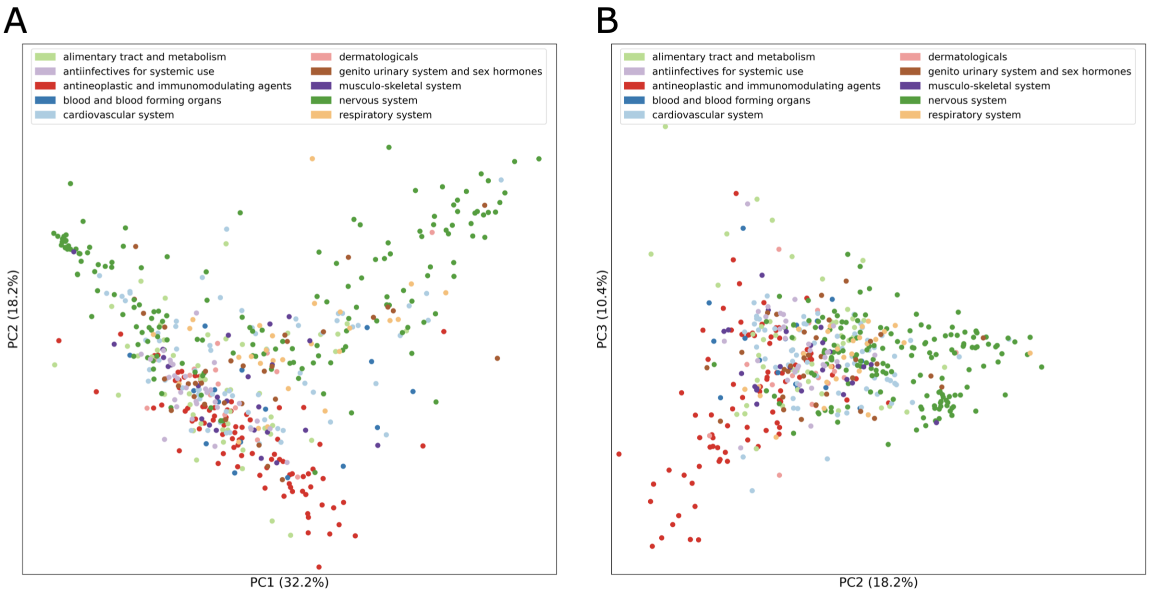


**Figure S5. PCA plot showing the relationship between drugs based on their association with diseases. (A)** PC1 (32.2% of the variance) vs PC2 (18.2% of the variance). **(B)** PC2 (18.2% of the variance) vs PC3 (10.4% of the variance).

Figures S6-S17 show heatmaps of ANDES similarity scores for diseases and drugs in different therapeutic classes based on Anatomical Therapeutic Chemical (ATC) codes. Each figure shows drugs in one ATC 1st level code. The intensity of color indicates the extent to which drug-disease associations exceed $z$-score=1.64 (corresponding to p-value=0.05).

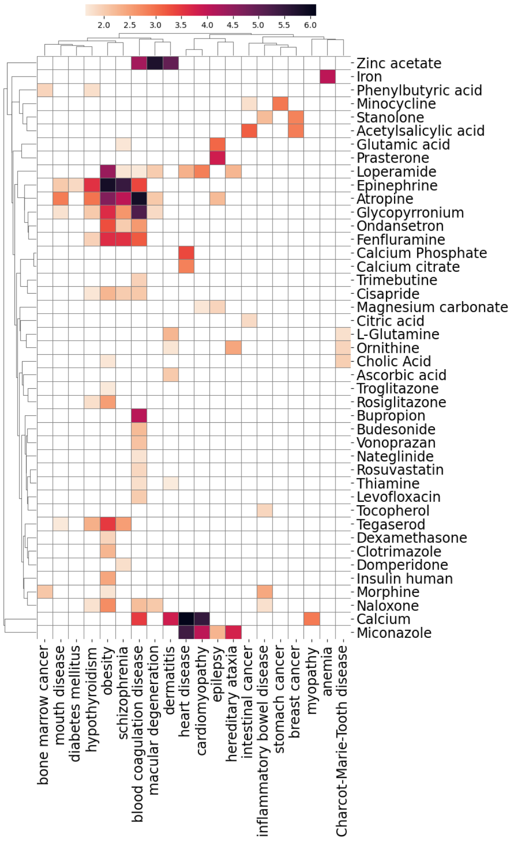


**Figure S6. A: Alimentary tract and metabilism**

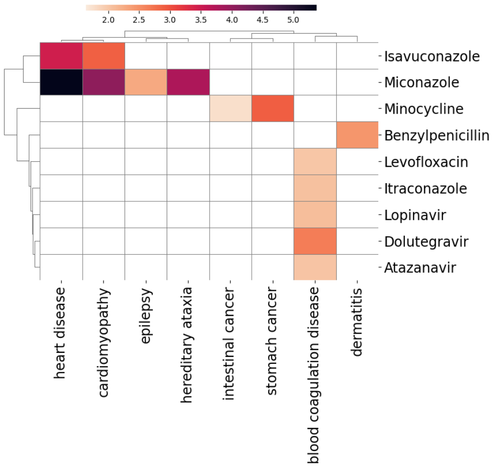


**Figure S7. J: Antiinfective for systemic use**

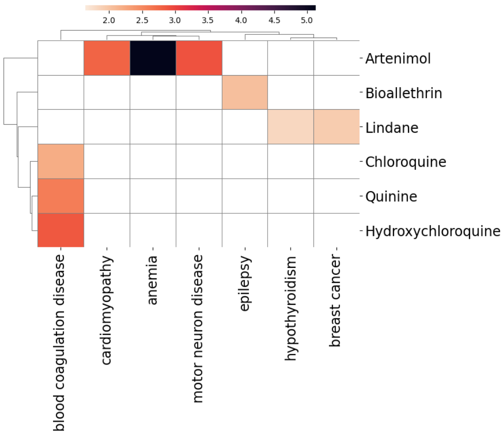


**Figure S8. P: Antiparasitic prducts, insecticides and repellents**

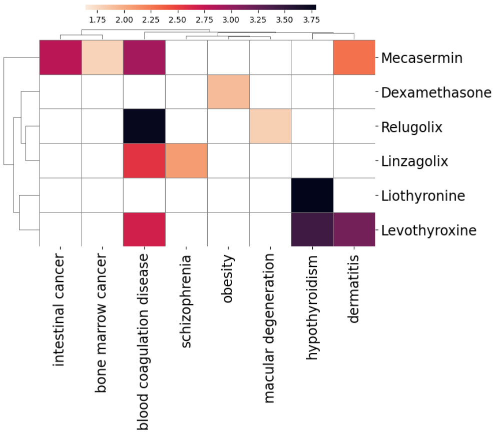


**Figure S9. H: Systemic hormonal preparations, excluding sex hormones and insulins**

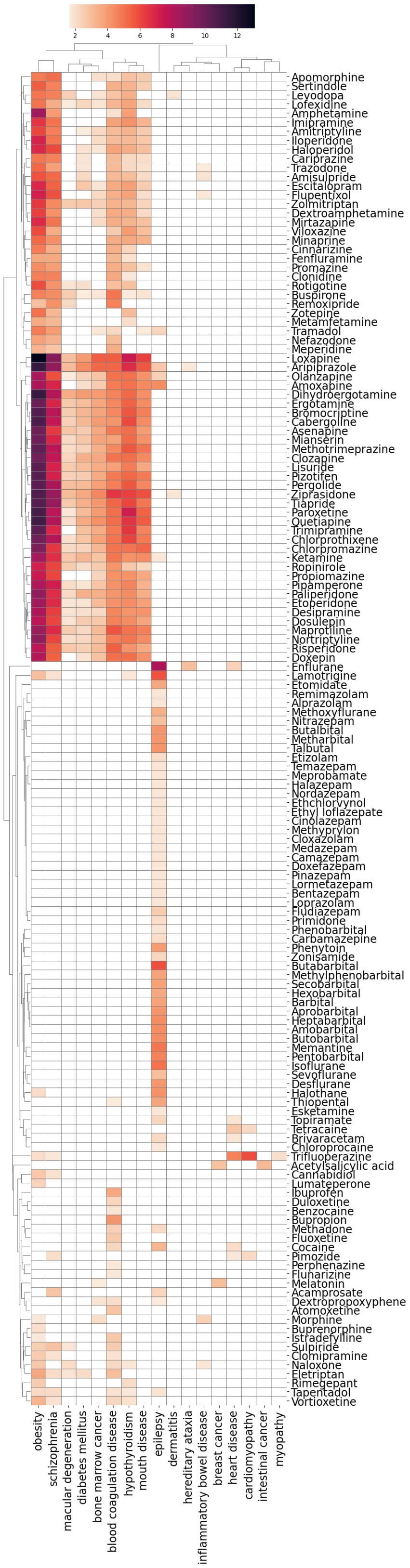

**Figure S10. N: Nervous system**

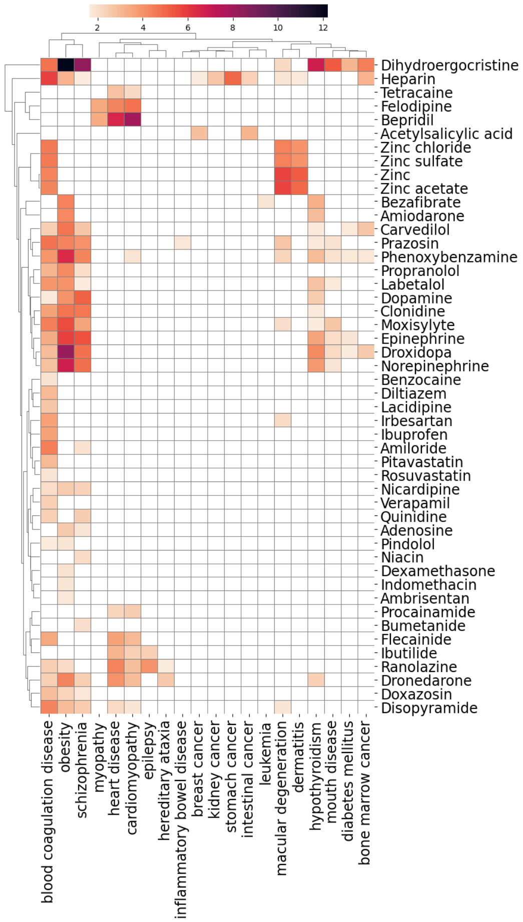


**Figure S11. C: Cardiovascular system**

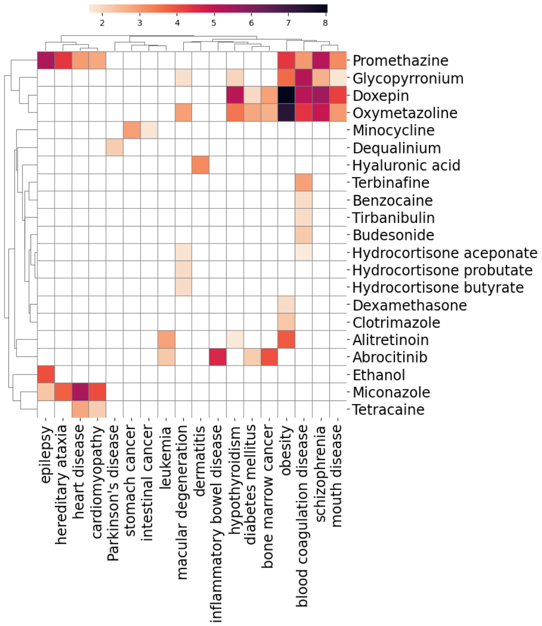


**Figure S12. D: Dermatologicals**

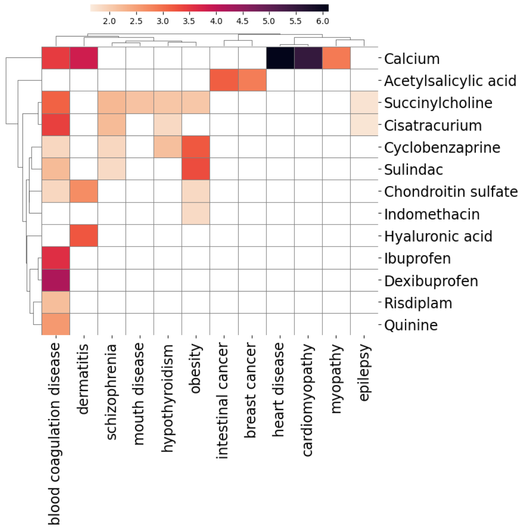


**Figure S13. M: Musculo-skeletal system**

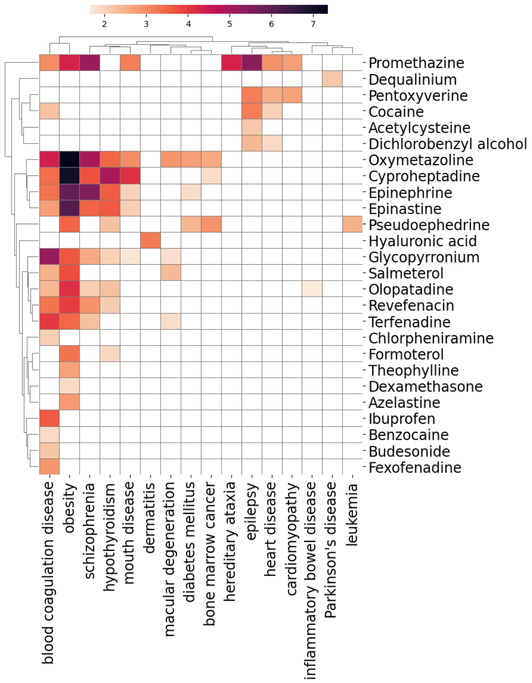


**Figure S14. R: Respiratory system**

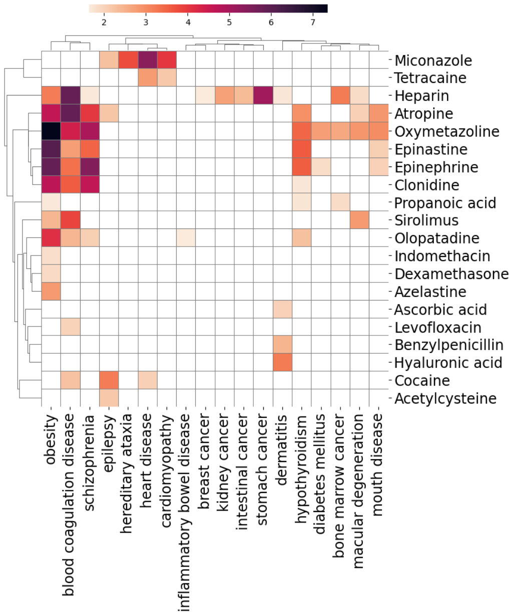


**Figure S15. S: Sensory organs**

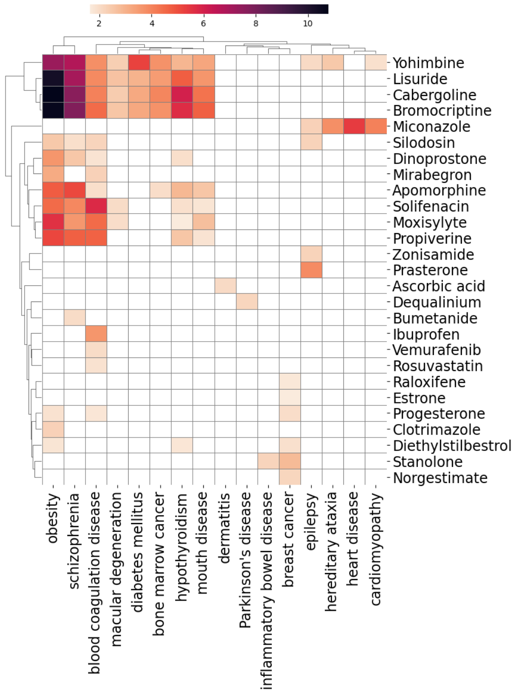


**Figure S16. G: Genito urinary system and sex hormones**

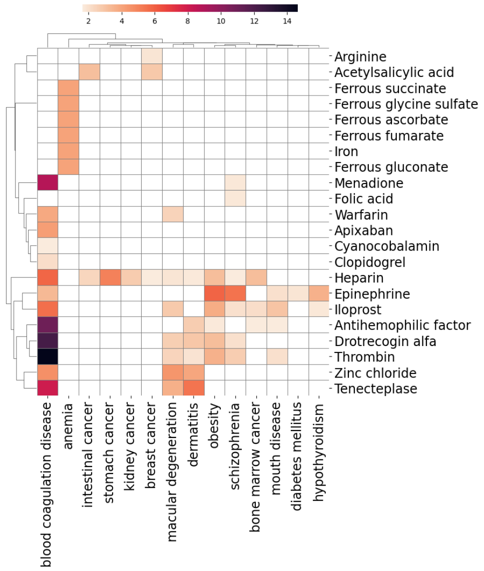


**Figure S17. B: Blood and blood forming organs**

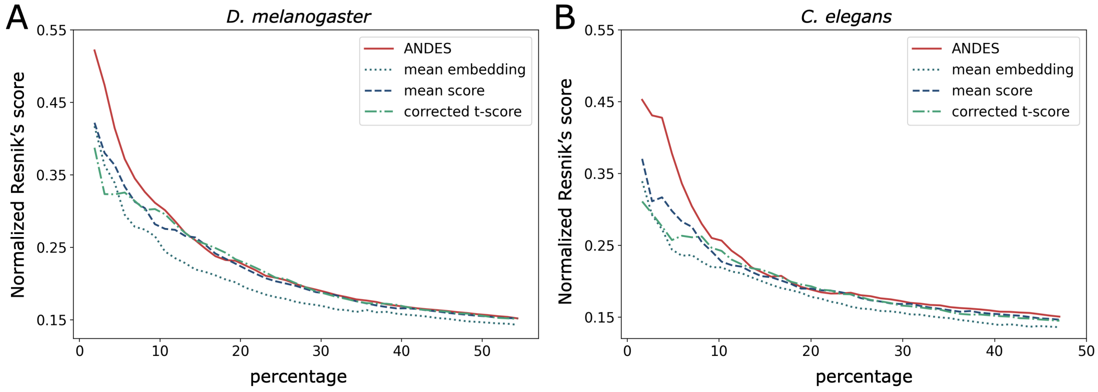


**Figure S18. Cumulative average of the Resnik's score walking down the ranked list. (A)** *H. sapiens* and *D. melanogaster*. **(B)** *H. sapiens* and *C. elegans*.

Figures S19-S41 show the heatmaps of ANDES similarity scores for human disease and mouse phenotype gene sets. Each figure shows one human disease slim term and all of its child terms in the Disease Ontology. For each human disease, the top 10 mouse phenotypes predicted to be functionally similar are shown. The intensity of color indicates the extent to which disease-phenotype associations exceed $z$-score=1.64 (corresponding to p-value=0.05).

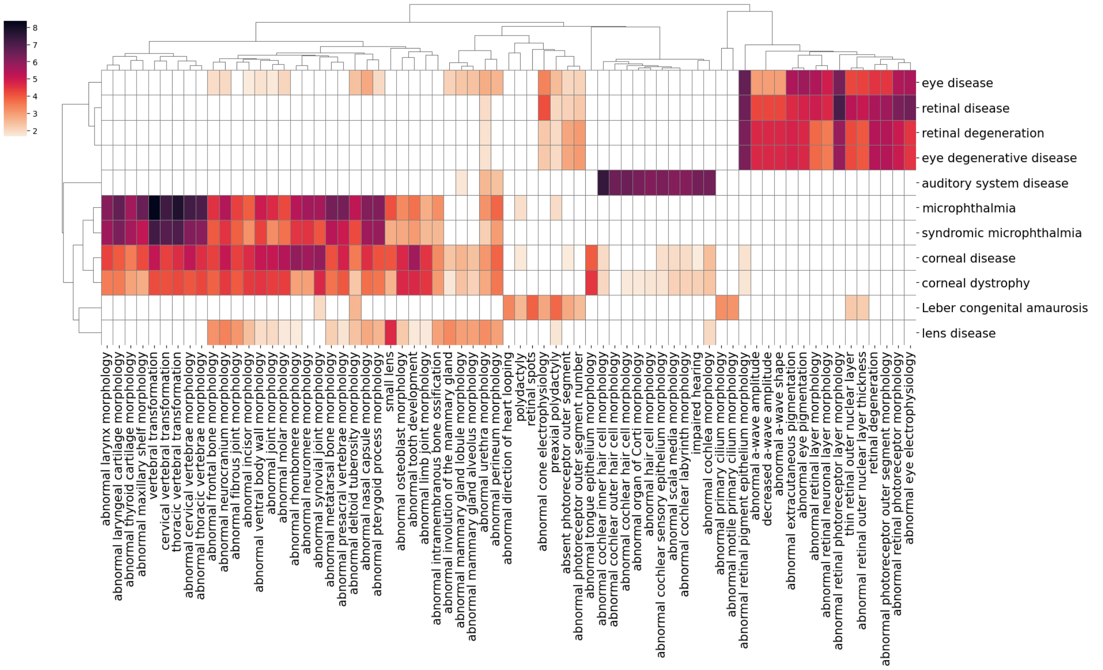


**Figure S19. DOID:0050155 (sensory system disease)**

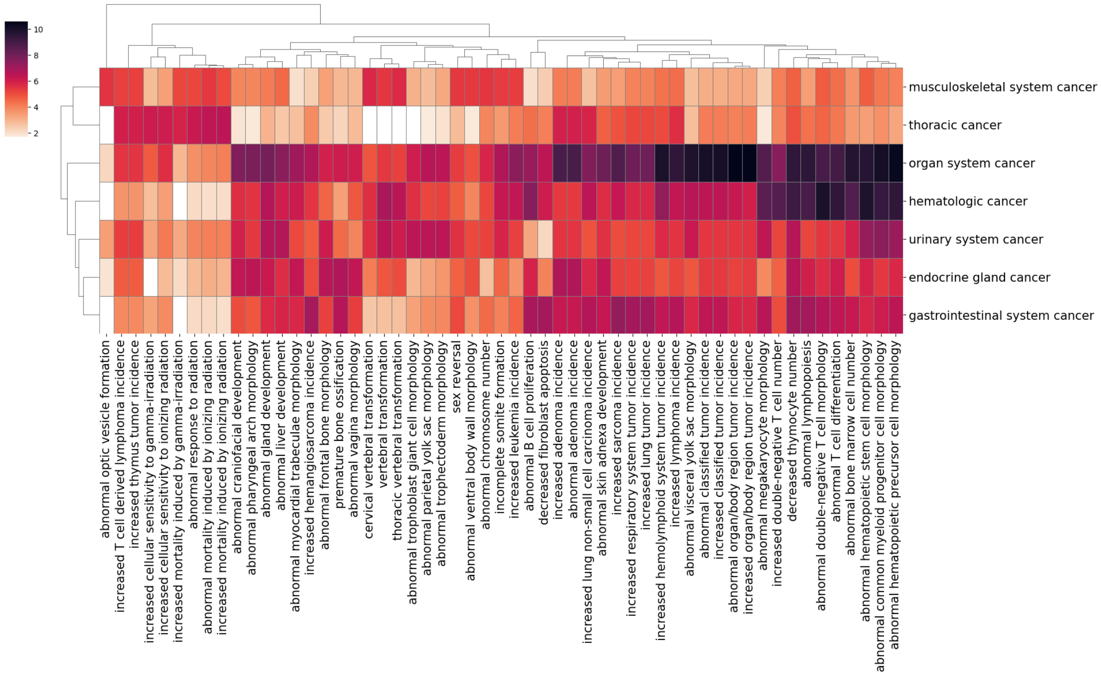


**Figure S20. DOID:0050686 (organ system cancer)**

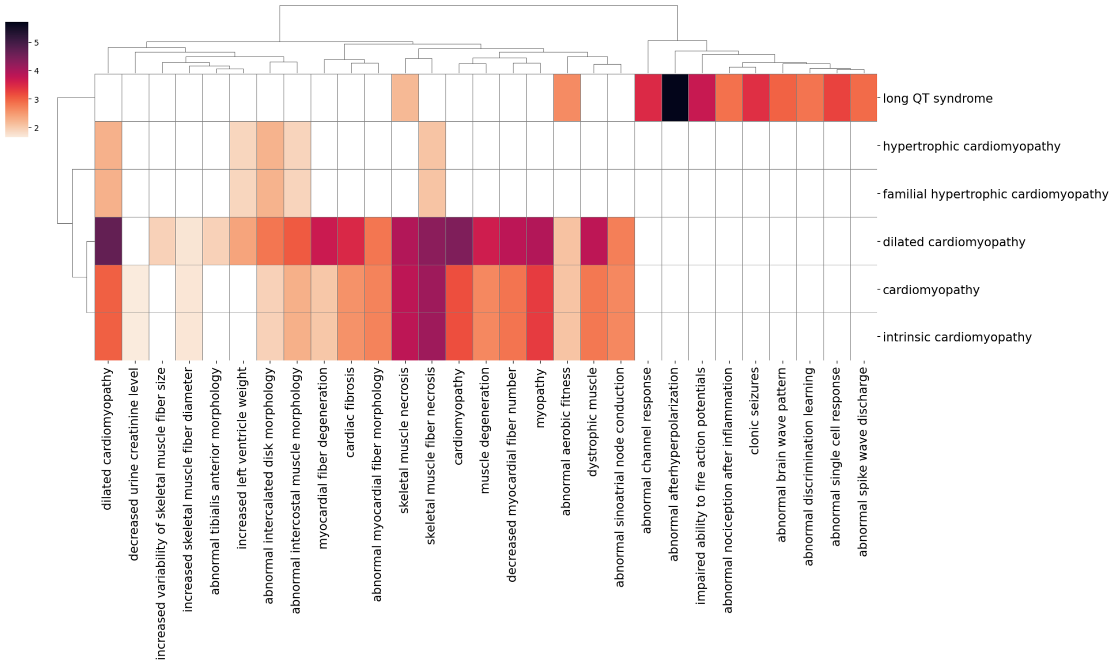


**Figure S21. DOID:0050700 (cardiomyopathy)**

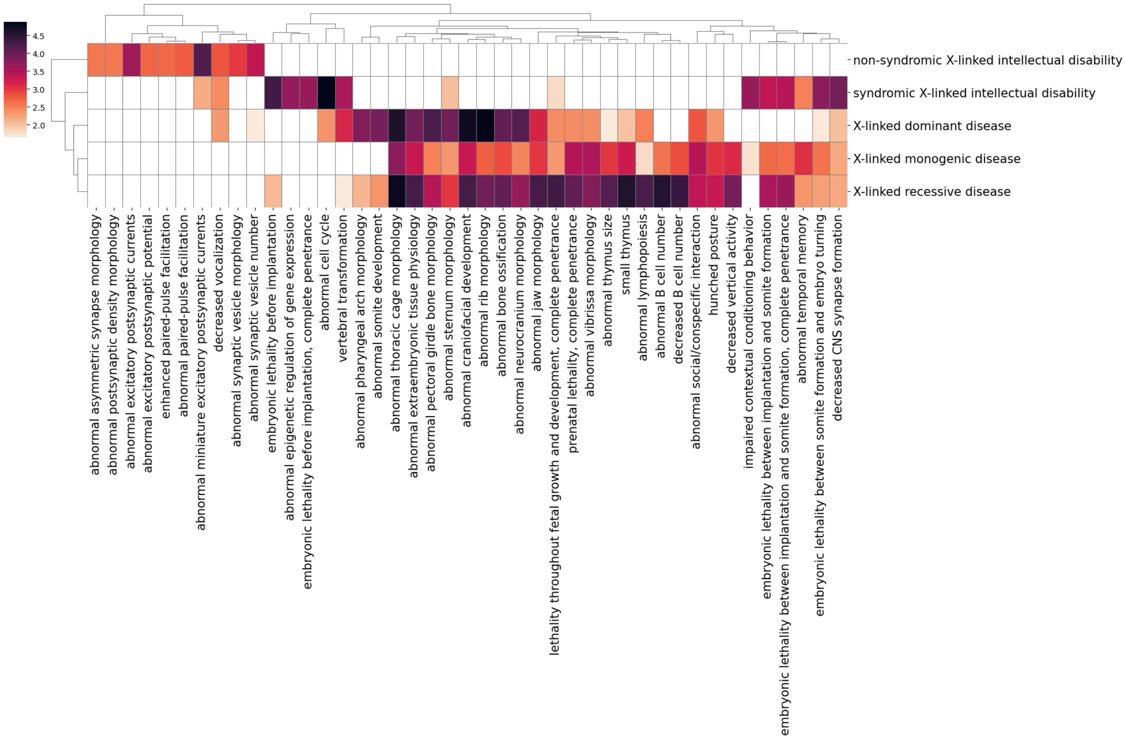


**Figure S22. DOID:0050735 (X-linked monogenic disease)**

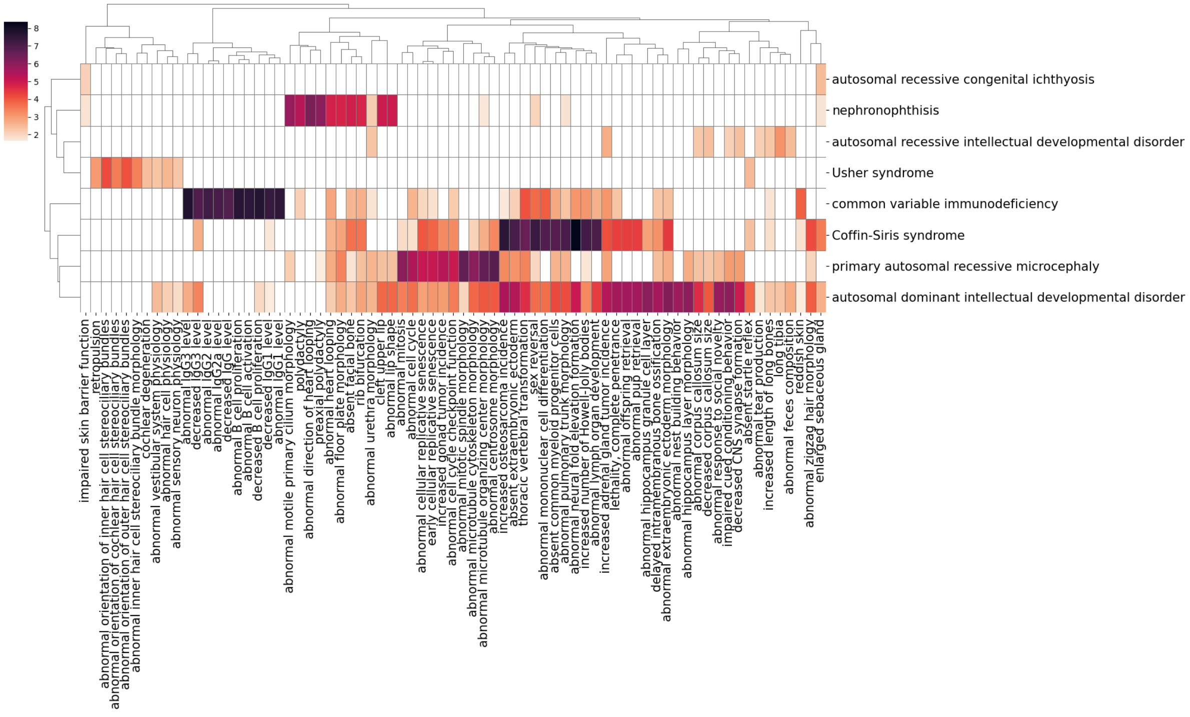


**Figure S23. DOID:0050739 (autosomal genetic disease)**

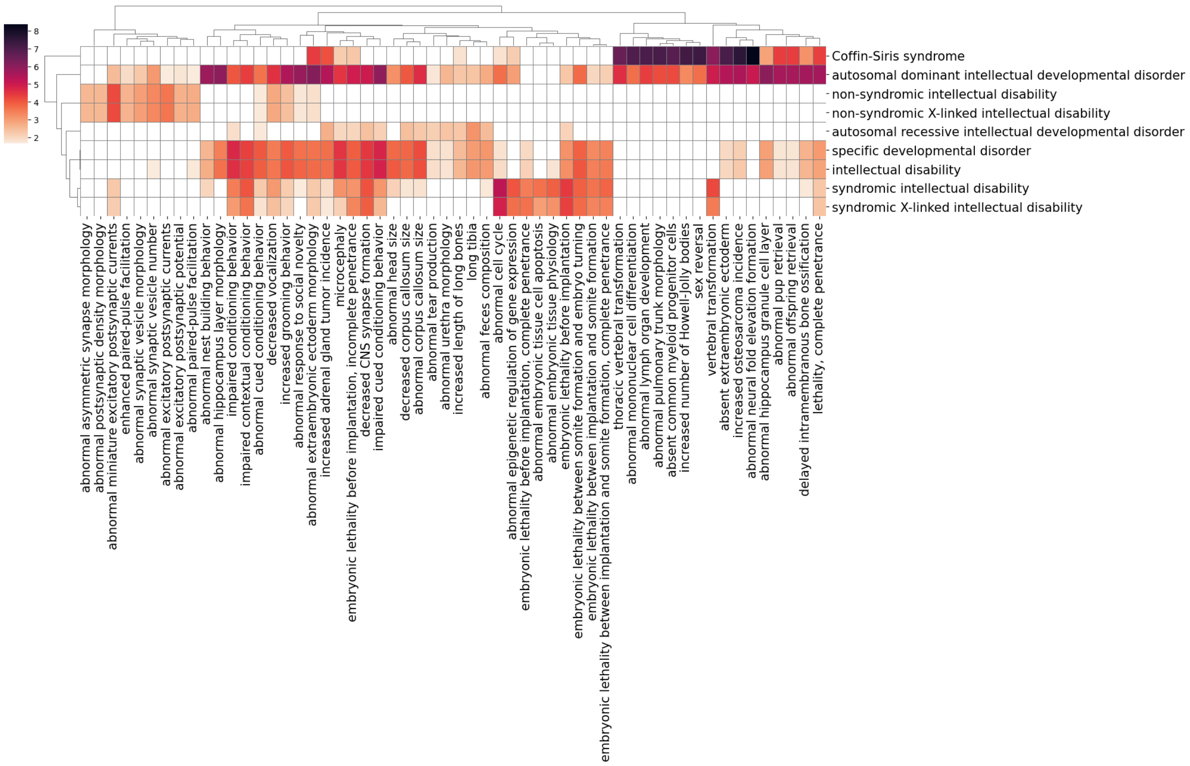


**Figure S24. DOID:0060038 (specific developmental disorder)**

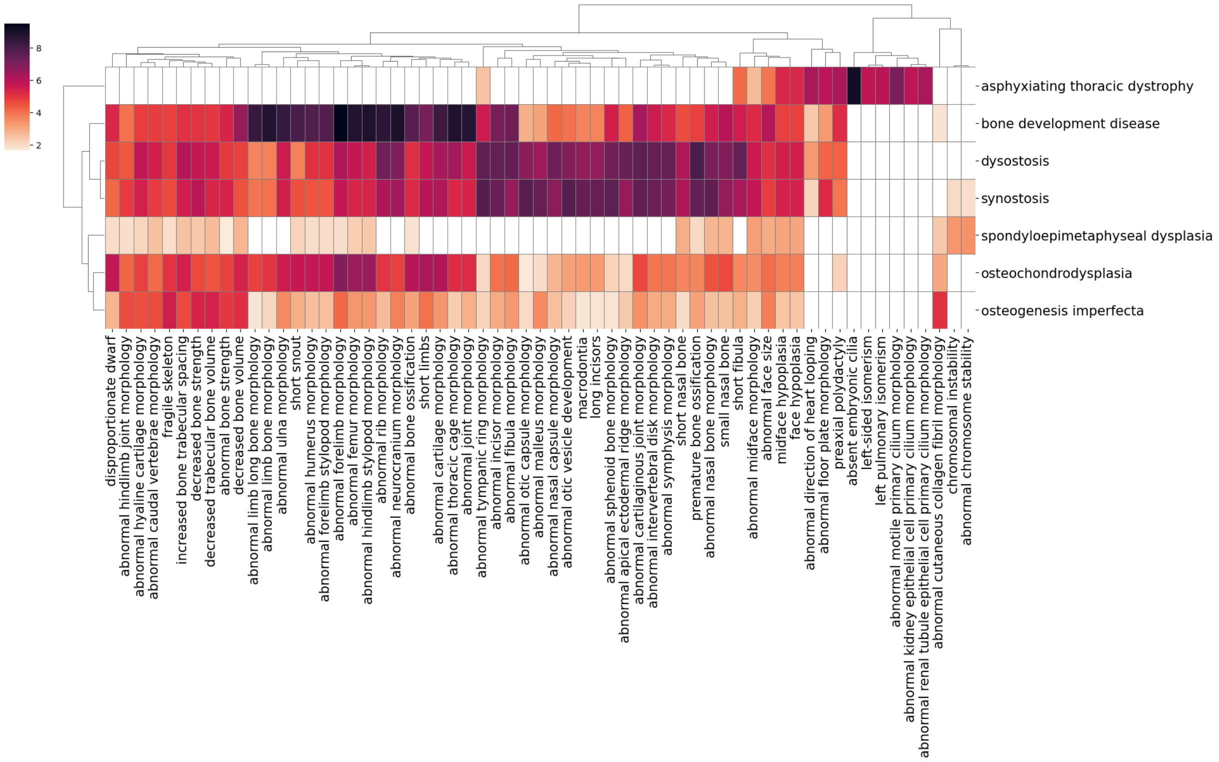


**Figure S25. DOID:0080006 (bone development disease)**

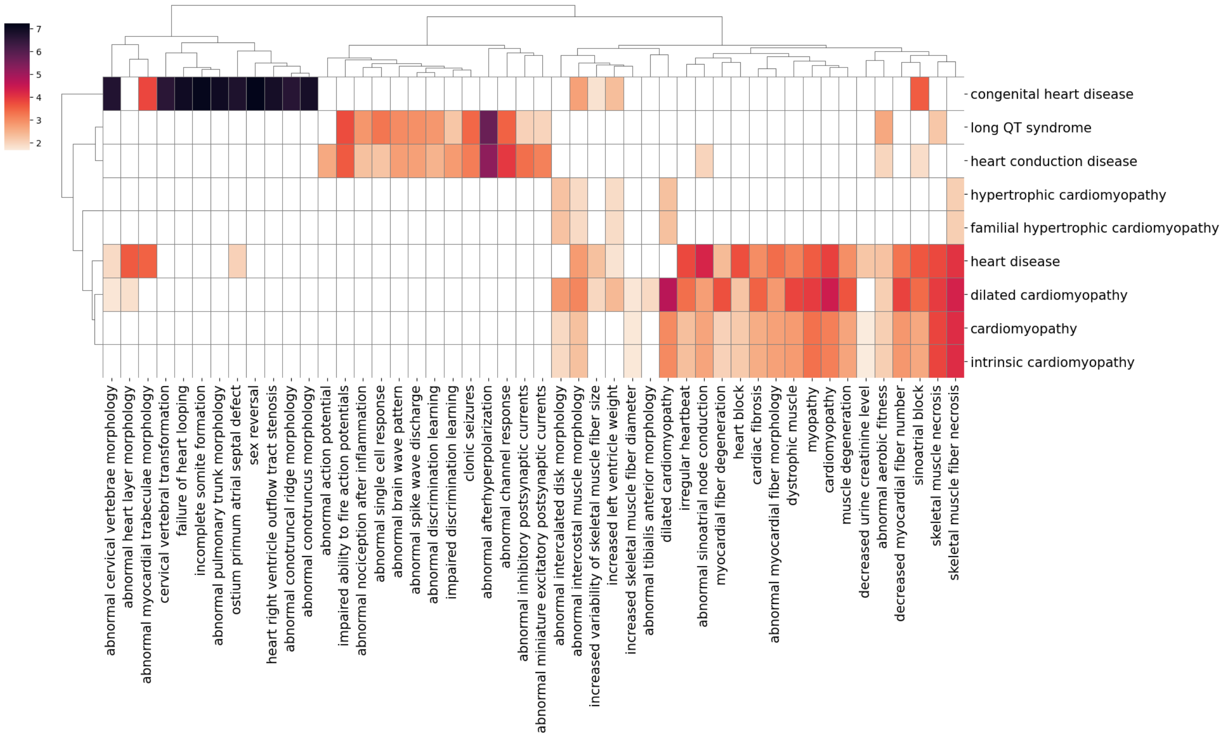


**Figure S26. DOID:114 (heart disease)**

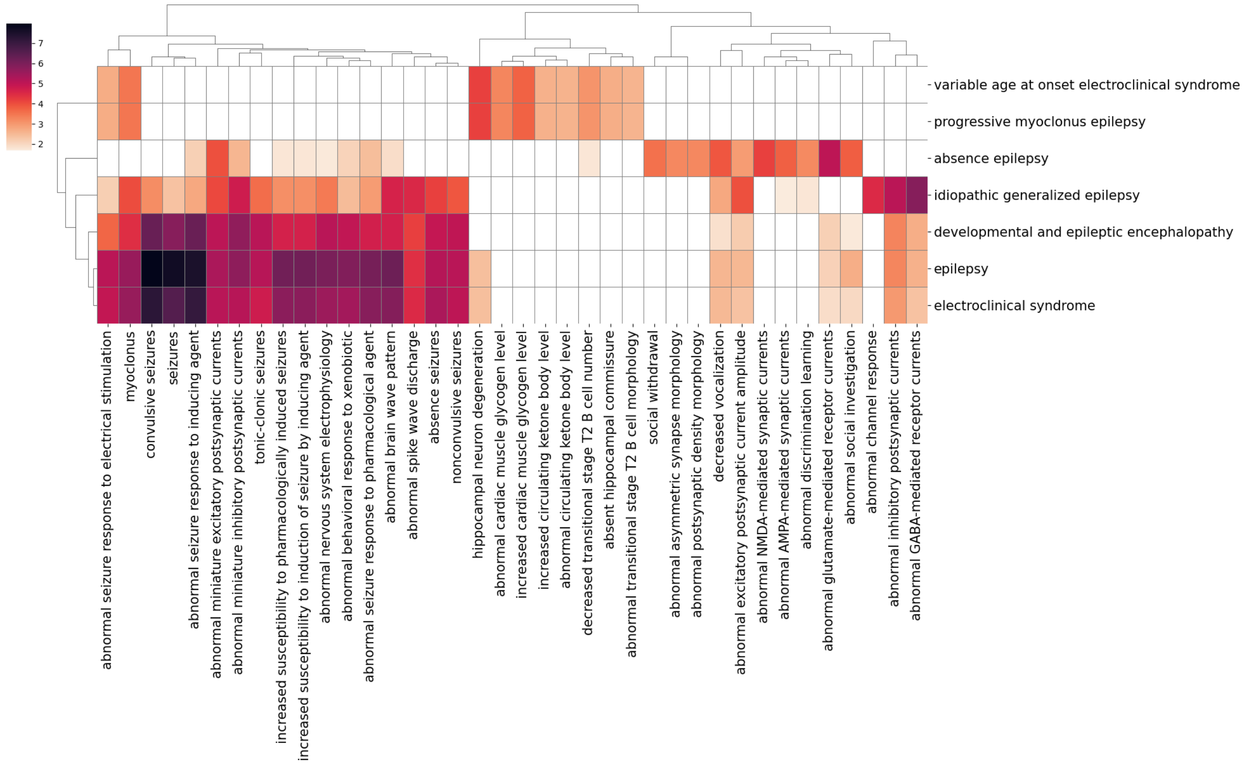


**Figure S27. DOID:1826 (epilepsy)**

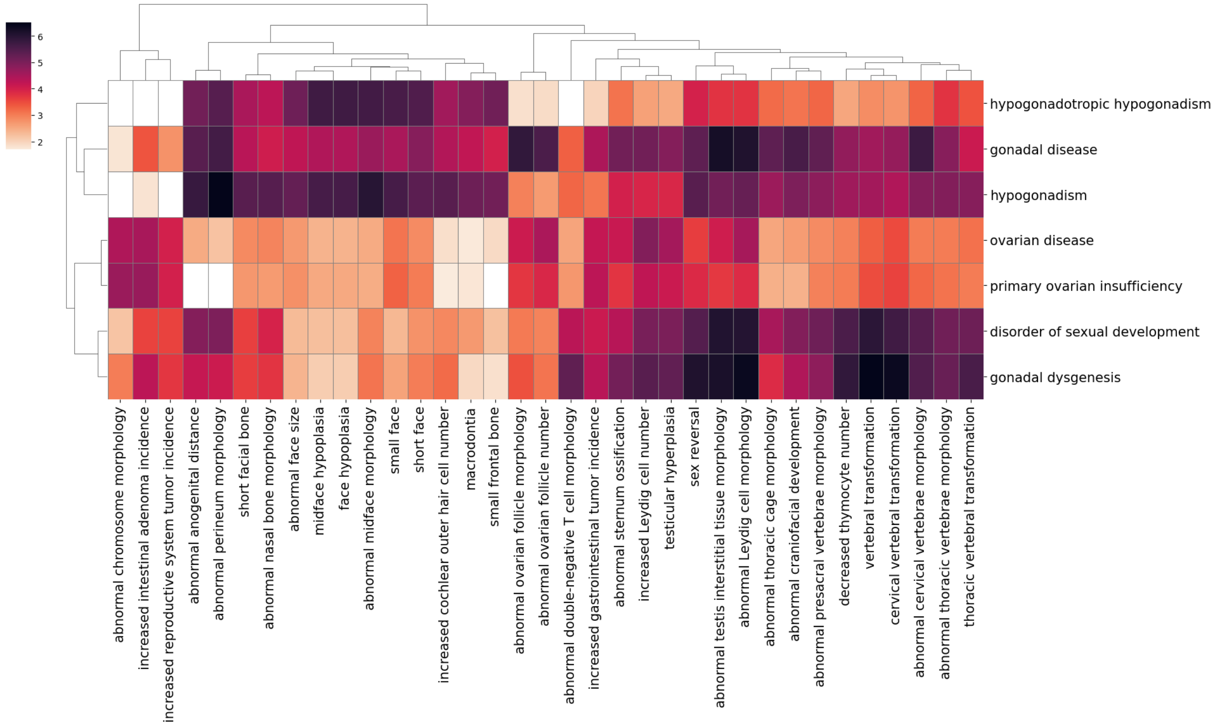


**Figure S28. DOID:2277 (gonadal disease)**

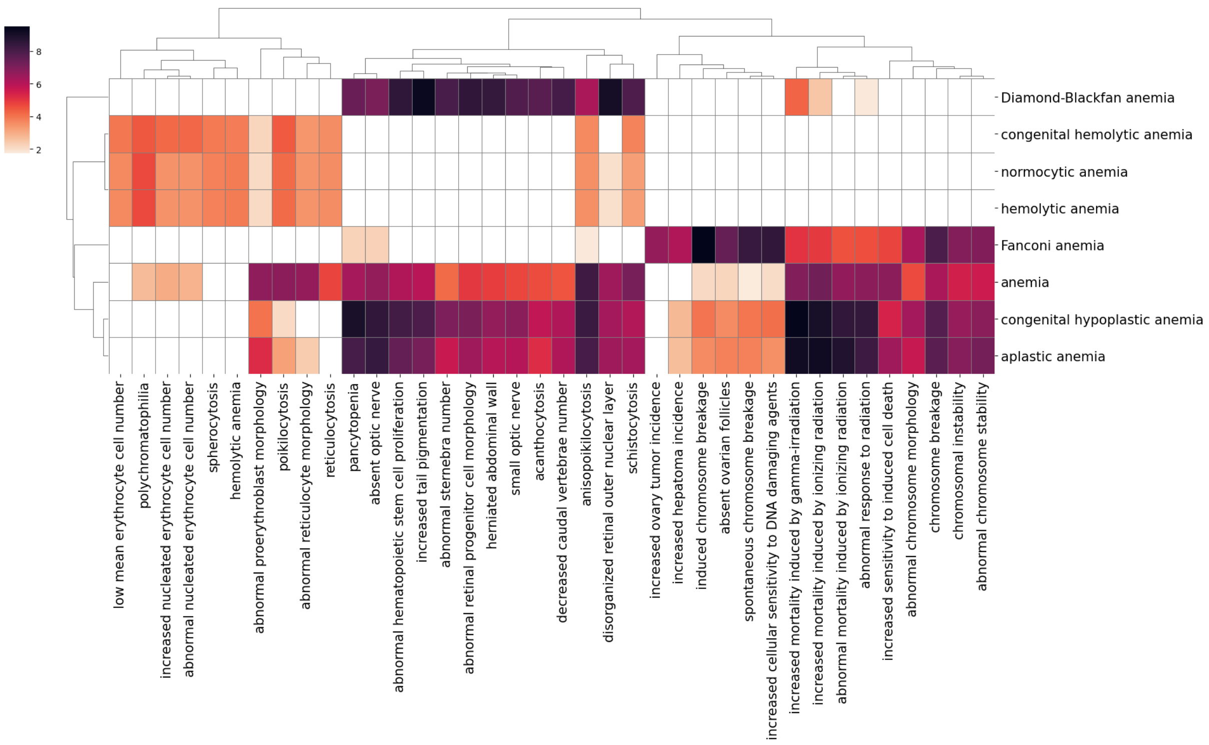


**Figure S29. DOID:2355 (anemia)**

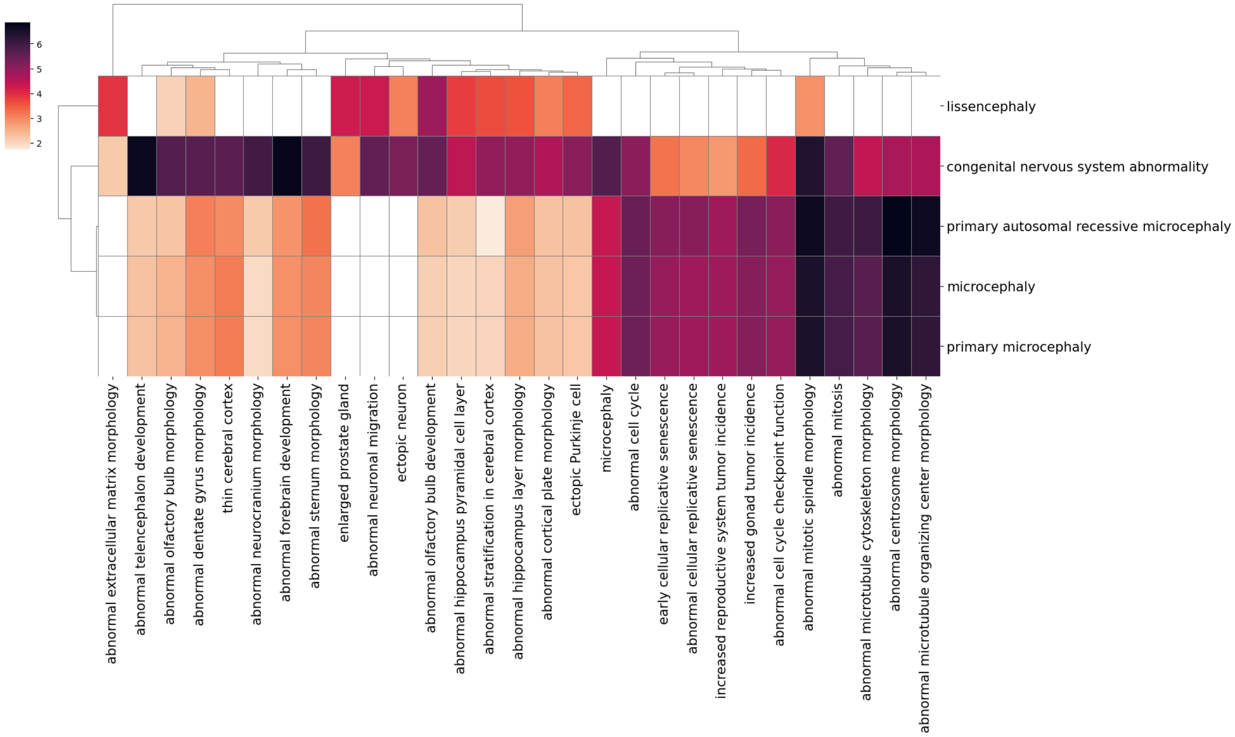


**Figure S30. DOID:2490 (congenital nervous system abnormality)**

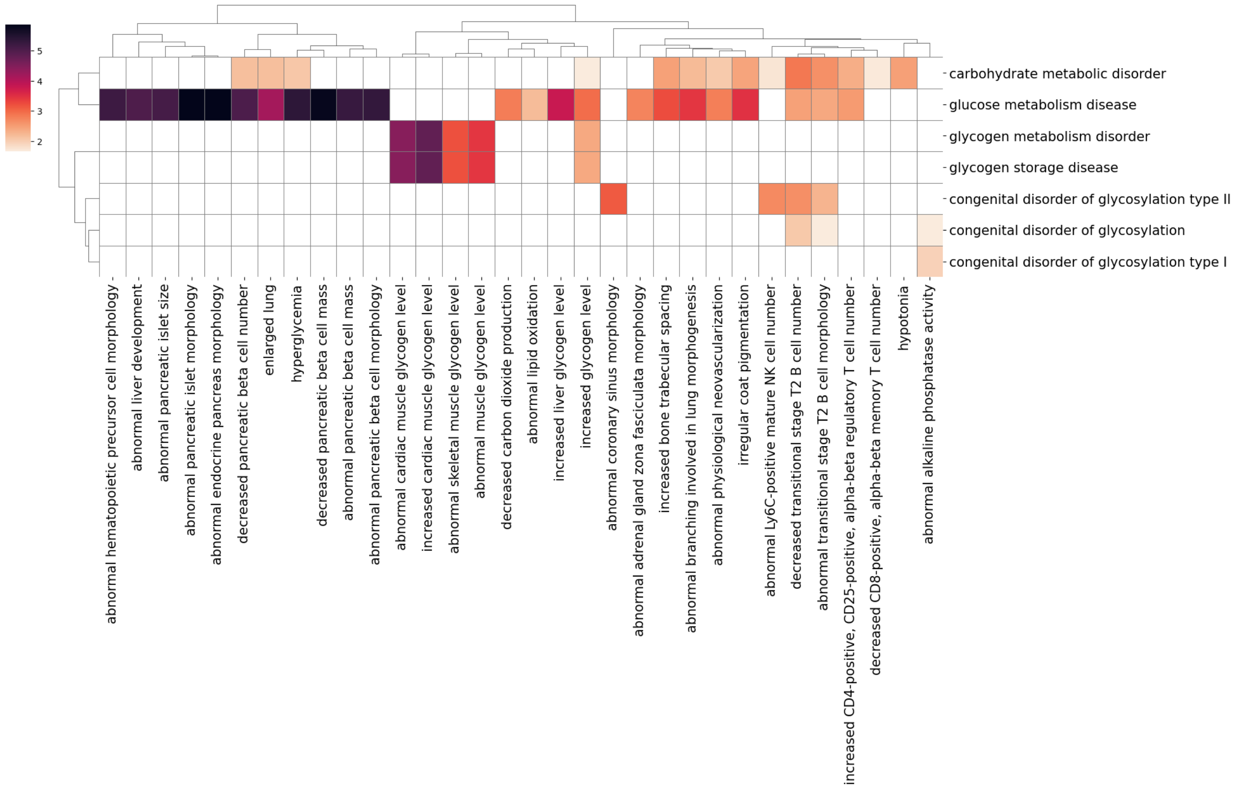


**Figure S31. DOID:2978 (carbohydrate metabolic disorder)**

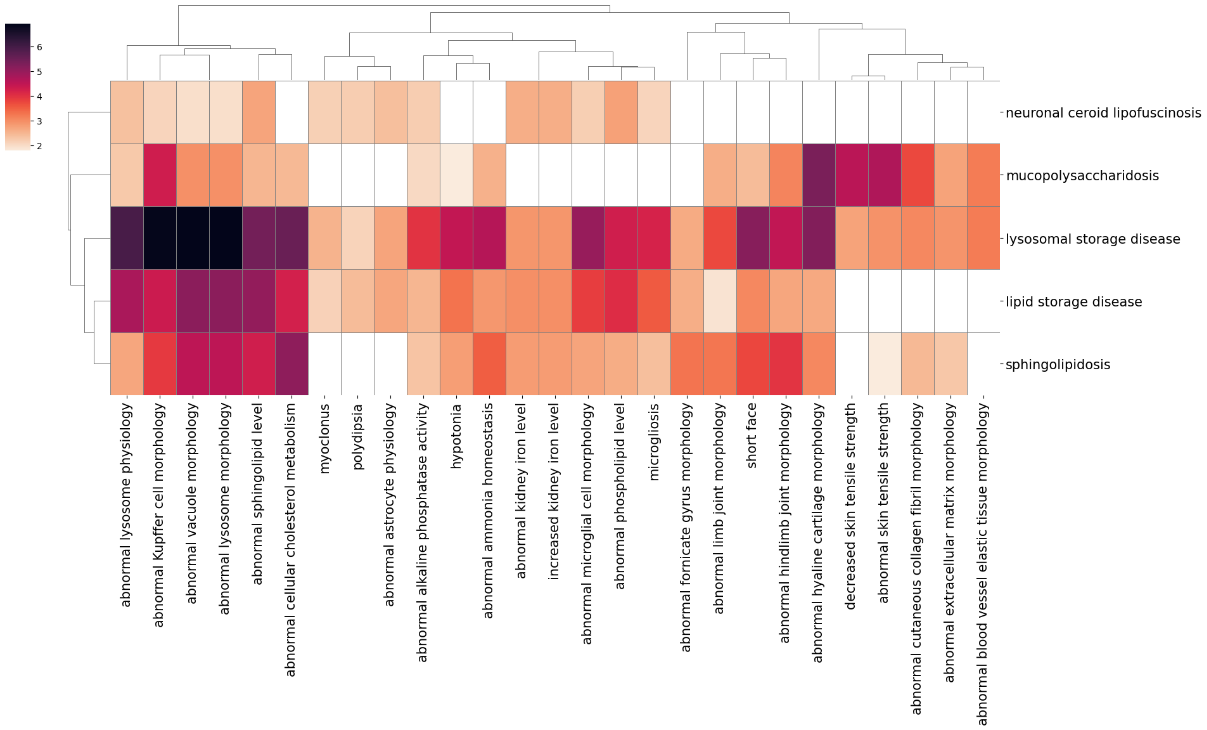


**Figure S32. DOID:3211 (lysosomal storage disease)**

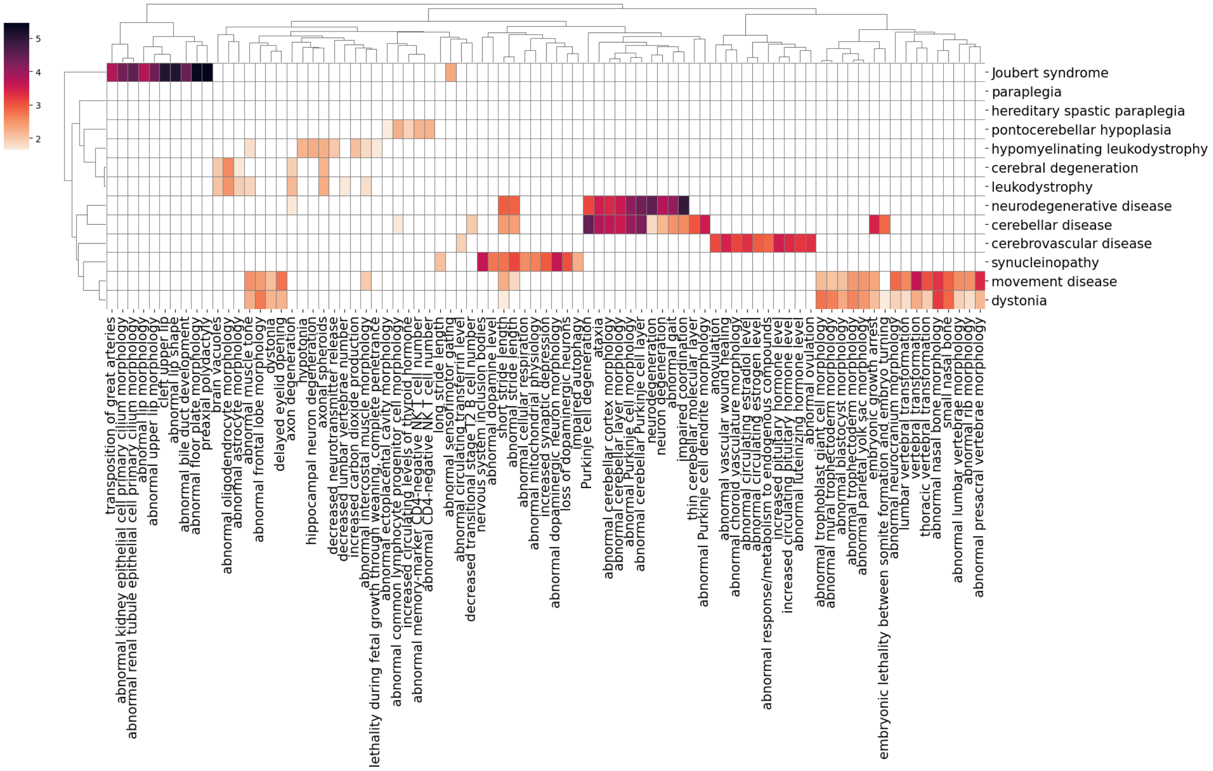


**Figure S33. DOID:331 (central nervous system disease)**

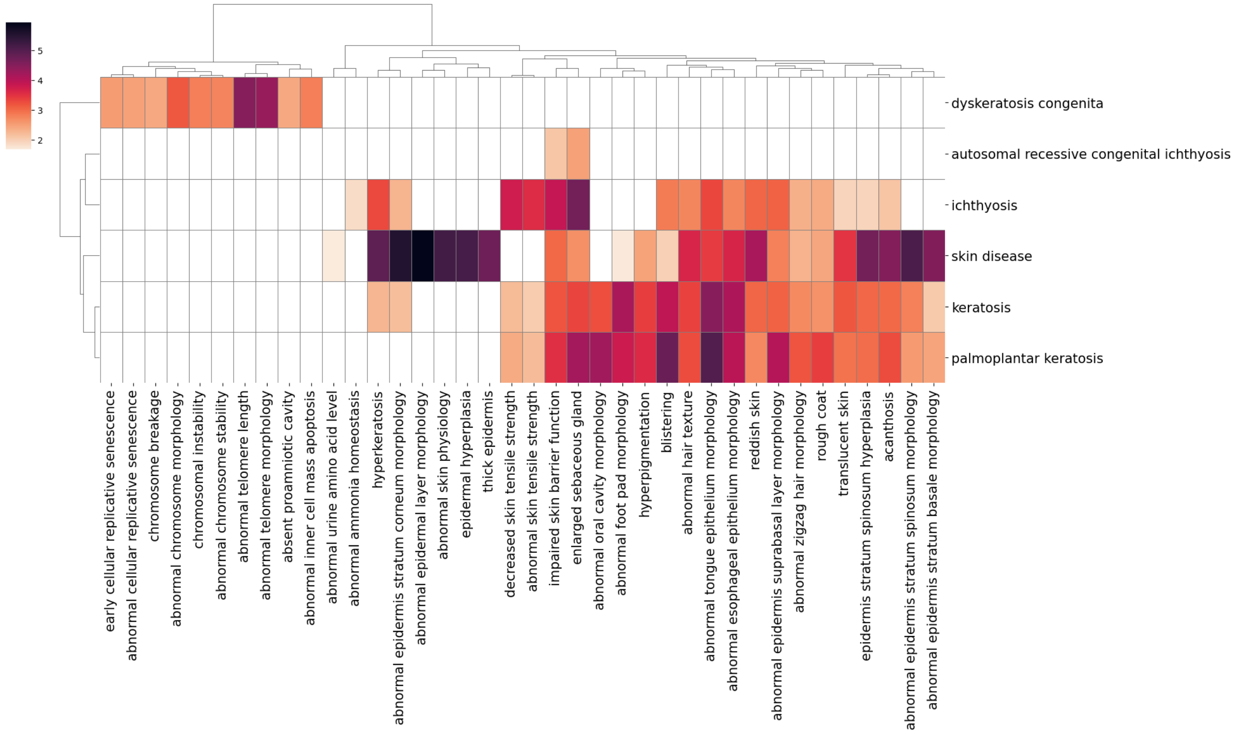


**Figure S34. DOID:37 (skin disease)**

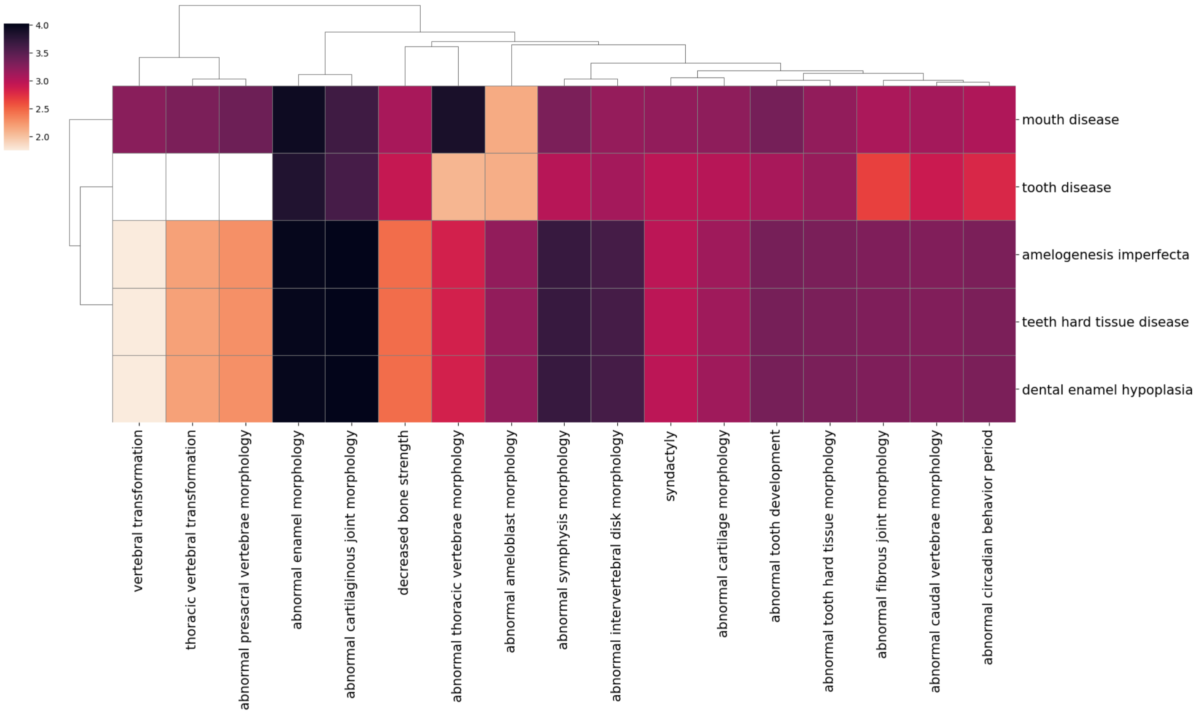


**Figure S35. DOID:403 (mouth disease)**

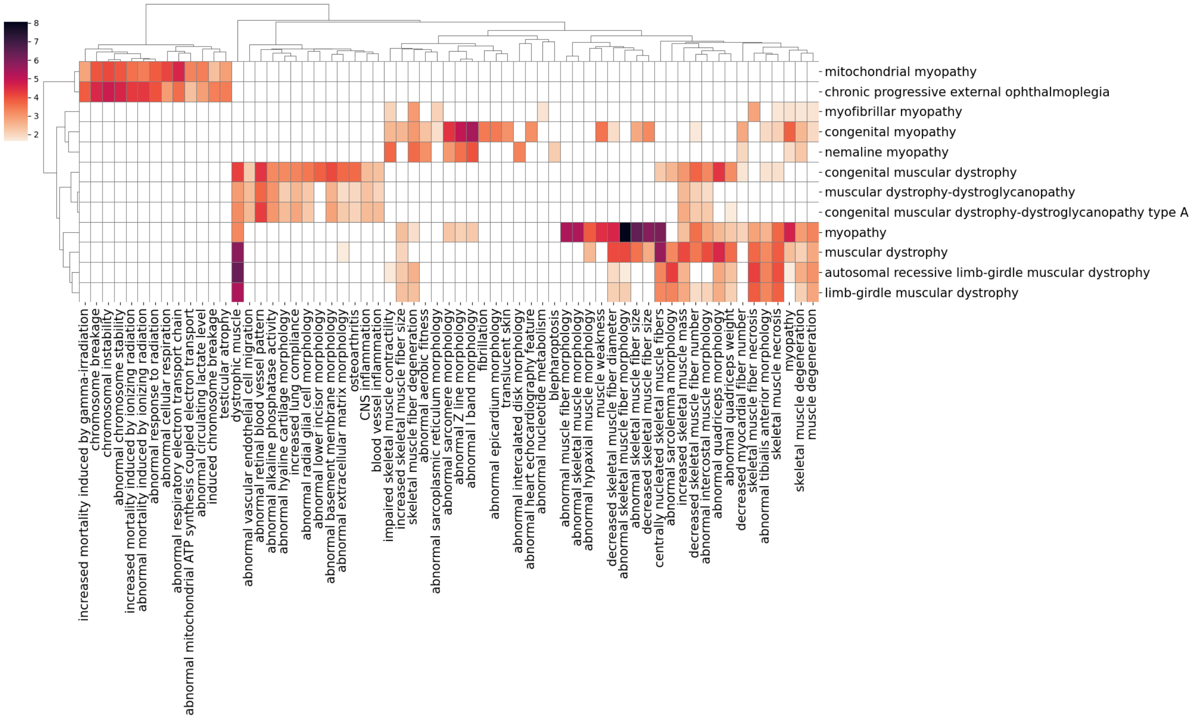


**Figure S36. DOID:423 (myopathy)**

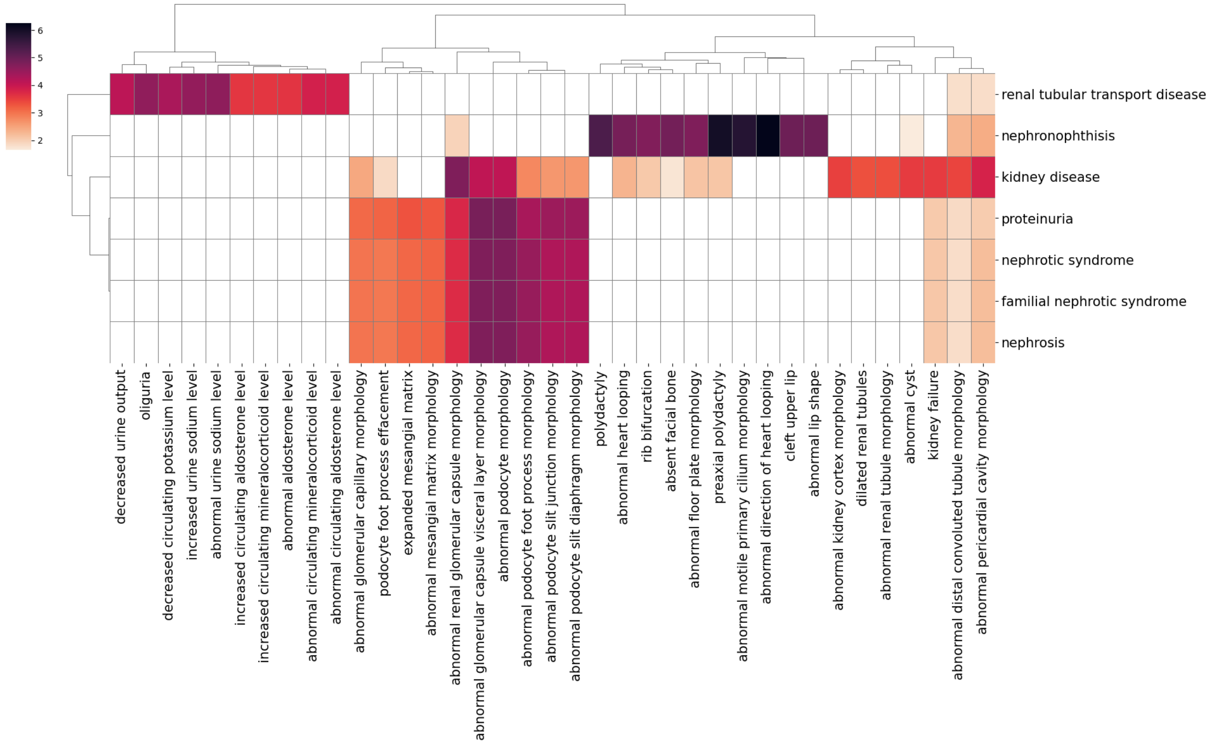


**Figure S37. DOID:557 (kidney disease)**

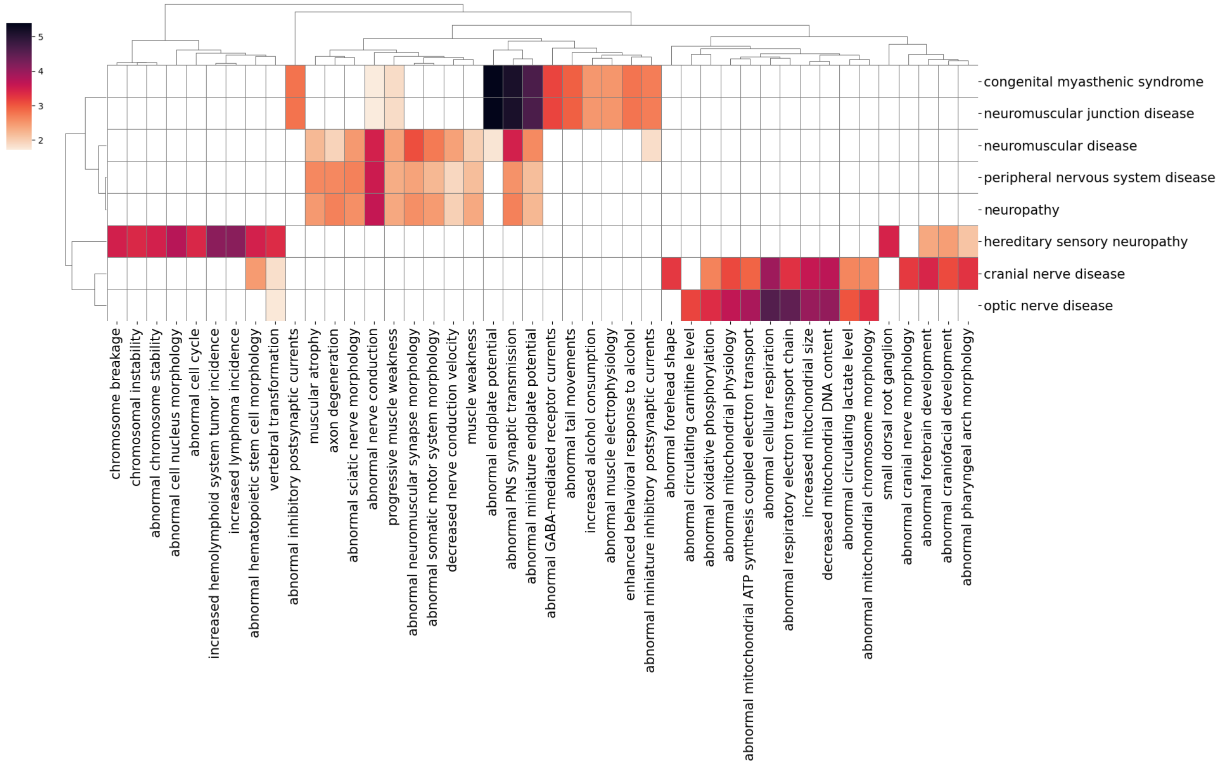


**Figure S38. DOID:574 (peripheral nervous system disease)**

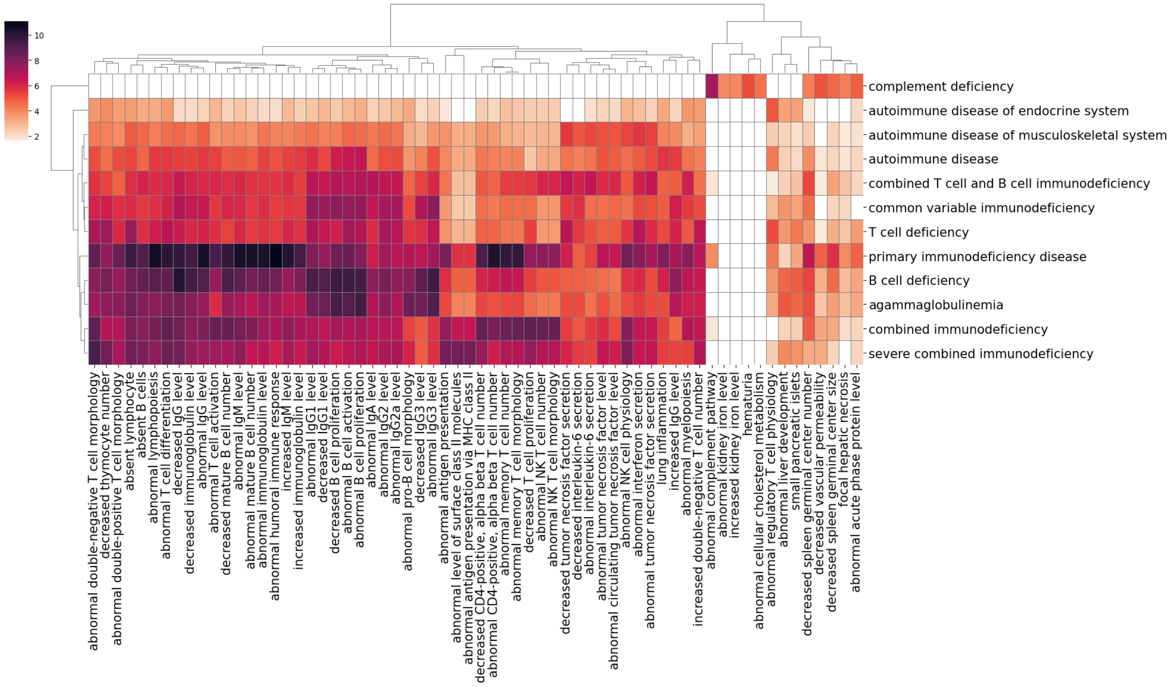


**Figure S39. DOID:612 (primary immunodeficiency disease)**

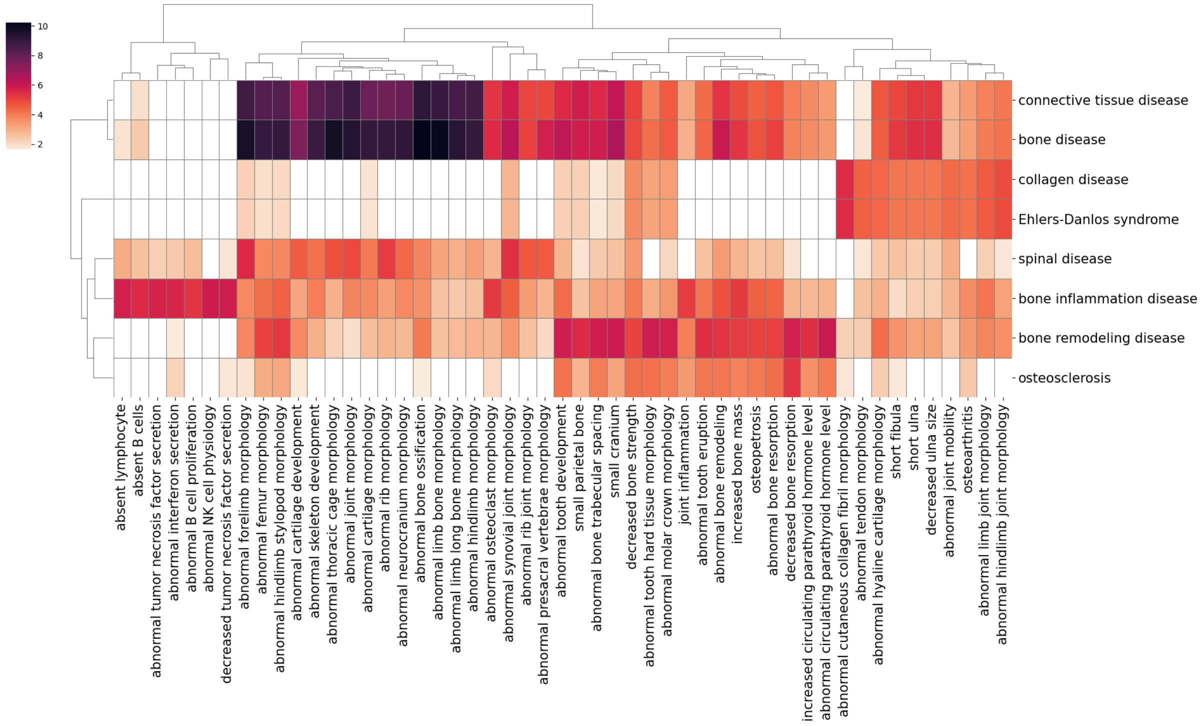


**Figure S40. DOID:65 (connective tissue disease)**

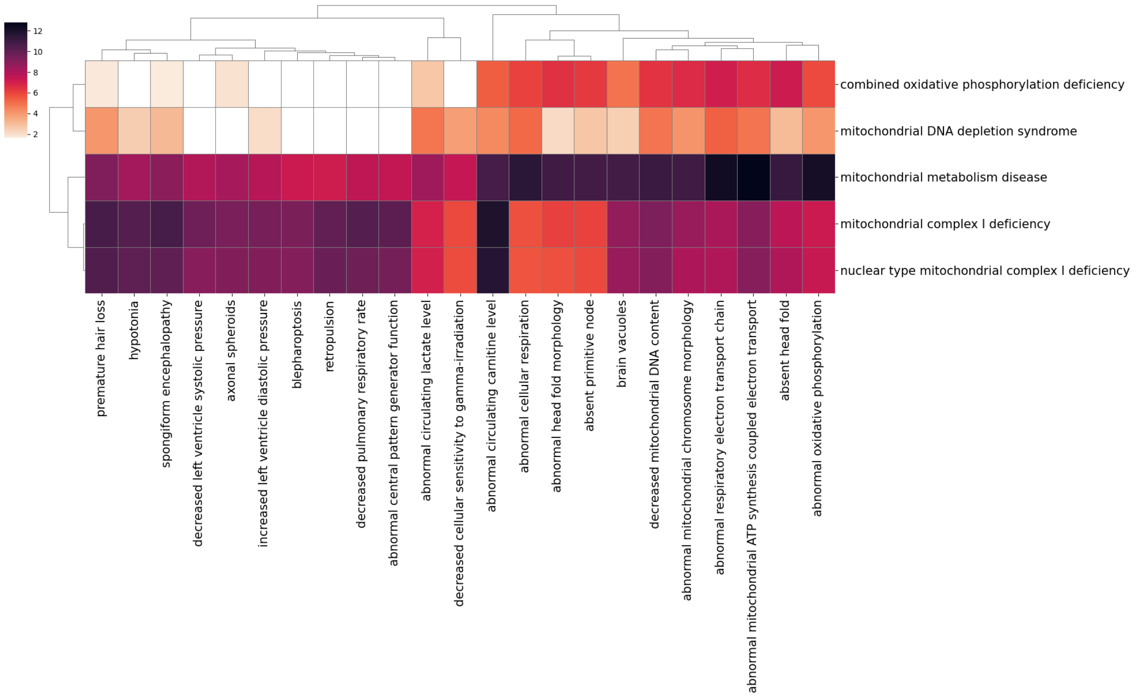


**Figure S41. DOID:700 (mitochondrial metabolism disease)**

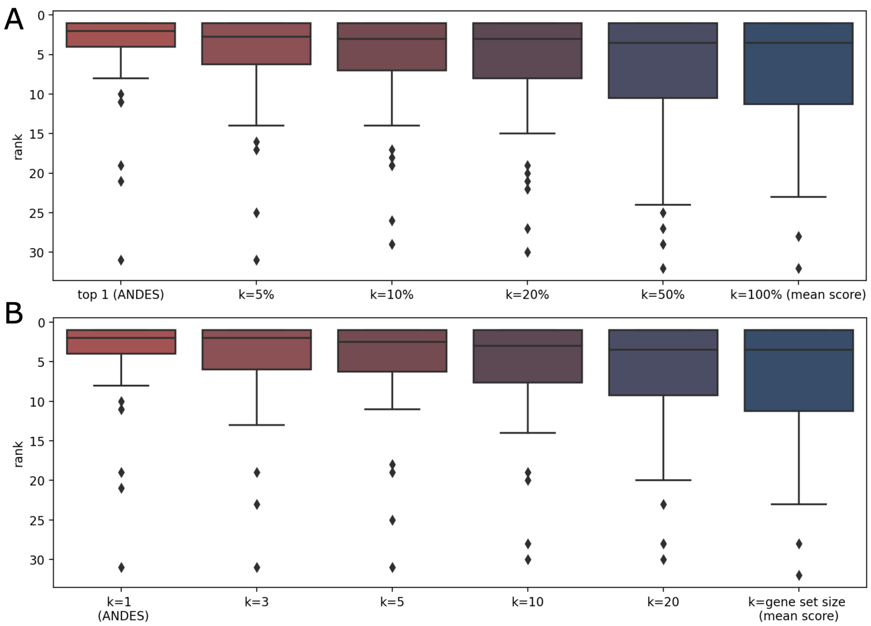


**Figure S42. Using more than the top matching genes as input to ANDES does not improve the performance of matching GO and KEGG terms. (A)** Boxplots of the ranking of the correct matching GO term for 50 KEGG terms using the top $k\%$ matches (relative to the gene set size) show that the performance of estimating gene set relationships between KEGG and GO decreases as $k$ increases, until it converges to the performance using $k = 100\%$ (which is mathematically equivalent to the mean score method). The average gene set size is 81, so $k = 5\%$ is on average equivalent to using the top 4 genes as matches. Best-match and mean score are two extreme cases of using top $k\%$ matches. **(B)** Ranking of the correct matching GO term for KEGG terms using top $k$ match, where $k$ is a fixed number instead of a ratio relative to gene set size. Choosing the top 1 (ANDES) still has the best performance.

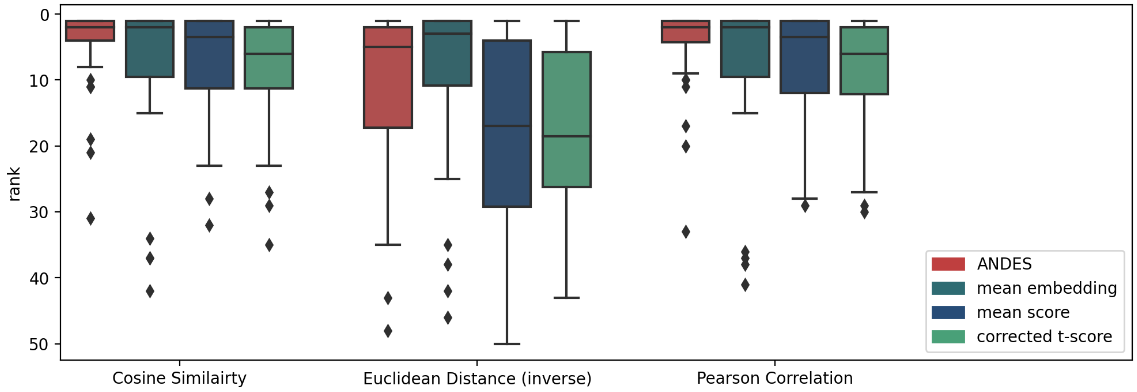


**Figure S43. KEGG-GO gene set matching performance when using other similarity metrics with ANDES.** Besides using cosine similarity, pairwise similarities are calculated using (inverse) Euclidean distance and Pearson correlation. Euclidean distance is inverted as $(\frac{1}{1+dist})$ to convert it to a similarity measurement. Boxplots show performance comparisons for ANDES, mean embedding, mean score, and corrected t-score. Similarity metrics that are not affected by embedding magnitude (cosine similarity and Pearson correlation) show better performance than magnitude-dependent ones such as Euclidean distance.

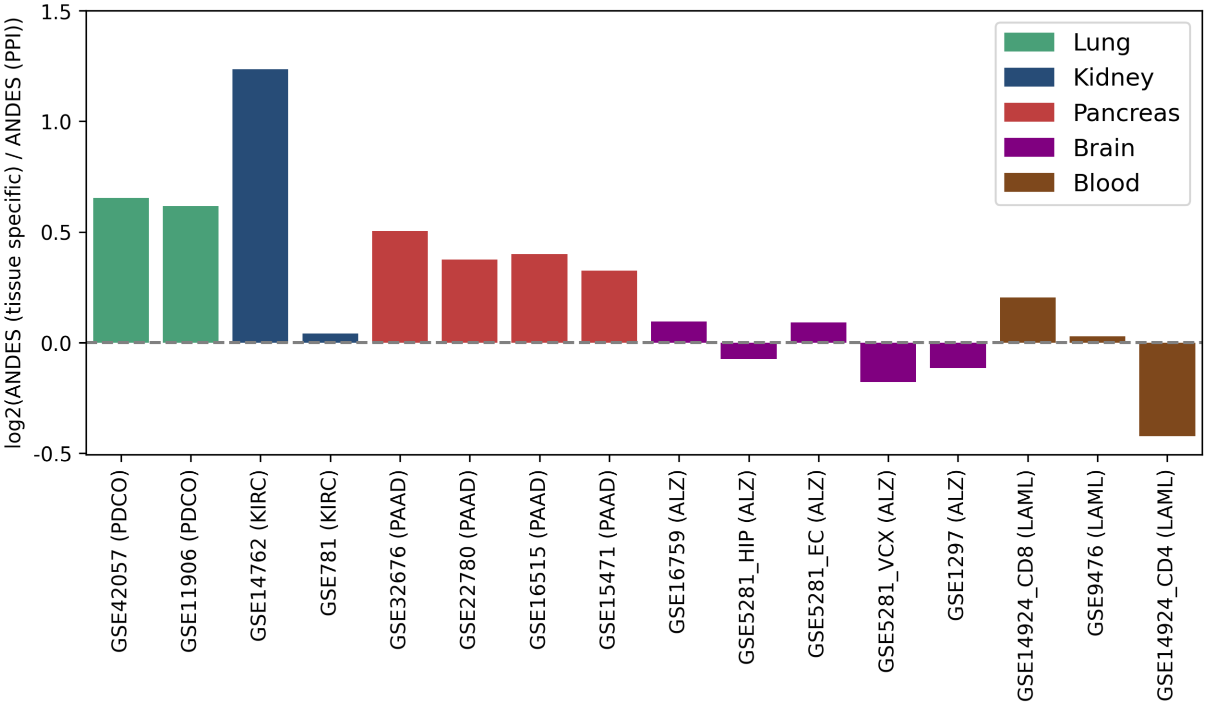


**Figure S44. Gene set enrichment analysis with a tissue-specific functional network improves performance compared to using a consensus PPI network.** Performance comparison between ANDES using relevant tissue-specific functional networks versus a PPI network in retrieving annotated KEGG terms. For each of 5 diseases (PDCO (Chronic Obstructive Pulmonary Disease), KIRC (Kidney Renal Clear Cell Carcinoma), PAAD (Pancreatic Adenocarcinoma), ALZ (Alzheimer's Disease) and LAML (Acute Myeloid Leukemia)), corresponding tissue-specific networks (lung, kidney, pancreas, brain, and blood) are used. The bar plot shows log2 AUPRC performance improvement using a tissue-specific functional network versus the consensus PPI network. The grey dotted line represents identical performance between the tissue-specific and PPI networks.

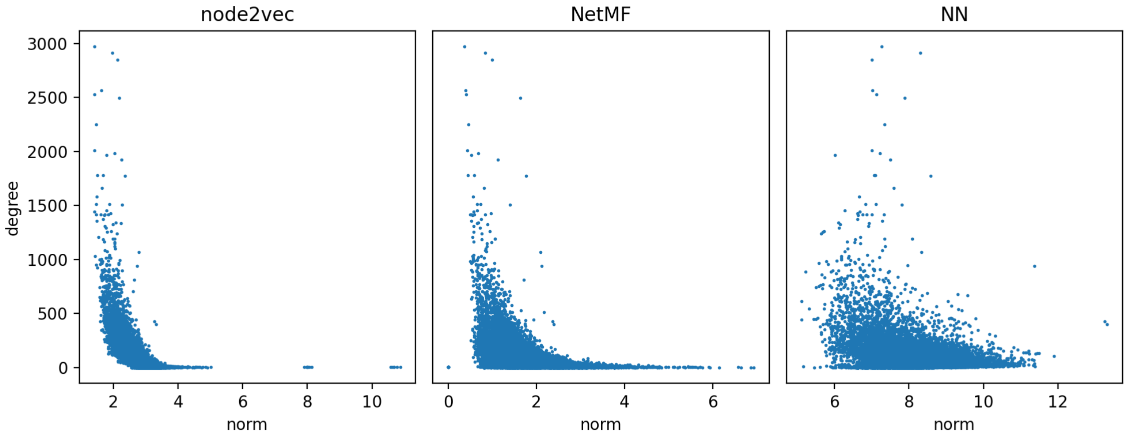


**Figure S45. PPI network node degree and embedding norms are negatively correlated.** Each point represents a gene, and the embeddings are generated from three methods. Regardless of embedding method used, there is a negative correlation between input node degree and the magnitude of the initial vector (node2vec: -0.642; NetMF: -0.420; NN: -0.231).
