## Supplemental Code for "ANDES: a novel best-match approach for enhancing gene set analysis in embedding spaces": README.md

# Binaries

This directory is where you can put any compiled binaries or scripts that you may need for the project, either from code in `src/` or from external resources. Make sure to include a simple bullet point explaining what each binary does / where it's from.
