## Supplemental Code for "ANDES: a novel best-match approach for enhancing gene set analysis in embedding spaces": demo.ipynb

```
# %% [code]
import numpy as np
import pandas as pd
from functools import partial
from multiprocessing import Pool

from collections import defaultdict
from sklearn import metrics

import src.load_data as ld
import src.set_analysis_func as func
import src.expression_analysis_func as expression_analysis_func


# %% [code]
# load embedding
node_vectors = np.loadtxt(
    './data/embedding/node2vec_consensus.csv', delimiter=',')
node_list = []
with open('./data/embedding/consensus_node.txt', 'r') as f:
    for line in f:
        node_list.append(line.strip())
        
S = metrics.pairwise.cosine_similarity(node_vectors, node_vectors)

# %% [code]
# create gene to embedding id mapping
g_node2index = {j:i for i,j in enumerate(node_list)}
g_index2node = {i:j for i,j in enumerate(node_list)}
g_node2index = defaultdict(lambda:-1, g_node2index)

# %% [code]
# load gene set data
GO_data = ld.load_gmt(
    './data/gene_sets/hsa_experimental_eval_BP_propagated.gmt')

GO2indices = ld.term2indexes(
    GO_data, g_node2index, upper=300, lower=10)

# %% [code]
# generate background gene list
GO_all_genes = set()
for x in GO_data:
    GO_all_genes = GO_all_genes.union(GO_data[x])
    
GO_all_genes = GO_all_genes.intersection(node_list)
GO_all_indices = [g_node2index[x] for x in GO_all_genes]

# %% [code]
f = partial(func.andes, matrix=S, g1_term2index=GO2indices, 
            g2_term2index=GO2indices, g1_population=GO_all_indices, 
            g2_population=GO_all_indices)

# %% [code]
# return ANDES raw and corrected score
f(('GO:0043648', 'GO:0006805'))

# %% [code]


# %% [markdown]
## ANDES as GSEA

# %% [code]
#load ranked list generated from gene expression data
ranked_list = pd.read_csv('./data/expression/GSE3467_rank.txt',
                          sep='\t', index_col=0, header=None)
ranked_list = [str(y) for y in ranked_list.index]
ranked_list = [g_node2index[y] for y in ranked_list if y in node_list]

# %% [code]
#directly load gene expression data
data = pd.read_csv('./data/expression/GSE3467.txt', skiprows=1, sep='\t')
condition = open('./data/expression/GSE3467.txt', 'r').readline().strip().split('\t')
ranked_list = expression_analysis_func.expression_data_to_ranked_list(data, condition)
ranked_list = [g_node2index[y] for y in ranked_list if y in node_list]

# %% [code]
f = partial(func.gsea_andes, ranked_list=ranked_list, matrix=S, 
            term2indices=GO2indices, 
            annotated_indices=GO_all_indices)

# %% [code]
geneset_terms = ['GO:0071466', 'GO:0006805', 'GO:0009410']

# %% [code]
f = partial(func.gsea_andes, ranked_list=ranked_list, matrix=S, 
            term2indices=GO2indices,
            annotated_indices=GO_all_indices)

zscores = [f(x)[1] for x in geneset_terms]

# %% [code]
def get_empirical_background(i):
    shuffled_ranked_list = expression_analysis_func.expression_data_to_ranked_list_label_shuffled(data, condition, seed=i)
    shuffled_ranked_list = [g_node2index[y] for y in shuffled_ranked_list if y in node_list]
    f = partial(func.gsea_andes, ranked_list=shuffled_ranked_list, 
                matrix=S, term2indices=GO2indices,
                annotated_indices=GO_all_indices, ite=100)
    rets = [f(x)[1] for x in geneset_terms]
    return rets

# %% [code]
with Pool(10) as p:
    rets = p.map(get_empirical_background, [i for i in range(100)])
background_scores = np.array(rets)

# %% [code]
empirical_pvalue = 1-np.sum(background_scores.T < np.array(zscores).reshape(-1,1), axis=1)/100

# %% [code]
```
