## Supplemental Code for "ANDES: a novel best-match approach for enhancing gene set analysis in embedding spaces": env.yml

```
name: ANDES
channels:
  - defaults
dependencies:
  - pip=23.3=py38h06a4308_0
  - python=3.8.18=h955ad1f_0
  - pip:
    - joblib==1.3.2
    - numpy==1.24.4
    - pandas==2.0.3
    - python-dateutil==2.8.2
    - pytz==2023.3.post1
    - scikit-learn==1.3.2
    - scipy==1.10.1
    - six==1.16.0
    - threadpoolctl==3.2.0
    - tzdata==2023.3
```
