## Supplemental Code for "ANDES: a novel best-match approach for enhancing gene set analysis in embedding spaces": drug_target.ipynb

```
# %% [code]
import pandas as pd
import numpy as np
import networkx as nx
import umap
import pickle

from collections import defaultdict
from sklearn import metrics
import matplotlib.pyplot as plt
import seaborn as sns
from matplotlib.pyplot import figure
import matplotlib.patches as mpatches
from collections import Counter

import helper as helper

# %% [code]
disease_slim_term = []
with open('./data/drug_info/DO_slim.txt', 'r') as f:
    for line in f:
        tokens = line.split('\t')
        disease_slim_term.append(tokens[0])

# %% [code]
drug_term2name = helper.term2name('./data/gmt/drugbank.2301.gmt')
omim_term2name = helper.term2name('./data/gmt/omim.20231030.gmt')

# %% [code]
result = pd.read_csv('./results/drug_disease/drug_disease_fixed_seed.csv', index_col=0)

# %% [code]
with open('./data/drug_info/drug_bank_atc.pickle', 'rb') as handle:
    drug2atc = pickle.load(handle)
    
with open('./data/drug_info/atc_name.pickle', 'rb') as handle:
    atc2name = pickle.load(handle)

# %% [code]
atcs_type = []
atcs2term_disctinct = defaultdict(set)
for x in list(result.index):
    codes = [y[:1] for y in drug2atc[x]]
    if len(codes)>0:
        code_counter = Counter(codes)
        sorted_codes = sorted(code_counter, key=lambda y: code_counter[y])
        atcs_type.append(sorted_codes[-1])
        atcs2term_disctinct[sorted_codes[-1]].add(x)
    else:
        atcs_type.append('')

# %% [code]
atcs2terms = defaultdict(set)
atcs_type_multi = []
for x in list(result.index):
    codes = [y[:1] for y in drug2atc[x]]
    atcs_type_multi.append(set(codes))
    for y in codes:
        atcs2terms[y].add(x)

# %% [code]
from sklearn.decomposition import PCA
pca = PCA(n_components=0.95)
pca_embedding = pca.fit_transform(result.values)

# %% [code]
colormap = {'L': '#e31a1c', 'B': '#1f78b4', '': '#808080', 'A': '#b2df8a', 
            'S': '#33a02c', 'V': '#808080', 'D': '#fb9a99', 'C': '#a6cee3',
            'N': '#33a02c', 'R': '#fdbf6f', 'M': '#6a3d9a', 'G': '#b15928',
            'J': '#cab2d6', 'H': '#808080', 'P': '#808080'}

# %% [code]
fig = plt.figure(figsize=(15,15), dpi=400) 
ax = fig.add_subplot(projection='3d')
colors = [colormap[x] for x in atcs_type]
used_indices = []
selected_atc_codes = ['B', 'S', 'V', 'D', 'R', 'M', 'G', 'J', 'H', 'P', 'A', 'C' ,'N','L']
colored_atc = []
for x in colormap.keys():
    if x=='':
        continue
    terms = atcs2term_disctinct[x]
    if len(terms)<=11:
        continue
    colored_atc.append(x)
    indices = [list(result.index).index(y) for y in terms]
    used_indices+=indices
    ax.scatter(pca_embedding[:,1][indices], 
               pca_embedding[:,2][indices],
               pca_embedding[:,0][indices],
                s=30, c=colormap[x], alpha=1)

ax.set_xlim(-24, 25)
ax.set_ylim(-27, 25)
ax.set_zlim(-26, 30)

ax.set_xlabel('PC2 (18.2%)', fontsize=20)
ax.set_ylabel('PC3 (10.4%)', fontsize=20)
ax.set_zlabel('PC1 (32.2%)', fontsize=20)

# ax.view_init(elev=20., azim=-35, roll=0)
for tick in ax.xaxis.get_major_ticks():
    tick.tick1line.set_visible(False)
    tick.tick2line.set_visible(False)
    tick.label1.set_visible(False)
    tick.label2.set_visible(False)
    
for tick in ax.yaxis.get_major_ticks():
    tick.tick1line.set_visible(False)
    tick.tick2line.set_visible(False)
    tick.label1.set_visible(False)
    tick.label2.set_visible(False)
    
for tick in ax.zaxis.get_major_ticks():
    tick.tick1line.set_visible(False)
    tick.tick2line.set_visible(False)
    tick.label1.set_visible(False)
    tick.label2.set_visible(False)
    
patches = [mpatches.Patch(color=colormap[x], label=atc2name[x].lower()) 
           for x in sorted(colored_atc, key=lambda x: atc2name[x])]
plt.legend(handles = patches, ncol=2, prop={'size': 15.5},
           bbox_to_anchor=(0.99, 1))
ax.xaxis.pane.fill = False
ax.yaxis.pane.fill = False
ax.zaxis.pane.fill = False

# %% [code]
omim_slim_terms = list(set(disease_slim_term).intersection(result.columns))

# %% [code]
for x in atcs2terms.keys():
    print(atc2name[x])
    terms = list(atcs2terms[x])
    sub_matrix = result.loc[terms][omim_slim_terms]
    sub_matrix = sub_matrix.loc[:,((sub_matrix>1.64).sum(axis=0) != 0)]
    sub_matrix = sub_matrix.loc[((sub_matrix>1.64).sum(axis=1) != 0)]
    a_fig=sns.clustermap(sub_matrix.values, 
                yticklabels=[drug_term2name[x].split(',')[0] for x in sub_matrix.index], 
                xticklabels=[omim_term2name[x] for x in sub_matrix.columns], 
                   figsize=(8, 5+sub_matrix.shape[0]/5), 
                         dendrogram_ratio=0.1, cbar_pos=(0.2, 0.94, .5, .015),
                linewidths=0.5, linecolor='grey', mask=1-(sub_matrix.values>1.64),
               vmin=1.64, cmap='rocket_r',cbar_kws={"orientation": "horizontal"})
    a_fig.ax_heatmap.set_xticklabels(a_fig.ax_heatmap.get_xmajorticklabels(), fontsize = 17)
    a_fig.ax_heatmap.set_yticklabels(a_fig.ax_heatmap.get_ymajorticklabels(), fontsize = 17)
    col = a_fig.ax_col_dendrogram.get_position()
    a_fig.ax_col_dendrogram.set_position([col.x0, col.y0, col.width, col.height*0.2])
    plt.show()


# %% [code]


# %% [code]
```
