## Supplemental Code for "ANDES: a novel best-match approach for enhancing gene set analysis in embedding spaces": GSEA.ipynb

```
# %% [code]
import pandas as pd
import numpy as np
import sklearn.metrics as metrics
import matplotlib.pyplot as plt

from collections import defaultdict
import scipy
from sklearn.metrics import average_precision_score
from sklearn.metrics import precision_score

import helper as helper

# %% [code]
kegg2genes = defaultdict(list)
with open('./data/CPDB_pathways_genes.tab', 'r') as f:
    f.readline()
    for line in f:
        tokens = line.strip().split('\t')
        database = tokens[2]
        term = tokens[1]
        genes = tokens[3]
        if database != 'KEGG':
            continue
        genes = genes.split(',')
        term = term.split(':')[1]
        kegg2genes[term]=genes

# %% [code]
geo2kegg = {}
with open('./data/geo2kegg.txt', 'r') as f:
    for line in f:
        tokens = line.strip().split(' ')
        geo2kegg[tokens[0]] = tokens[-1]

# %% [code]
malaterms = {}
for x in set(geo2kegg.values()):
    mapped_terms = helper.parse_mala_file('./data/malakegg/'+x+'.txt')
    malaterms[x] = mapped_terms
    
all_malaterms = set()
for x in malaterms.keys():
    all_malaterms = all_malaterms.union([y[0] for y in malaterms[x]])
len(all_malaterms)

# %% [markdown]
# compare with GSEA

# %% [code]
andes_gsea_score = pd.read_csv('./results/enrichment_analysis/andes_gsea_consensus_node2vec_fixed_seed.csv', index_col=0)
andes_gsea_score = andes_gsea_score.abs()


# %% [code]
terms = list(set(andes_gsea_score.columns).intersection(all_malaterms))
geos = set(andes_gsea_score.index)
geos = [x for x in geos if np.sum(andes_gsea_score.loc[x])!=0]

# %% [code]
andes_gsea_score = andes_gsea_score[terms]

# %% [code]
andes_gsea_auprcs = []
gsea_s_auprcs = []
gspa_s_auprcs = []
for i, x in enumerate(geos):   
    term = geo2kegg[x]
    mala_labels = malaterms[term]
    mala_labels = [x for x,y in mala_labels if y>0]
    print(x, len(mala_labels))
    andes_gsea_s = list(andes_gsea_score.loc[x])
    gspa_s = helper.get_GSPA_score(x, terms)
    gsea_s = helper.get_GSEA_score(x, terms)
    
    gspa_s = np.array(gspa_s)
    gspa_s = np.abs(gspa_s)
    gspa_s[np.isinf(gspa_s)] = 0
    
    gsea_s = np.array(gsea_s)
    gsea_s = np.abs(gsea_s)
    gsea_s[np.isinf(gsea_s)] = 0
  
    labels = [1 if x in mala_labels else 0 for x in terms]

    andes_gsea_auprc = average_precision_score(labels, andes_gsea_s)
    gsea_s_auprc = average_precision_score(labels, gsea_s)
    gspa_s_auprc = average_precision_score(labels, gspa_s)
 
    andes_gsea_auprcs.append(andes_gsea_auprc)
    gsea_s_auprcs.append(gsea_s_auprc)
    gspa_s_auprcs.append(gspa_s_auprc)
    

# %% [code]
andes_gsea_auprcs = np.array(andes_gsea_auprcs)
gsea_s_auprcs = np.array(gsea_s_auprcs)
gspa_s_auprcs = np.array(gspa_s_auprcs)

# %% [code]
order = list(np.argsort(andes_gsea_auprcs))

# %% [code]
plt.figure(1,figsize=(5,8), dpi=100) 
plt.scatter(gsea_s_auprcs[order], [i for i in range(len(andes_gsea_auprcs))],
            marker='<', c = '#49a078', label='GSEA')
plt.scatter(gspa_s_auprcs[order], [i for i in range(len(andes_gsea_auprcs))],
            marker='>', c='#274c77', label='GSPA')

plt.scatter(andes_gsea_auprcs[order], [i for i in range(len(andes_gsea_auprcs))],
            label='ANDES', color='#bf3f3f')

plt.yticks([i for i in range(len(geos))], 
           [ x+' ('+str(geo2kegg[x])+')' for x in np.array(geos)[order]])
plt.tick_params(axis='both', which='major', labelsize=6)
plt.xlabel('AUPRC', fontsize=8)
plt.xticks([0.0, 0.2, 0.4, 0.6, 0.8], [0.0, 0.2, 0.4, 0.6, 0.8])
plt.xlim(-0.01, 0.9)
plt.legend(prop={'size': 8})

plt.tight_layout()

# %% [markdown]
# compare with function graph

# %% [code]
andes_gsea_func_score = pd.read_csv('./results/enrichment_analysis/andes_gsea_functional_network.txt', index_col=0)
andes_gsea_func_score = andes_gsea_func_score.abs()
andes_gsea_func_score = andes_gsea_func_score[terms]

# %% [code]
andes_gsea_func_auprcs = []
for i, x in enumerate(andes_gsea_func_score.index):   
    term = geo2kegg[x]
    mala_labels = malaterms[term]
    mala_labels = [x for x,y in mala_labels if y>0]
    print(x, len(mala_labels))
    andes_gsea_func_s = list(andes_gsea_func_score.loc[x])
  
    labels = [1 if x in mala_labels else 0 for x in terms]

    andes_gsea_func_auprc = average_precision_score(labels, andes_gsea_func_s)
 
    andes_gsea_func_auprcs.append(andes_gsea_func_auprc)
    

# %% [code]
andes_gsea_func_auprcs = np.array(andes_gsea_func_auprcs)

# %% [code]
selected_geos = [
    'GSE42057 (PDCO)', 'GSE11906 (PDCO)', 'GSE14762 (KIRC)', 'GSE781 (KIRC)', 'GSE32676 (PAAD)',
    'GSE22780 (PAAD)', 'GSE16515 (PAAD)', 'GSE15471 (PAAD)', 'GSE16759 (ALZ)', 'GSE5281_HIP (ALZ)', 
    'GSE5281_EC (ALZ)', 'GSE5281_VCX (ALZ)', 'GSE1297 (ALZ)', 'GSE14924_CD8 (LAML)', 
    'GSE9476 (LAML)', 'GSE14924_CD4 (LAML)']

# %% [code]
selected_ppi_scores = [andes_gsea_auprcs[geos.index(x.split(' ')[0])] for x in selected_geos]
selected_func_scores = [andes_gsea_func_auprcs[list(andes_gsea_func_score.index).index(x.split(' ')[0])] for x in selected_geos]

# %% [code]
import matplotlib.patches as mpatches
colors = ['#49a078', '#49a078', '#274c77', '#274c77', '#bf3f3f', '#bf3f3f', '#bf3f3f', '#bf3f3f']+\
         ['#800080', '#800080', '#800080', '#800080', '#800080', '#7e481c', '#7e481c', '#7e481c']
fig, ax = plt.subplots(figsize=(10,4), dpi=300)
plt.bar([i for i in range(16)], np.log2(np.array(selected_func_scores)/np.array(selected_ppi_scores)),
        color=colors)
plt.xticks([i for i in range(16)], selected_geos, rotation=90)
patches = [mpatches.Patch(color='#49a078', label='Lung'),
           mpatches.Patch(color='#274c77', label='Kidney'),
           mpatches.Patch(color='#bf3f3f', label='Pancreas'),
           mpatches.Patch(color='#800080', label='Brain'),
           mpatches.Patch(color='#7e481c', label='Blood')]
plt.legend(handles=patches, prop={'size': 12})
plt.yticks([-0.5, 0, 0.5, 1, 1.5,], ['-0.5', '0.0', '0.5', '1.0', '1.5'])
plt.xlim(-0.5, 15.5)
plt.axhline(y=0, color='grey', linestyle='dashed')
plt.ylabel('log2(ANDES (tissue specific) / ANDES (PPI))')

# %% [markdown]
# compare with hypergeomtric

# %% [code]
andes_set_score = pd.read_csv('./results/enrichment_analysis/andes_set_consensus_node2vec_fdr_only_fixed_seed.csv', index_col=0)
andes_set_score = andes_set_score.abs()

# %% [code]
hyper_score = pd.read_csv('./results/enrichment_analysis/hypergeom_score_fdr_only.csv', index_col=0)

# %% [code]
terms = list(set(andes_set_score.columns).intersection(all_malaterms))
geos = set(andes_set_score.index)
geos = [x for x in geos if np.sum(andes_set_score.loc[x])!=0]

# %% [code]
andes_set_score = andes_set_score[terms]
hyper_score = hyper_score[terms]

# %% [code]
andes_set_auprcs = []
hyper_auprcs = []

for i, x in enumerate(geos):   
    if x not in andes_set_score.index:
        continue
    term = geo2kegg[x]
    mala_labels = malaterms[term]
    mala_labels = [x for x,y in mala_labels if y>0]

    andes_set_s = list(andes_set_score.loc[x])
    hyper_s = list(1-hyper_score.loc[x])
   
    labels = [1 if x in mala_labels else 0 for x in terms]
    
    andes_set_auprc = average_precision_score(labels, andes_set_s) 
    hyper_auprc = average_precision_score(labels, hyper_s)
   
    andes_set_auprcs.append(andes_set_auprc)
    hyper_auprcs.append(hyper_auprc)
    

# %% [code]
andes_set_auprcs = np.array(andes_set_auprcs)
hyper_auprcs = np.array(hyper_auprcs)

# %% [code]
order = list(np.argsort(andes_set_auprcs))

# %% [code]
plt.figure(1,figsize=(5,8), dpi=100) 
plt.scatter(hyper_auprcs[order], [i for i in range(len(hyper_auprcs))],
            marker='d', c='#bebebe', label='hypergeom')

plt.scatter(andes_set_auprcs[order], [i for i in range(len(andes_set_auprcs))],
            label='ANDES', color='#bf3f3f')

plt.yticks([i for i in range(len(geos))], 
           [ x+' ('+str(geo2kegg[x])+')' for x in np.array(geos)[order]])
plt.tick_params(axis='both', which='major', labelsize=6)
plt.xlabel('AUPRC', fontsize=8)
plt.xticks([0.0, 0.2, 0.4, 0.6, 0.8], [0.0, 0.2, 0.4, 0.6, 0.8])
plt.xlim(-0.01, 0.9)
plt.legend(prop={'size': 8})

plt.tight_layout()

# %% [code]


# %% [code]
```
