## Supplemental Code for "ANDES: a novel best-match approach for enhancing gene set analysis in embedding spaces": helper.py

```
import pandas as pd
import numpy as np

def term2name(file_name):
    term2name = {}
    with open(file_name, 'r') as f:
        for line in f:
            tokens = line.split('\t')
            tokens = [x.strip() for x in tokens]
            term2name[tokens[0]]=tokens[1]
            
    return term2name

def paths_to_root(graph, term, root):
    if term == root:
        return [[term]]
    parents = [x[0] for x in list(graph.in_edges(term))]
    ret = []
    for x in parents:
        for y in paths_to_root(graph, x, root):
            ret.append(y+[term])
    return ret

def get_sub_matrix_single_selected(term, df, x_slim2terms, ordered_all_x_terms, ordered_all_y_terms, k=5):
    x_terms = x_slim2terms[term]
    ordered_x_terms = [x for x in ordered_all_x_terms if x in x_terms]
    sub_m = df.loc[ordered_x_terms]
    sum_m_values = sub_m.values
    
    columns = list(df.columns)
    
    selected_columns = set()
    for i in range(sum_m_values.shape[0]):
        top_k = sorted(range(sum_m_values.shape[1]), key=lambda j: sum_m_values[i][j])[-1*k:]
        for y in top_k:
            selected_columns.add(columns[y])
            
    return sub_m[list(selected_columns)]

def load_model_org_data(org1, org2):

    best_avg = pd.read_csv('./results/model_org/'+org1+'_'+org2+'/best_average_slim_p0_fixed_seed.csv', 
                     delimiter=',', index_col=0)

    t_score = pd.read_csv('./results/model_org/'+org1+'_'+org2+'/t_score_slim_p0_fixed_seed.csv', 
                     delimiter=',', index_col=0)

    mean_emb = pd.read_csv('./results/model_org/'+org1+'_'+org2+'/mean_emb_slim_value_fixed_seed.csv', 
                          delimiter=',', index_col=0)

    mean_value = pd.read_csv('./results/model_org/'+org1+'_'+org2+'/mean_value_slim_value_fixed_seed.csv', 
                          delimiter=',', index_col=0)
    
    shared_terms = list(set(best_avg.index).intersection(best_avg.columns))
    
    best_avg_matrix = best_avg.loc[shared_terms][shared_terms].values
    t_score_matrix = t_score.loc[shared_terms][shared_terms].values
    mean_emb_matrix = mean_emb.loc[shared_terms][shared_terms].values
    mean_value_matrix = mean_value.loc[shared_terms][shared_terms].values
    
    return [best_avg_matrix, mean_emb_matrix,  t_score_matrix, mean_value_matrix], shared_terms


def get_model_org_rank(matrices, axis):
    # axis determine the direction 
    # for example the rank of hsa terms in mmu 0
    # the rank of mmu terms in hsa is 1
    orders = [x.argsort(axis=axis) for x in matrices]
    ranks = [x.argsort(axis=axis) for x in orders]
    ranks = [x.shape[0]-np.diagonal(x) for x in ranks]
    
    return ranks


def load_kegg_go_data(postfix, selected_row_terms, selected_column_terms):

    best_avg = pd.read_csv('./results/kegg_go/best_average_p0_'+postfix+'.csv', 
                     delimiter=',', index_col=0)

    t_score = pd.read_csv('./results/kegg_go/t_score_p0_'+postfix+'.csv', 
                     delimiter=',', index_col=0)

    mean_emb = pd.read_csv('./results/kegg_go/mean_emb_values_'+postfix+'.csv', 
                          delimiter=',', index_col=0)

    mean_value = pd.read_csv('./results/kegg_go/mean_value_values_'+postfix+'.csv', 
                          delimiter=',', index_col=0)
    
    best_avg_matrix = best_avg.loc[selected_row_terms][selected_column_terms].values
    t_score_matrix = t_score.loc[selected_row_terms][selected_column_terms].values
    mean_emb_matrix = mean_emb.loc[selected_row_terms][selected_column_terms].values
    mean_value_matrix = mean_value.loc[selected_row_terms][selected_column_terms].values
    
    return [best_avg_matrix, mean_emb_matrix,  t_score_matrix, mean_value_matrix]


def load_kegg_go_matrix_data(postfix, selected_row_terms, selected_column_terms):
    
    best_avg = pd.read_csv('./results/kegg_go/best_average_p0_'+postfix+'.csv', 
                     delimiter=',', index_col=0)


    t_score = pd.read_csv('./results/kegg_go/t_score_p0_'+postfix+'.csv', 
                     delimiter=',', index_col=0)


    mean_value = pd.read_csv('./results/kegg_go/mean_value_values_'+postfix+'.csv', 
                          delimiter=',', index_col=0)
    
    best_avg_matrix = best_avg.loc[selected_row_terms][selected_column_terms].values
    t_score_matrix = t_score.loc[selected_row_terms][selected_column_terms].values
    mean_value_matrix = mean_value.loc[selected_row_terms][selected_column_terms].values
    
    return[best_avg_matrix,  t_score_matrix, mean_value_matrix]


def load_kegg_go_k_data(postfix, selected_row_terms, selected_column_terms, ks):
    dfs = []
    for x in ks:
        dfs.append(pd.read_csv('./results/kegg_go/top_'+x+'_average_p0_consensus_'+postfix+'_fixed_seed.csv',
                               delimiter=',', index_col=0))
        
    matrices = [x.loc[selected_row_terms][selected_column_terms].values for x in dfs]
    
    return matrices

def generate_kegg_go_result(matrices, label_matrix):
    result_lists = [[] for i in range(len(matrices))]
    for i in range(label_matrix.shape[0]):
        for j,m in enumerate(matrices):
            label_indices = np.nonzero(label_matrix[i])
            
            order = matrices[j][i].argsort()
            ranks = order.argsort()
            result_lists[j].append(np.mean(ranks[list(label_indices)]))
            
    return result_lists          
    
def parse_mala_file(file):
    ret = []
    with open(file, 'r') as f:
        f.readline()
        for line in f:
            tokens = line.strip().split(',')
            term = tokens[0][1:-1]
            score = float(tokens[2])
            ret.append((term, score))            
    return ret

def get_GSPA_score(geo_name, terms):
    data = pd.read_csv('./results/enrichment_analysis/GSPA/'+geo_name+'_GSPA_results_our_ppi.csv', index_col='Gene Set')
    for x in set(terms).difference(data.index):
        data.loc[x] = [0,0,0,0]
    data = data.loc[terms]
    data = data.loc[terms]
    return np.array(data['NES'])

def get_GSEA_score(geo_name, terms):
    data = pd.read_csv('./results/enrichment_analysis/GSEA/'+geo_name+'_result.csv', index_col='Term', delimiter=',')
    for x in set(terms).difference(data.index):
        data.loc[x] = ['', 0,0,0,0,0,0,0,0]
    data = data.loc[terms]
    return np.array(data['NES']) 
```
