## Supplemental Code for "ANDES: a novel best-match approach for enhancing gene set analysis in embedding spaces": KEGG-GO-analysis.ipynb

```
# %% [code]
import pandas as pd
import numpy as np
import sklearn.metrics as metrics
import pickle
import umap

from collections import defaultdict
import matplotlib.pyplot as plt
import seaborn as sns
from matplotlib.pyplot import figure

import helper as helper

# %% [code]
matched_pairs = []
with open('./data/matched_pair.txt', 'r') as f:
    for line in f:
        tokens = line.strip().split('\t')
        matched_pairs.append((tokens[0], tokens[1]))

# %% [code]
selected_kegg_terms = sorted(list(set([x for x,y in matched_pairs])))
selected_go_terms = sorted(list(set([y for x,y in matched_pairs])))
print(len(selected_kegg_terms), len(selected_go_terms))

# %% [code]
labels = np.zeros((len(selected_kegg_terms), len(selected_go_terms)))
for i,x in enumerate(selected_kegg_terms):
    for j,y in enumerate(selected_go_terms):
        if (x,y) in matched_pairs:
            labels[i][j] = 1

# %% [markdown]
# figure 2A

# %% [code]
netmf_data = helper.load_kegg_go_data('consensus_netmf_fixed_seed', selected_kegg_terms, selected_go_terms)
node2vec_data = helper.load_kegg_go_data('consensus_node2vec_fixed_seed', selected_kegg_terms, selected_go_terms)
etna_data = helper.load_kegg_go_data('consensus_etna_fixed_seed', selected_kegg_terms, selected_go_terms)
netmf_data_results = helper.generate_kegg_go_result(netmf_data, labels)
node2vec_data_results = helper.generate_kegg_go_result(node2vec_data, labels)
etna_data_results = helper.generate_kegg_go_result(etna_data, labels)

netmf_data_results = pd.DataFrame(50-(np.array(netmf_data_results)).T,
                                  columns=['best_average',  'mean embedding',
                                           'corrected t-score', 'mean score'])

node2vec_data_results = pd.DataFrame(50-(np.array(node2vec_data_results)).T,
                                  columns=['best_average',  'mean embedding',
                                           'corrected t-score', 'mean score'])

etna_data_results = pd.DataFrame(50-(np.array(etna_data_results)).T,
                                  columns=['best_average', 'mean embedding',
                                           'corrected t-score', 'mean score' ])

# %% [code]
figure1a_df_y = list(node2vec_data_results['best_average'])+\
              list(node2vec_data_results['mean embedding'])+\
              list(node2vec_data_results['mean score'])+\
              list(node2vec_data_results['corrected t-score'])+\
              list(netmf_data_results['best_average'])+\
              list(netmf_data_results['mean embedding'])+\
              list(netmf_data_results['mean score'])+\
              list(netmf_data_results['corrected t-score'])+\
              list(etna_data_results['best_average'])+\
              list(etna_data_results['mean embedding'])+\
              list(etna_data_results['mean score'])+\
              list(etna_data_results['corrected t-score'])

figure1a_df_x = list([0 for i in range(node2vec_data_results.shape[0])])+\
                list([1 for i in range(node2vec_data_results.shape[0])])+\
                list([2 for i in range(node2vec_data_results.shape[0])])+\
                list([3 for i in range(node2vec_data_results.shape[0])])+\
                list([5 for i in range(netmf_data_results.shape[0])])+\
                list([6 for i in range(netmf_data_results.shape[0])])+\
                list([7 for i in range(netmf_data_results.shape[0])])+\
                list([8 for i in range(netmf_data_results.shape[0])])+\
                list([10 for i in range(etna_data_results.shape[0])])+\
                list([11 for i in range(etna_data_results.shape[0])])+\
                list([12 for i in range(etna_data_results.shape[0])])+\
                list([13 for i in range(etna_data_results.shape[0])])
figure1a_df = pd.DataFrame([figure1a_df_x, figure1a_df_y]).T
figure1a_df = figure1a_df.rename(columns={0:'x', 1:'y'})
figure1a_df['x'] = np.array(figure1a_df['x']).astype(int)

# %% [code]

import matplotlib.patches as mpatches
plt.figure(1,figsize=(12,4), dpi=200) 
figure1a_pal = {0:'#bf3f3f', 1:'#2e6b72', 2:'#274c77', 3:'#49a078', 4:'#ffffff',
                5:'#bf3f3f', 6:'#2e6b72', 7:'#274c77', 8:'#49a078', 9:'#ffffff',
                10:'#bf3f3f', 11:'#2e6b72', 12:'#274c77', 13:'#49a078', 14:'#ffffff',
                15:'#ffffff', 16:'#ffffff', 17:'#ffffff',18:'#ffffff'}
patch0 = mpatches.Patch(color='#bf3f3f', label='ANDES')
patch1 = mpatches.Patch(color='#2e6b72', label='mean embedding')
patch2 = mpatches.Patch(color='#274c77', label='mean score')
patch3 = mpatches.Patch(color='#49a078', label='corrected t-score')

a = sns.boxplot(x='x', y='y', data=figure1a_df, palette=figure1a_pal, order=np.arange(18))
a.invert_yaxis() 
a.set_xticks([1.5, 6.5, 11.5], labels=['node2vec', 'NetMF', 'NN'])
a.set_xlabel('')
a.set_ylabel('')
plt.ylabel('rank')
plt.legend(handles = [patch0,patch1,patch2,patch3], loc='lower right' )

# %% [markdown]
# figure 2C

# %% [code]
jaccard_data = helper.load_kegg_go_matrix_data('consensus_jaccard_fixed_seed', selected_kegg_terms, selected_go_terms)
degree_data = helper.load_kegg_go_matrix_data('consensus_degree_fixed_seed', selected_kegg_terms, selected_go_terms)

jaccard_data_results = helper.generate_kegg_go_result(jaccard_data, labels)
degree_data_results = helper.generate_kegg_go_result(degree_data, labels)

jaccard_data_results = pd.DataFrame(50-(np.array(jaccard_data_results)).T,
                                      columns=['best_average',
                                               'corrected t-score', 'mean score' ])
degree_data_results = pd.DataFrame(50-(np.array(degree_data_results)).T,
                                      columns=['best_average',
                                               'corrected t-score', 'mean score' ])

# %% [code]
figure1b_df_y =list(jaccard_data_results['best_average'])+\
              list(jaccard_data_results['mean score'])+\
              list(jaccard_data_results['corrected t-score'])+\
              list(degree_data_results['best_average'])+\
              list(degree_data_results['mean score'])+\
              list(degree_data_results['corrected t-score'])

figure1b_df_x = list([0 for i in range(jaccard_data_results.shape[0])])+\
                list([1 for i in range(jaccard_data_results.shape[0])])+\
                list([2 for i in range(jaccard_data_results.shape[0])])+\
                list([4 for i in range(degree_data_results.shape[0])])+\
                list([5 for i in range(degree_data_results.shape[0])])+\
                list([6 for i in range(degree_data_results.shape[0])])
                
figure1b_df = pd.DataFrame([figure1b_df_x, figure1b_df_y]).T
figure1b_df = figure1b_df.rename(columns={0:'x', 1:'y'})
figure1b_df['x'] = np.array(figure1b_df['x']).astype(int)

# %% [code]
plt.figure(1,figsize=(3,4), dpi=200) 
figure1b_pal = {0:'#bf3f3f', 1:'#274c77', 2:'#49a078', 3:'#ffffff',
                4:'#bf3f3f', 5:'#274c77', 6:'#49a078', 7:'#ffffff',
                8:'#bf3f3f', 9:'#274c77', 10:'#49a078', 11:'#ffffff',
                15:'#ffffff'}
b = sns.boxplot(x='x', y='y', data=figure1b_df, palette=figure1b_pal, order=np.arange(7))
b.invert_yaxis() 
b.set_xlim(-1,7)
b.set_xticks([1, 5,], labels=['jaccard', 'degree'])
b.set_xlabel('')
b.set_ylabel('rank')

# %% [markdown]
# Sfigure 1

# %% [code]
netmf_data = helper.load_kegg_go_data('consensus_netmf_overlap_fixed_seed', selected_kegg_terms, selected_go_terms)
node2vec_data = helper.load_kegg_go_data('consensus_node2vec_overlap_fixed_seed', selected_kegg_terms, selected_go_terms)
etna_data = helper.load_kegg_go_data('consensus_ETNA_overlap_fixed_seed', selected_kegg_terms, selected_go_terms)
netmf_data_results = helper.generate_kegg_go_result(netmf_data, labels)
node2vec_data_results = helper.generate_kegg_go_result(node2vec_data, labels)
etna_data_results = helper.generate_kegg_go_result(etna_data, labels)

# %% [code]
netmf_data_results = pd.DataFrame(50-(np.array(netmf_data_results)).T,
                                  columns=['best_average',  'mean embedding',
                                           'corrected t-score', 'mean score'])

node2vec_data_results = pd.DataFrame(50-(np.array(node2vec_data_results)).T,
                                  columns=['best_average',  'mean embedding',
                                           'corrected t-score', 'mean score'])

etna_data_results = pd.DataFrame(50-(np.array(etna_data_results)).T,
                                  columns=['best_average', 'mean embedding',
                                           'corrected t-score', 'mean score' ])

# %% [code]
figure1a_df_y = list(node2vec_data_results['best_average'])+\
              list(node2vec_data_results['mean embedding'])+\
              list(node2vec_data_results['mean score'])+\
              list(node2vec_data_results['corrected t-score'])+\
              list(netmf_data_results['best_average'])+\
              list(netmf_data_results['mean embedding'])+\
              list(netmf_data_results['mean score'])+\
              list(netmf_data_results['corrected t-score'])+\
              list(etna_data_results['best_average'])+\
              list(etna_data_results['mean embedding'])+\
              list(etna_data_results['mean score'])+\
              list(etna_data_results['corrected t-score'])

figure1a_df_x = list([0 for i in range(node2vec_data_results.shape[0])])+\
                list([1 for i in range(node2vec_data_results.shape[0])])+\
                list([2 for i in range(node2vec_data_results.shape[0])])+\
                list([3 for i in range(node2vec_data_results.shape[0])])+\
                list([5 for i in range(netmf_data_results.shape[0])])+\
                list([6 for i in range(netmf_data_results.shape[0])])+\
                list([7 for i in range(netmf_data_results.shape[0])])+\
                list([8 for i in range(netmf_data_results.shape[0])])+\
                list([10 for i in range(etna_data_results.shape[0])])+\
                list([11 for i in range(etna_data_results.shape[0])])+\
                list([12 for i in range(etna_data_results.shape[0])])+\
                list([13 for i in range(etna_data_results.shape[0])])
figure1a_df = pd.DataFrame([figure1a_df_x, figure1a_df_y]).T
figure1a_df = figure1a_df.rename(columns={0:'x', 1:'y'})
figure1a_df['x'] = np.array(figure1a_df['x']).astype(int)

# %% [code]

import matplotlib.patches as mpatches
plt.figure(1,figsize=(12,4), dpi=200) 
figure1a_pal = {0:'#bf3f3f', 1:'#2e6b72', 2:'#274c77', 3:'#49a078', 4:'#ffffff',
                5:'#bf3f3f', 6:'#2e6b72', 7:'#274c77', 8:'#49a078', 9:'#ffffff',
                10:'#bf3f3f', 11:'#2e6b72', 12:'#274c77', 13:'#49a078', 14:'#ffffff',
                15:'#ffffff', 16:'#ffffff', 17:'#ffffff',18:'#ffffff'}
patch0 = mpatches.Patch(color='#bf3f3f', label='ANDES')
patch1 = mpatches.Patch(color='#2e6b72', label='mean embedding')
patch2 = mpatches.Patch(color='#274c77', label='mean score')
patch3 = mpatches.Patch(color='#49a078', label='corrected t-score')

a = sns.boxplot(x='x', y='y', data=figure1a_df, palette=figure1a_pal, order=np.arange(18))
a.invert_yaxis() 
a.set_xticks([1.5, 6.5, 11.5], labels=['node2vec', 'NetMF', 'NN'])
a.set_xlabel('')
a.set_ylabel('')
plt.ylabel('rank')
plt.legend(handles = [patch0,patch1,patch2,patch3], loc='lower right' )

# %% [markdown]
# Sfigure 2

# %% [code]
node2vec = helper.load_kegg_go_matrix_data('consensus_node2vec_fixed_seed', selected_kegg_terms, selected_go_terms)
heat01 = helper.load_kegg_go_matrix_data('heat_kernel_distinct_0.1_fixed_seed', selected_kegg_terms, selected_go_terms)
comm01 = helper.load_kegg_go_matrix_data('comm_kernel_distinct_0.1_fixed_seed', selected_kegg_terms, selected_go_terms)
jaccard = helper.load_kegg_go_matrix_data('consensus_jaccard_fixed_seed', selected_kegg_terms, selected_go_terms)
degree = helper.load_kegg_go_matrix_data('consensus_degree_fixed_seed', selected_kegg_terms, selected_go_terms)

node2vec_data_results = helper.generate_kegg_go_result(node2vec, labels)
heat01_data_results = helper.generate_kegg_go_result(heat01, labels)
comm01_data_results = helper.generate_kegg_go_result(comm01, labels)
jaccard_data_results = helper.generate_kegg_go_result(jaccard, labels)
degree_data_results = helper.generate_kegg_go_result(degree, labels)

# %% [code]
node2vec_data_results = pd.DataFrame(50-(np.array(node2vec_data_results)).T, 
                                     columns=['best_average',
                                               'corrected t-score', 'mean score' ])
heat01_data_results = pd.DataFrame(50-(np.array(heat01_data_results)).T, 
                                     columns=['best_average',
                                               'corrected t-score', 'mean score' ])
comm01_data_results = pd.DataFrame(50-(np.array(comm01_data_results)).T, 
                                     columns=['best_average',
                                               'corrected t-score', 'mean score' ])
jaccard_data_results = pd.DataFrame(50-(np.array(jaccard_data_results)).T, 
                                     columns=['best_average',
                                               'corrected t-score', 'mean score' ])
degree_data_results = pd.DataFrame(50-(np.array(degree_data_results)).T, 
                                     columns=['best_average',
                                               'corrected t-score', 'mean score' ])

# %% [code]
figure1c_df_y =list(node2vec_data_results['best_average'])+\
              list(node2vec_data_results['mean score'])+\
              list(node2vec_data_results['corrected t-score'])+\
              list(jaccard_data_results['best_average'])+\
              list(jaccard_data_results['mean score'])+\
              list(jaccard_data_results['corrected t-score'])+\
              list(heat01_data_results['best_average'])+\
              list(heat01_data_results['mean score'])+\
              list(heat01_data_results['corrected t-score'])+\
              list(comm01_data_results['best_average'])+\
              list(comm01_data_results['mean score'])+\
              list(comm01_data_results['corrected t-score'])+\
              list(degree_data_results['best_average'])+\
              list(degree_data_results['mean score'])+\
              list(degree_data_results['corrected t-score'])

figure1c_df_x = list([0 for i in range(node2vec_data_results.shape[0])])+\
                list([1 for i in range(node2vec_data_results.shape[0])])+\
                list([2 for i in range(node2vec_data_results.shape[0])])+\
                list([4 for i in range(heat01_data_results.shape[0])])+\
                list([5 for i in range(heat01_data_results.shape[0])])+\
                list([6 for i in range(heat01_data_results.shape[0])])+\
                list([8 for i in range(comm01_data_results.shape[0])])+\
                list([9 for i in range(comm01_data_results.shape[0])])+\
                list([10 for i in range(comm01_data_results.shape[0])])+\
                list([12 for i in range(jaccard_data_results.shape[0])])+\
                list([13 for i in range(jaccard_data_results.shape[0])])+\
                list([14 for i in range(jaccard_data_results.shape[0])])+\
                list([16 for i in range(degree_data_results.shape[0])])+\
                list([17 for i in range(degree_data_results.shape[0])])+\
                list([18 for i in range(degree_data_results.shape[0])])
                
figure1c_df = pd.DataFrame([figure1c_df_x, figure1c_df_y]).T
figure1c_df = figure1c_df.rename(columns={0:'x', 1:'y'})
figure1c_df['x'] = np.array(figure1c_df['x']).astype(int)

# %% [code]
import matplotlib.patches as mpatches
plt.figure(1,figsize=(12,4), dpi=200) 
figure1c_pal = {0:'#bf3f3f', 1:'#274c77', 2:'#49a078', 3:'#ffffff',
                4:'#bf3f3f', 5:'#274c77', 6:'#49a078', 7:'#ffffff',
                8:'#bf3f3f', 9:'#274c77', 10:'#49a078', 11:'#ffffff',
                12:'#bf3f3f', 13:'#274c77', 14:'#49a078', 15:'#ffffff',
                16:'#bf3f3f', 17:'#274c77', 18:'#49a078', 19:'#ffffff'}
b = sns.boxplot(x='x', y='y', data=figure1c_df, palette=figure1c_pal, order=np.arange(4,19))
b.invert_yaxis() 
b.set_xlim(-1,19)
b.set_xticks([1, 5, 9, 13], labels=[ 'jaccard', 'heat diffusion',  'exp diffusion', 'degree'])
# b.set_xticks([1, 5, 9, 13, 17], labels=['node2vec', 'heat diffus', 
#                                         'exp diffus', 'jaccard', 'degree'])
b.set_xlabel('')
b.set_ylabel('rank')

patch0 = mpatches.Patch(color='#bf3f3f', label='ANDES')
patch1 = mpatches.Patch(color='#2e6b72', label='mean embedding')
patch2 = mpatches.Patch(color='#274c77', label='mean score')
patch3 = mpatches.Patch(color='#49a078', label='corrected t-score')

plt.legend(handles = [patch0,patch2,patch3], loc='lower right' )

# %% [markdown]
# Sfigure 42

# %% [code]
top_k_results = helper.load_kegg_go_k_data('node2vec', selected_kegg_terms, selected_go_terms, ['5','10','20','50'])
top_k_results = helper.generate_kegg_go_result(top_k_results, labels)
top_k_results = pd.DataFrame(50-(np.array(top_k_results)).T, columns=['5', '10','20','50'])

# supplement figure
figure1d_df_y =list(node2vec_data_results['best_average'])+\
              list(top_k_results['5'])+\
              list(top_k_results['10'])+\
              list(top_k_results['20'])+\
              list(top_k_results['50'])+\
              list(node2vec_data_results['mean score'])
              

figure1d_df_x = list([0 for i in range(node2vec_data_results.shape[0])])+\
                list([1 for i in range(top_k_results.shape[0])])+\
                list([2 for i in range(top_k_results.shape[0])])+\
                list([3 for i in range(top_k_results.shape[0])])+\
                list([4 for i in range(top_k_results.shape[0])])+\
                list([5 for i in range(node2vec_data_results.shape[0])])
                
figure1d_df = pd.DataFrame([figure1d_df_x, figure1d_df_y]).T
figure1d_df = figure1d_df.rename(columns={0:'x', 1:'y'})
figure1d_df['x'] = np.array(figure1d_df['x']).astype(int)

plt.figure(1,figsize=(12,4), dpi=200) 
figure1d_pal = {0:'#bf3f3f', 1:'#a1414b', 2:'#82414a', 3:'#644455',
                4:'#46496b', 5:'#274c77'}
d = sns.boxplot(x='x', y='y', data=figure1d_df, palette=figure1d_pal, order=np.arange(0,6))
d.invert_yaxis() 
d.set_xlim(-0.5,5.5)
d.set_xticks([0,1,2,3,4,5], labels=['top 1 (ANDES)', 'k=5%', 'k=10%', 'k=20%', 'k=50%', 'k=100% (mean score)'])
 
d.set_xlabel('')
d.set_ylabel('rank')

# %% [markdown]
# Sfigure 43

# %% [code]
cosine = helper.load_kegg_go_data('consensus_node2vec_fixed_seed', selected_kegg_terms, selected_go_terms)
euclidean = helper.load_kegg_go_data('consensus_node2vec_euclidean_fixed_seed', selected_kegg_terms, selected_go_terms)
pearson = helper.load_kegg_go_data('consensus_node2vec_correlation_fixed_seed', selected_kegg_terms, selected_go_terms)

cosine_data_results = helper.generate_kegg_go_result(cosine, labels)
euclidean_data_results = helper.generate_kegg_go_result(euclidean, labels)
pearson_data_results = helper.generate_kegg_go_result(pearson, labels)

cosine_data_results = pd.DataFrame(50-(np.array(cosine_data_results)).T,
                                      columns=['best_average',  'mean embedding',
                                               'corrected t-score', 'mean score' ])
euclidean_data_results = pd.DataFrame(50-(np.array(euclidean_data_results)).T,
                                      columns=['best_average',  'mean embedding',
                                               'corrected t-score', 'mean score' ])
pearson_data_results = pd.DataFrame(50-(np.array(pearson_data_results)).T,
                                      columns=['best_average',  'mean embedding',
                                               'corrected t-score', 'mean score' ])

# %% [code]
figure1c_df_y =list(cosine_data_results['best_average'])+\
              list(cosine_data_results['mean embedding'])+\
              list(cosine_data_results['mean score'])+\
              list(cosine_data_results['corrected t-score'])+\
              list(euclidean_data_results['best_average'])+\
              list(euclidean_data_results['mean embedding'])+\
              list(euclidean_data_results['mean score'])+\
              list(euclidean_data_results['corrected t-score'])+\
              list(pearson_data_results['best_average'])+\
              list(pearson_data_results['mean embedding'])+\
              list(pearson_data_results['mean score'])+\
              list(pearson_data_results['corrected t-score'])

figure1c_df_x = list([0 for i in range(cosine_data_results.shape[0])])+\
                list([1 for i in range(cosine_data_results.shape[0])])+\
                list([2 for i in range(cosine_data_results.shape[0])])+\
                list([3 for i in range(cosine_data_results.shape[0])])+\
                list([5 for i in range(euclidean_data_results.shape[0])])+\
                list([6 for i in range(euclidean_data_results.shape[0])])+\
                list([7 for i in range(euclidean_data_results.shape[0])])+\
                list([8 for i in range(euclidean_data_results.shape[0])])+\
                list([10 for i in range(pearson_data_results.shape[0])])+\
                list([11 for i in range(pearson_data_results.shape[0])])+\
                list([12 for i in range(pearson_data_results.shape[0])])+\
                list([13 for i in range(pearson_data_results.shape[0])])
                
figure1c_df = pd.DataFrame([figure1c_df_x, figure1c_df_y]).T
figure1c_df = figure1c_df.rename(columns={0:'x', 1:'y'})
figure1c_df['x'] = np.array(figure1c_df['x']).astype(int)

# %% [code]
import matplotlib.patches as mpatches
plt.figure(1,figsize=(12,4), dpi=200) 
figure1c_pal = {0:'#bf3f3f', 1:'#2e6b72', 2:'#274c77', 3:'#49a078', 4:'#ffffff',
                5:'#bf3f3f', 6:'#2e6b72', 7:'#274c77', 8:'#49a078', 9:'#ffffff',
                10:'#bf3f3f', 11:'#2e6b72', 12:'#274c77', 13:'#49a078', 14:'#ffffff',
                15:'#ffffff', 16:'#ffffff', 17:'#ffffff',18:'#ffffff'}
b = sns.boxplot(x='x', y='y', data=figure1c_df, palette=figure1c_pal, order=np.arange(18))
b.invert_yaxis() 
# b.set_xlim(-1,11)
b.set_xticks([1.5, 6.5, 11.5], labels=['Cosine Similairty', 'Euclidean Distance (inverse)', 'Pearson Correlation'])
b.set_xlabel('')
b.set_ylabel('rank')


patch0 = mpatches.Patch(color='#bf3f3f', label='ANDES')
patch1 = mpatches.Patch(color='#2e6b72', label='mean embedding')
patch2 = mpatches.Patch(color='#274c77', label='mean score')
patch3 = mpatches.Patch(color='#49a078', label='corrected t-score')
plt.legend(handles = [patch0,patch1,patch2,patch3], loc='lower right' )

# %% [code]
```
