## Supplemental Code for "ANDES: a novel best-match approach for enhancing gene set analysis in embedding spaces": model_organism_analysis.ipynb

```
# %% [code]
import pandas as pd
import numpy as np
import sklearn.metrics as metrics
import networkx as nx
import obonet
import random
import pickle

from collections import defaultdict

import matplotlib.pyplot as plt
import seaborn as sns
from matplotlib.pyplot import figure

import helper as helper


# %% [code]
org1='hsa'
org2='mmu'

# %% [code]
results, shared_terms = helper.load_model_org_data(org1, org2)
ranks = helper.get_model_org_rank(results, axis=0)
ranks = np.array(ranks)

# %% [code]
figure2a_y = ranks.flatten()/ranks.shape[1]
figure2a_x = list([0 for i in range(ranks.shape[1])])+\
                list([1 for i in range(ranks.shape[1])])+\
                list([2 for i in range(ranks.shape[1])])+\
                list([3 for i in range(ranks.shape[1])])
figure2a_df = pd.DataFrame([figure2a_x, figure2a_y]).T
figure2a_df = figure2a_df.rename(columns={0:'x', 1:'y'})
figure2a_df['x'] = np.array(figure2a_df['x']).astype(int)

# %% [code]
import matplotlib.patches as mpatches
plt.figure(1,figsize=(10,8), dpi=200) 
figure2a_pal = {0:'#bf3f3f', 1:'#2e6b72', 2:'#274c77', 3:'#49a078', 4:'#ffffff', 5:'#ffffff', 6:'#ffffff'}
patch0 = mpatches.Patch(color='#bf3f3f', label='ANDES')
patch1 = mpatches.Patch(color='#2e6b72', label='mean embedding')
patch2 = mpatches.Patch(color='#274c77', label='mean score')
patch3 = mpatches.Patch(color='#49a078', label='corrected t-score')
                
a = sns.boxplot(x='x', y='y', data=figure2a_df, palette=figure2a_pal, order=np.arange(6))
a.invert_yaxis() 
a.set_xticks([1.5], labels=[org1+'_'+org2])
a.set_xlabel('')
a.set_ylabel('normalized rank')
plt.legend(handles = [patch0,patch1,patch2,patch3], loc='lower right' )

# %% [code]
resnik_scores = pd.read_csv('./data/go_resnik_score.csv', index_col=0)
resnik_scores = resnik_scores[shared_terms].loc[shared_terms].values

# %% [code]
cum_resnik_scores = [resnik_scores.flatten()[list(np.argsort(x.flatten()))[::-1]] for x in results]
cum_resnik_scores = [np.cumsum(x)/np.arange(1,len(x)+1) for x in cum_resnik_scores]

# %% [code]
cum_resnik_score_size = len(cum_resnik_scores[0])
plot_indices = [i for i in range(20, cum_resnik_score_size//3, 100)]

figure(figsize=(9, 6), dpi=200)
plt.plot(np.array(plot_indices)/cum_resnik_score_size, cum_resnik_scores[0][plot_indices], 
         color='#bf3f3f', linewidth=2, label='ANDES')
plt.plot(np.array(plot_indices)/cum_resnik_score_size, cum_resnik_scores[1][plot_indices], 
         color='#49a078', linestyle='dashdot', linewidth=2, label='corrected t-score')
plt.plot(np.array(plot_indices)/cum_resnik_score_size, cum_resnik_scores[2][plot_indices], 
         color='#2e6b72', linestyle='dotted', linewidth=2, label='mean embedding')
plt.plot(np.array(plot_indices)/cum_resnik_score_size, cum_resnik_scores[3][plot_indices], 
         color='#274c77', linestyle='dashed', linewidth=2, label='mean score')


plt.xticks([0, 0.05, 0.10, 0.15, 0.20, 0.25, 0.30], 
             labels=['0', '5', '10', '15', '20', '25', '30'], fontsize=15)
plt.yticks([0.15,  0.25,  0.35,  0.45, 0.55], fontsize=15)
plt.legend(prop={'size': 15})
plt.ylabel('normalized resnik\'s score', fontsize=15)
plt.xlabel('percentage', fontsize=15)


# %% [code]
```
