## Supplemental Code for "ANDES: a novel best-match approach for enhancing gene set analysis in embedding spaces": mouse_pheno.ipynb

```
# %% [code]
import pandas as pd
import numpy as np
import obonet
import networkx as nx

from collections import defaultdict
from sklearn import metrics
import scipy.stats as stats
import matplotlib.pyplot as plt
import seaborn as sns
from matplotlib.pyplot import figure
import scipy

import helper as helper


# %% [code]
mouse_pheno = pd.read_csv('./results/mouse_pheno/p0_fixed_seed.csv', sep=',', index_col=0)

# %% [code]
omim_terms = list(mouse_pheno.index)
omim_term2name = helper.term2name('./data/gmt/omim.20231030.gmt')
obo_graph = obonet.read_obo('./data/ontology/doid.obo')
omim_graph = nx.DiGraph()
for x,y in obo_graph.edges():
    omim_graph.add_edge(y, x)
    
for term, name in obo_graph.nodes(data='name'):
    omim_term2name[term] = name
ordered_omim_nodes = list(nx.dfs_preorder_nodes(omim_graph, 'DOID:4'))
ordered_omim_terms = [x for x in ordered_omim_nodes if x in omim_terms]
print(len(ordered_omim_terms), len(omim_terms))
omim_indices = [omim_terms.index(x) for x in ordered_omim_terms]
ordered_omim_names = [omim_term2name[x] for x in ordered_omim_terms]

omim_name2term = dict()
for key, value in omim_term2name.items():
    omim_name2term[value]=key

# %% [code]
mmu_terms = list(mouse_pheno.columns)
mmu_term2name = helper.term2name('./data/gmt/mouse_pheno.202203.gmt')
obo_graph = obonet.read_obo('./data/ontology/MPheno_OBO.ontology')
mmu_graph = nx.DiGraph()
for x,y in obo_graph.edges():
    mmu_graph.add_edge(y, x)

for term, name in obo_graph.nodes(data='name'):
    mmu_term2name[term] = name
ordered_mmu_nodes = list(nx.dfs_preorder_nodes(mmu_graph, 'MP:0000001'))
ordered_mmu_terms = [x for x in ordered_mmu_nodes if x in mmu_terms]
print(len(ordered_mmu_terms), len(mmu_terms))
mmu_indices = [mmu_terms.index(x) for x in ordered_mmu_terms]
ordered_mmu_names = [mmu_term2name[x] for x in ordered_mmu_terms]

mmu_name2term = dict()
for key, value in mmu_term2name.items():
    mmu_name2term[value]=key

# %% [code]
disease_slim_term = []
with open('./data/drug_info/DO_slim.txt', 'r') as f:
    for line in f:
        tokens = line.split('\t')
        disease_slim_term.append(tokens[0])

# %% [code]
omim2slim = defaultdict(list)
omim_slim2terms = defaultdict(list)
omim_slim_terms = set()
for x in ordered_omim_terms:
    paths = helper.paths_to_root(omim_graph, x, 'DOID:4')
    mapped_slim = set()
    for p in paths:
        mapped_slim = mapped_slim.union(set(disease_slim_term).intersection(p))
        
    if len(mapped_slim) == 0:
        mapped_slim = set([y[3] for y in helper.paths_to_root(omim_graph, x, 'DOID:4') if len(y)>3])
        
    omim_slim_terms = omim_slim_terms.union(mapped_slim)
    if len(mapped_slim)>0:
        omim2slim[x] = list(mapped_slim)
    for y in mapped_slim:
        omim_slim2terms[y].append(x)

# %% [code]
ordered_omim_slim_terms = [x for x in ordered_omim_nodes if x in omim_slim_terms]
for x in ordered_omim_slim_terms:
    # if 'cancer' in omim_term2name[x]:
    #     continue
    print(x, omim_term2name[x])
    
    sub_matrix = helper.get_sub_matrix_single_selected(x, mouse_pheno, omim_slim2terms, 
                                                ordered_omim_nodes, ordered_mmu_nodes, k=10)
    sub_matrix = sub_matrix.loc[:,((sub_matrix>1.64).sum(axis=0) != 0)]
    

    if sub_matrix.shape[0]<5:
        continue
    if sub_matrix.shape[1]<5:
        continue
        
    print(sub_matrix.shape)

    a_fig = sns.clustermap(sub_matrix.values, 
                yticklabels=[omim_term2name[x] for x in sub_matrix.index], 
                xticklabels=[mmu_term2name[x] for x in sub_matrix.columns], 
                figsize=(20,10), linewidths=0.5, linecolor='grey', 
                mask=1-(sub_matrix.values>1.64), cmap='rocket_r',
                           cbar_pos=(0.1, .75, .02, .2))
    row = a_fig.ax_row_dendrogram.get_position()
    col = a_fig.ax_col_dendrogram.get_position()
    print(a_fig.ax_col_dendrogram.get_position())
    a_fig.ax_row_dendrogram.set_position([row.x0+row.width*0.8, row.y0, row.width*0.2, row.height])
    # a_fig.ax_col_dendrogram.set_position([col.x0, col.y0, col.width, 0.1])
    print(a_fig.ax_col_dendrogram.get_position())
    a_fig.ax_heatmap.set_xticklabels(a_fig.ax_heatmap.get_xmajorticklabels(), fontsize = 15)
    a_fig.ax_heatmap.set_yticklabels(a_fig.ax_heatmap.get_ymajorticklabels(), fontsize = 15)
    
    # plt.tight_layout()
    plt.show()
    plt.close()
    # break
    

# %% [code]
```
