## Supplemental Code for "ANDES: a novel best-match approach for enhancing gene set analysis in embedding spaces": README.md

# ANDES: Algorithm for Network Data Embedding and Similarity analysis
This repository contains the scripts to run the ANDES method and corresponding analysis.

## About
Embedding methods have emerged as a valuable class of approaches for distilling essential information from complex high-dimensional data into more accessible lower-dimensional spaces. Applications of embedding methods to biological data have demonstrated that gene embeddings can effectively capture physical, structural, and functional relationships between genes. This utility has largely been demonstrated by using gene embeddings for downstream machine learning tasks. Much less has been done to examine the embeddings directly. Limited efforts towards comparing gene sets typically opt to compare simple mean embeddings between sets.
Here, we propose a novel best-match approach that considers gene similarity while reconciling gene set diversity. We demonstrate that our method can better represent gene set similarity compared to existing methods in both single-species and cross-species settings. In addition, by employing our best-match concept on a gene embedding space made from protein-protein interactions, we developed a novel rank-based gene set enrichment analysis method that achieves state-of-the-art performance.

ANDES's method has two main functions:

  1. calculating gene sets similarity in a embedding space
  2. ranked-based GSEA using gene embedding information

These functions are implemented in `src/set_analysis_fun.py`. The demo
jupyter notebook (`demo.ipynb`) illustrates how to use these two funcions


## Usage
This project uses conda to manage the required packages and setup a virtual environment. Once conda is installed on your machine get started by setting up the virtual environment.

```sh
conda env create -f env.yml
conda activate ANDES
```

ANDES can be run through the command line as follows:

```sh
python src/andes.py --emb embedding_file.csv --genelist embedding_gene_ids.txt --geneset1 first_gene_set_database.gmt --geneset2 second_gene_set_database.gmt --out output_file.csv -n num_processor
```

ANDES performing GSEA can be run through the command line as follows:

```sh
python src/andes_gsea.py --emb embedding_file.csv --genelist embedding_gene_ids.txt --geneset gene_set_database.gmt --rankedlist ranked_genes.txt --out output_file.csv -n num_processor
```
