## Supplementary figures and images for "ANDES: a novel best-match approach for enhancing gene set analysis in embedding spaces"

### hsa00010.pdf

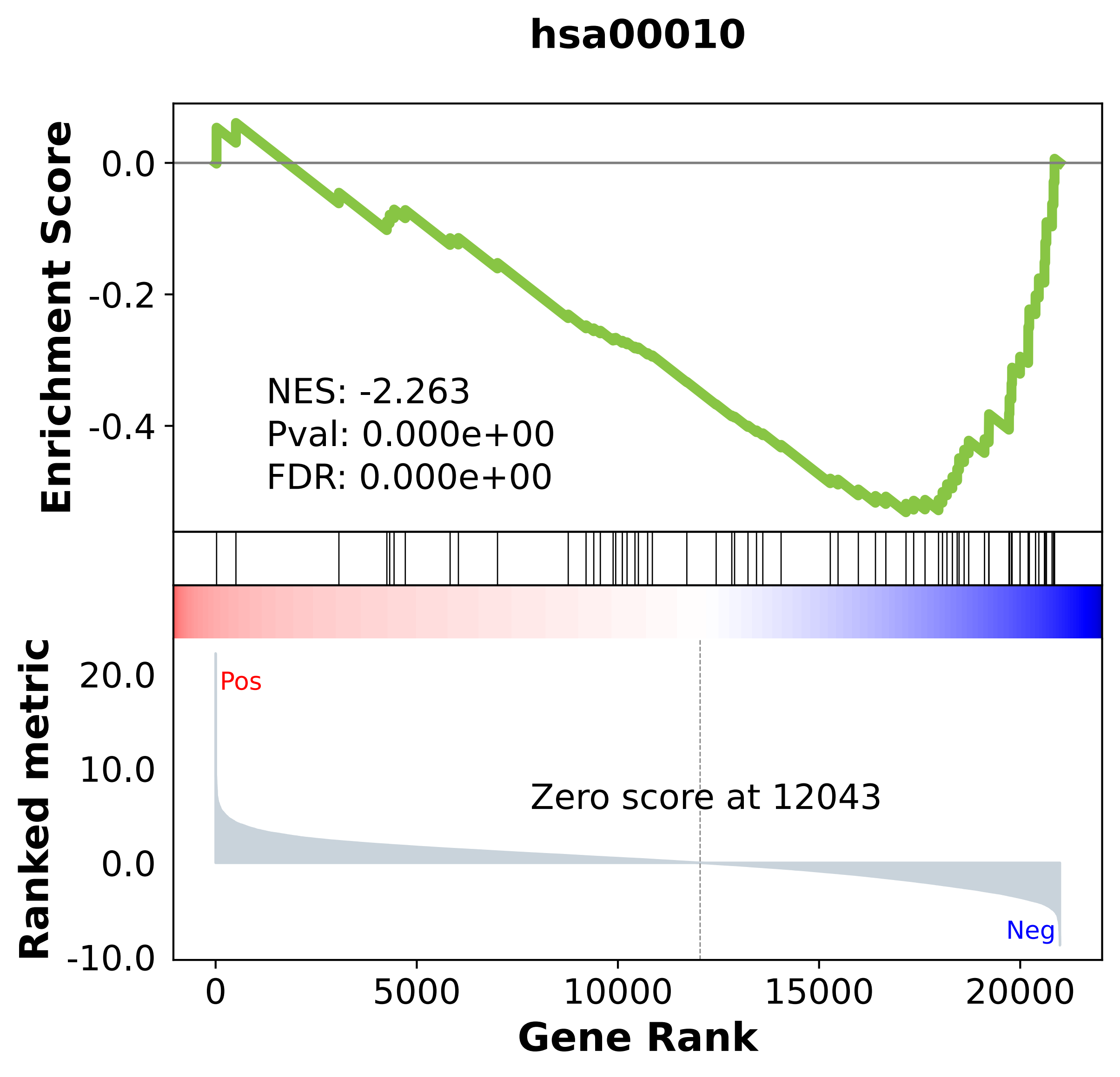

### hsa00020.pdf

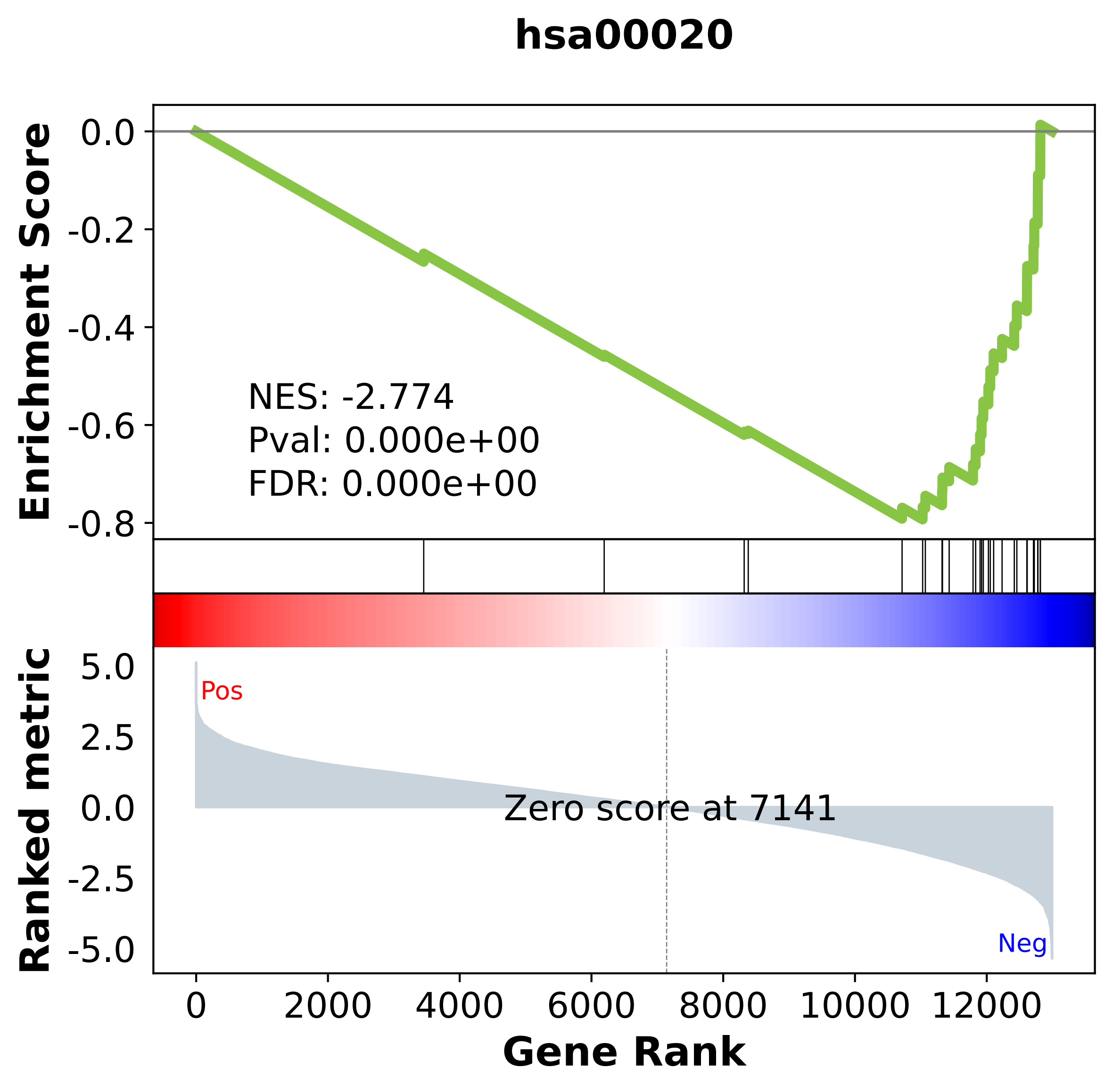

### hsa00030.pdf

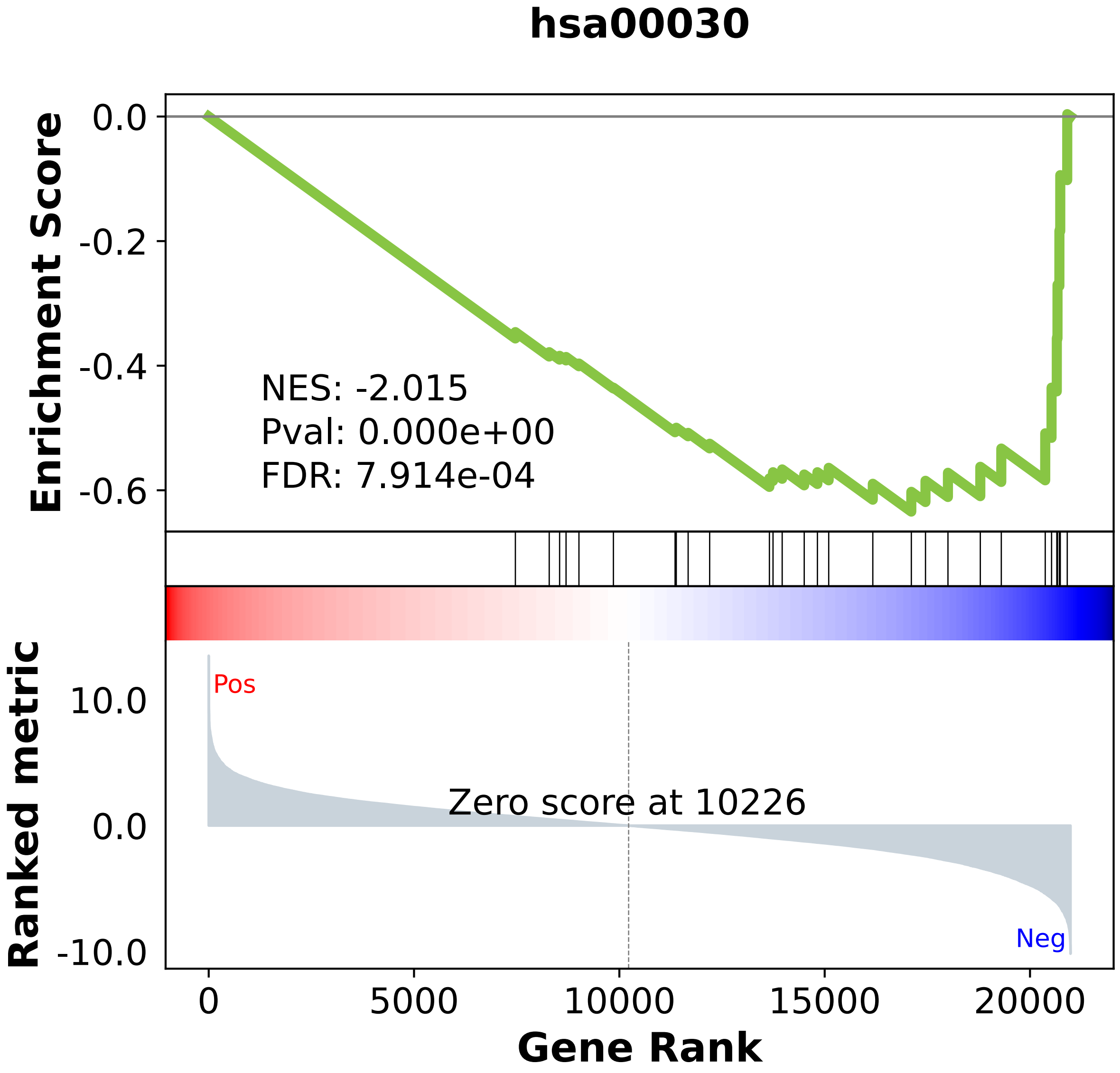

### hsa00040.pdf

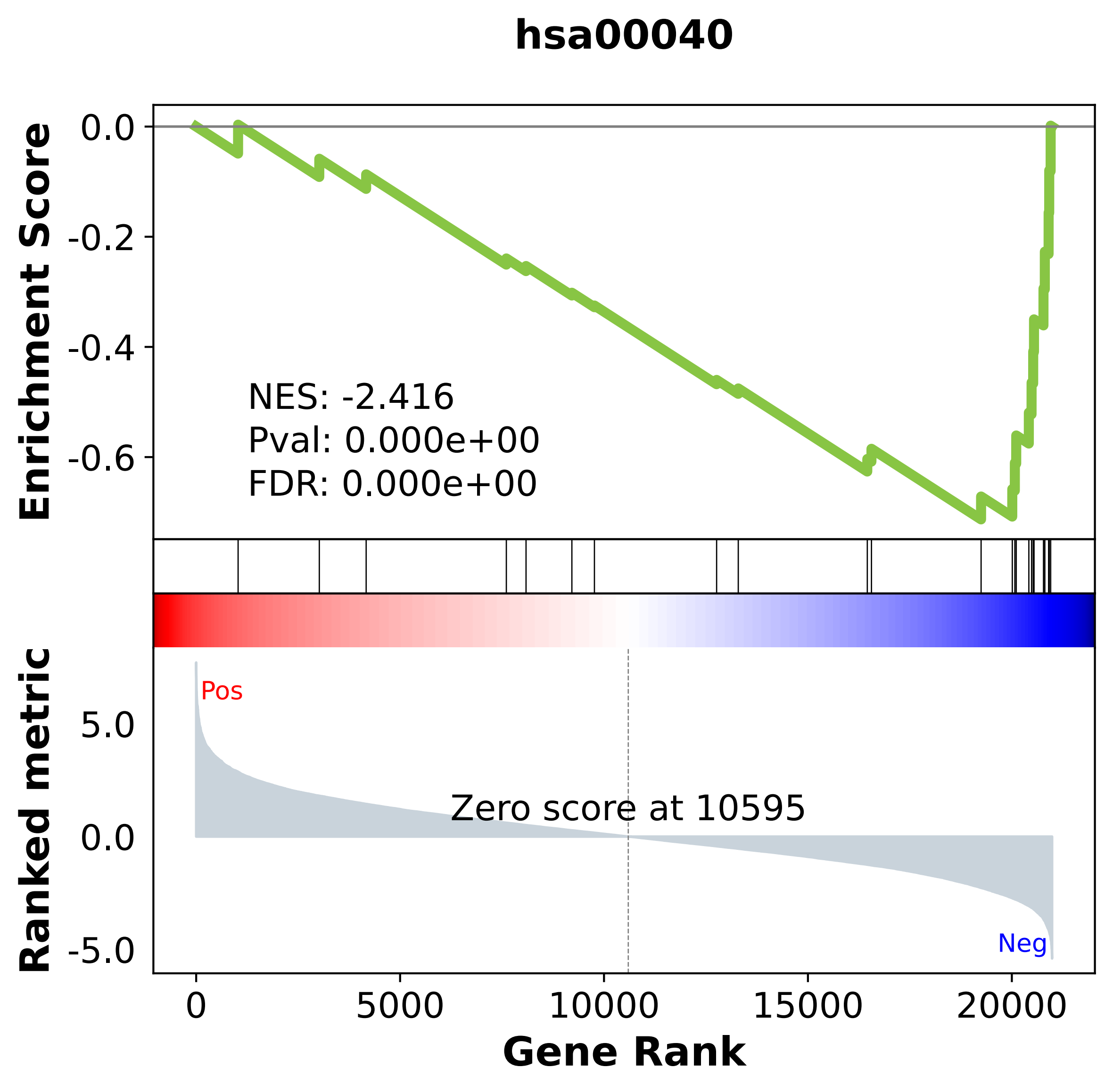

### hsa00051.pdf

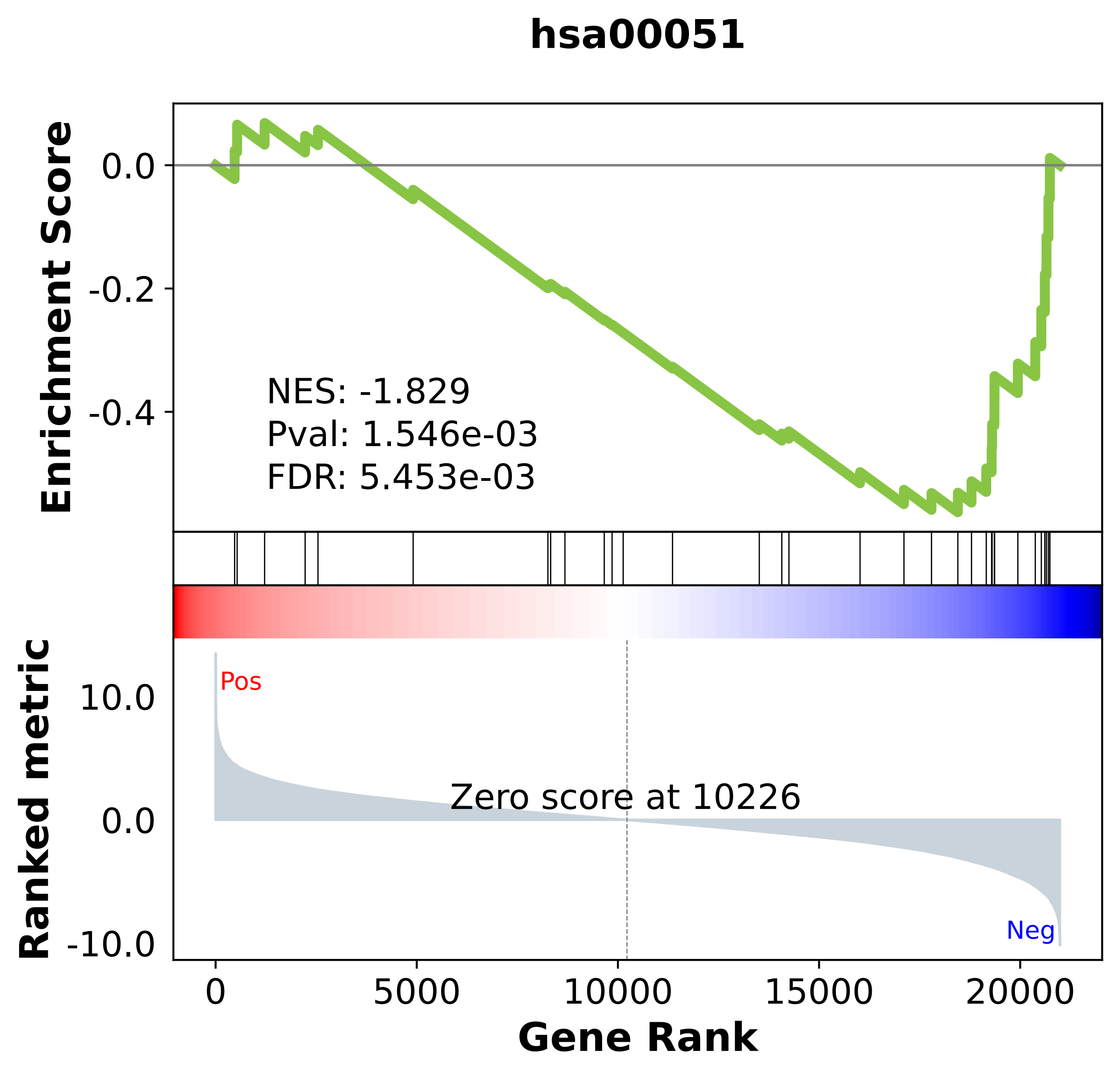

### hsa00052.pdf

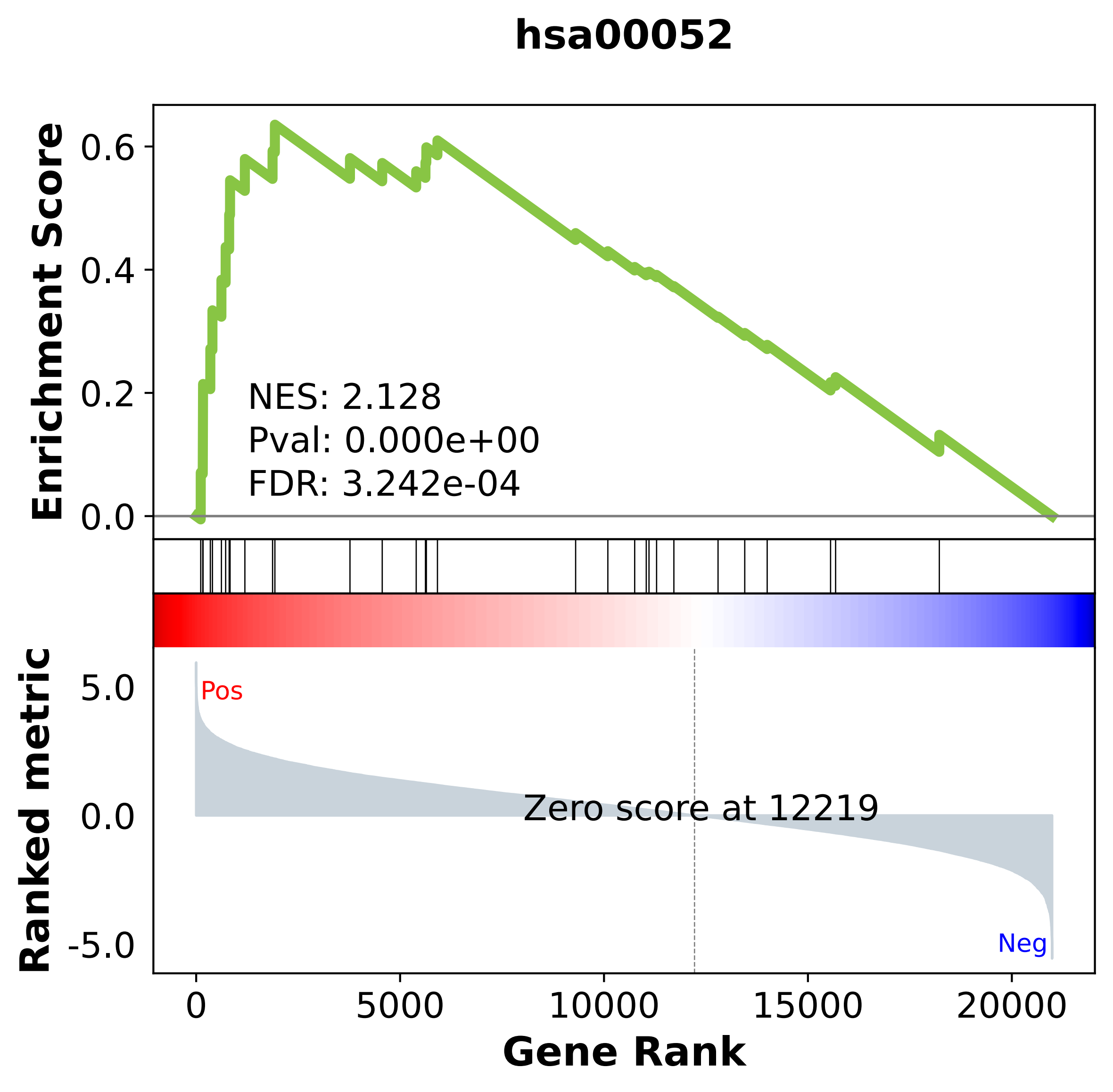

### hsa00053.pdf

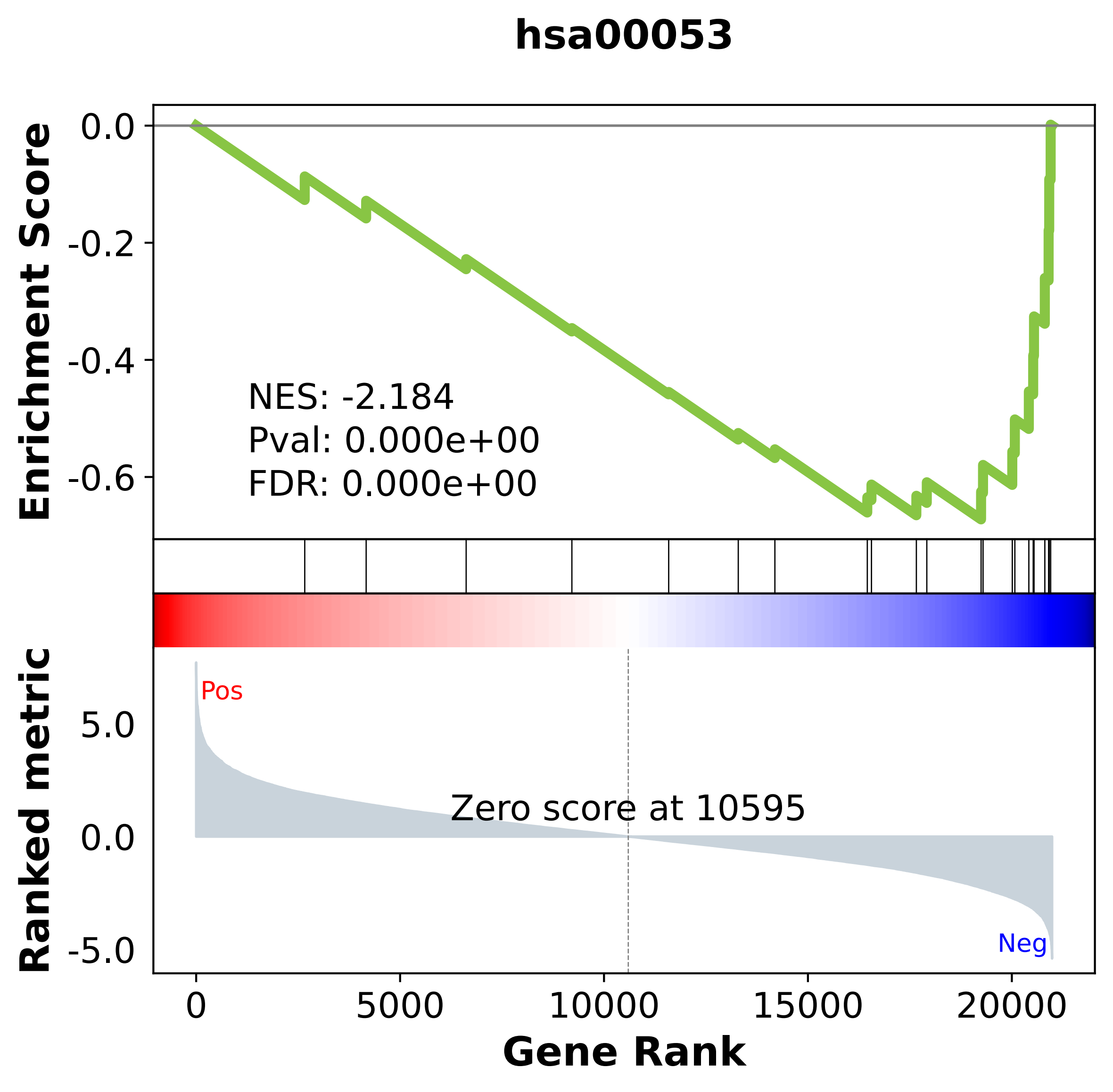

### hsa00062.pdf

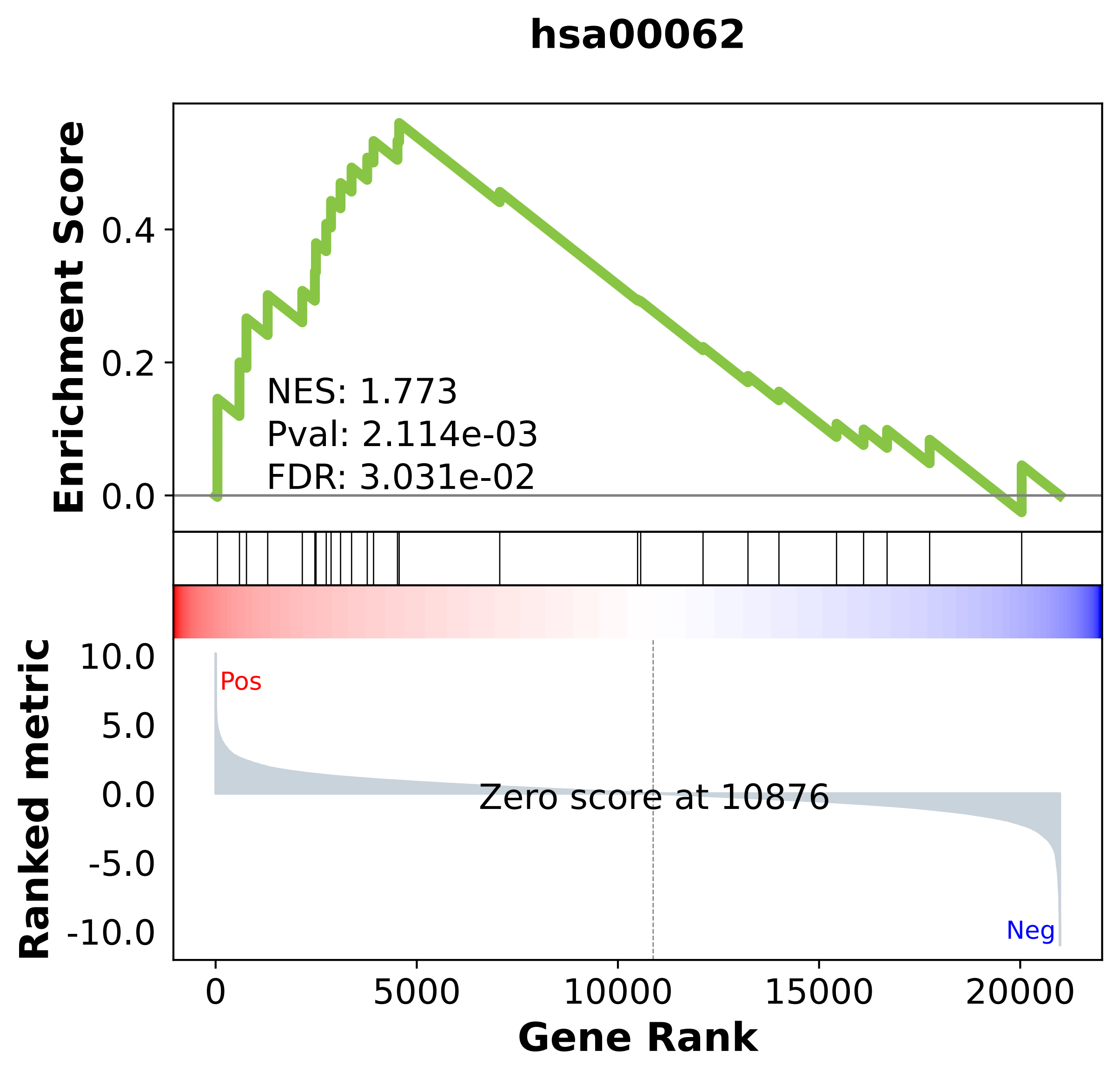

### hsa00071.pdf

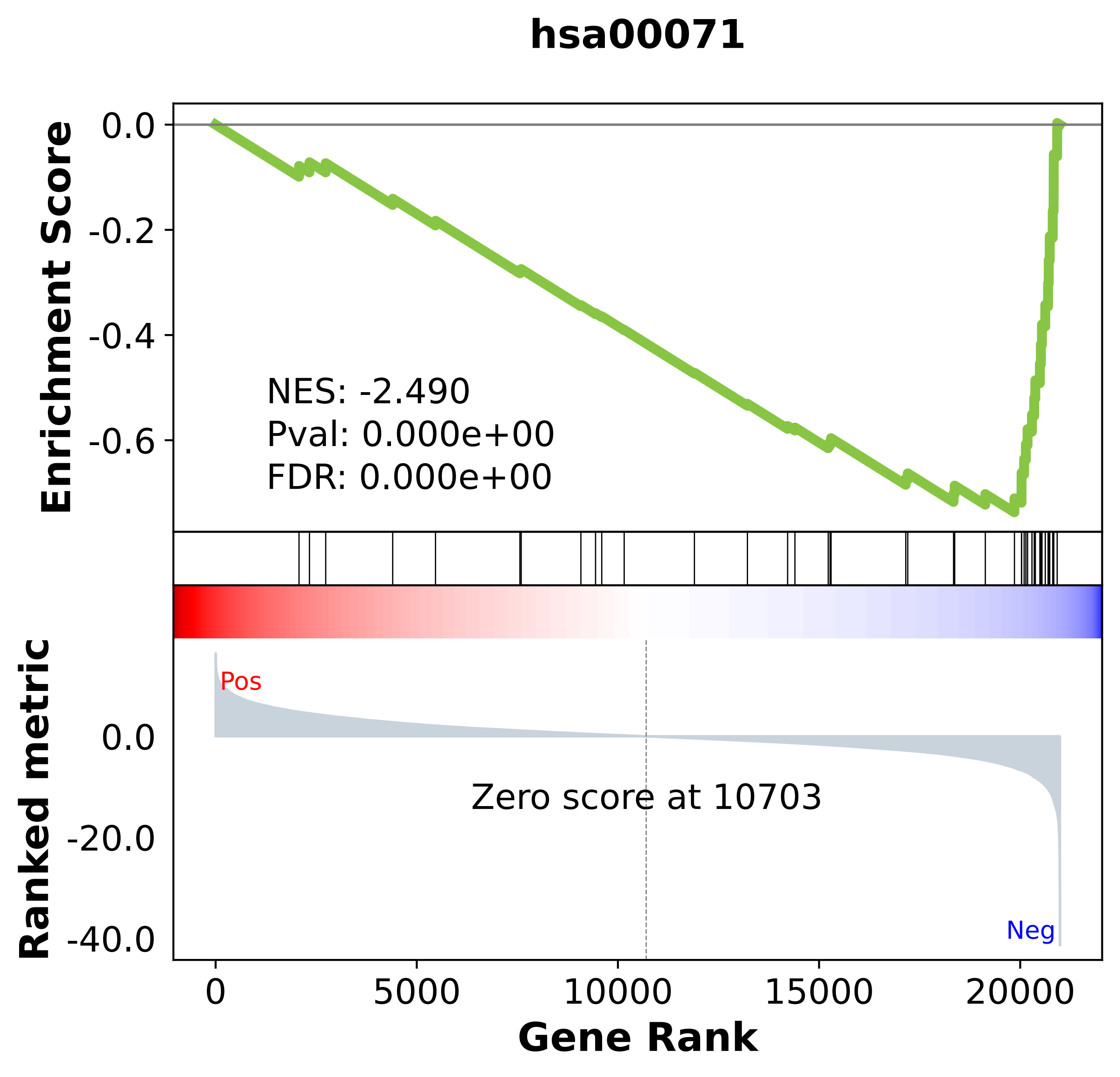

### hsa00100.pdf

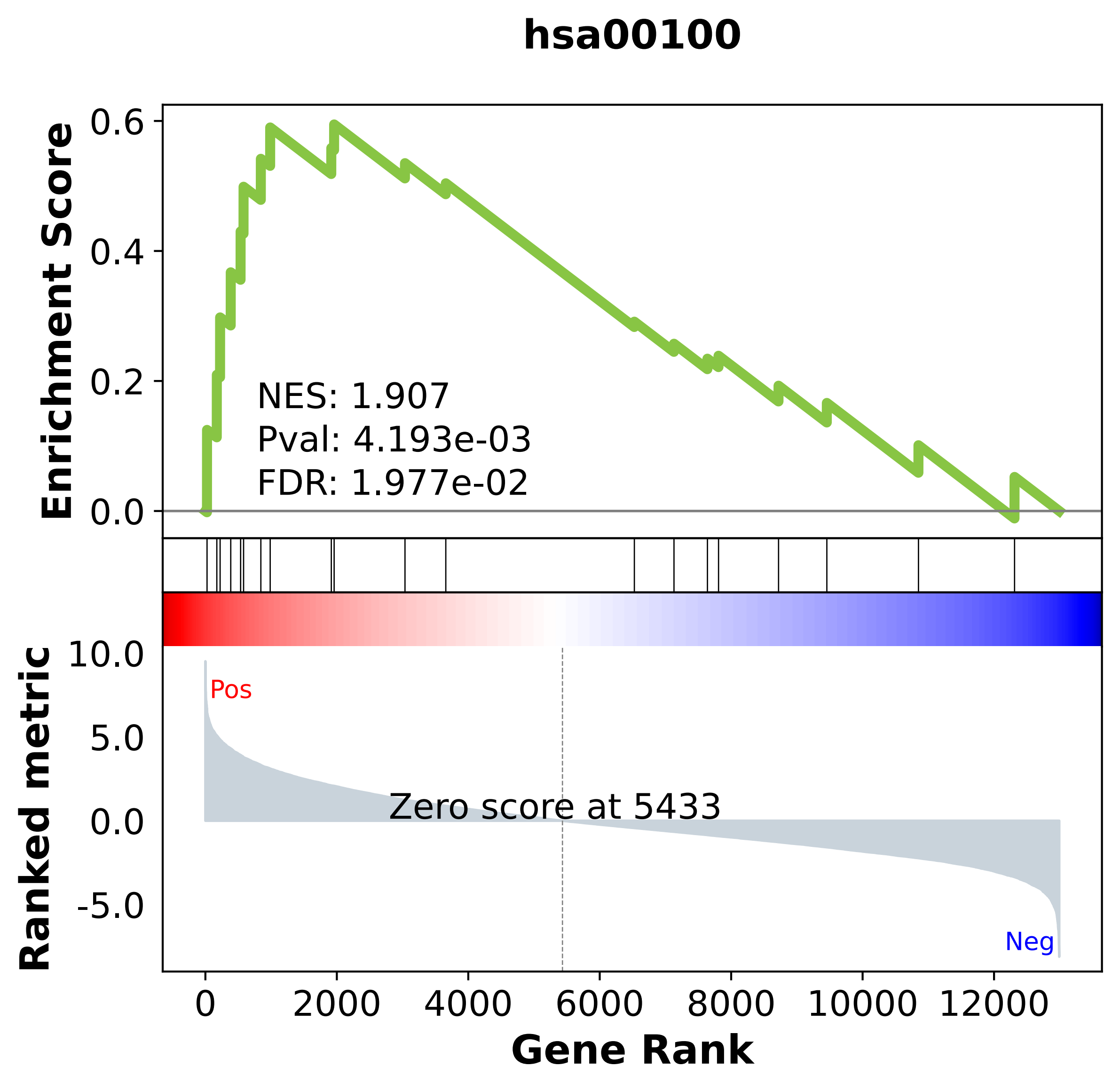

### hsa00120.pdf

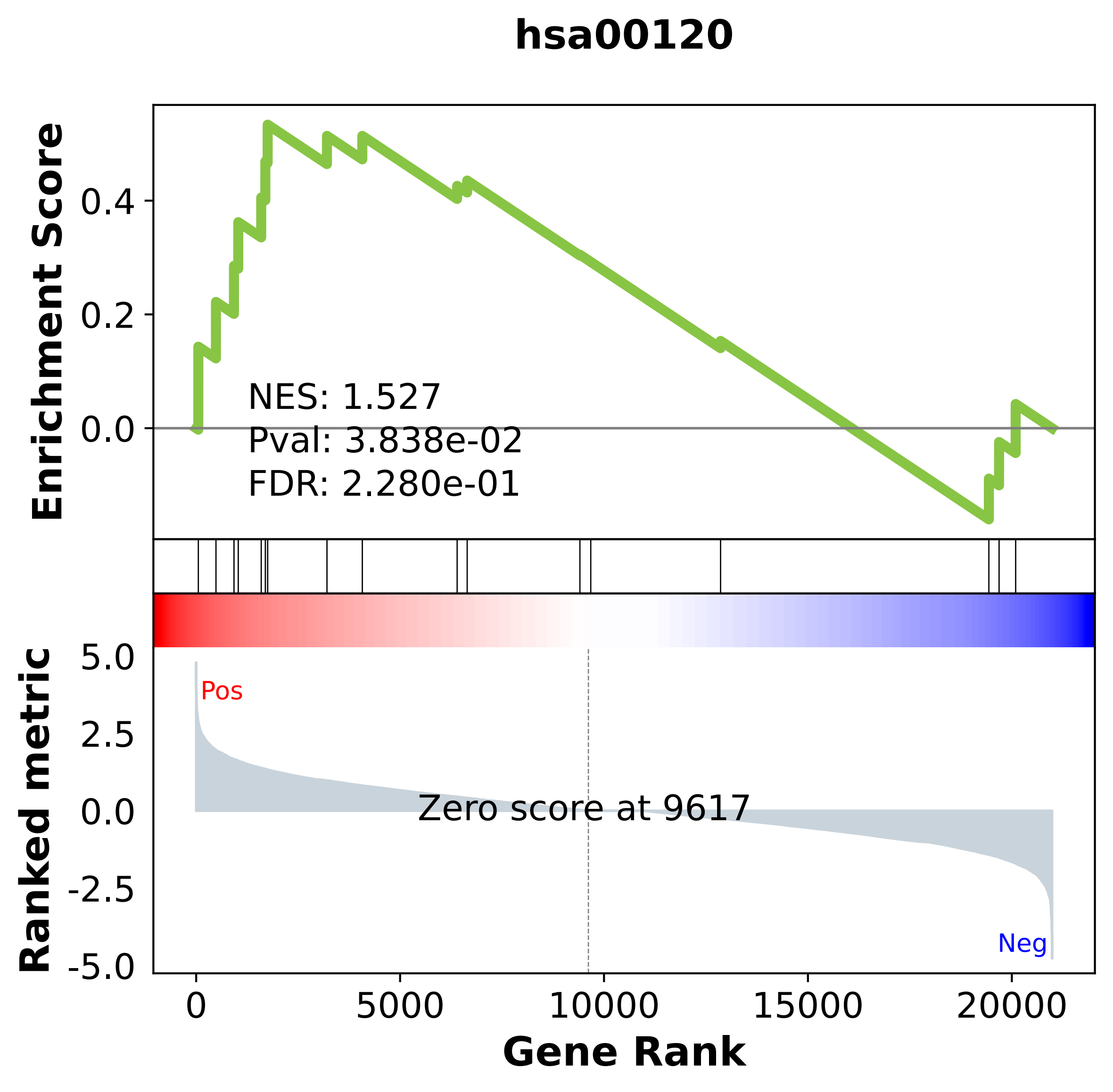

### hsa00140.pdf

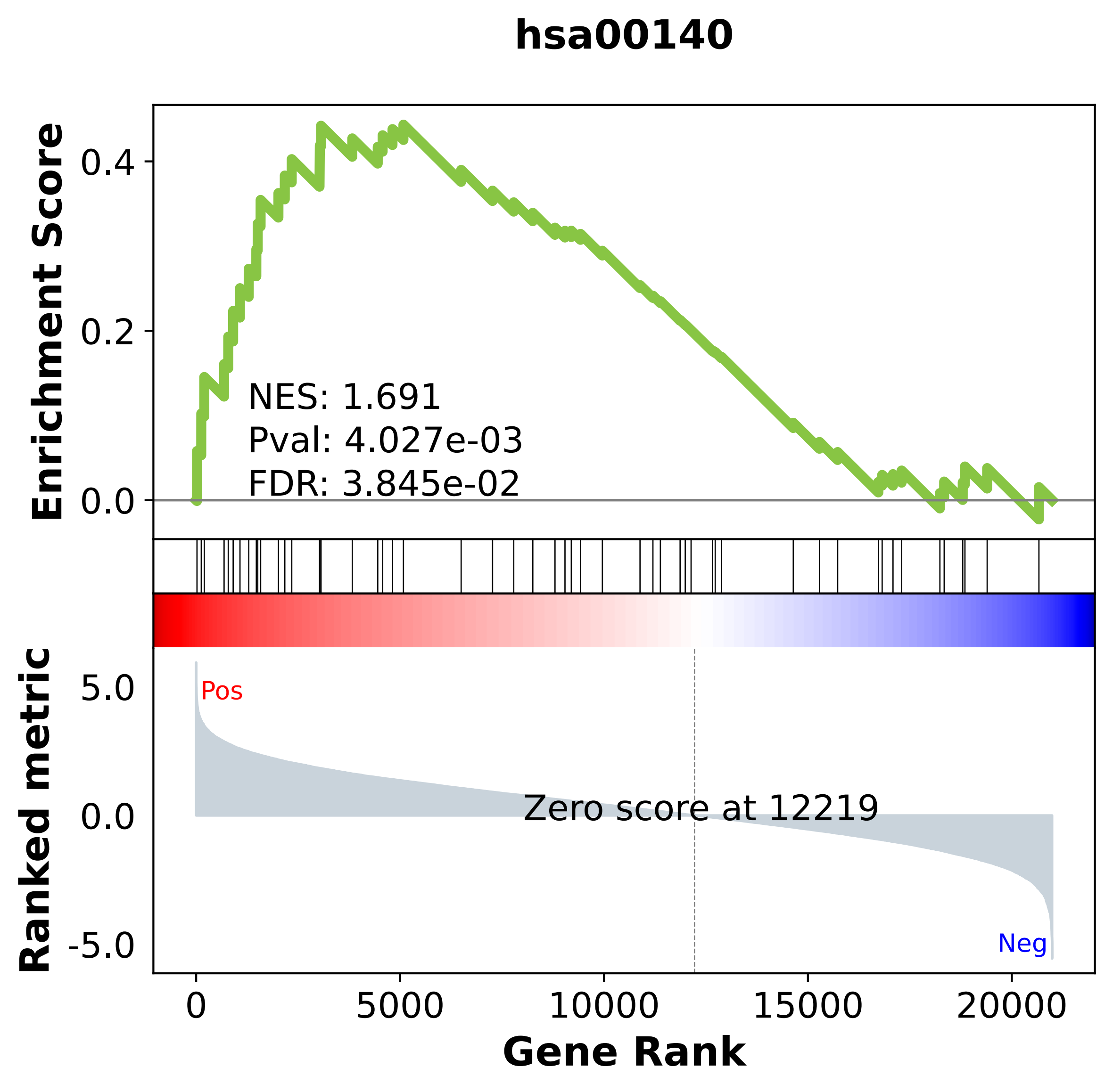

### hsa00190.pdf

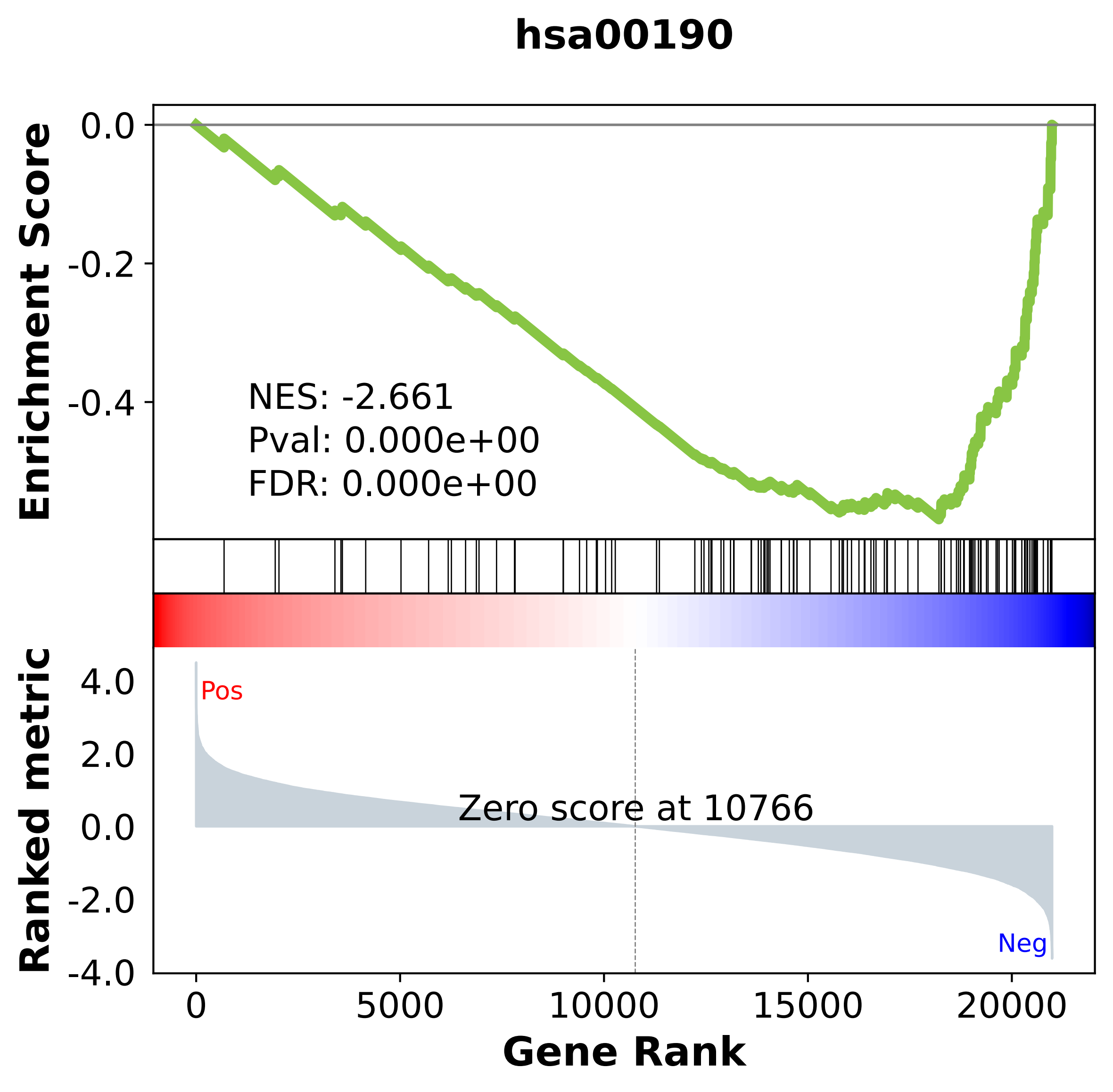

### hsa00270.pdf

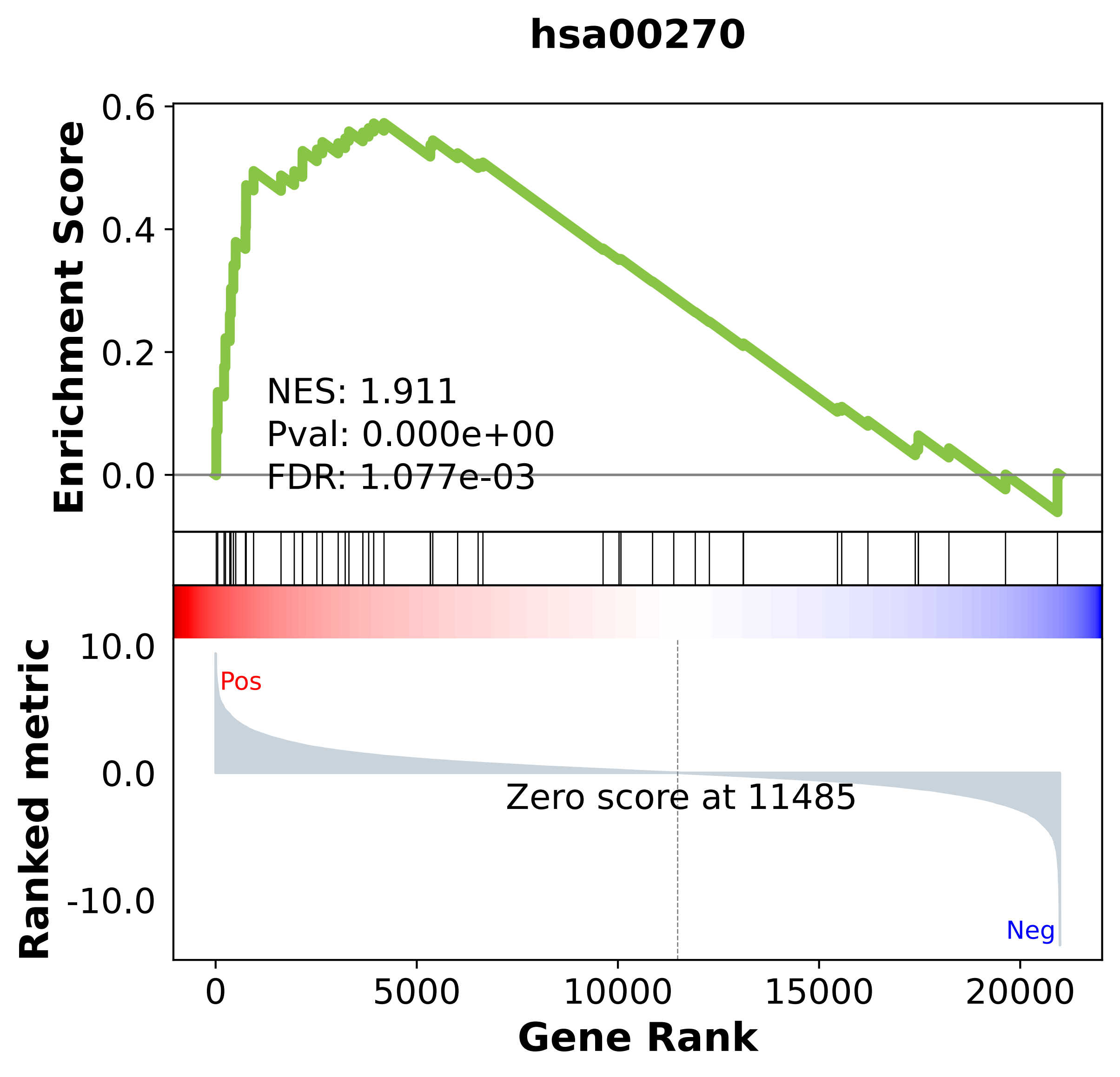

### hsa00280.pdf

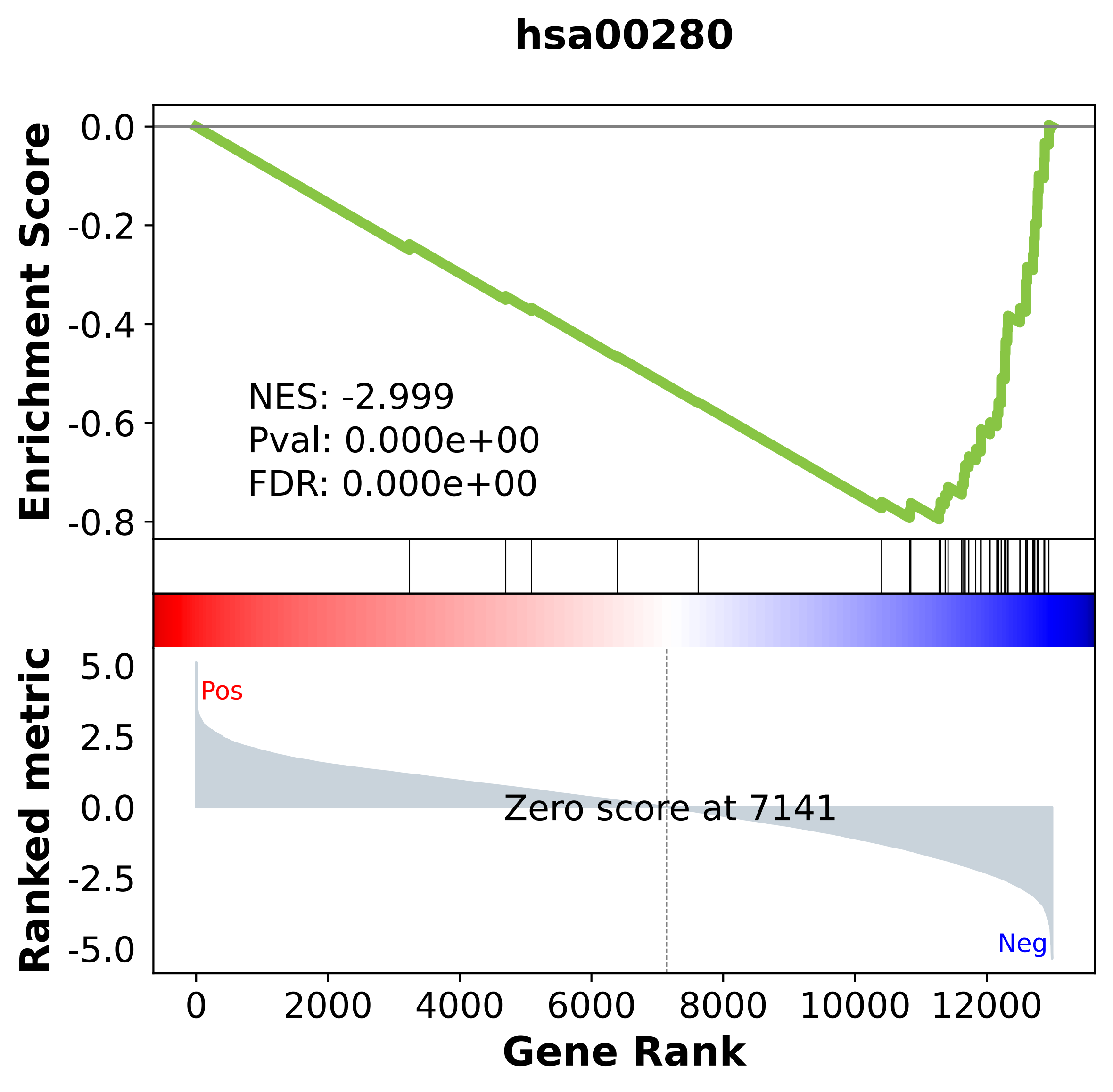

### hsa00330.pdf

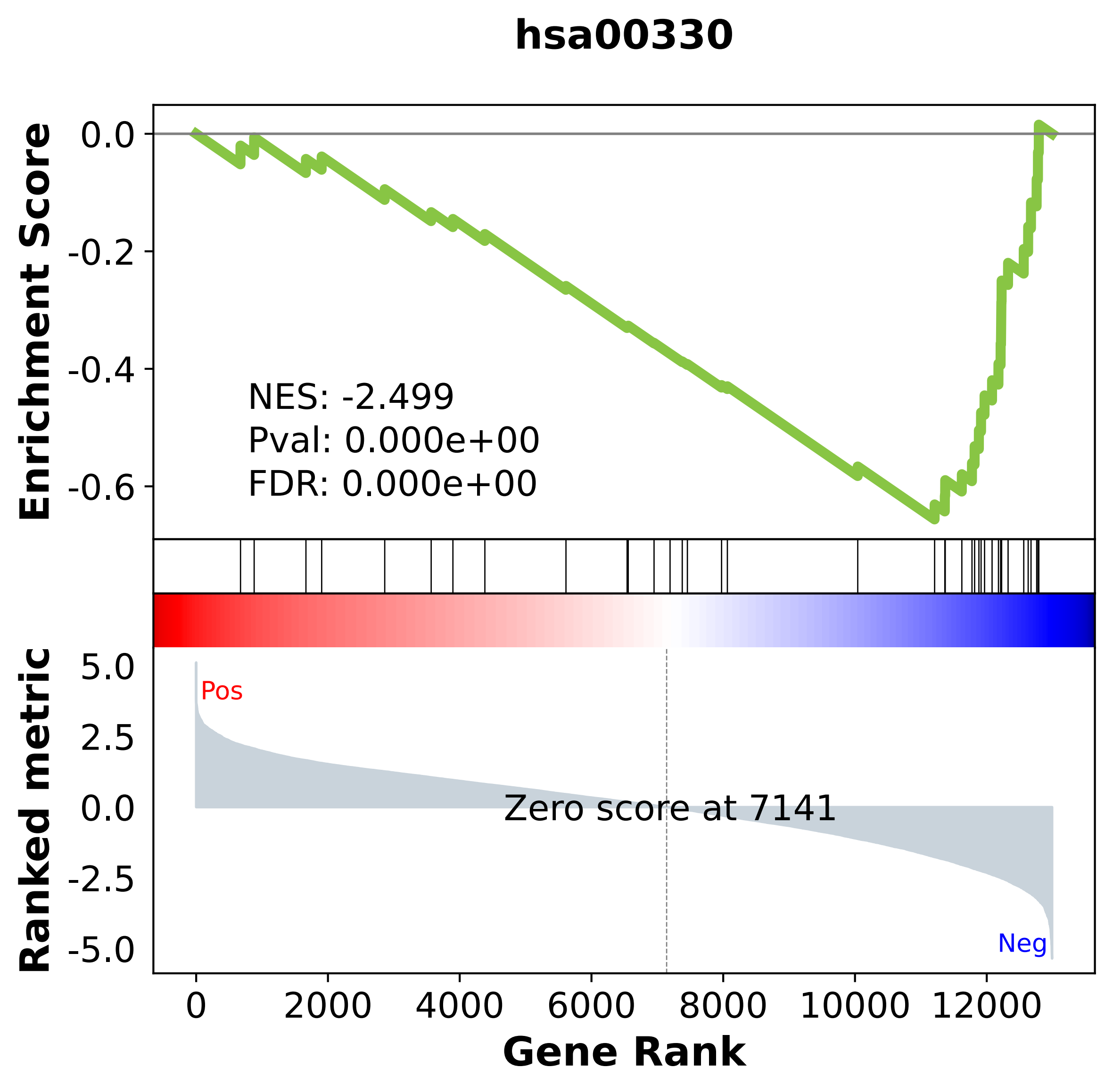

### hsa00350.pdf

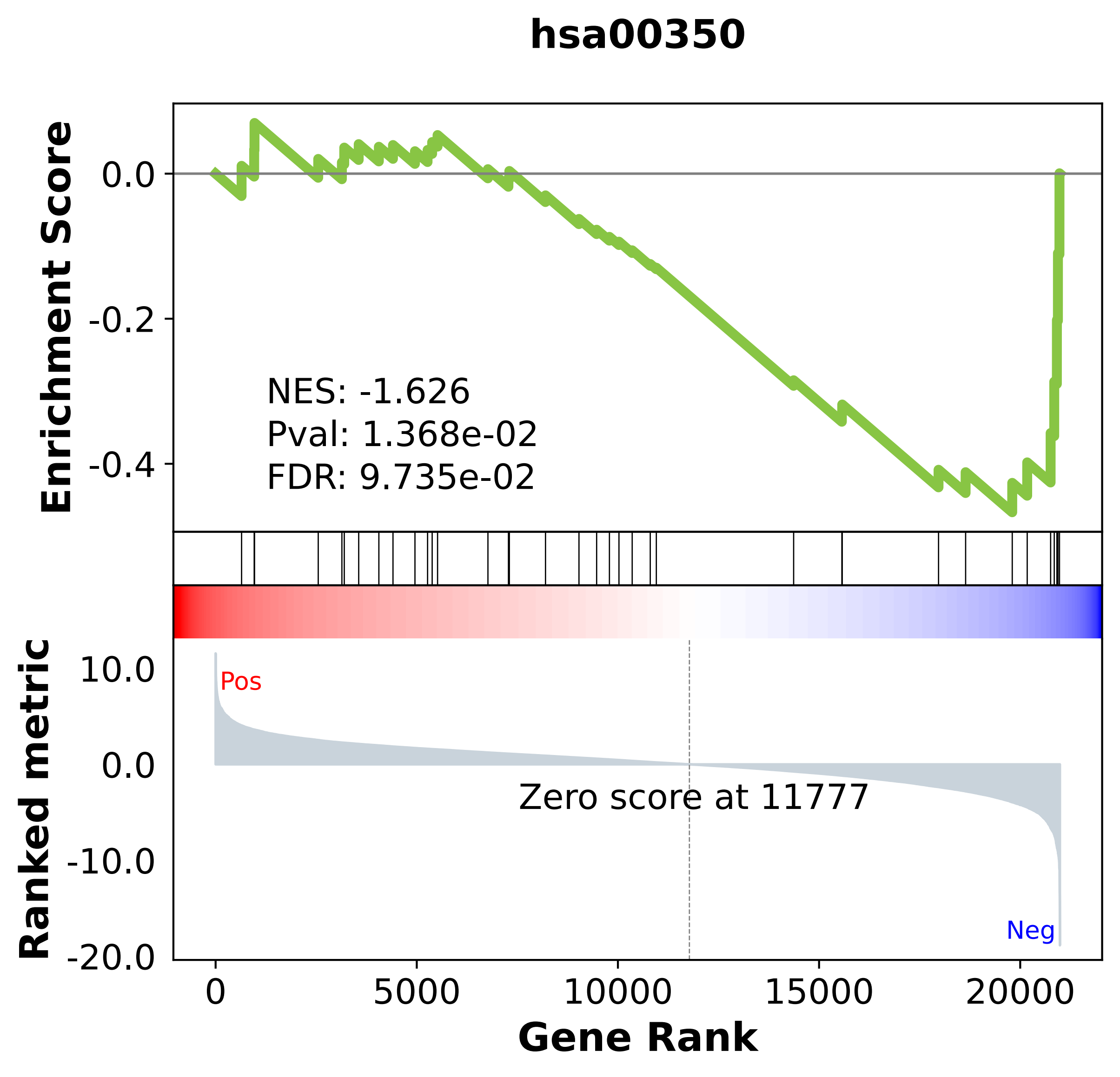

### hsa00360.pdf

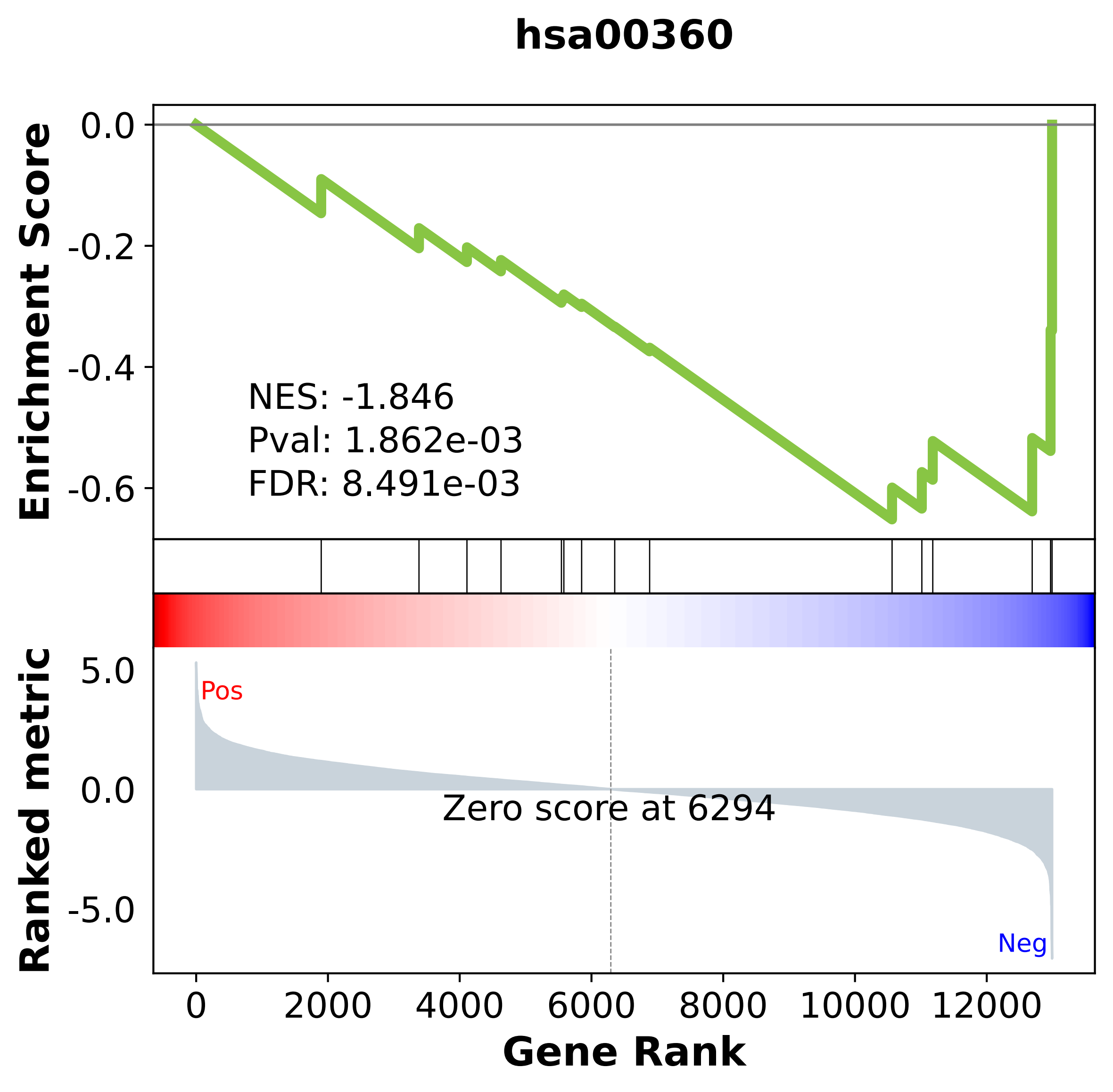

### hsa00410.pdf

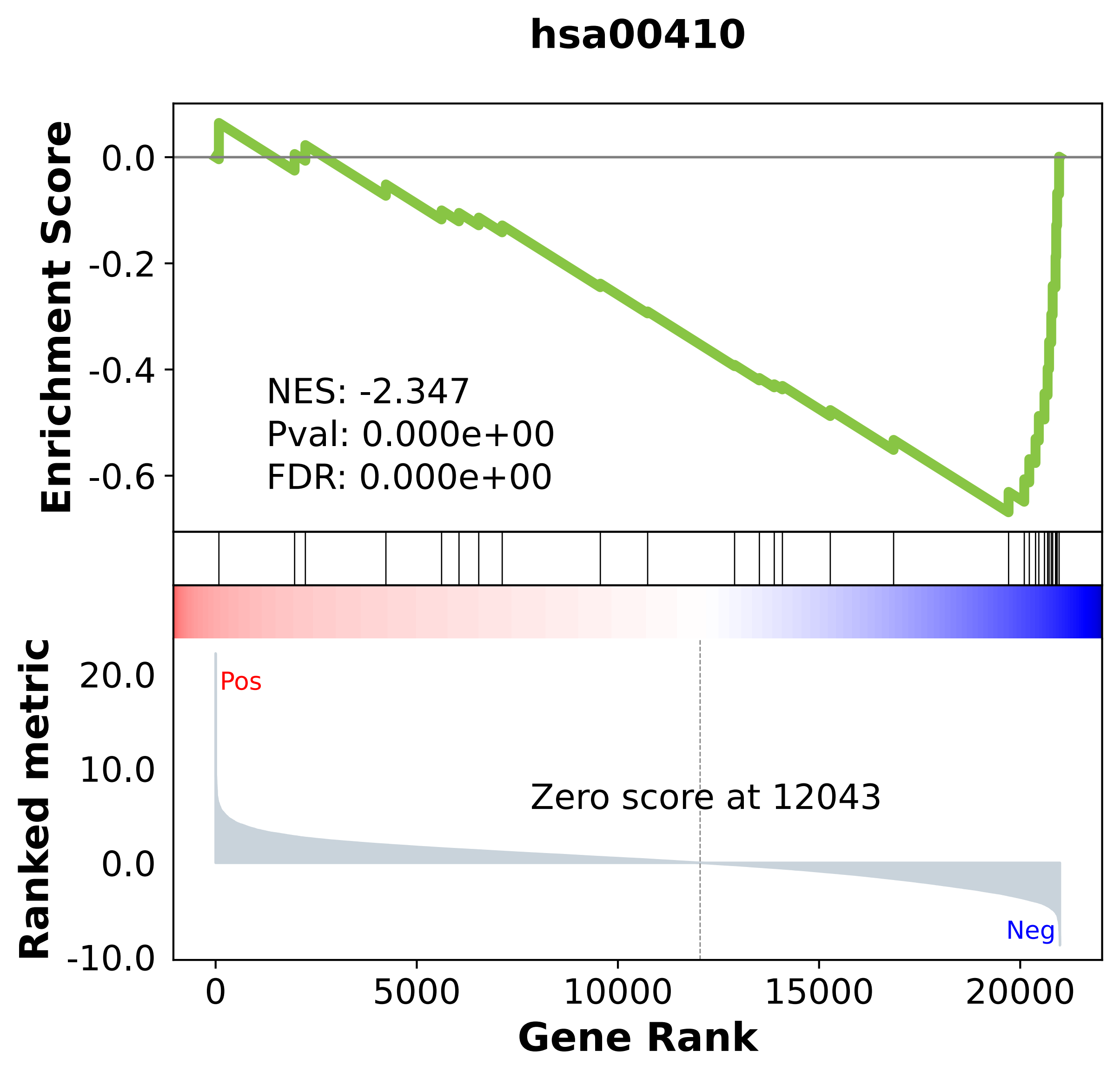

### hsa00480.pdf

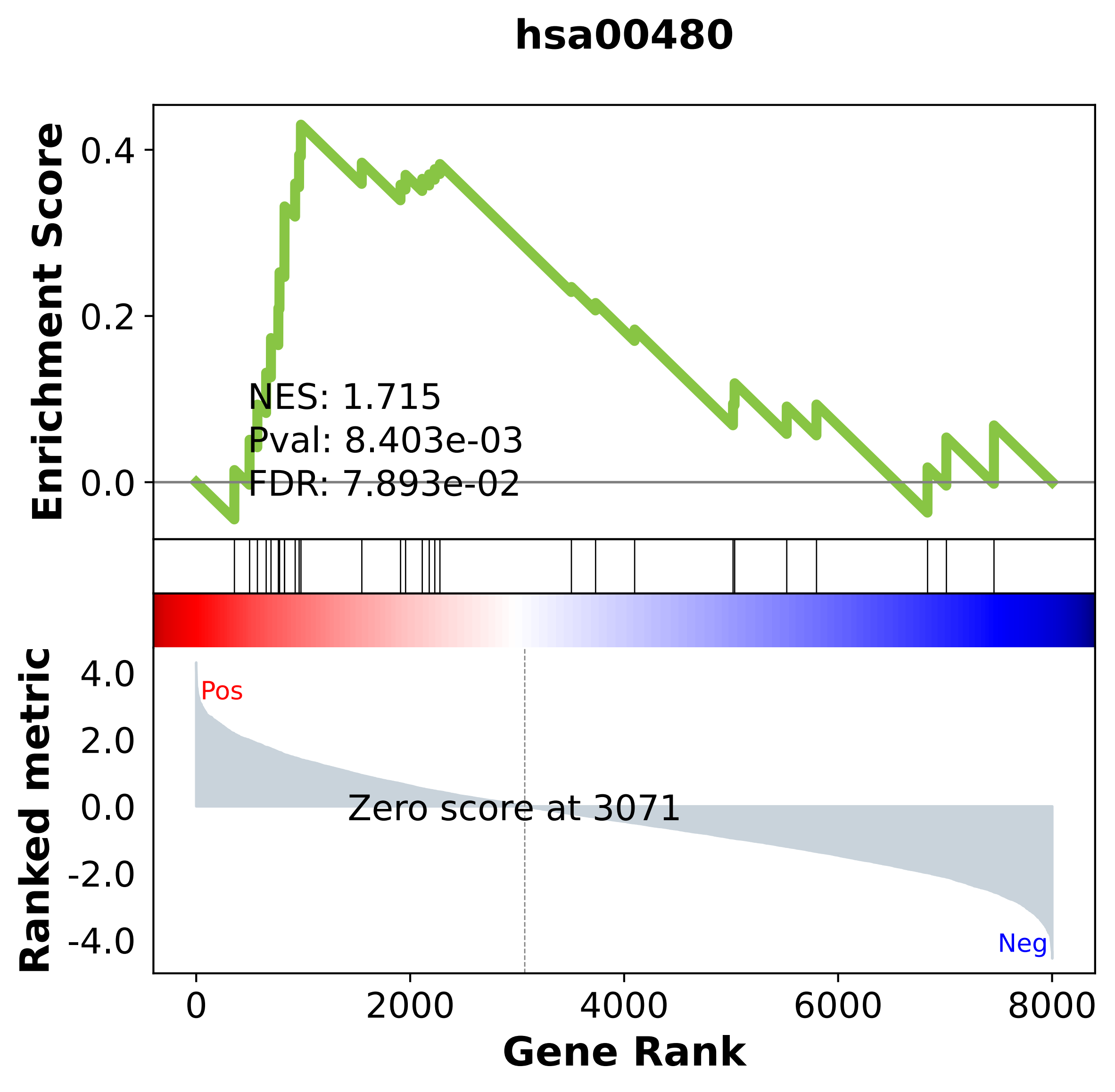

### hsa00510.pdf

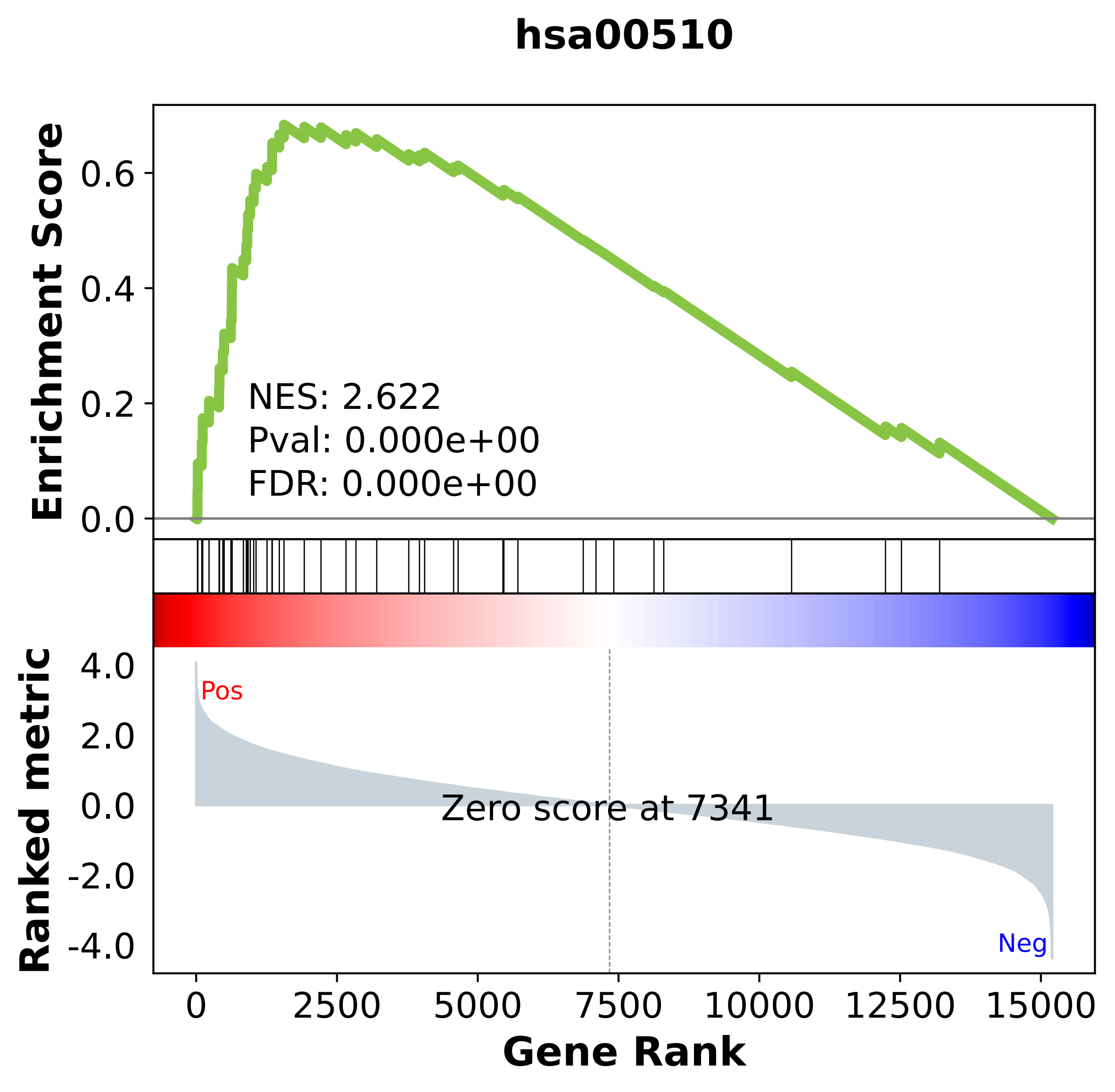

### hsa00512.pdf

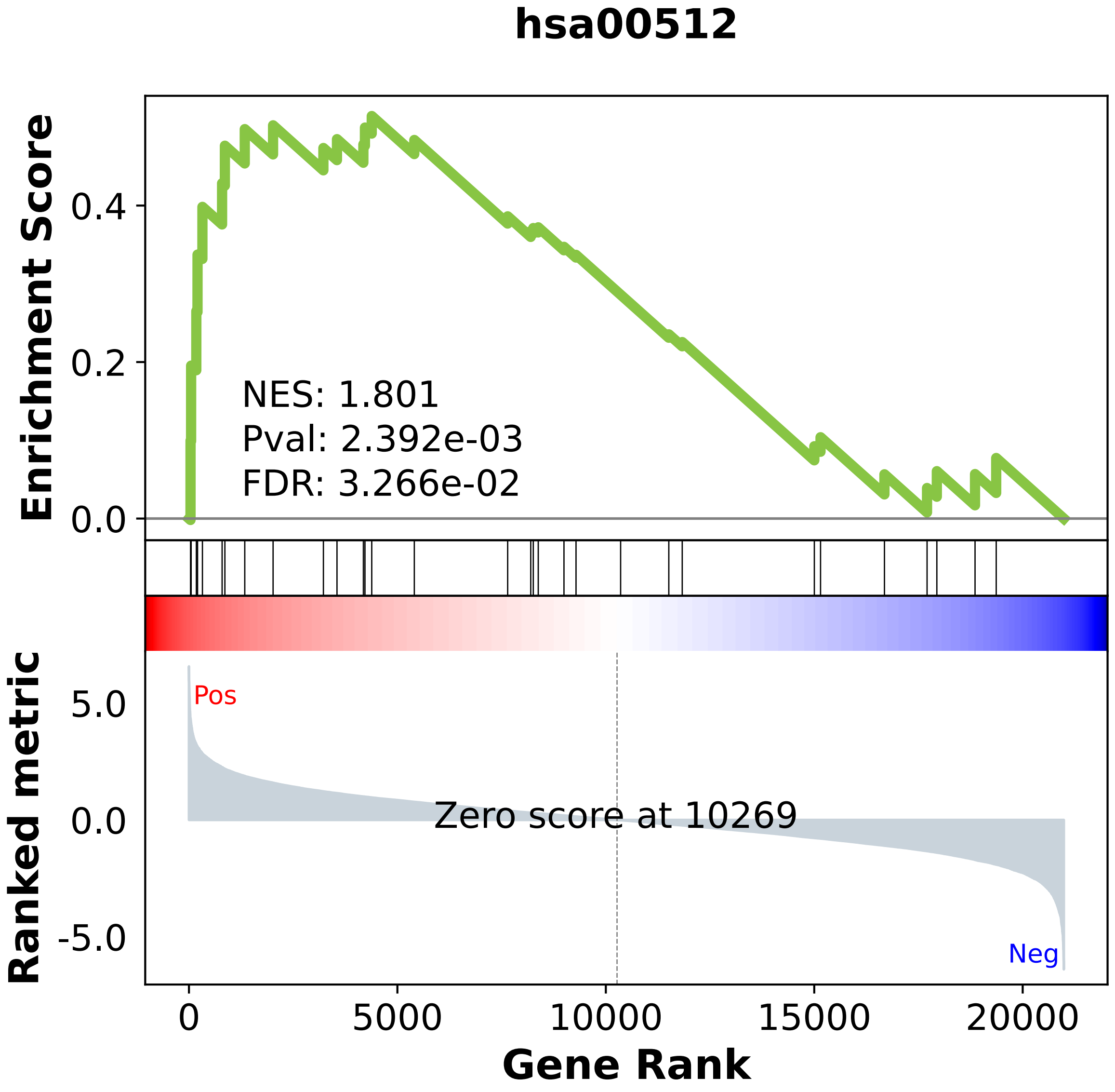

### hsa00513.pdf

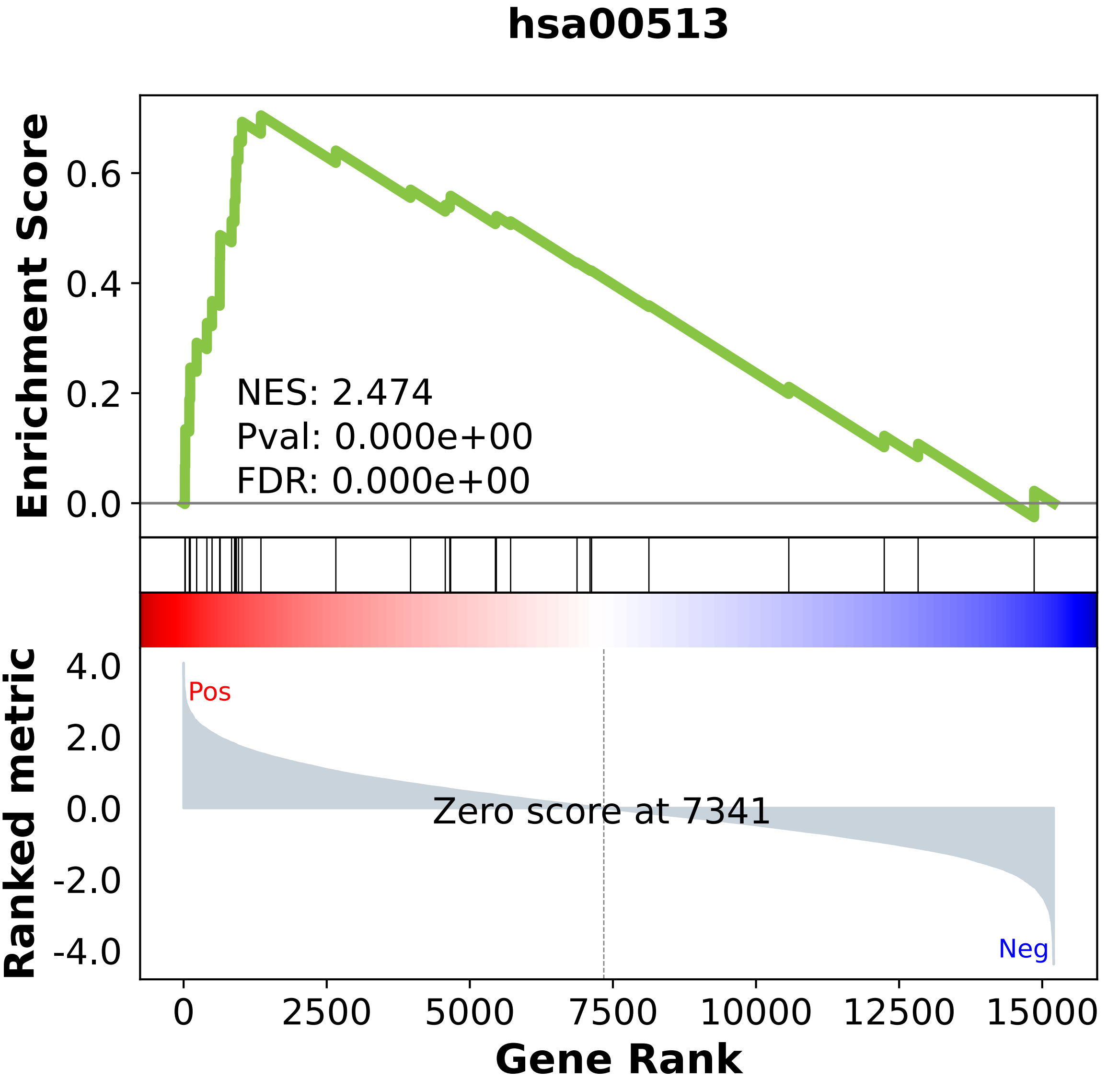

### hsa00520.pdf

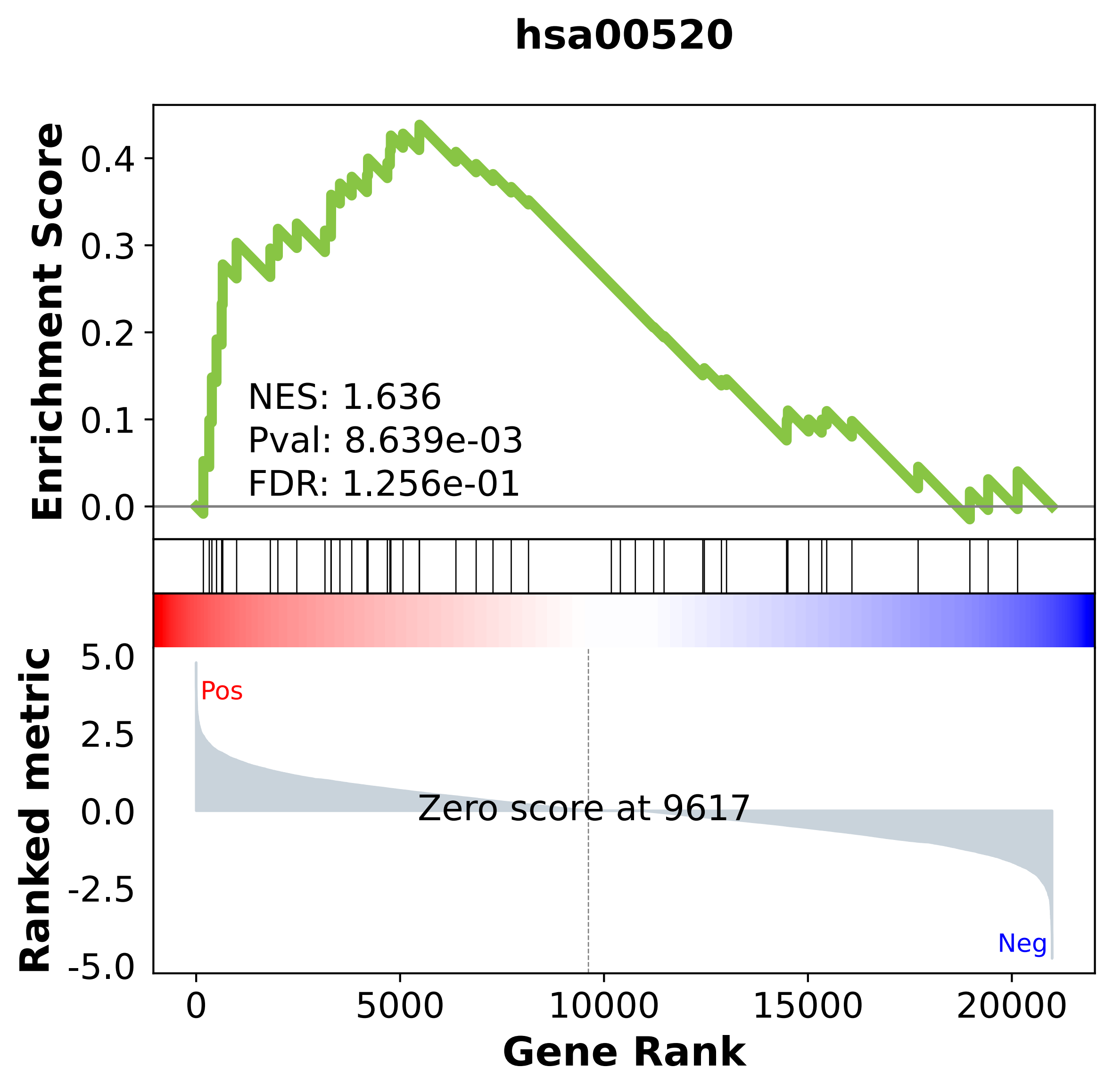

### hsa00531.pdf

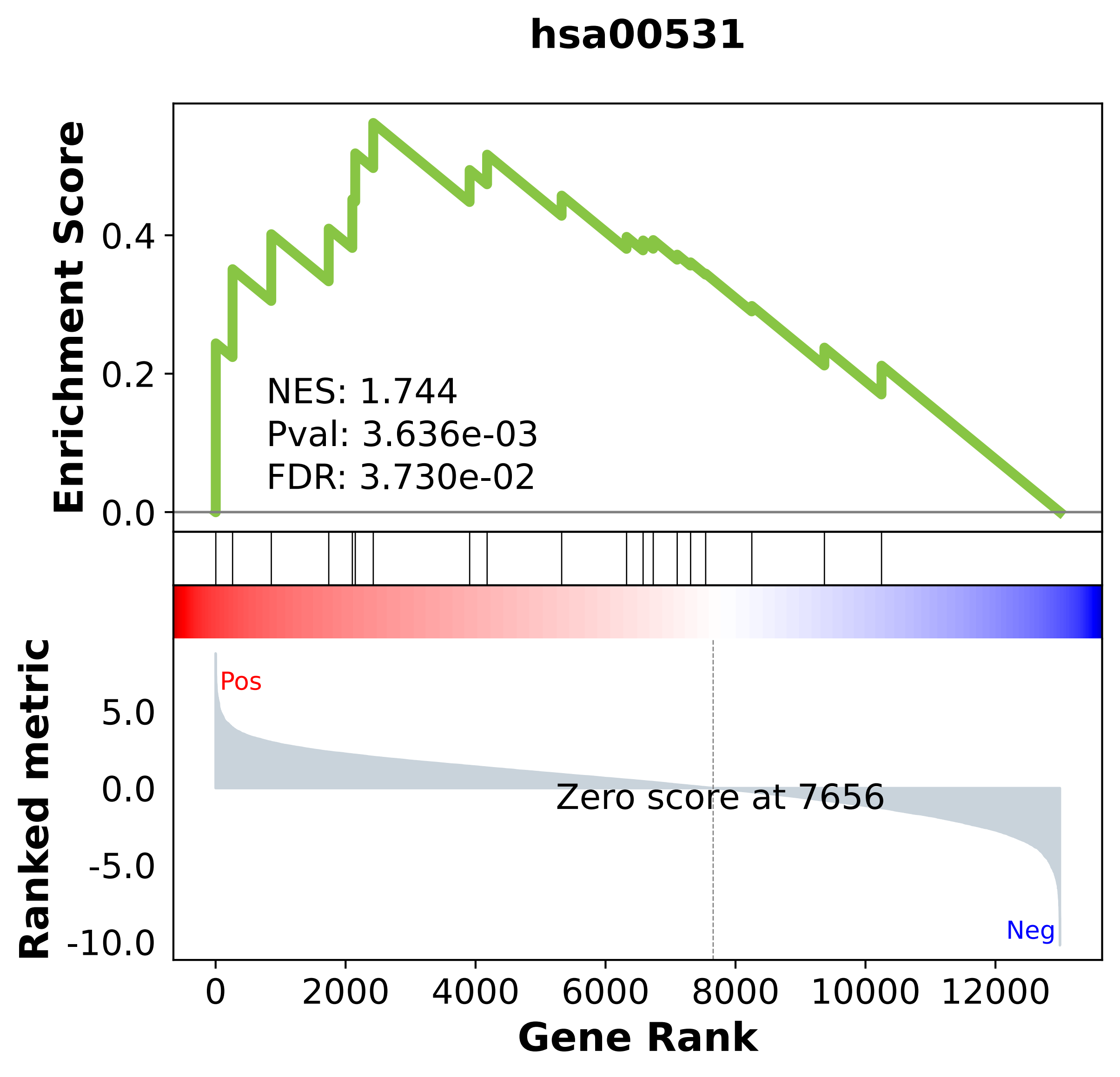

### hsa00532.pdf

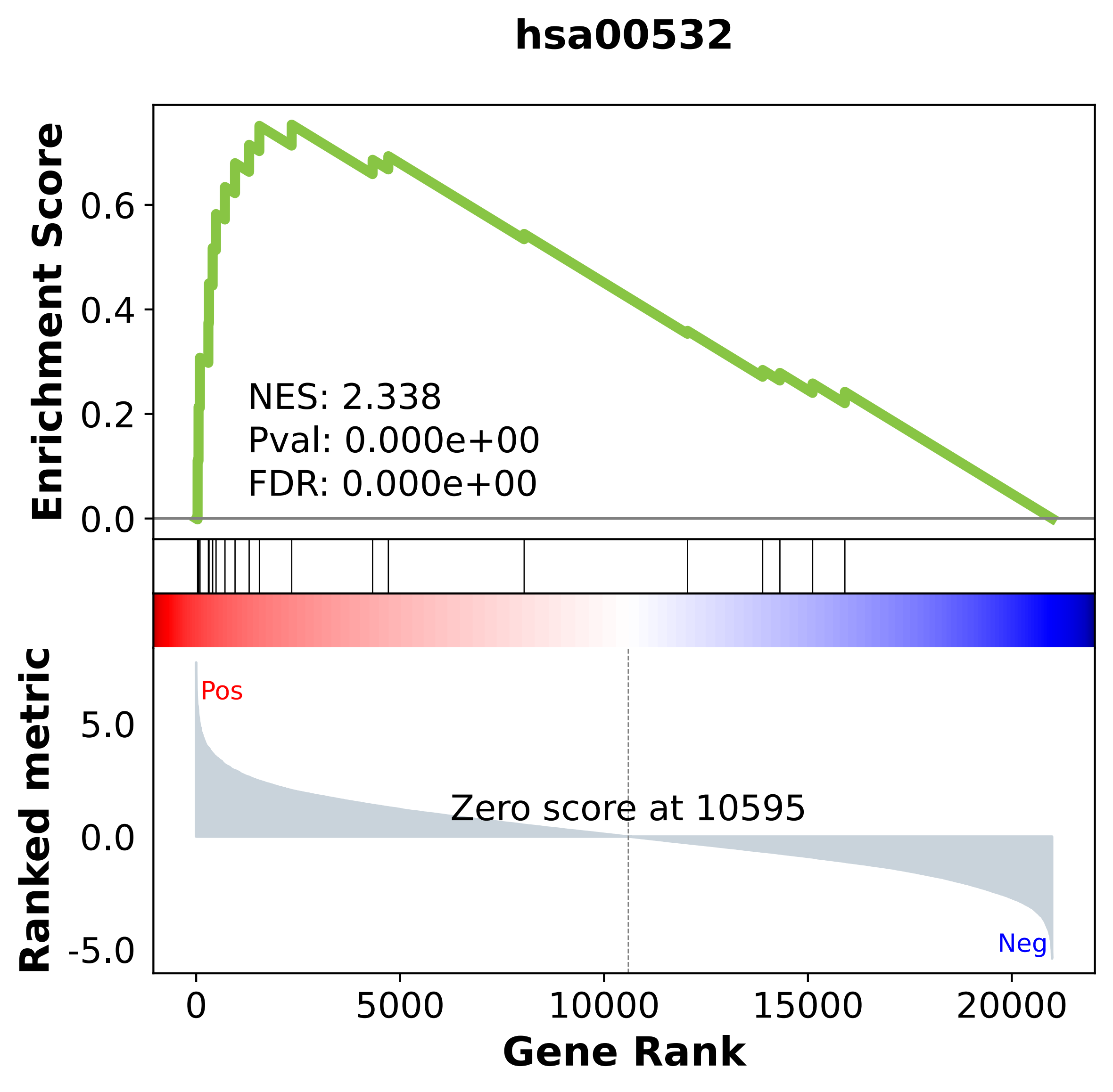

### hsa00561.pdf

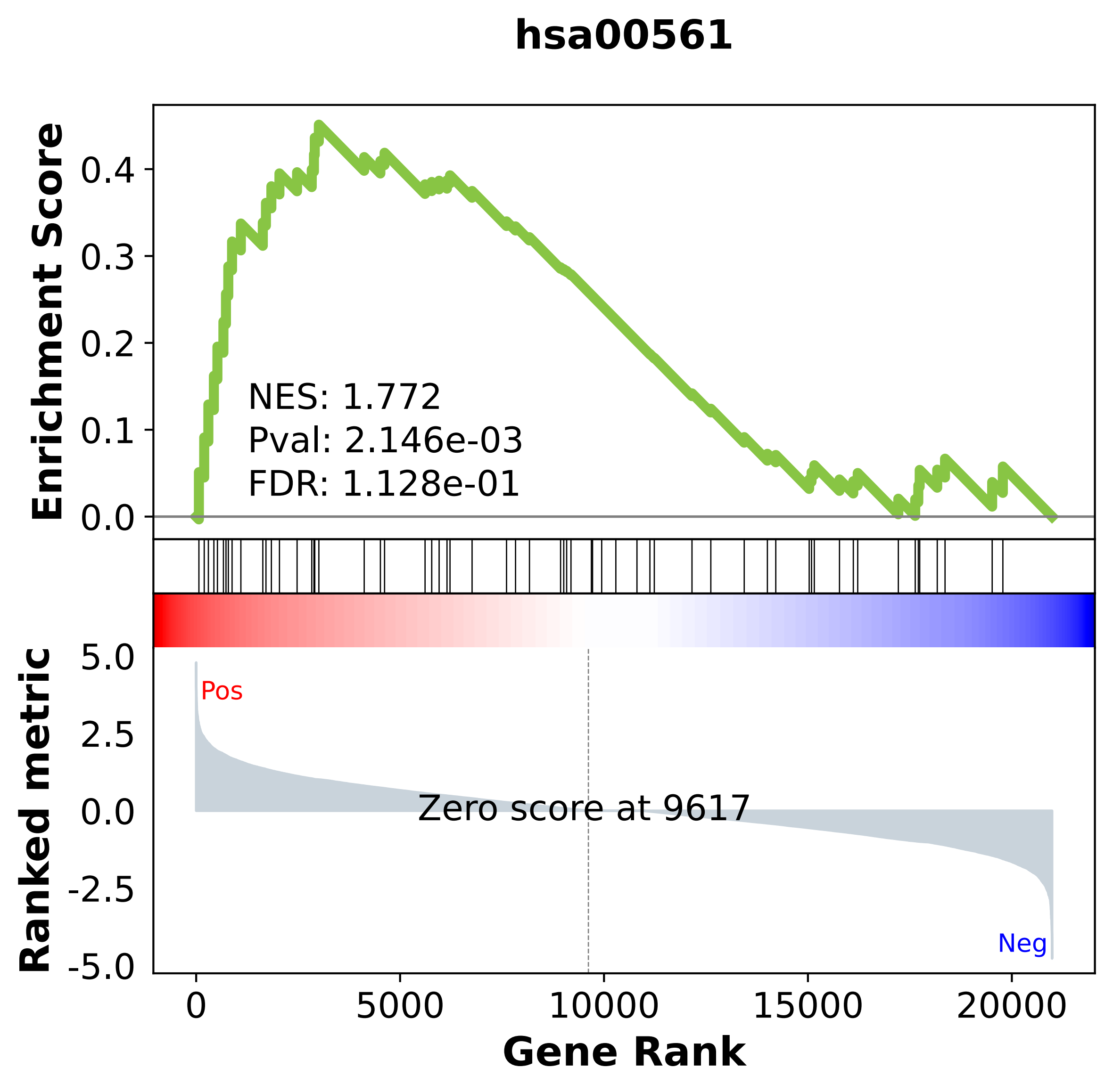

### hsa00563.pdf

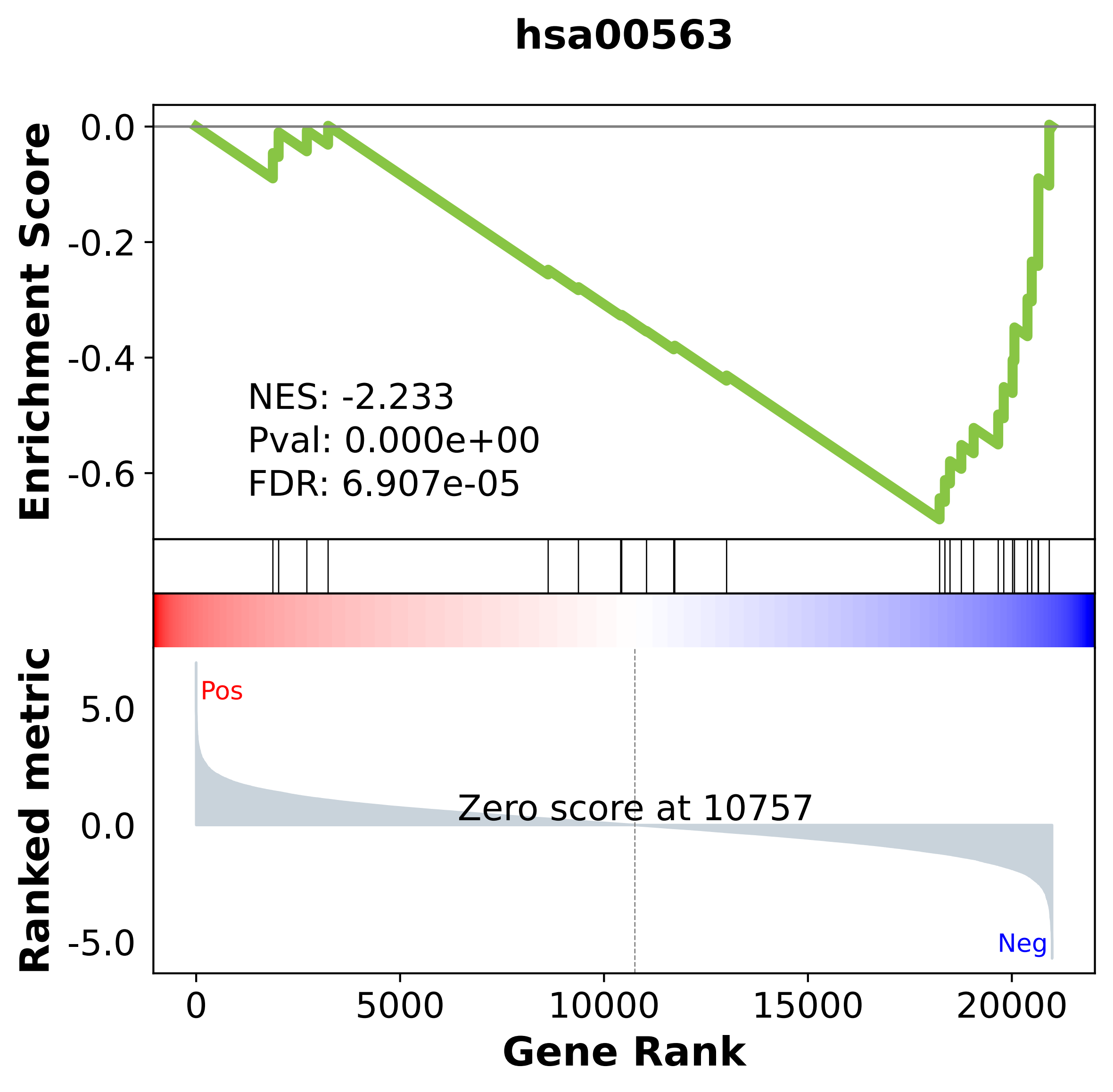

### hsa00564.pdf

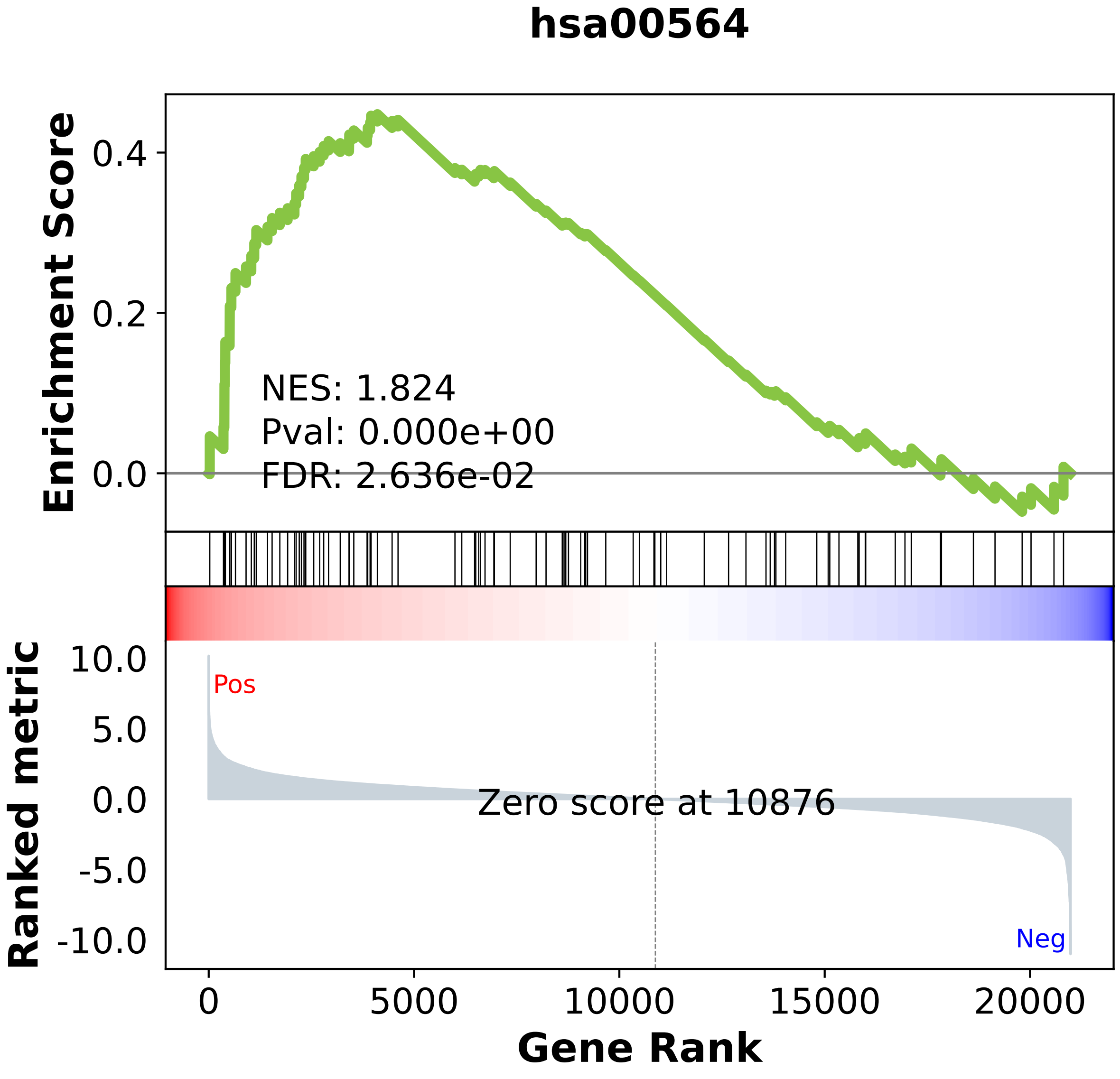

### hsa00590.pdf

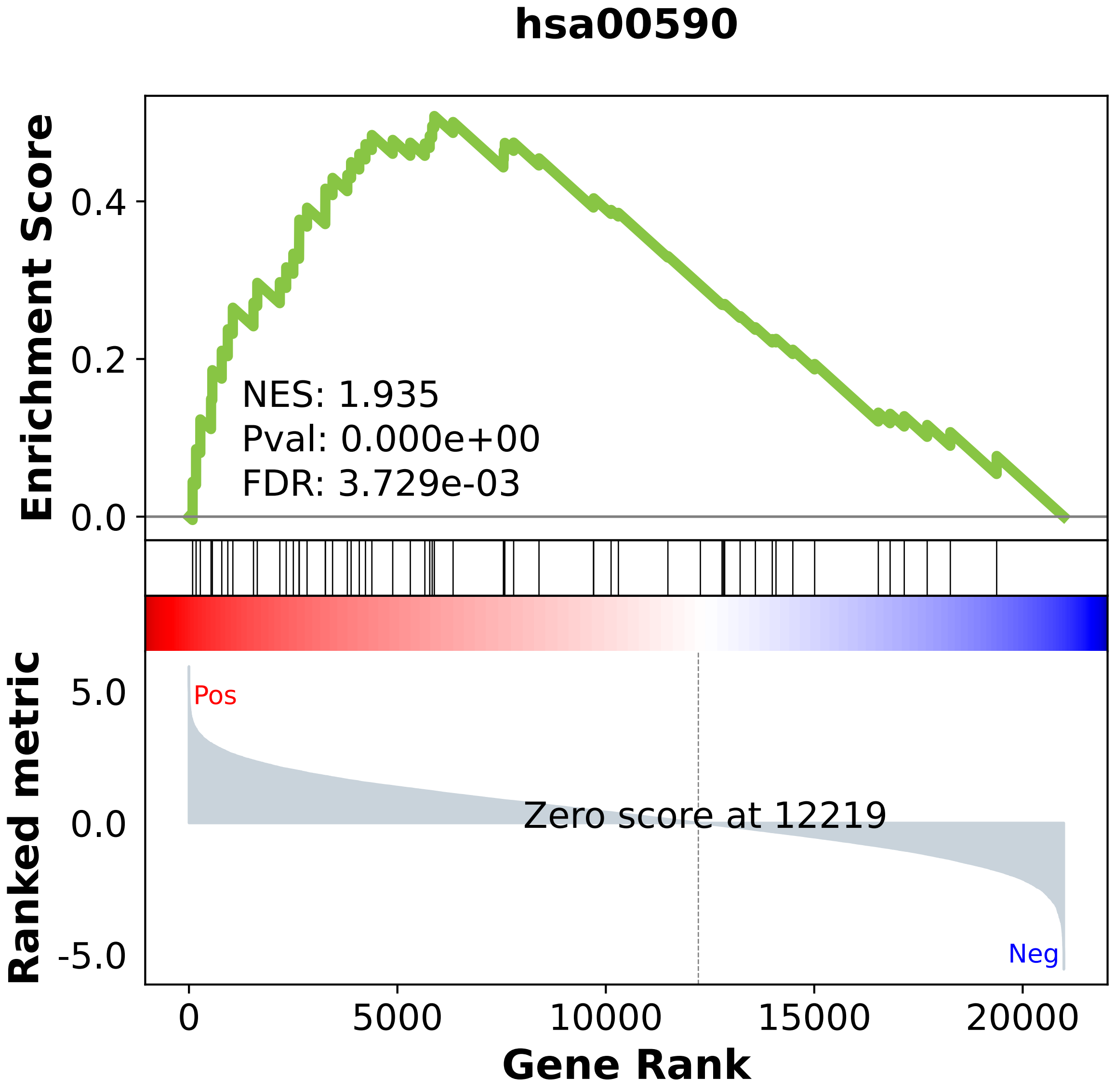

### hsa00620.pdf

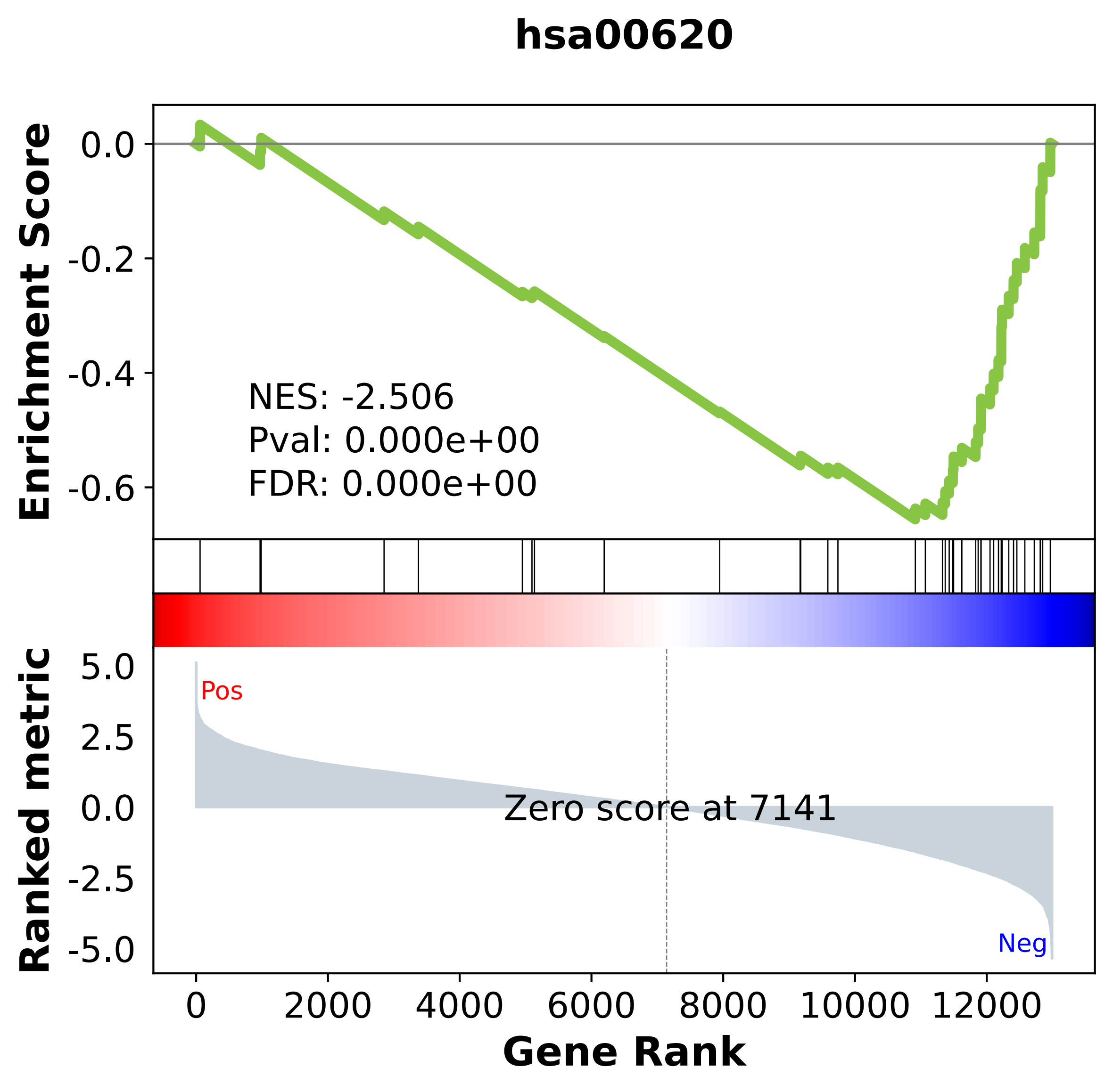

### hsa00630.pdf

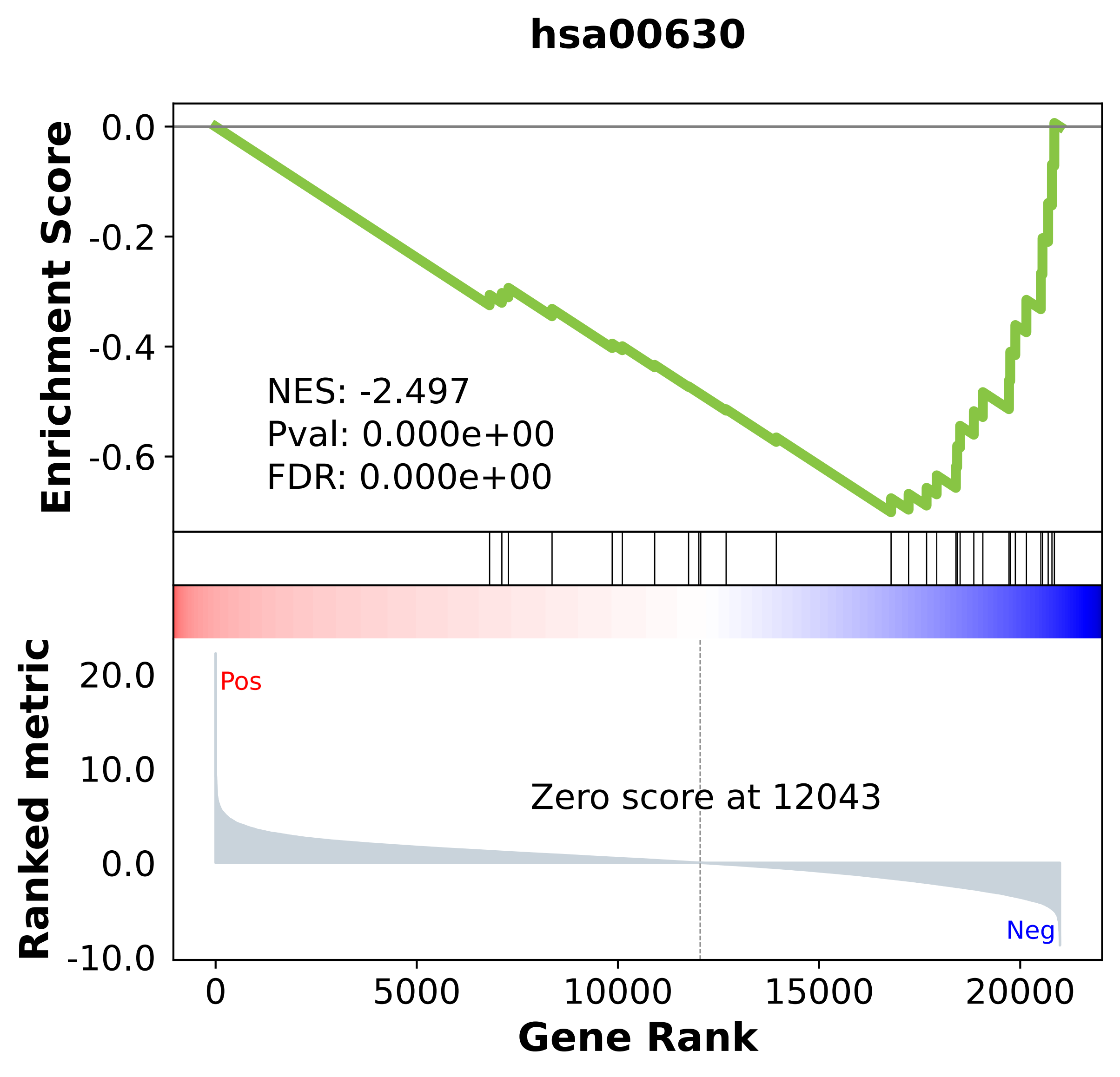

### hsa00640.pdf

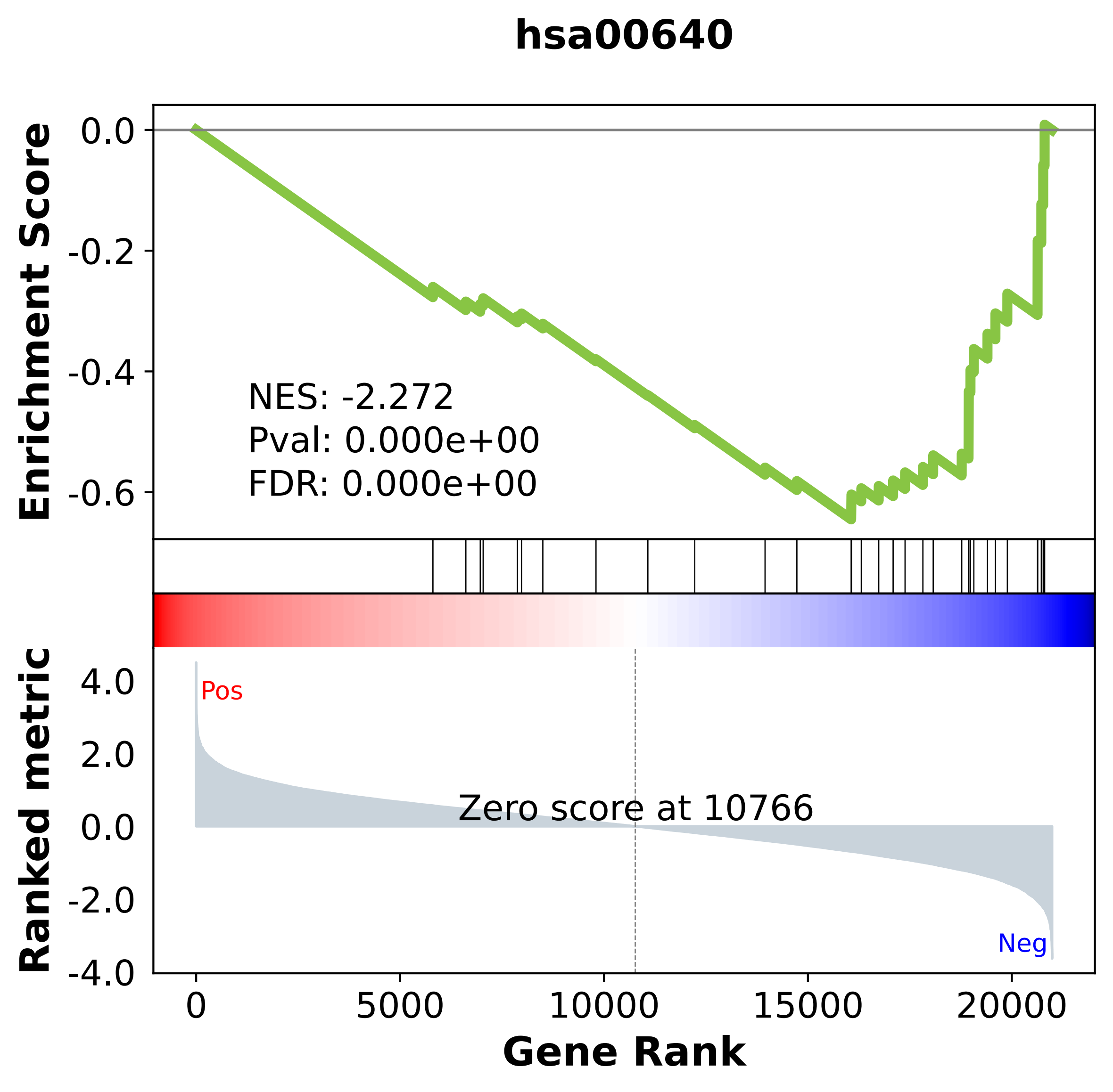

### hsa00650.pdf

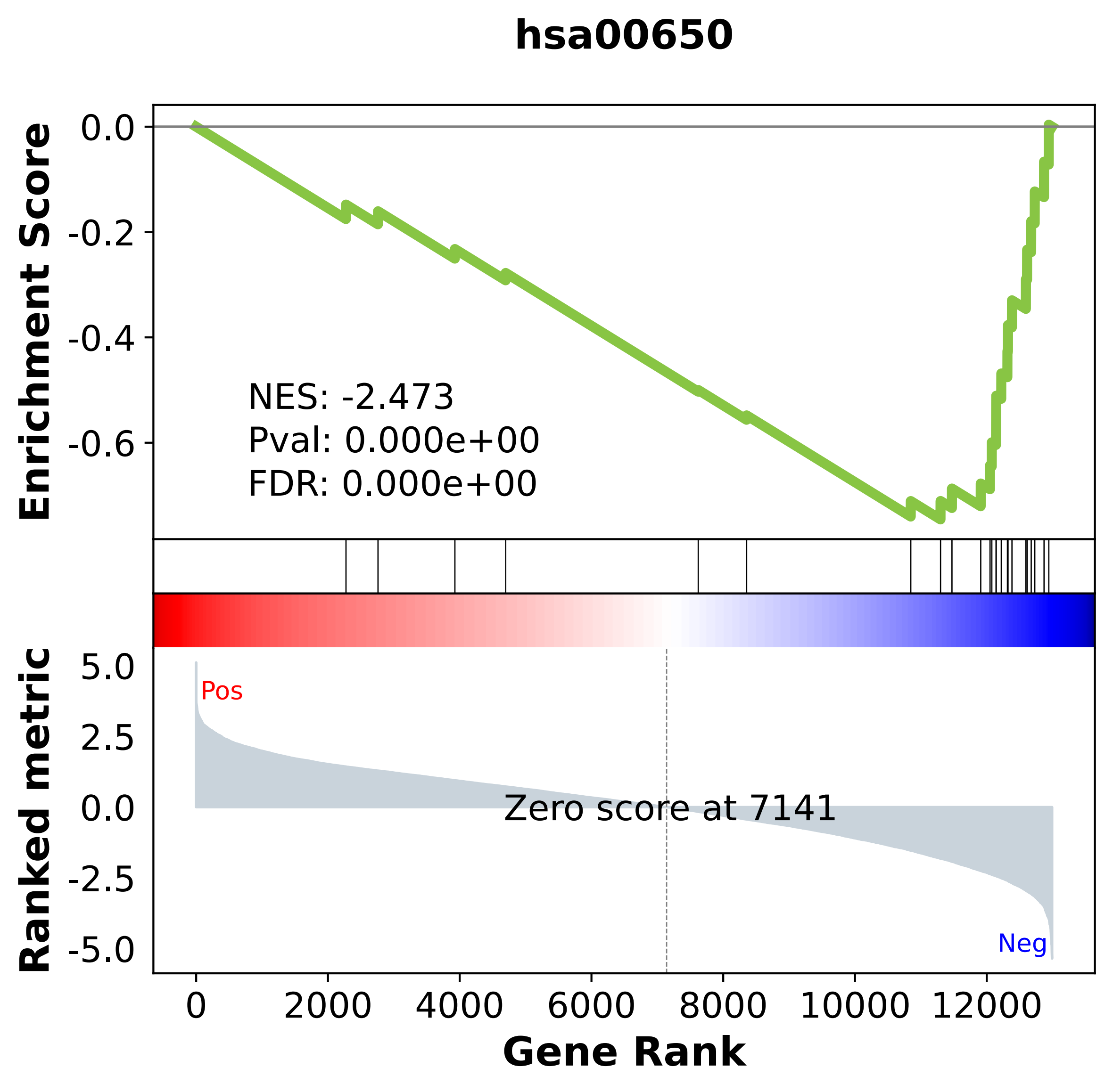

### hsa00670.pdf

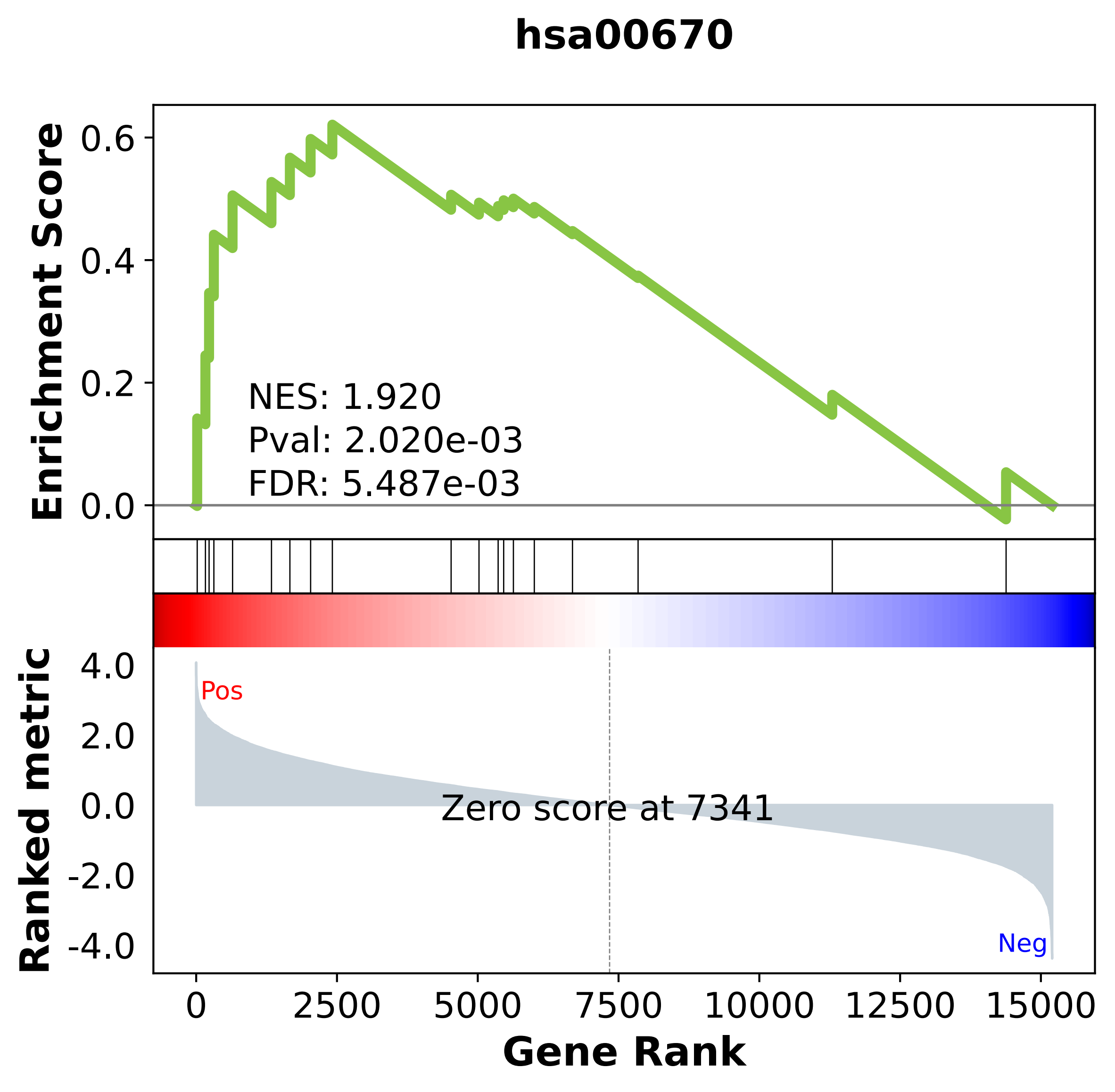

### hsa00730.pdf

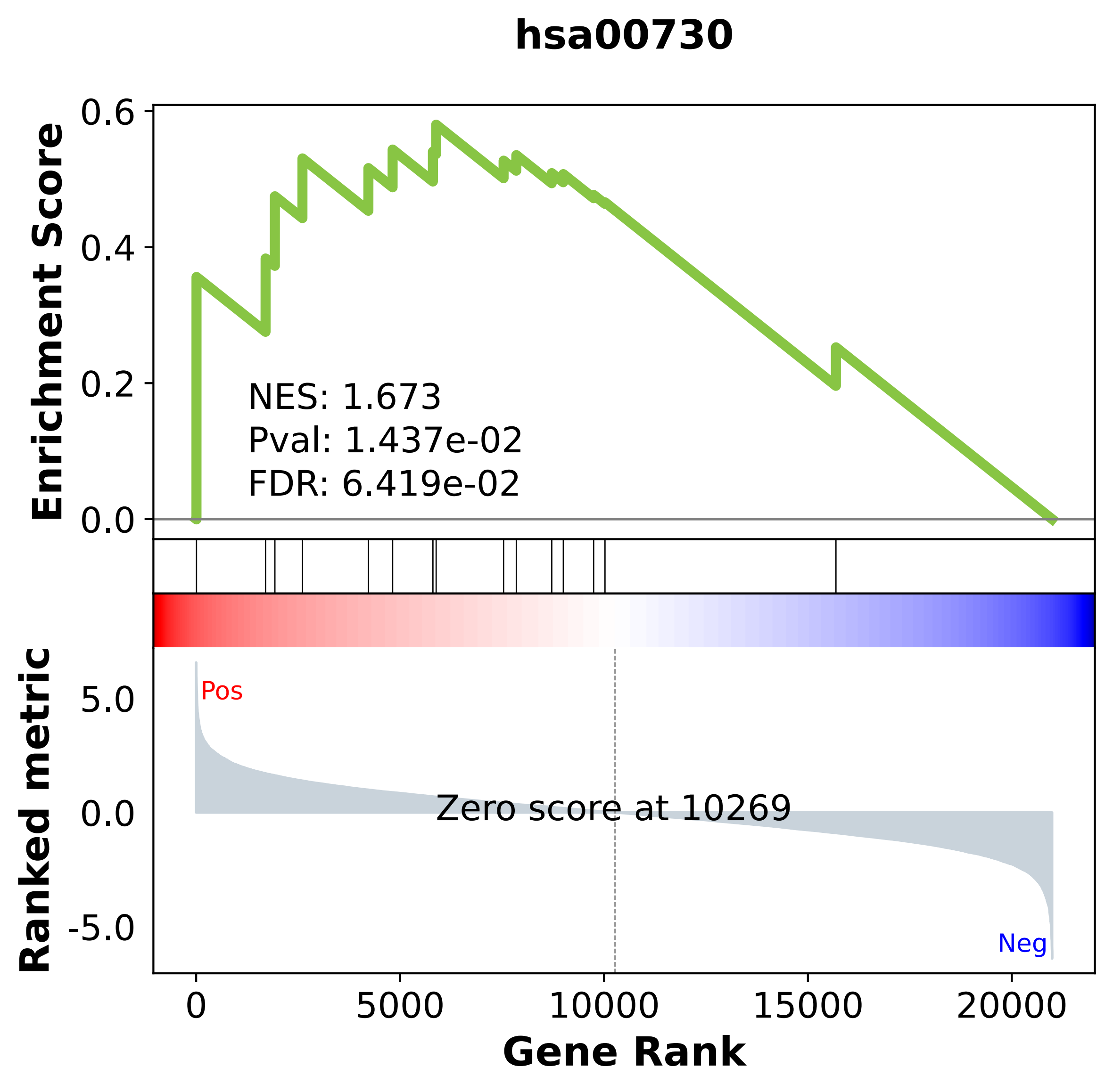

### hsa00830.pdf

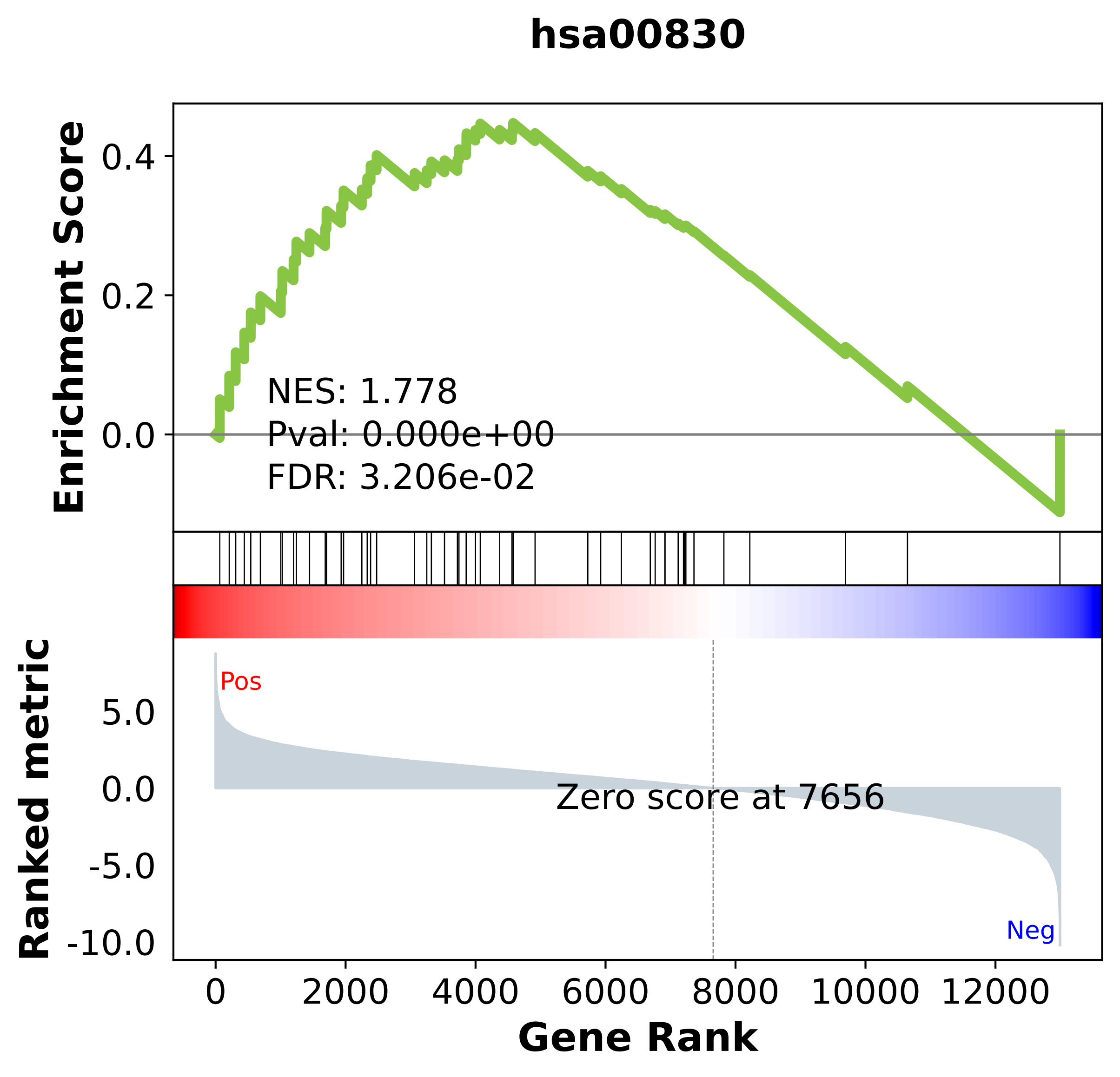

### hsa00860.pdf

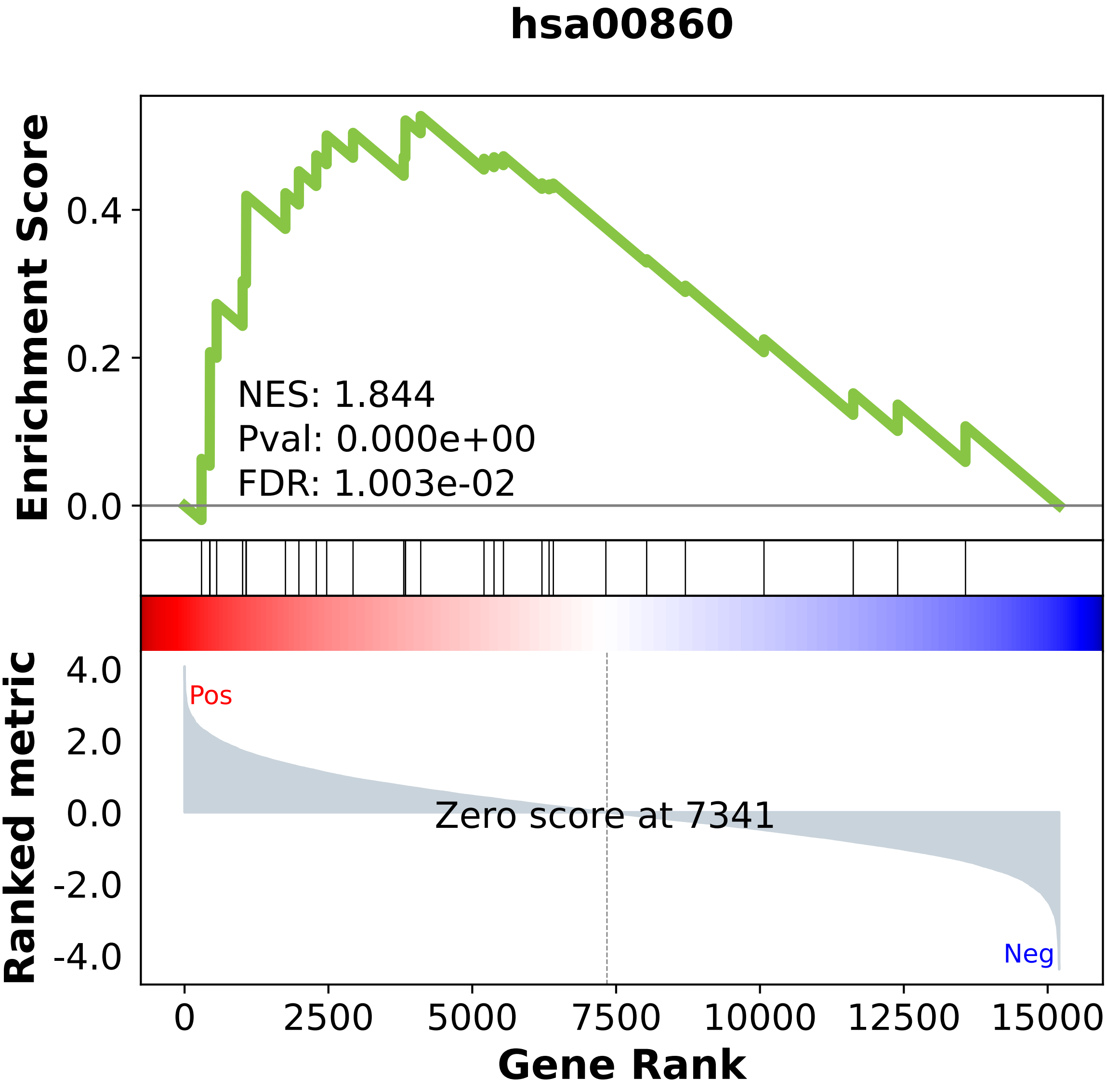

### hsa00900.pdf

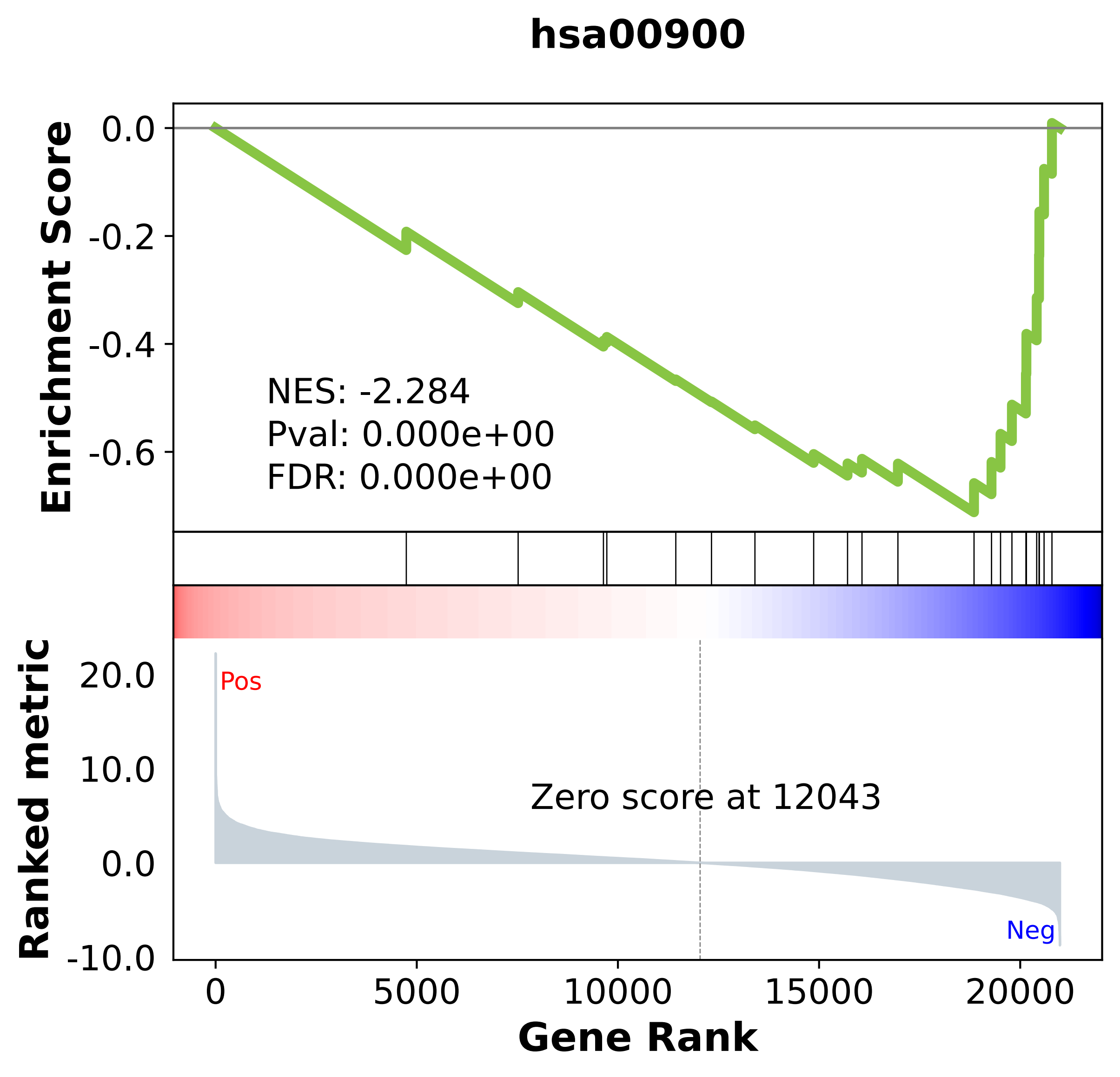

### hsa00910.pdf

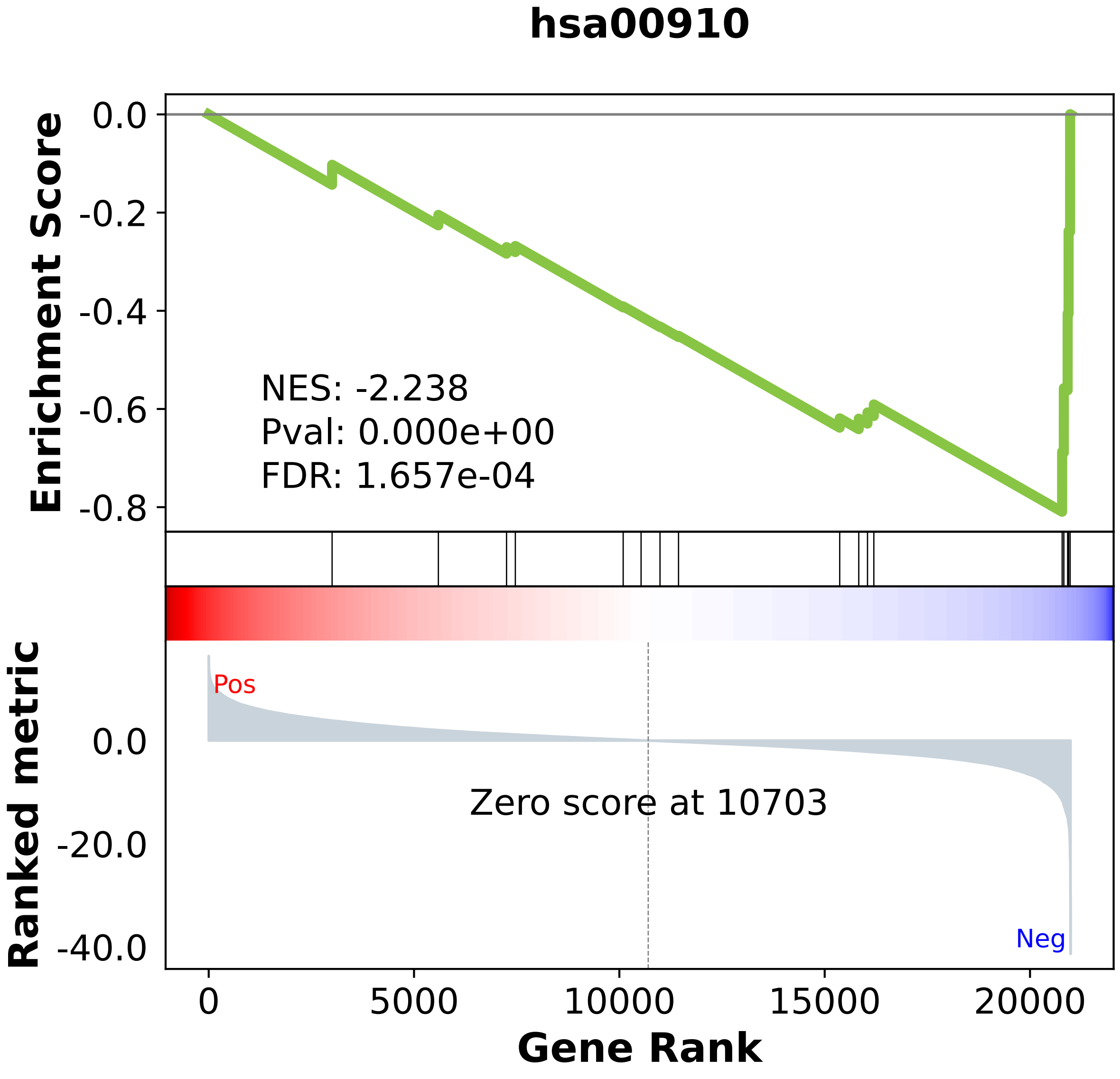

### hsa00970.pdf

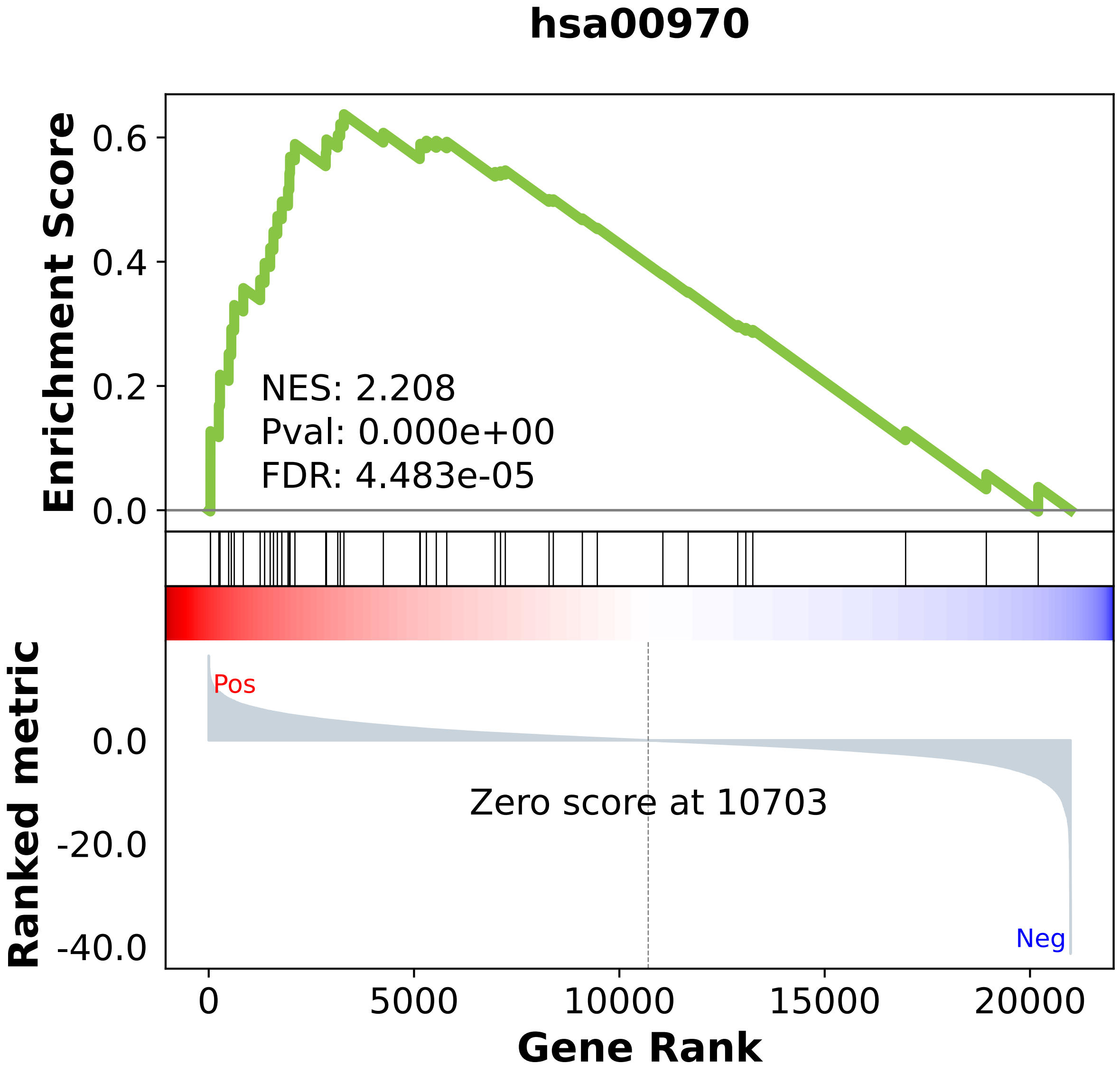

### hsa00980.pdf

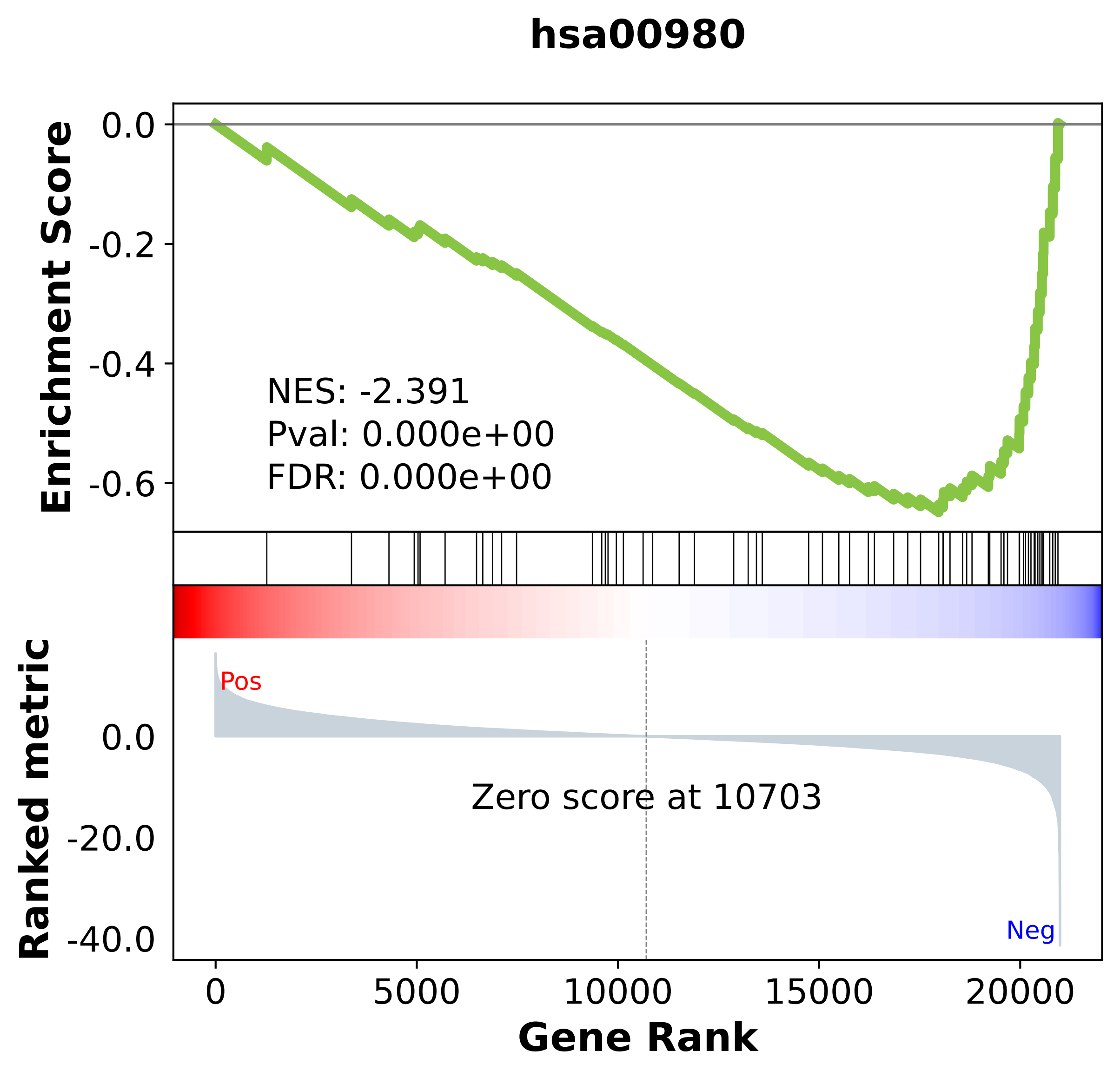

### hsa00982.pdf

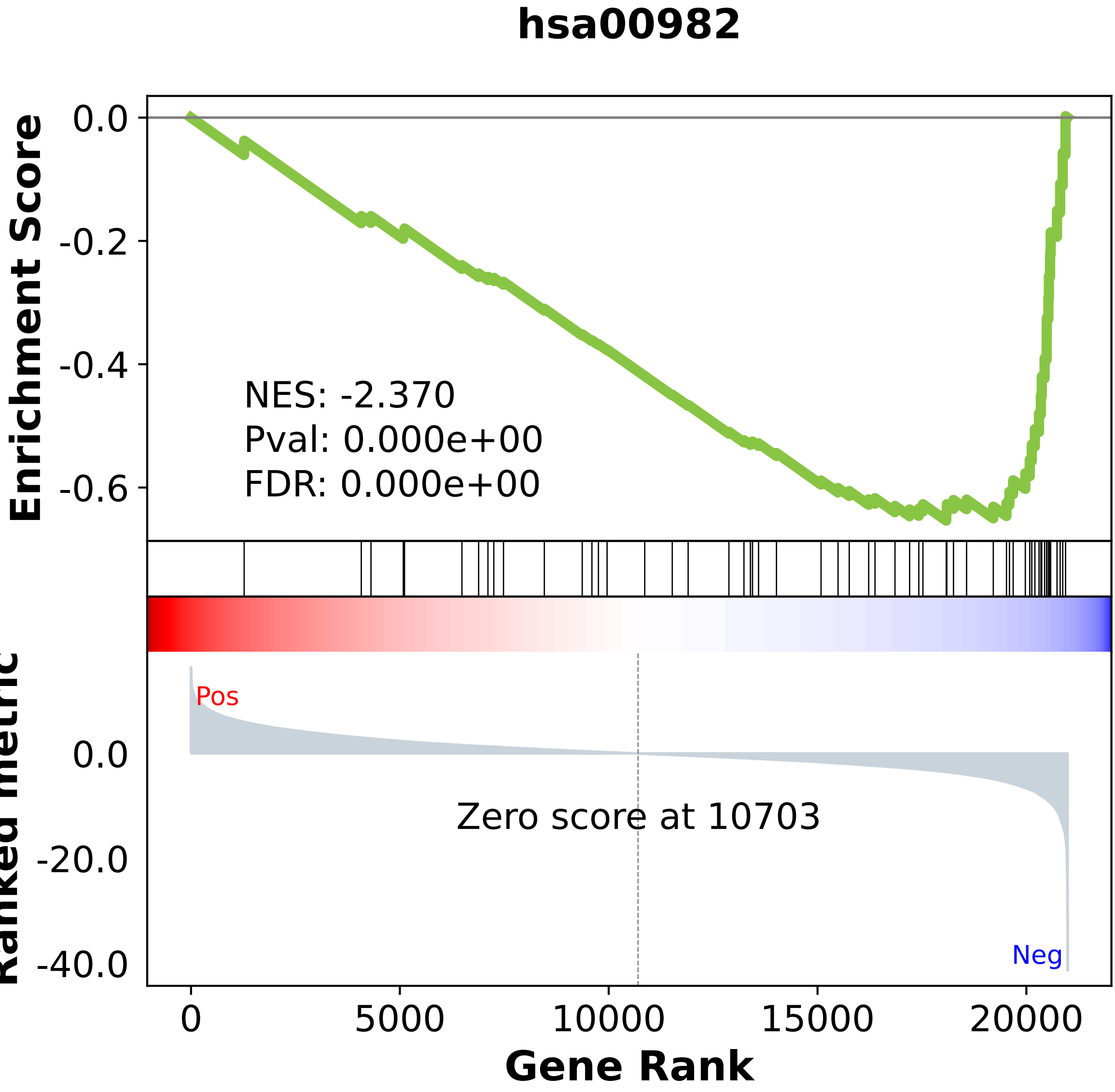

### hsa01040.pdf

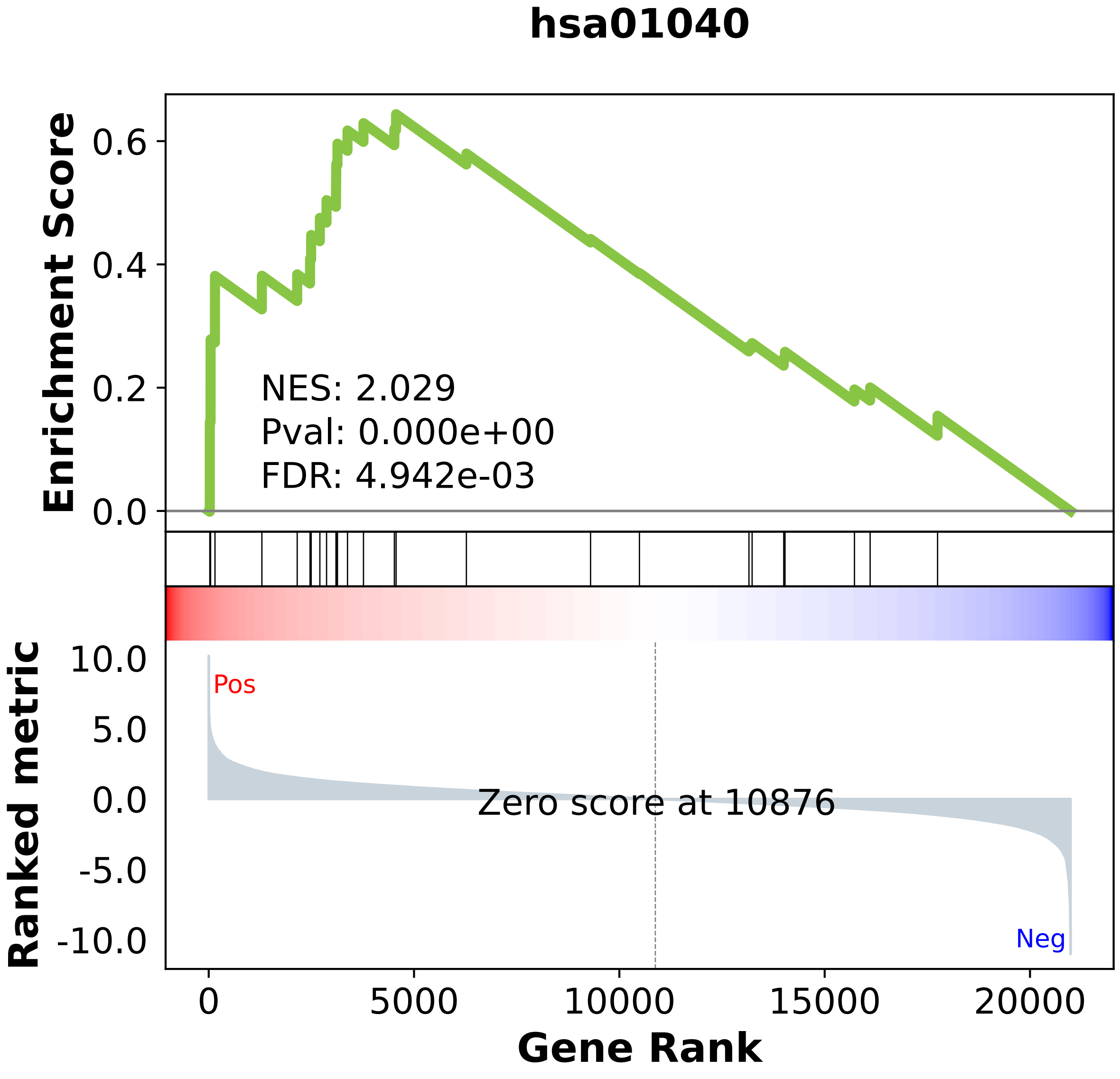

### hsa03008.pdf

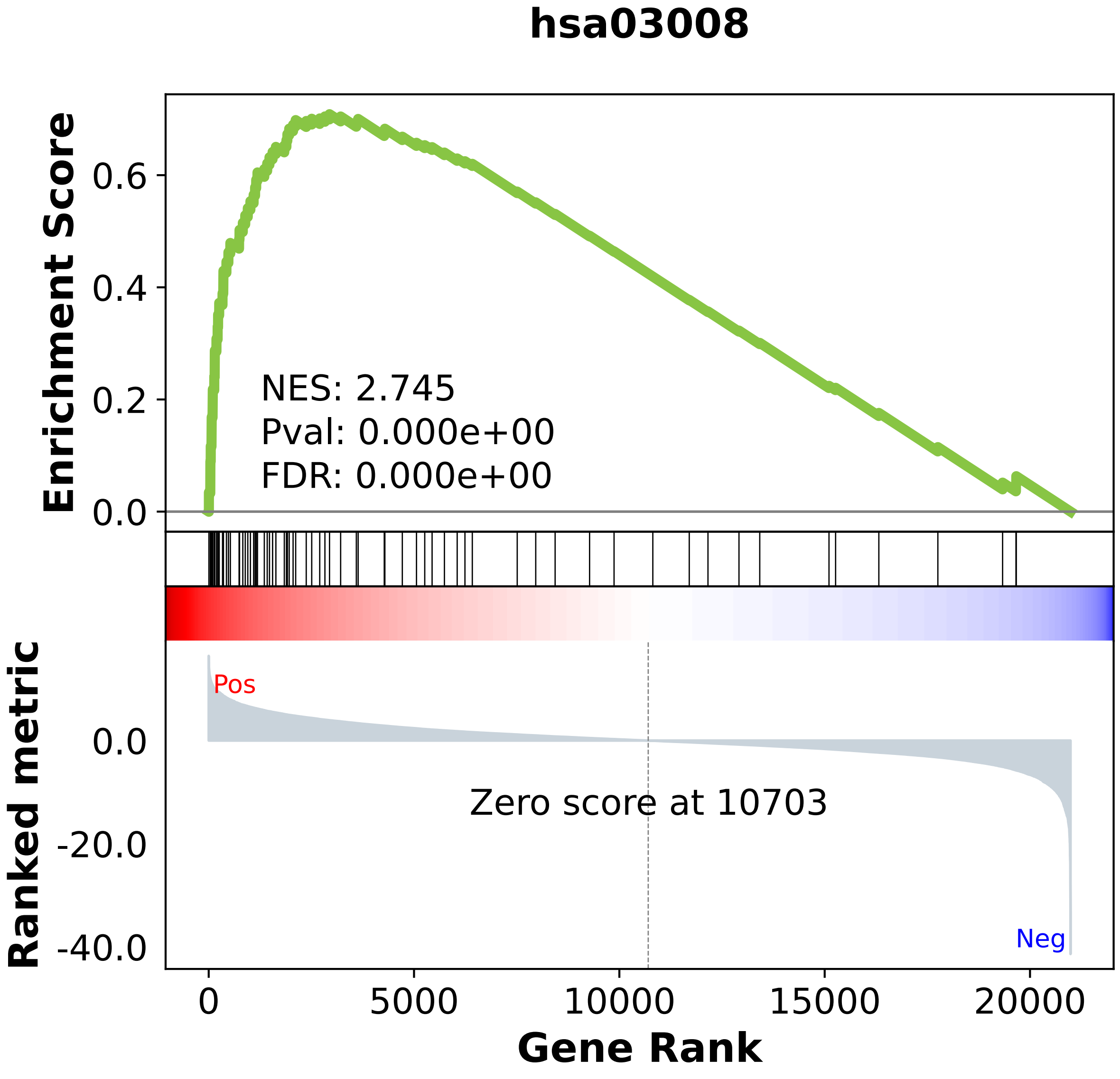

### hsa03010.pdf

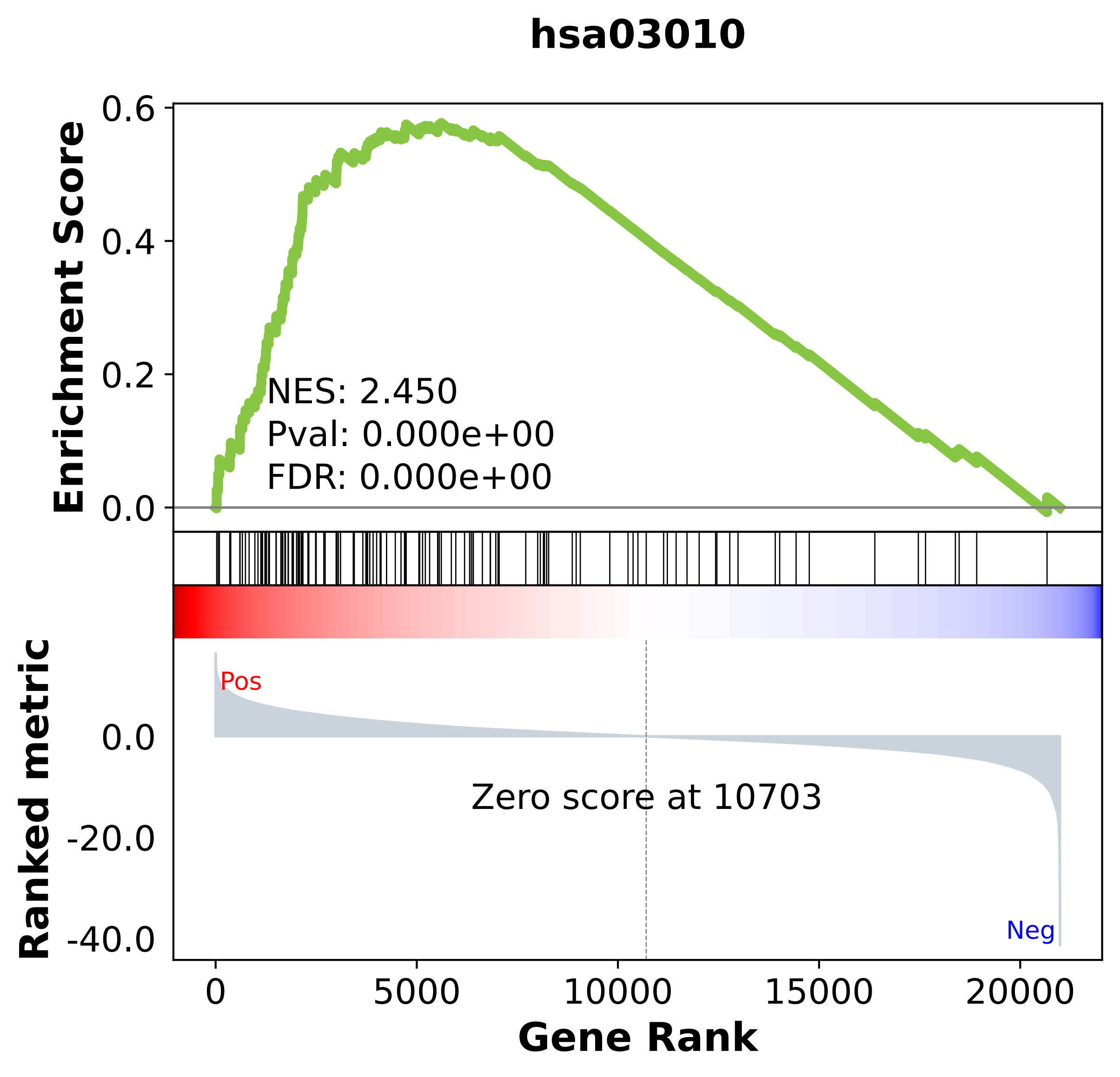

### hsa03013.pdf

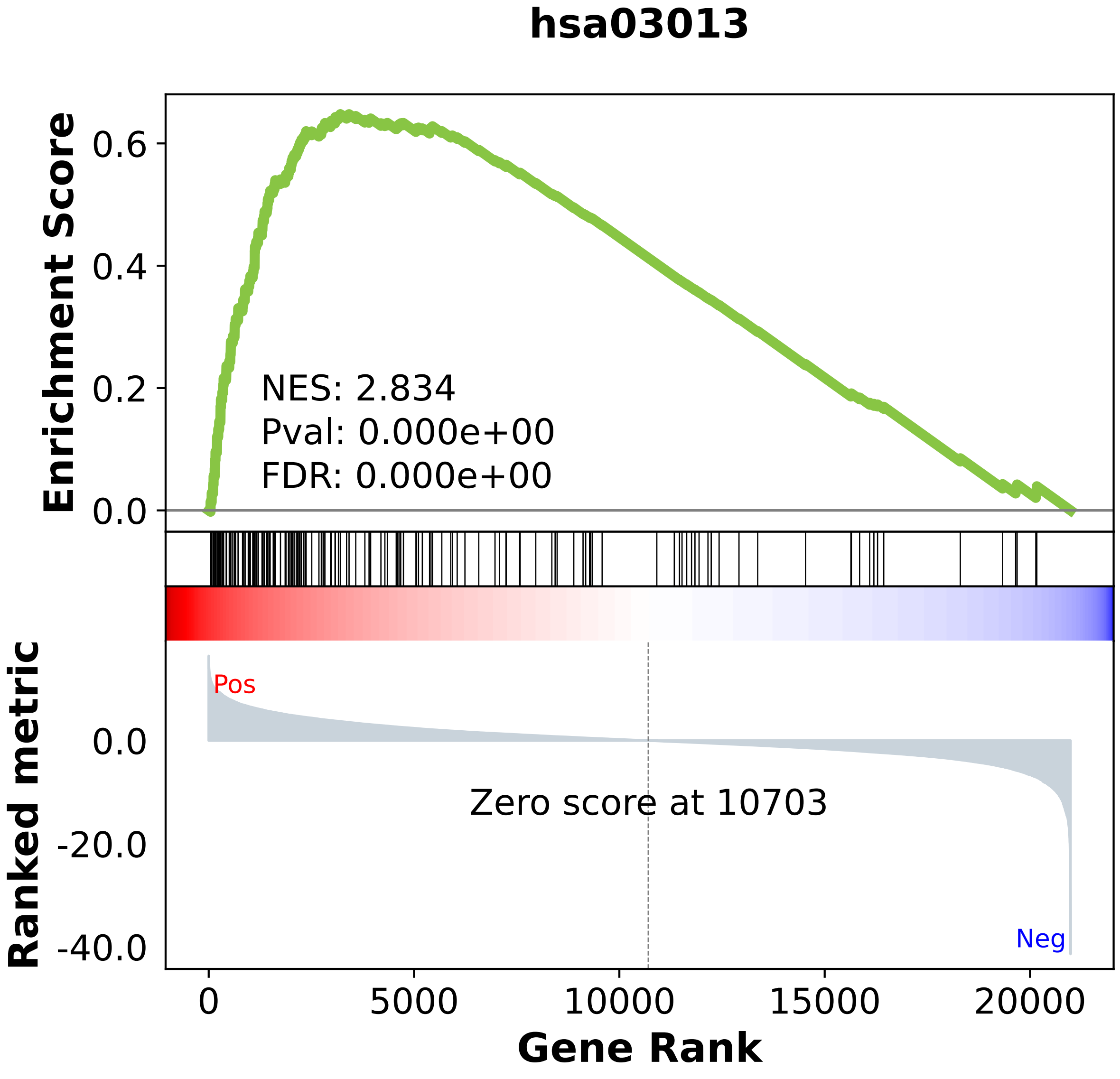

### hsa03015.pdf

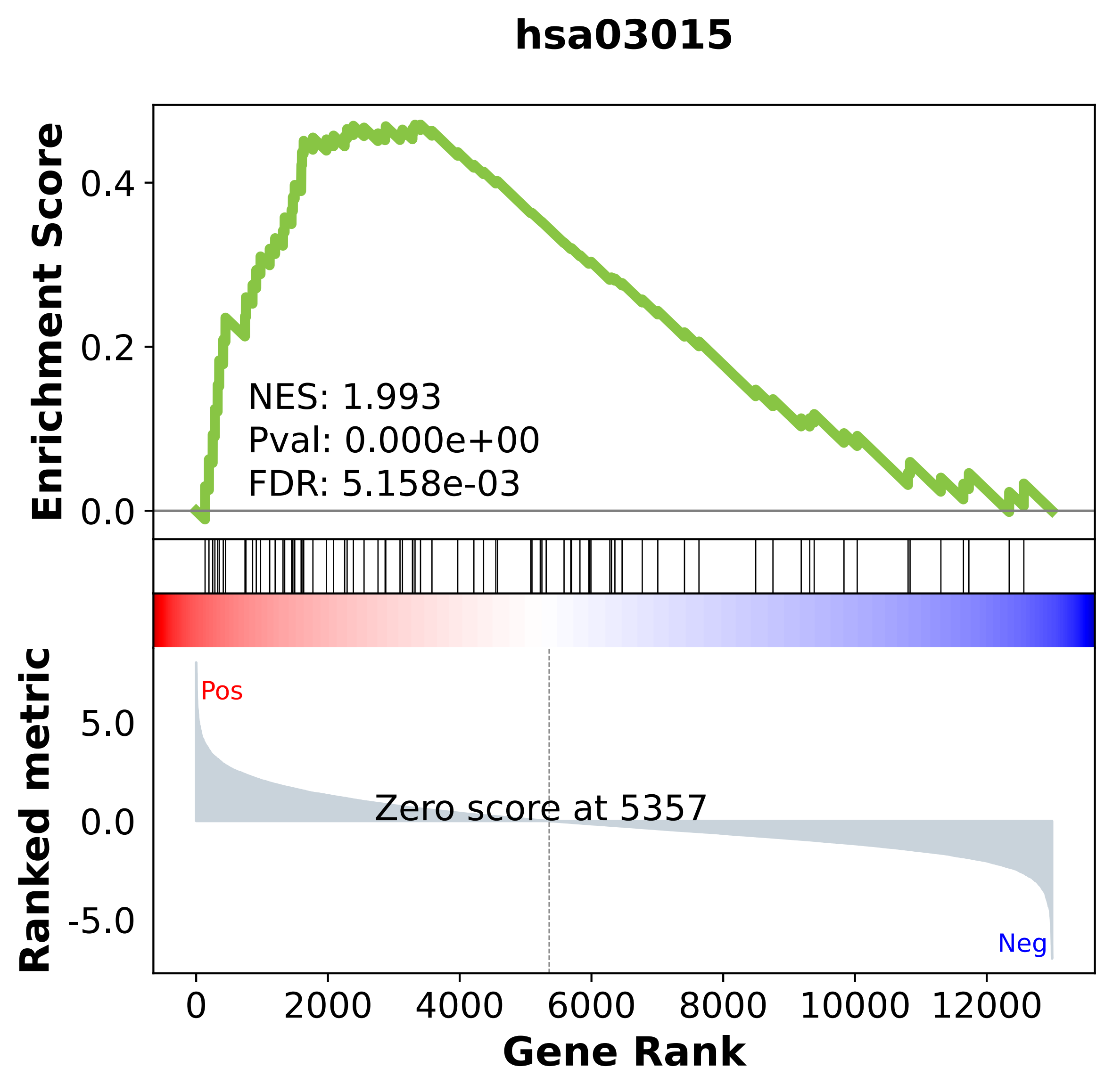

### hsa03018.pdf

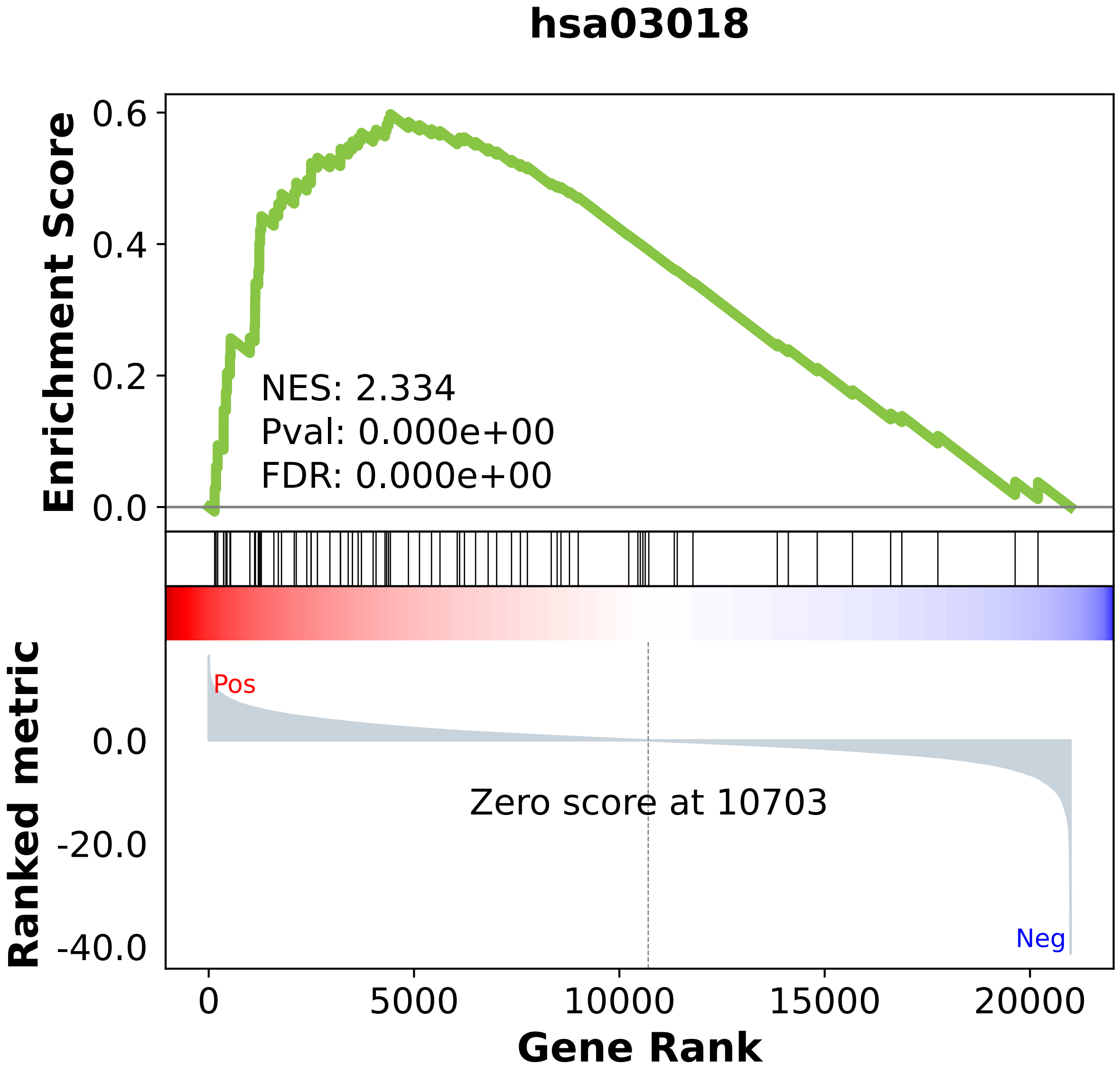

### hsa03020.pdf

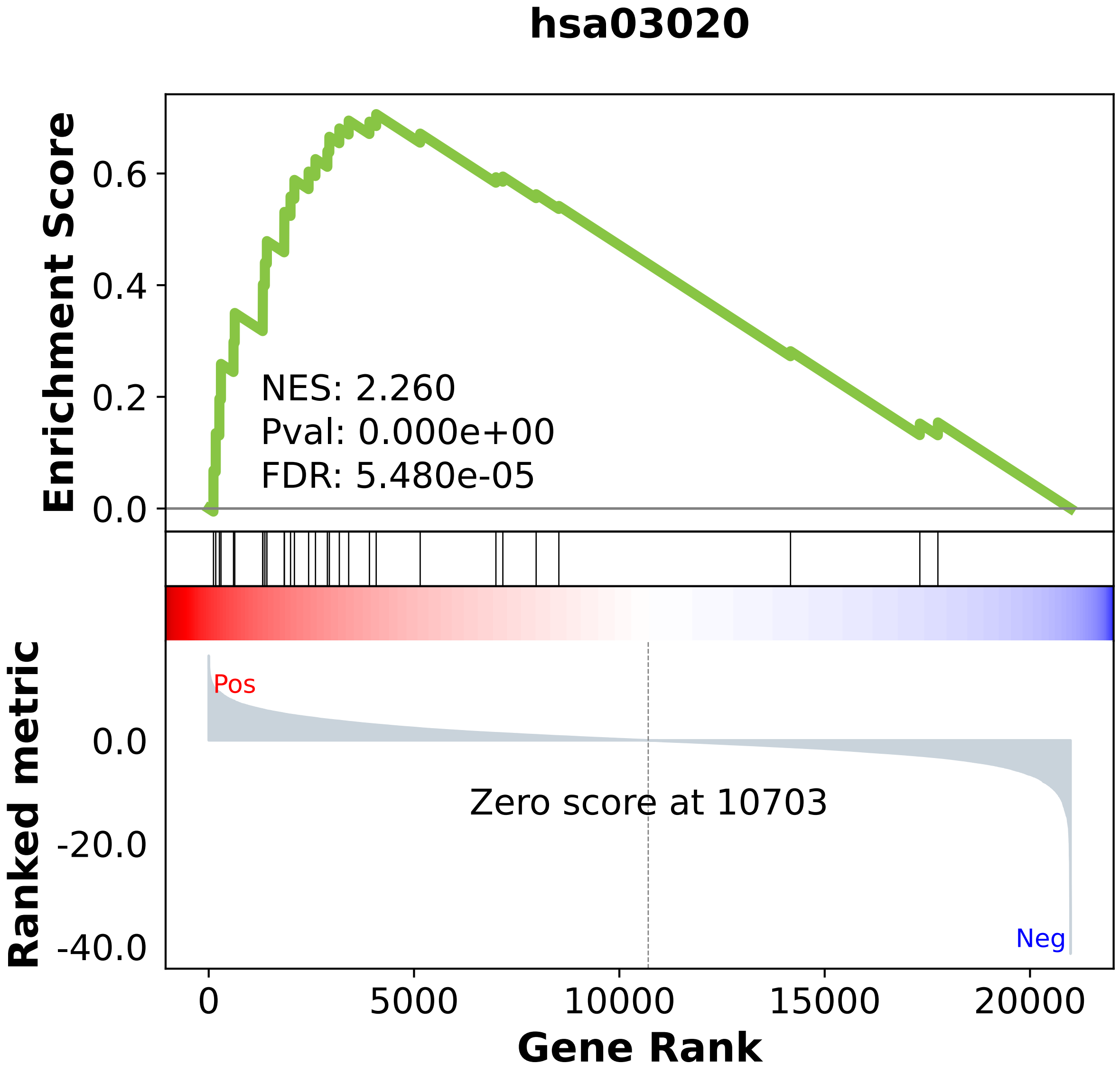

### hsa03022.pdf

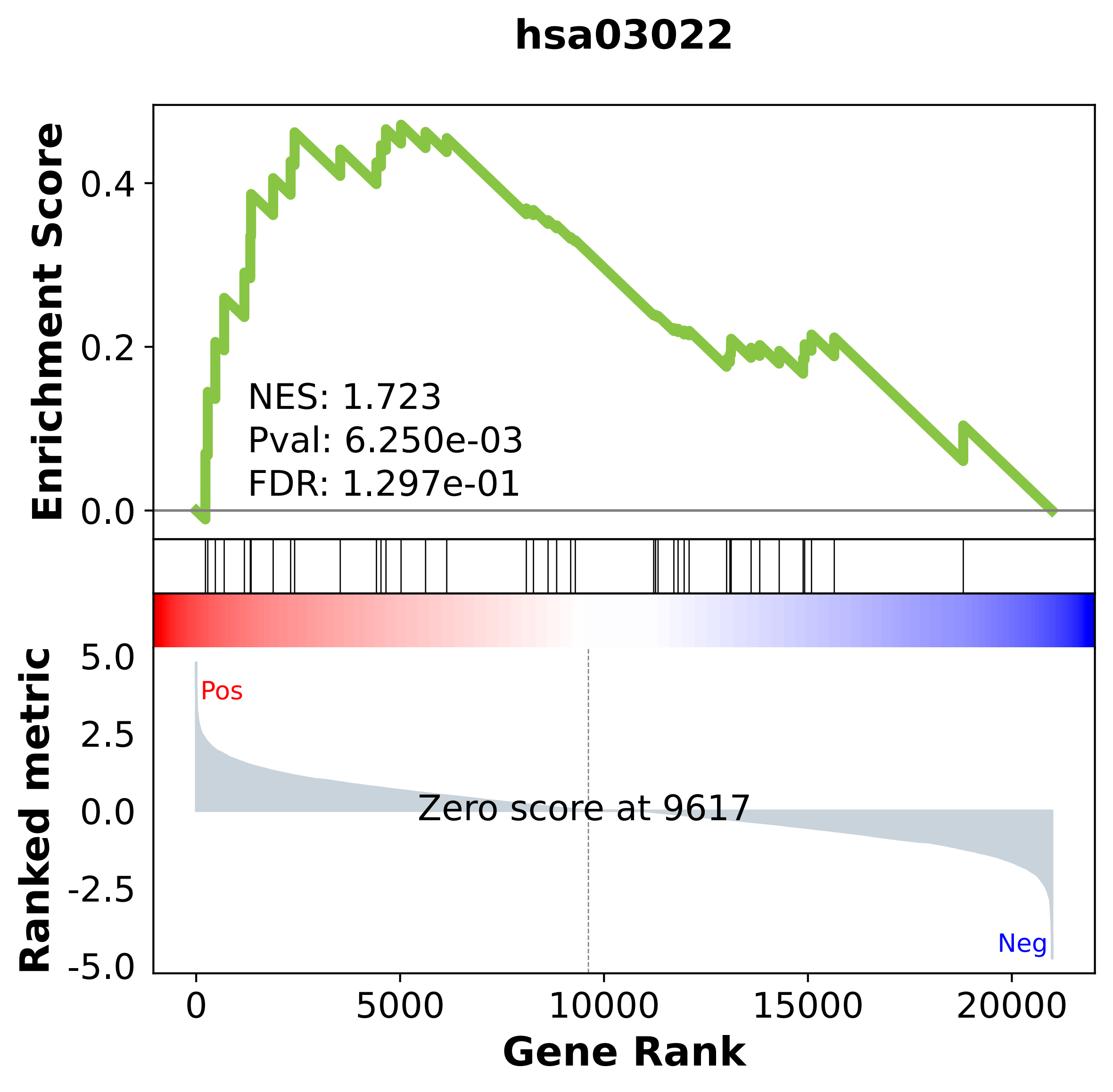

### hsa03030.pdf

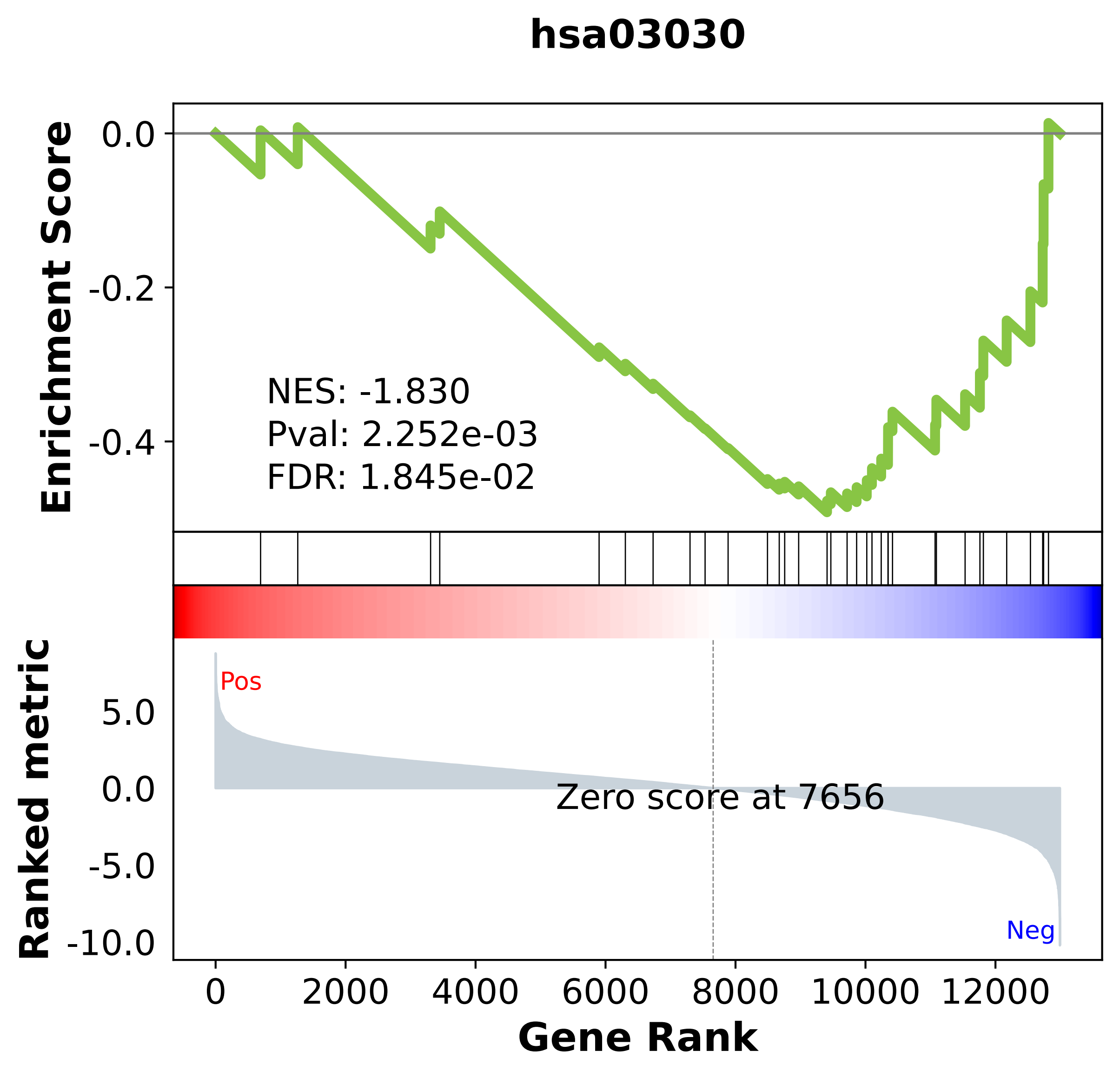

### hsa03040.pdf

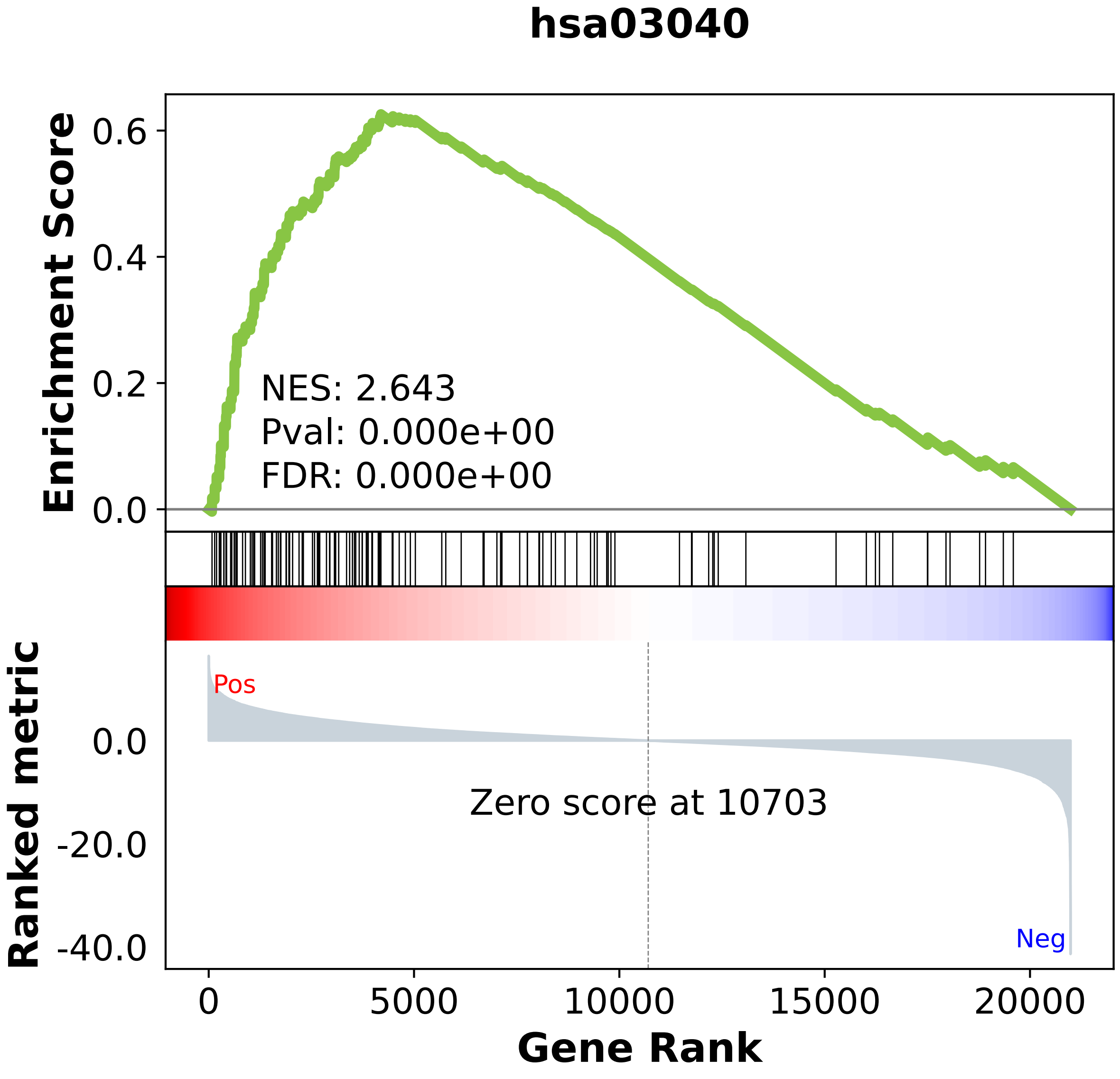

### hsa03050.pdf

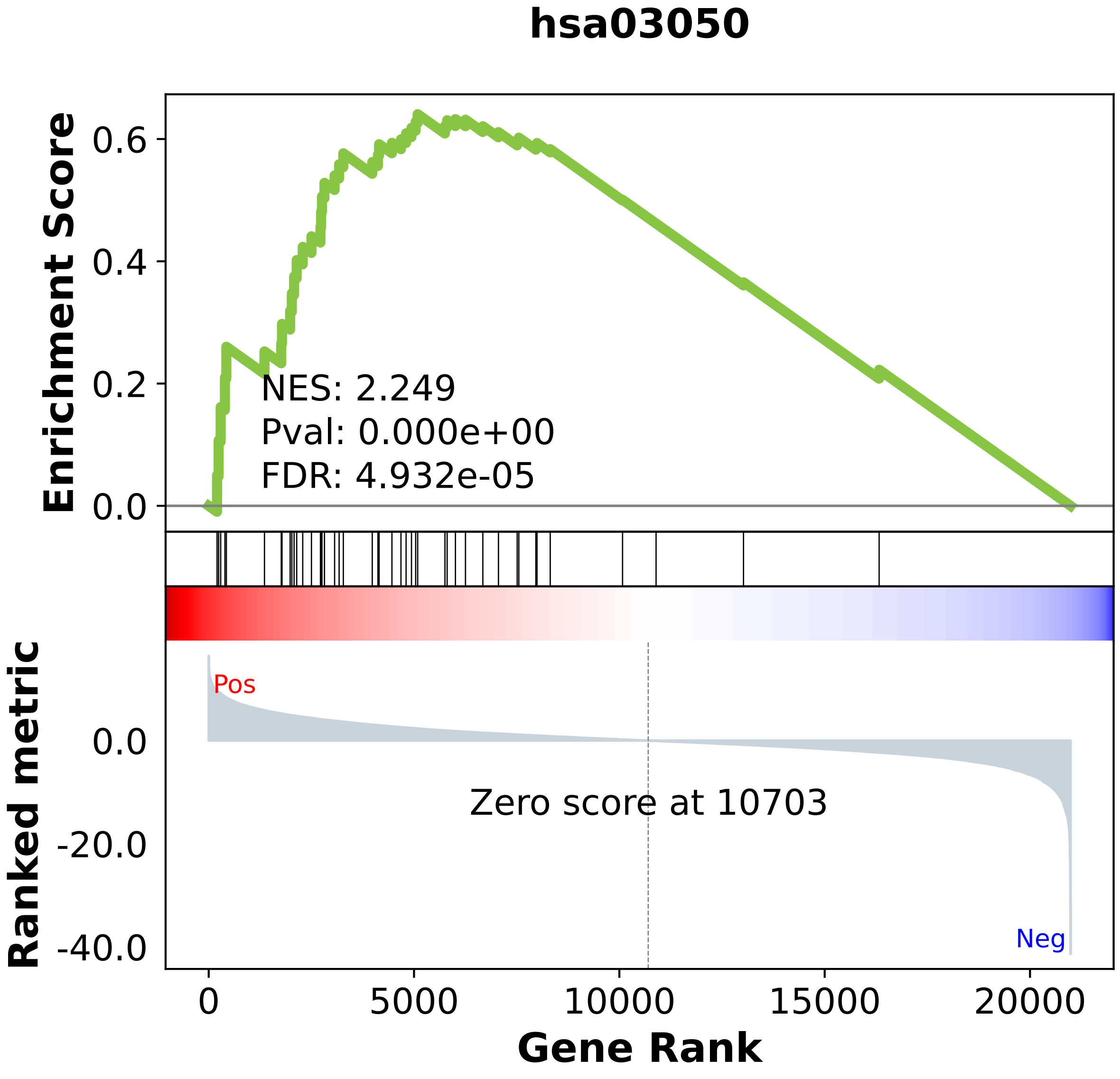

### hsa03060.pdf

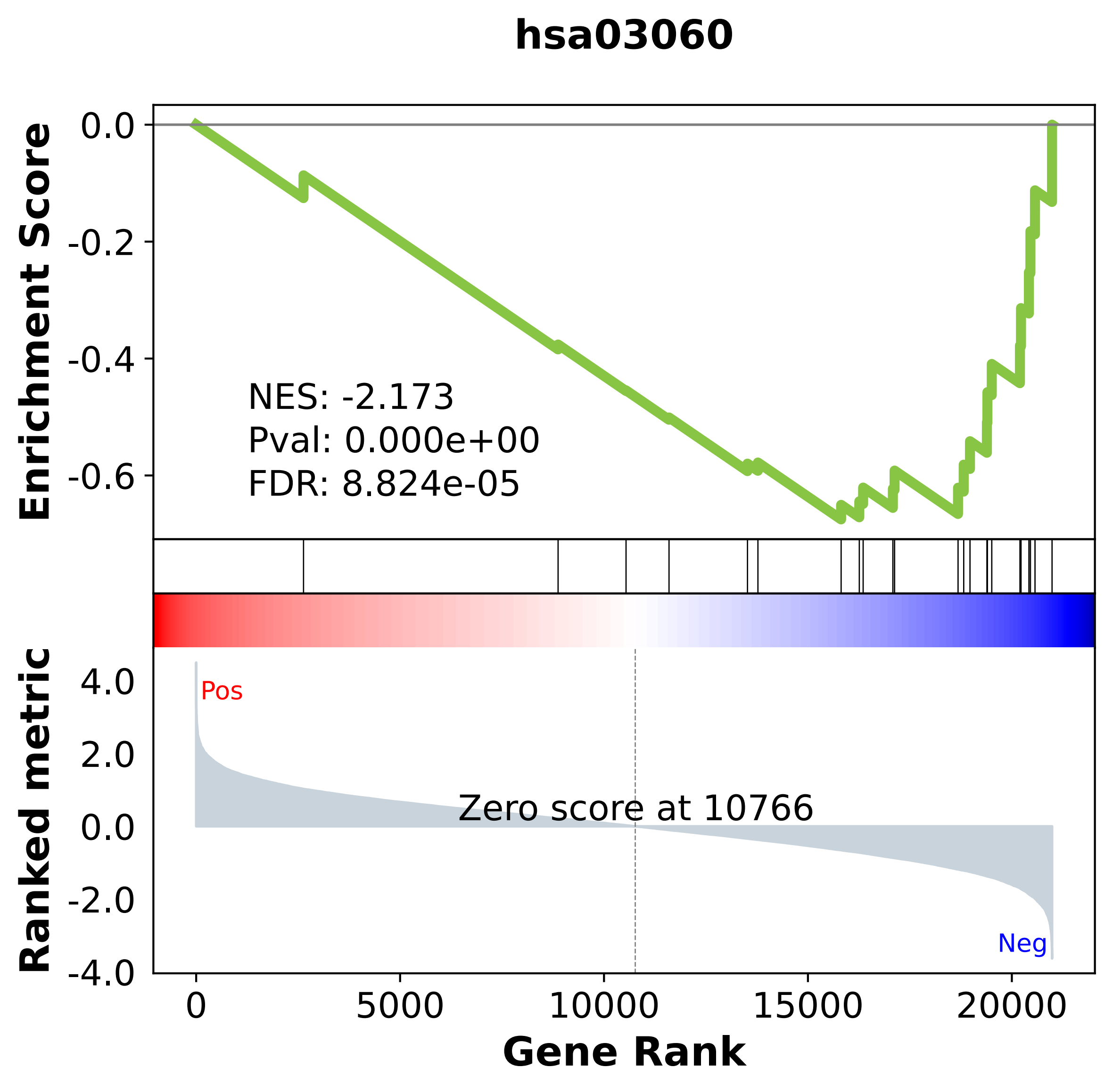

### hsa03320.pdf

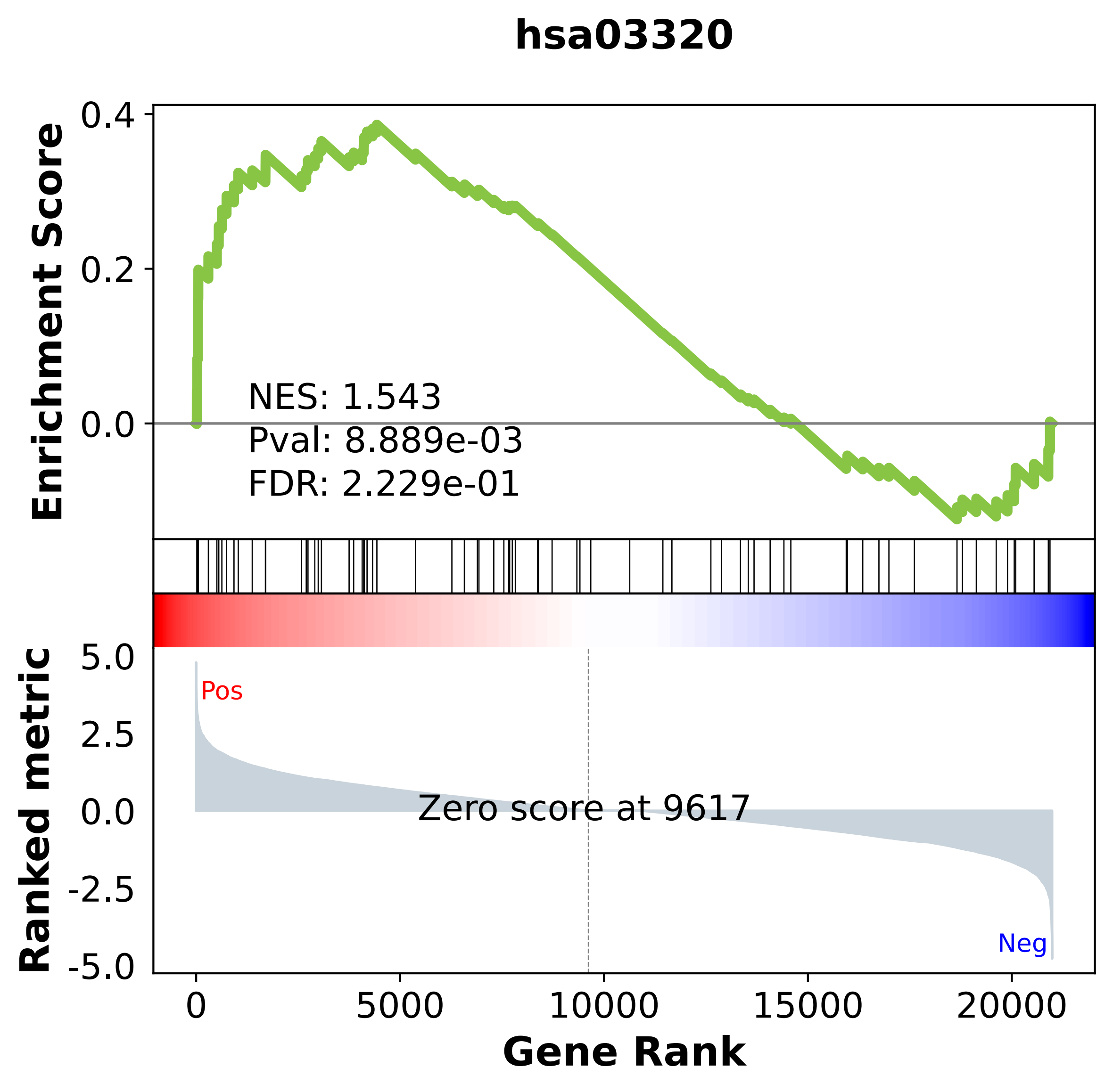

### hsa03410.pdf

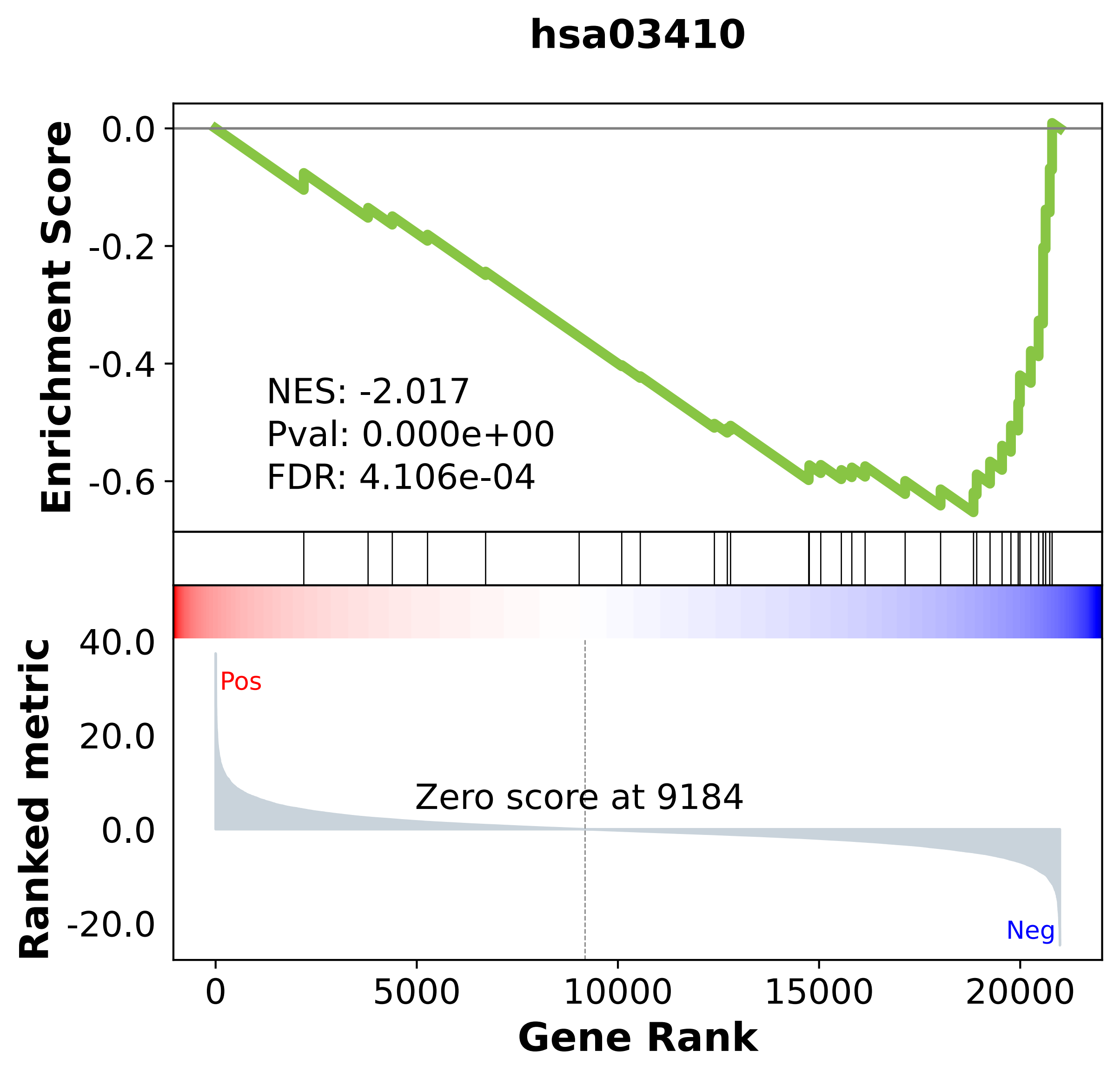

### hsa03420.pdf

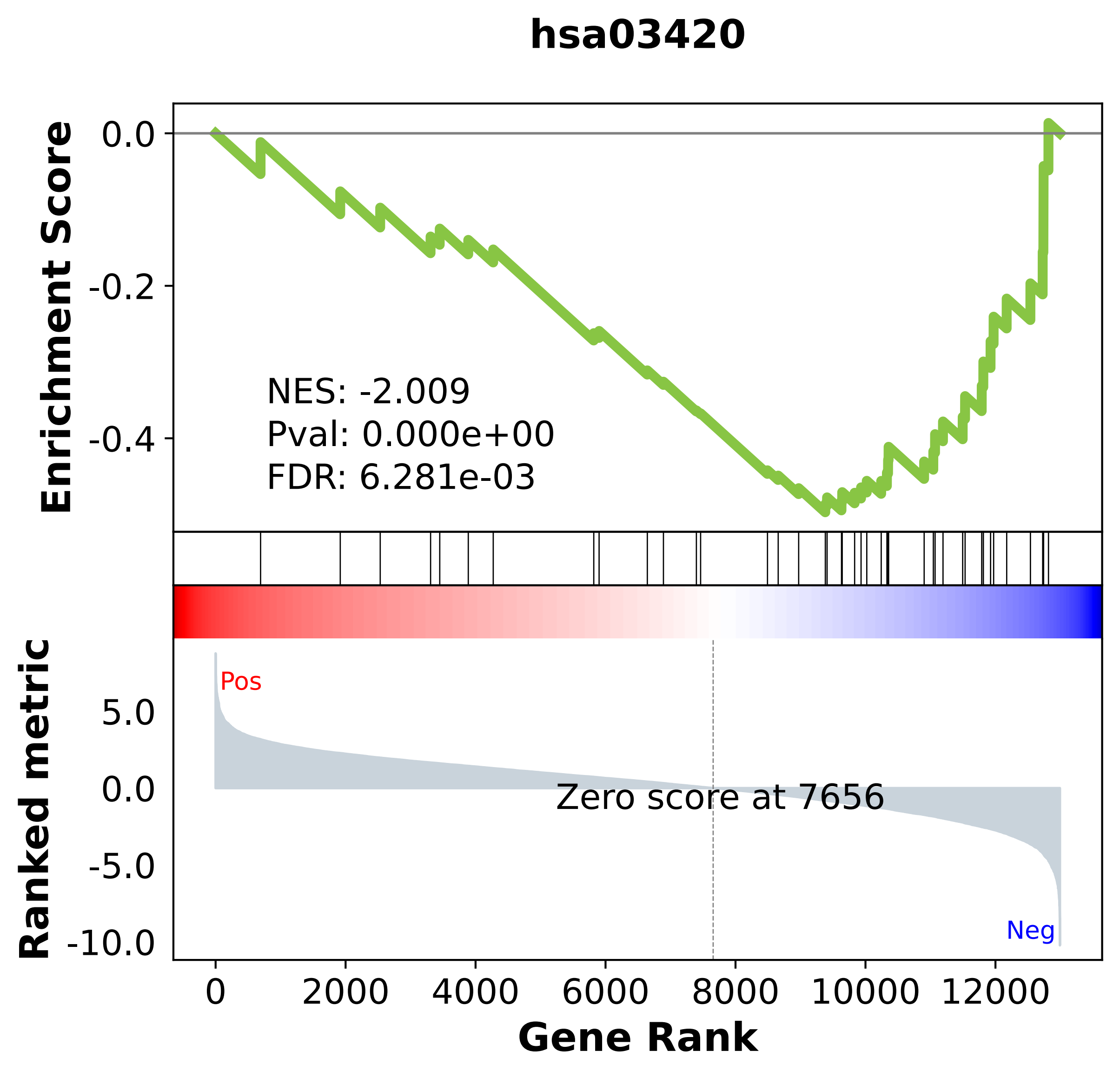

### hsa03430.pdf

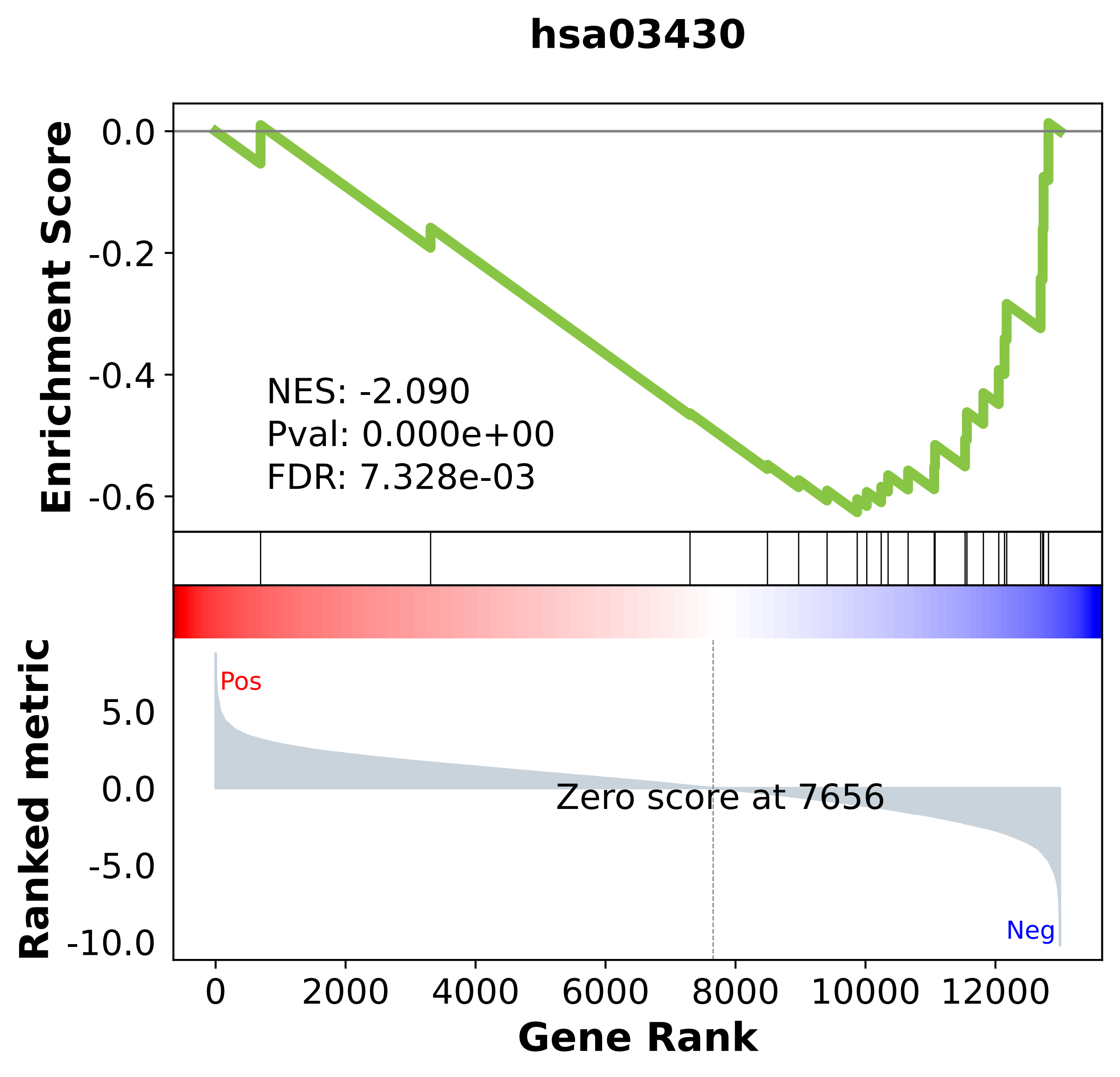

### hsa03440.pdf

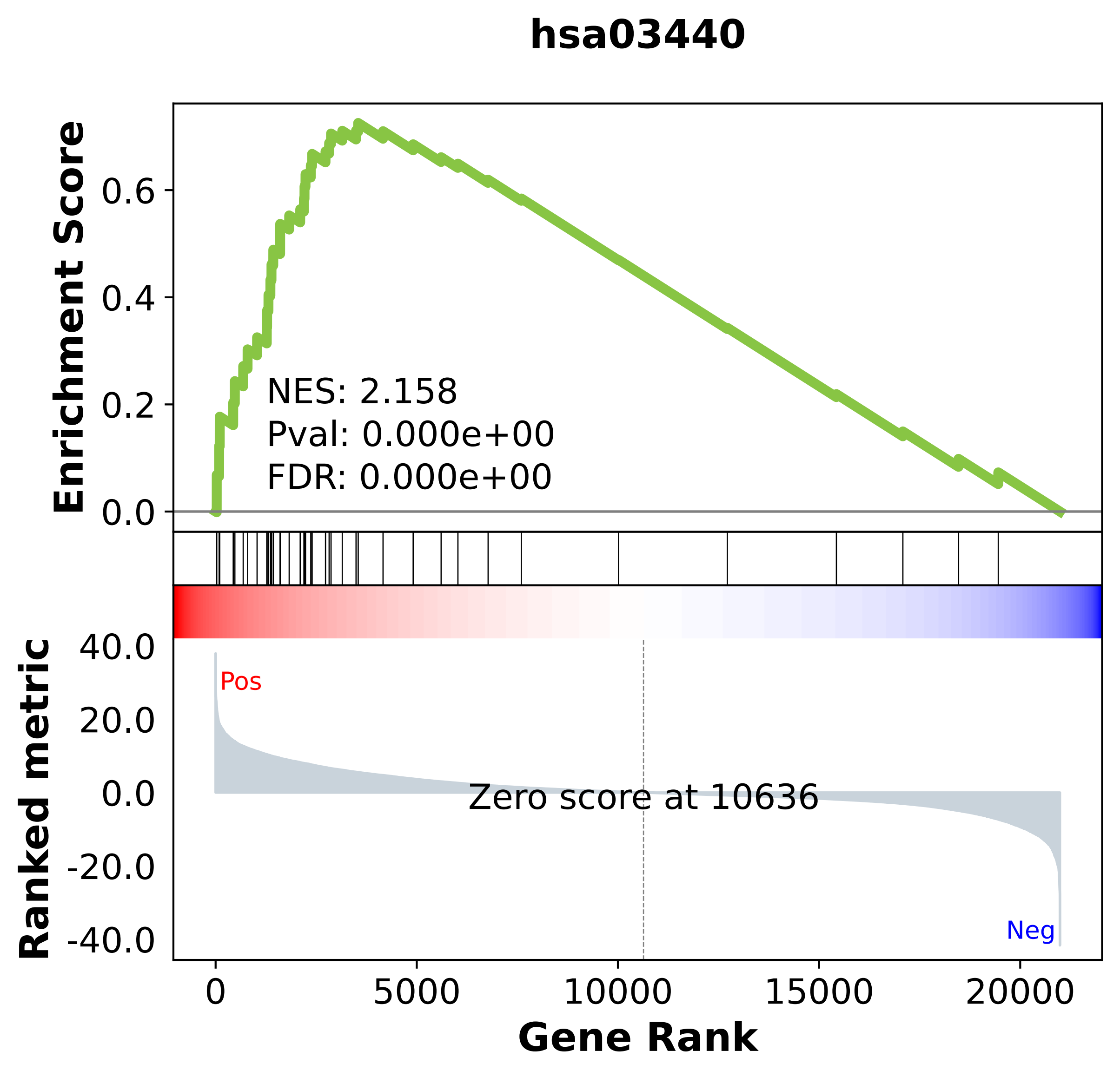

### hsa03460.pdf

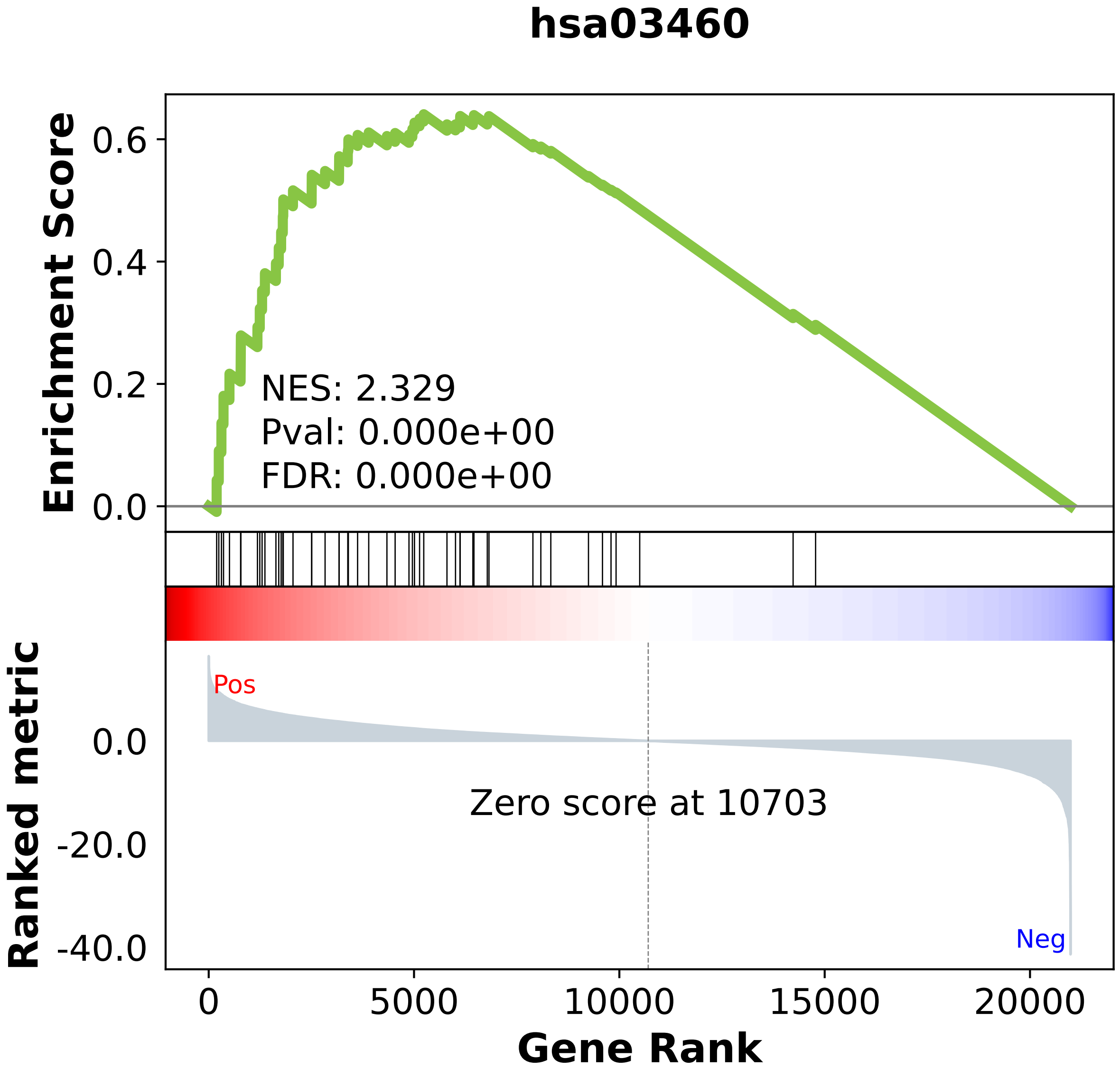

### hsa04012.pdf

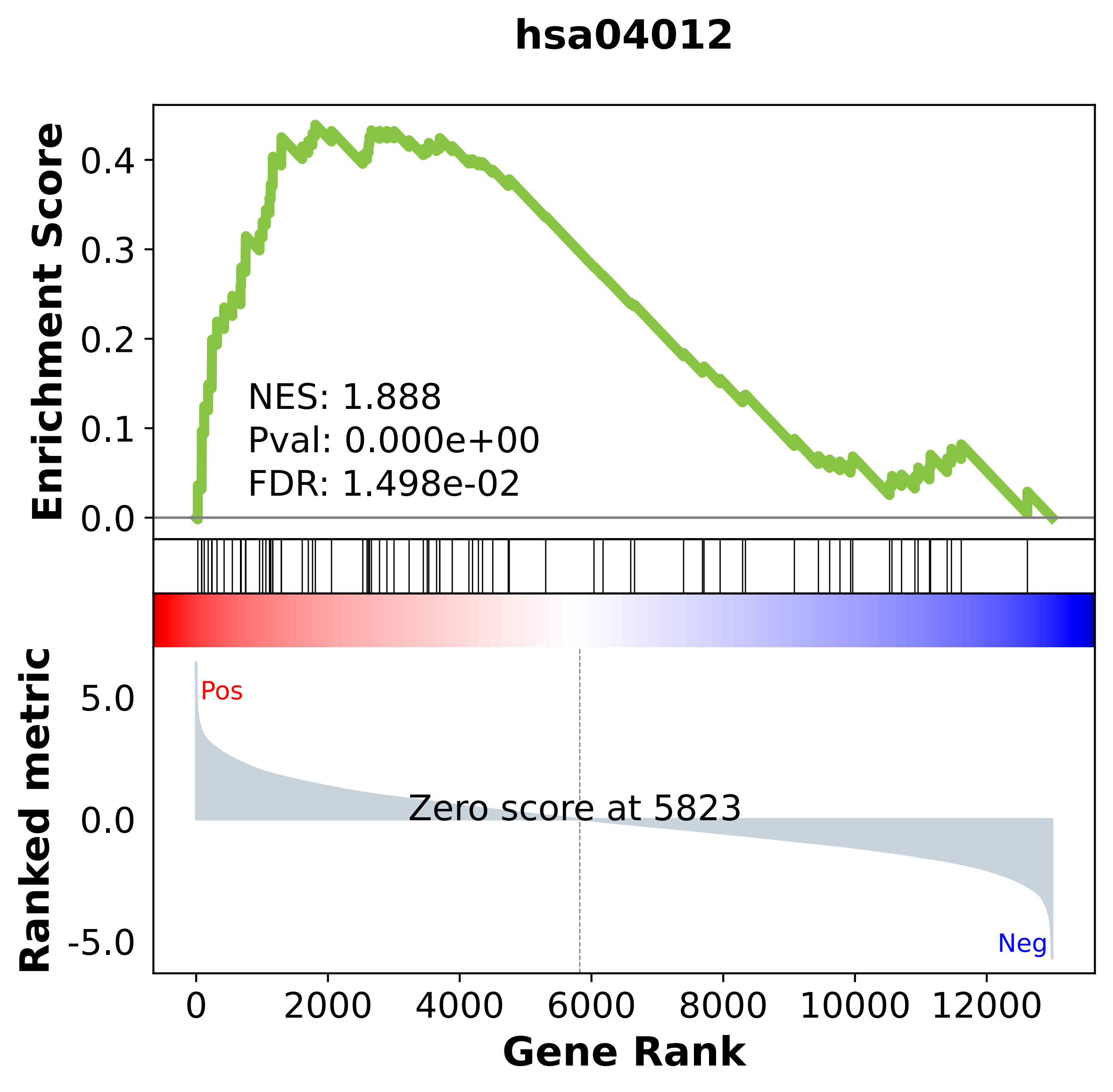

### hsa04020.pdf

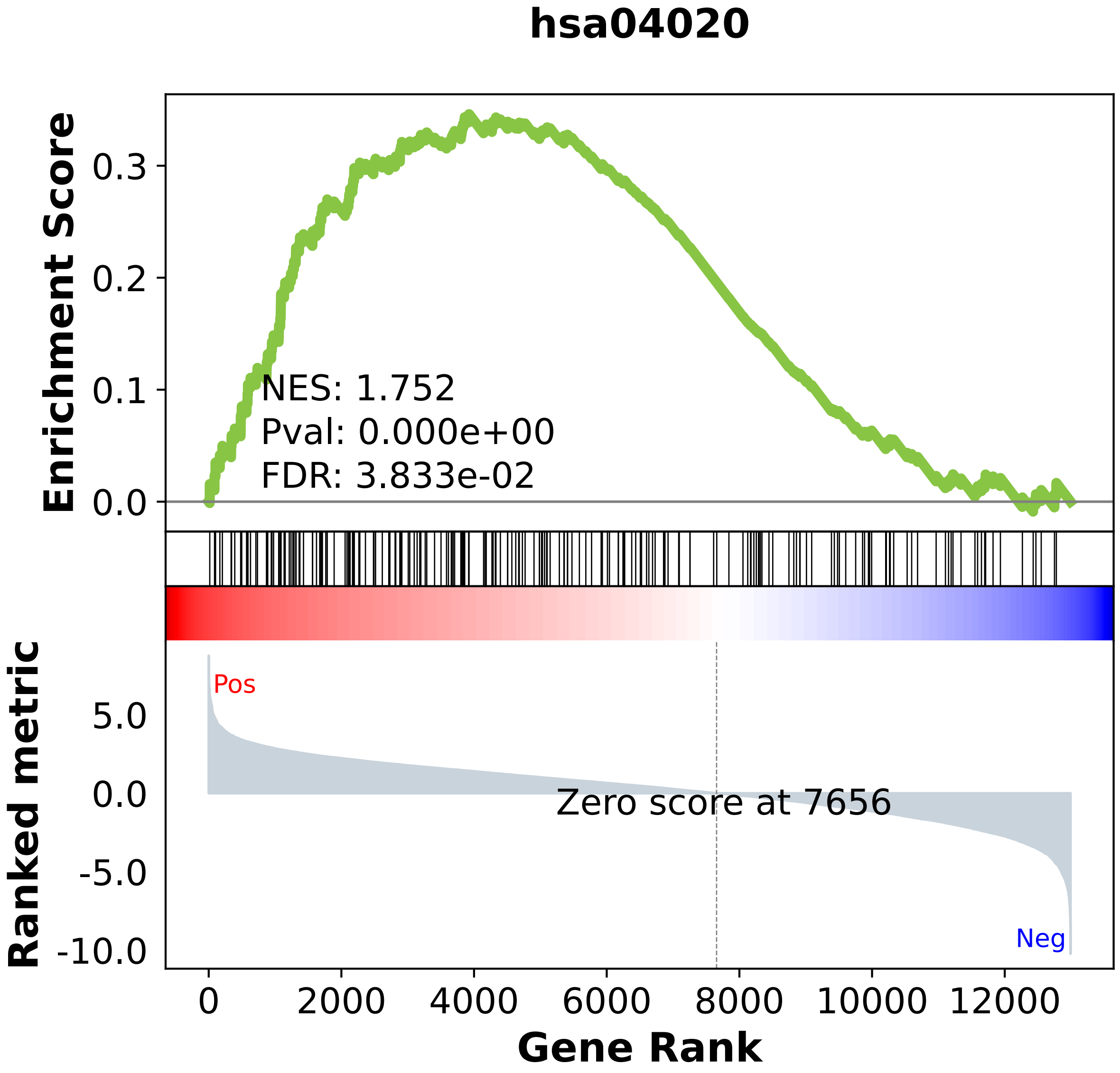

### hsa04060.pdf

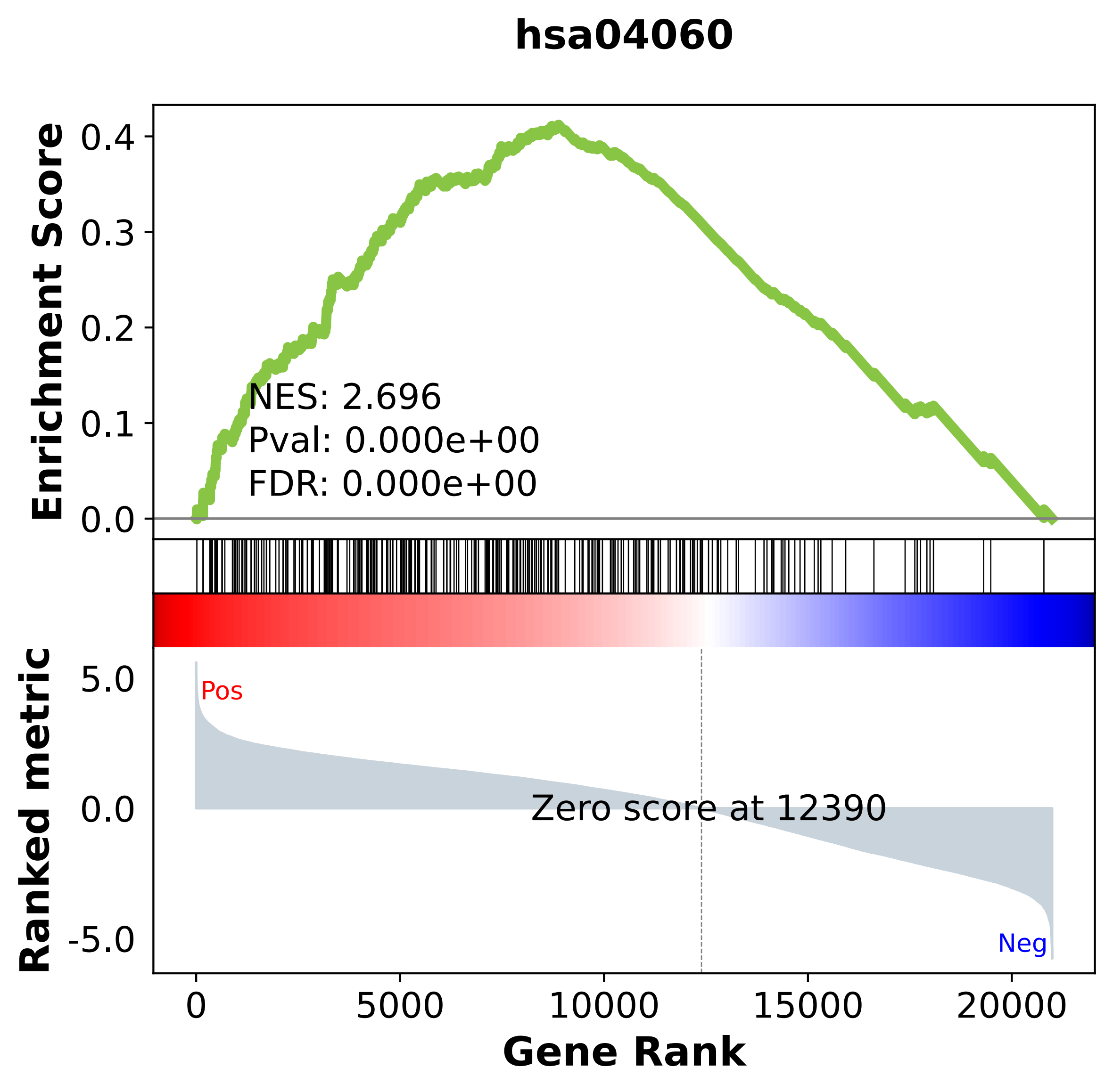

### hsa04061.pdf

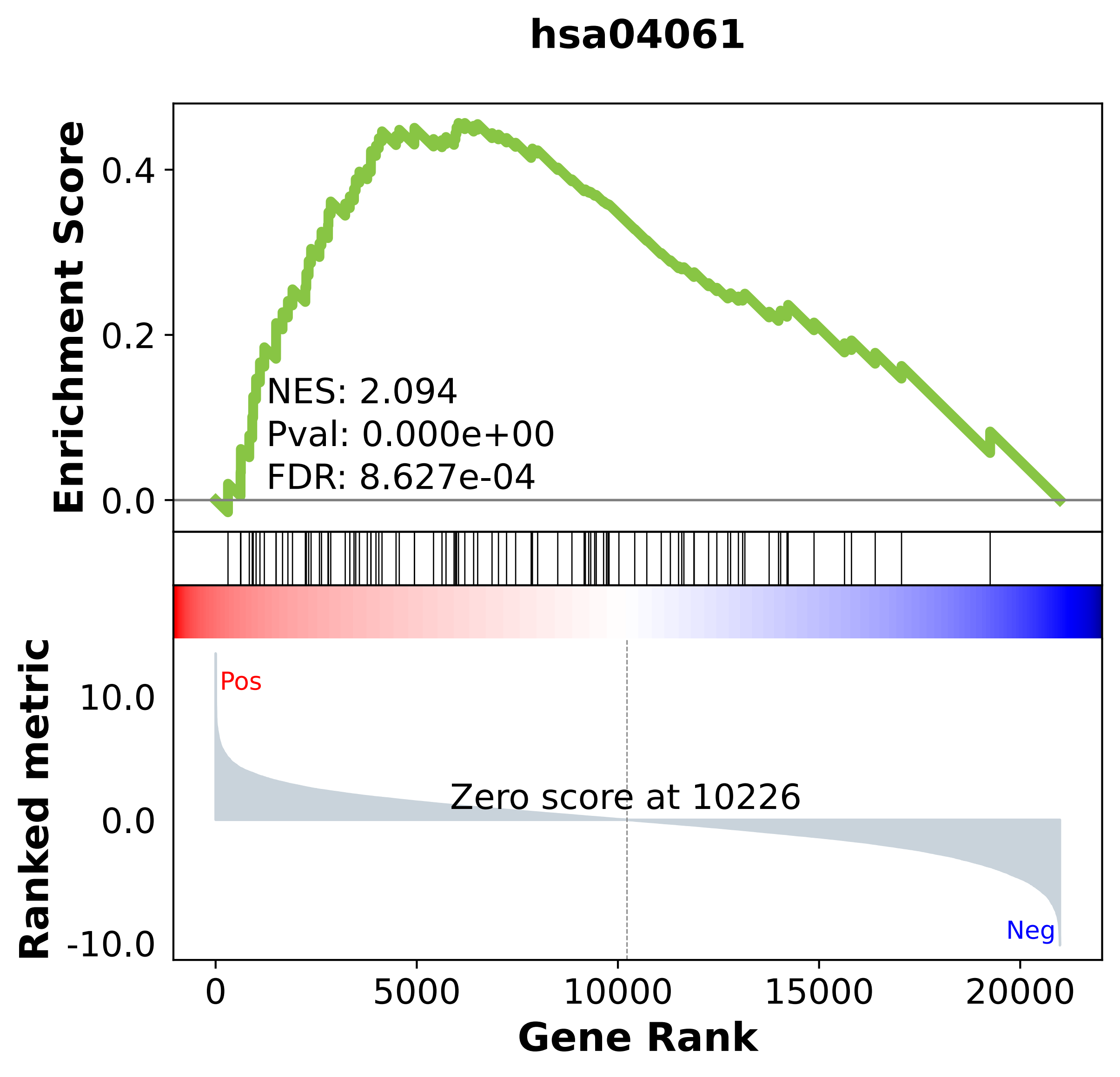

### hsa04062.pdf

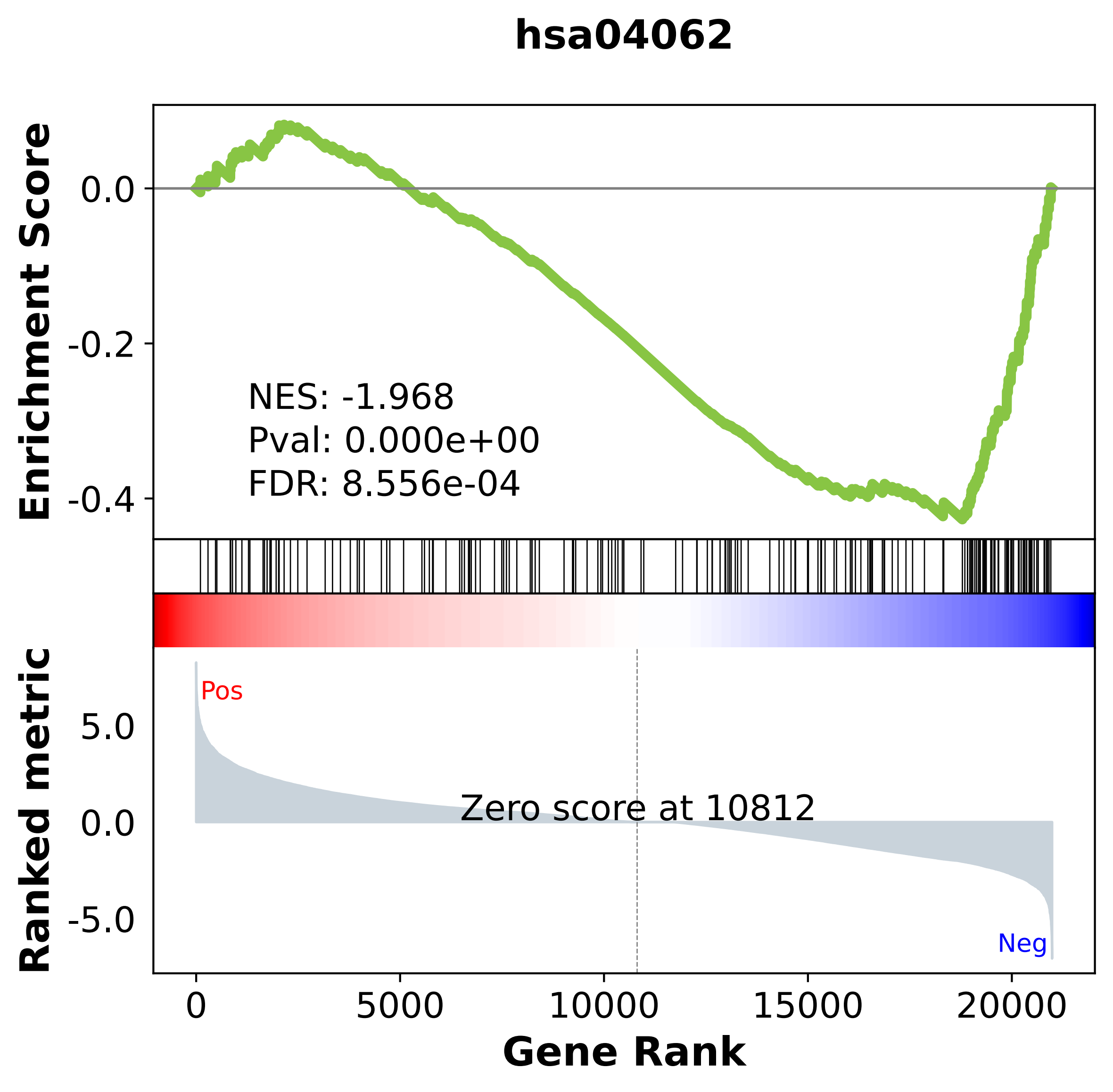

### hsa04064.pdf

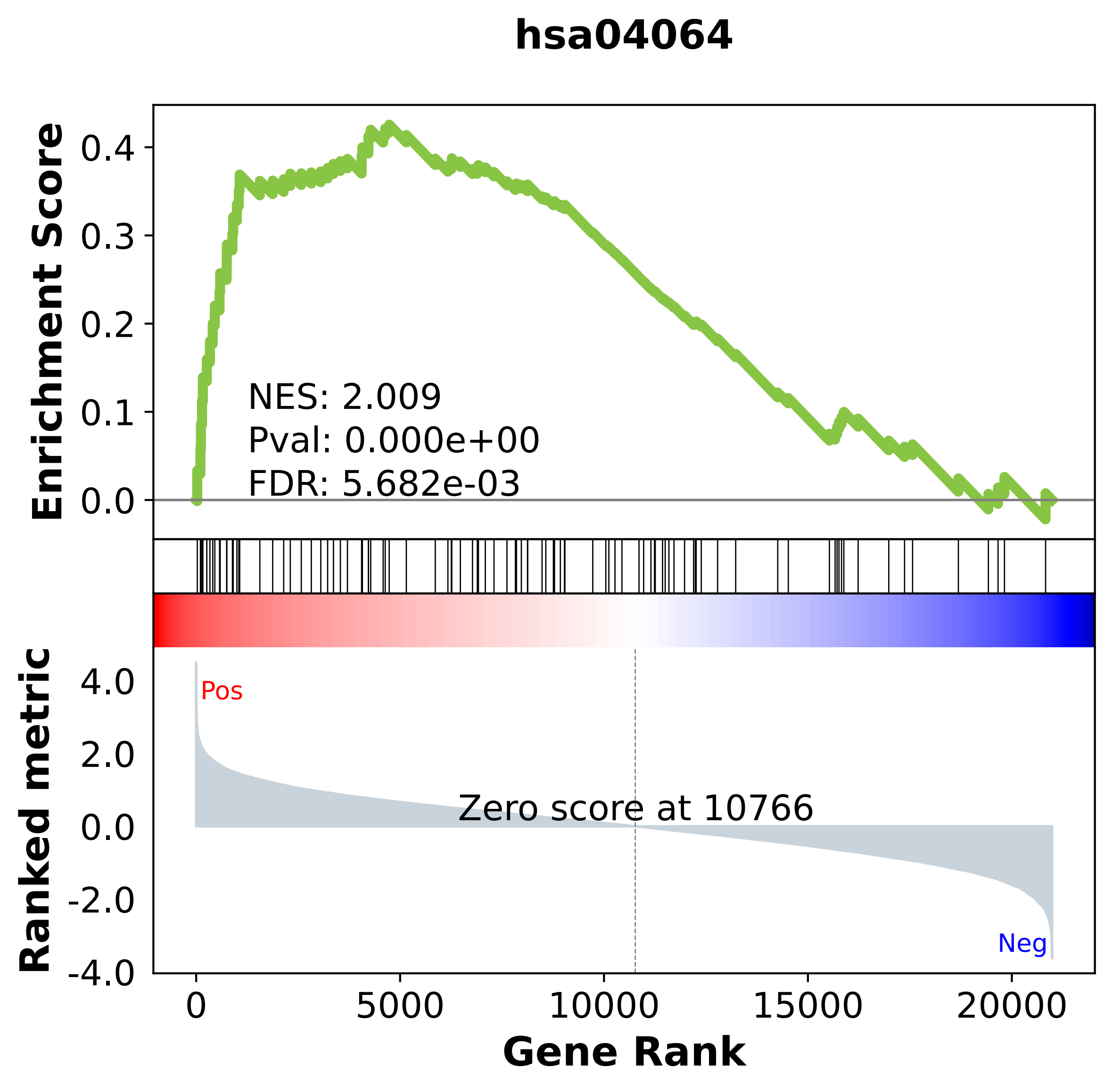

### hsa04066.pdf

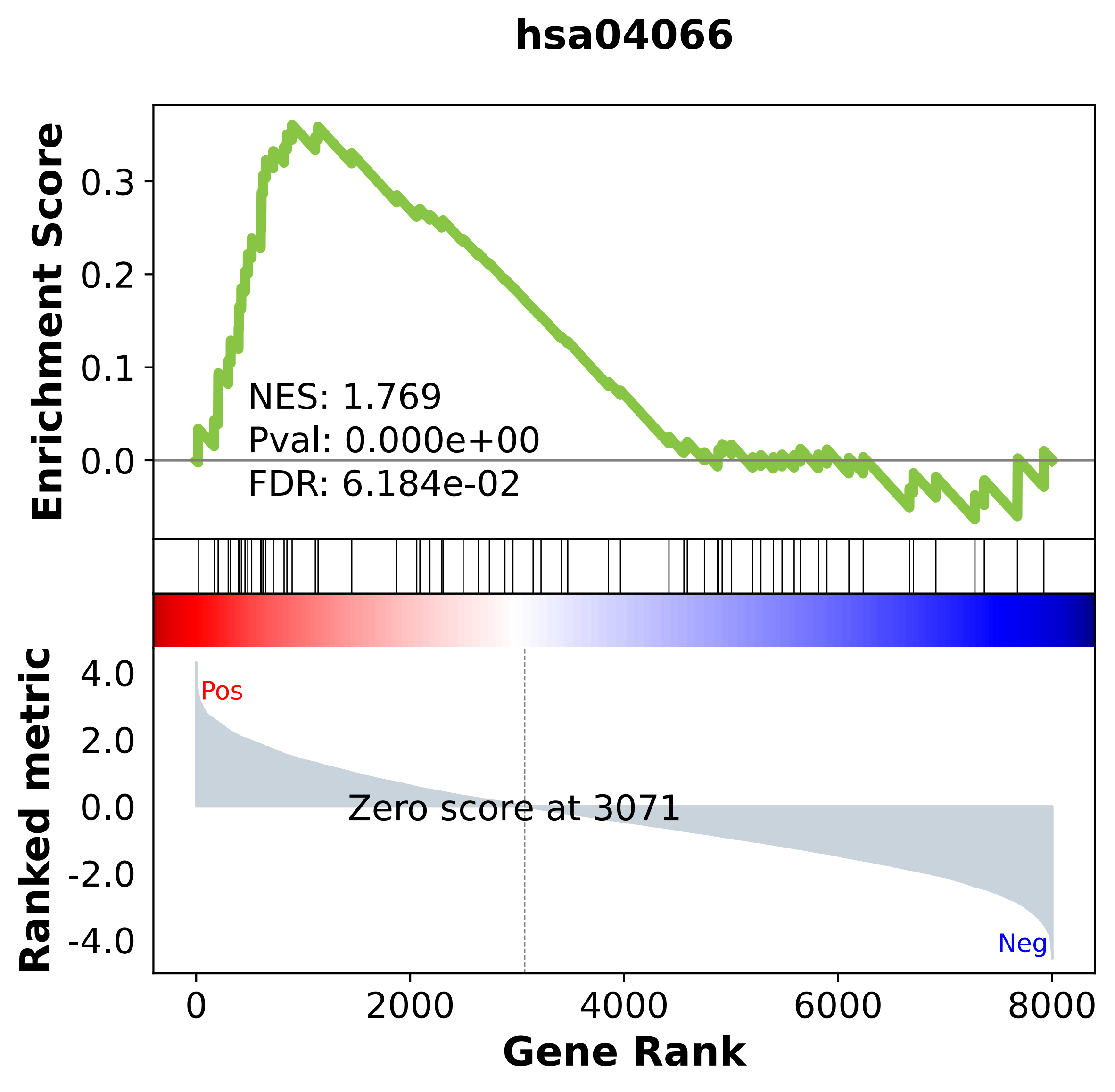

### hsa04068.pdf

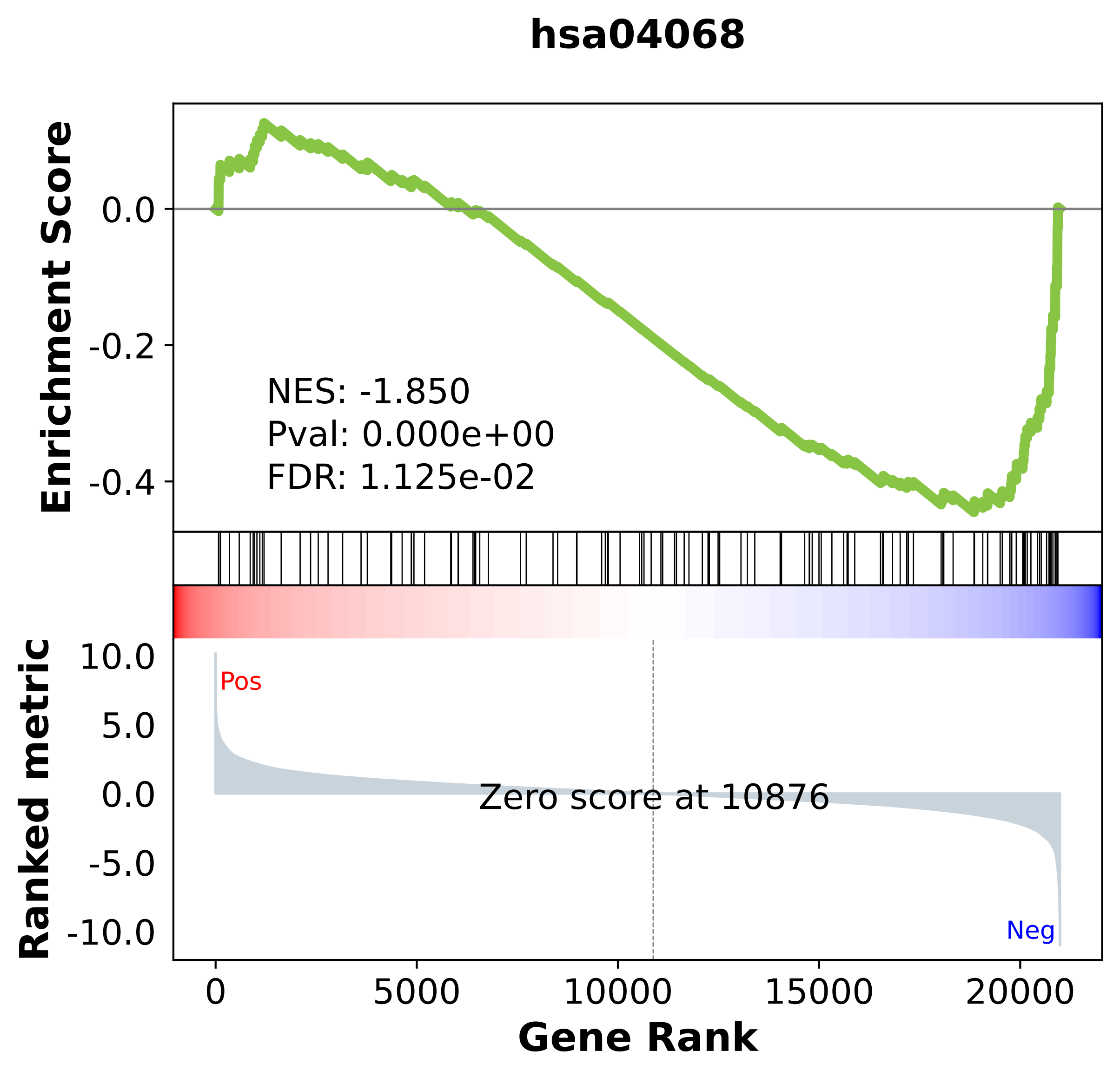

### hsa04070.pdf

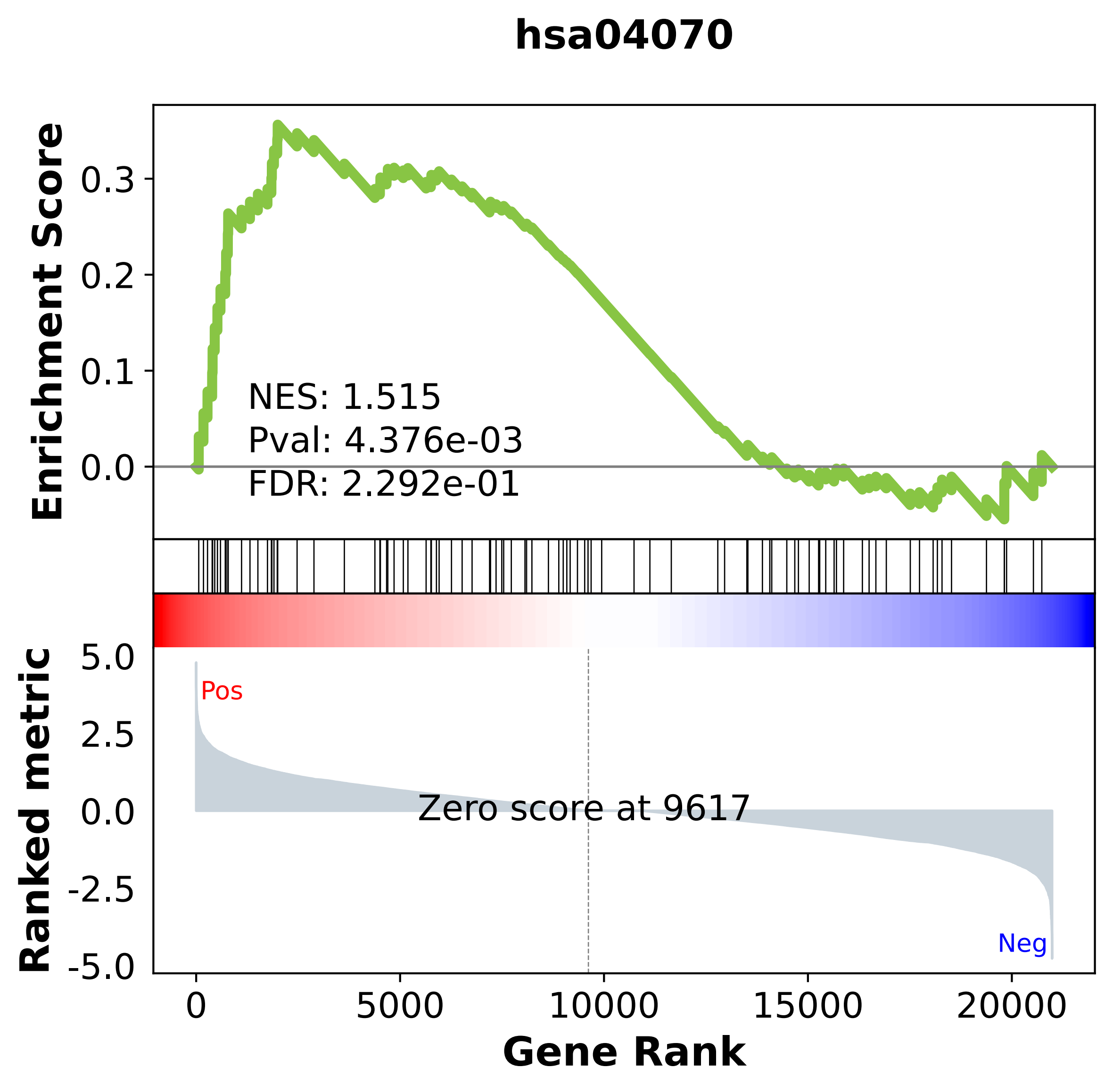

### hsa04080.pdf

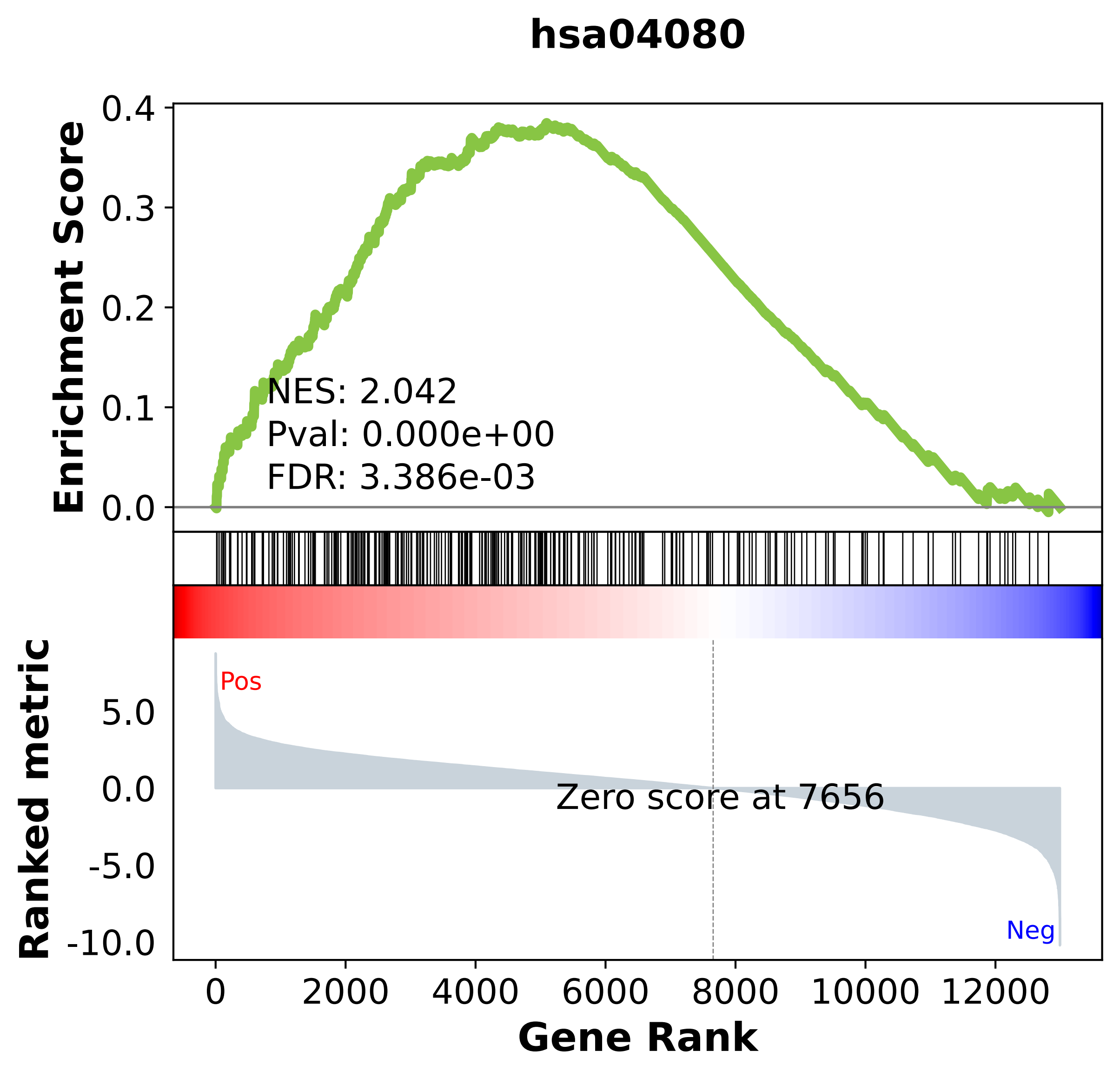

### hsa04110.pdf

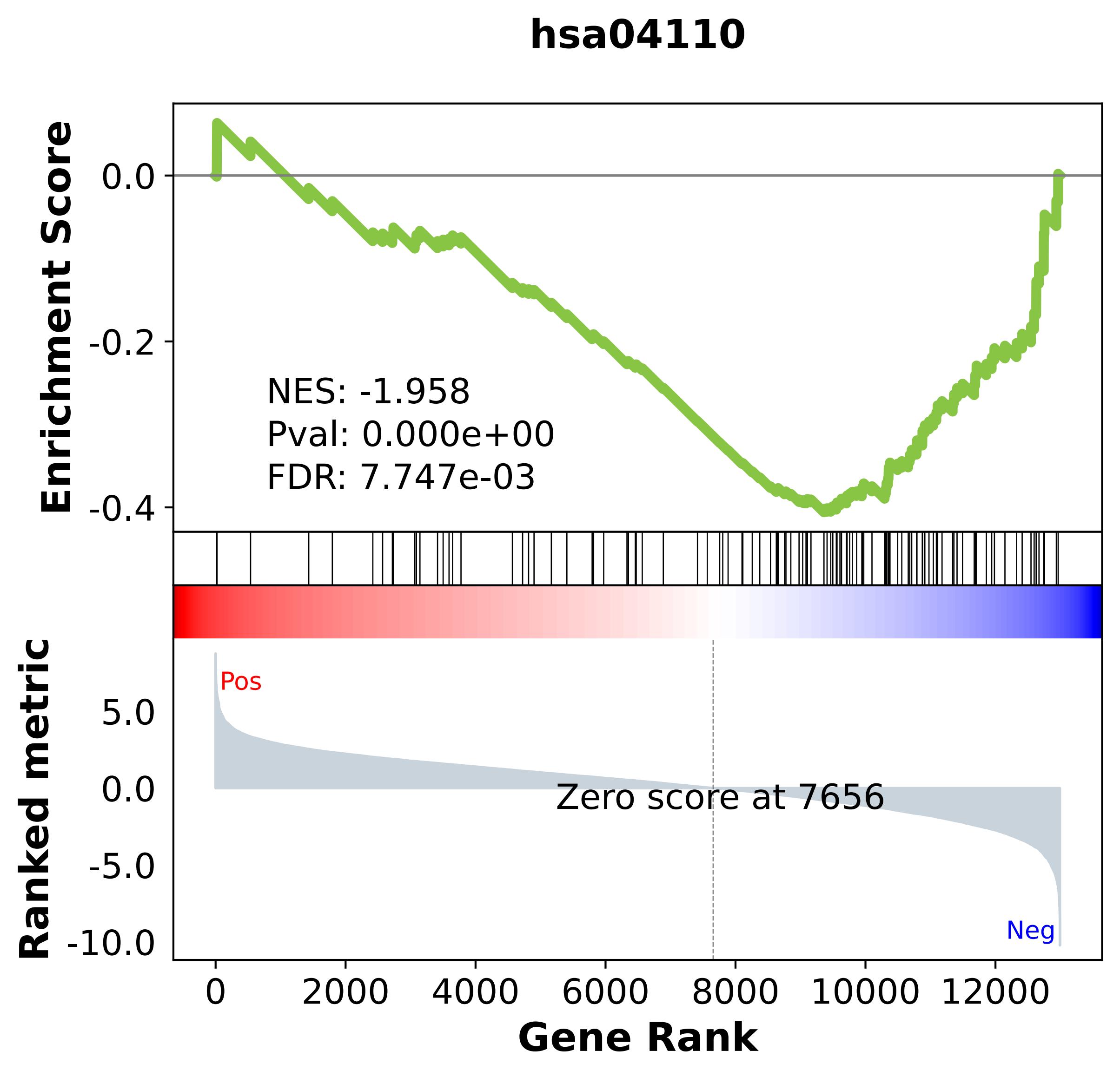

### hsa04114.pdf

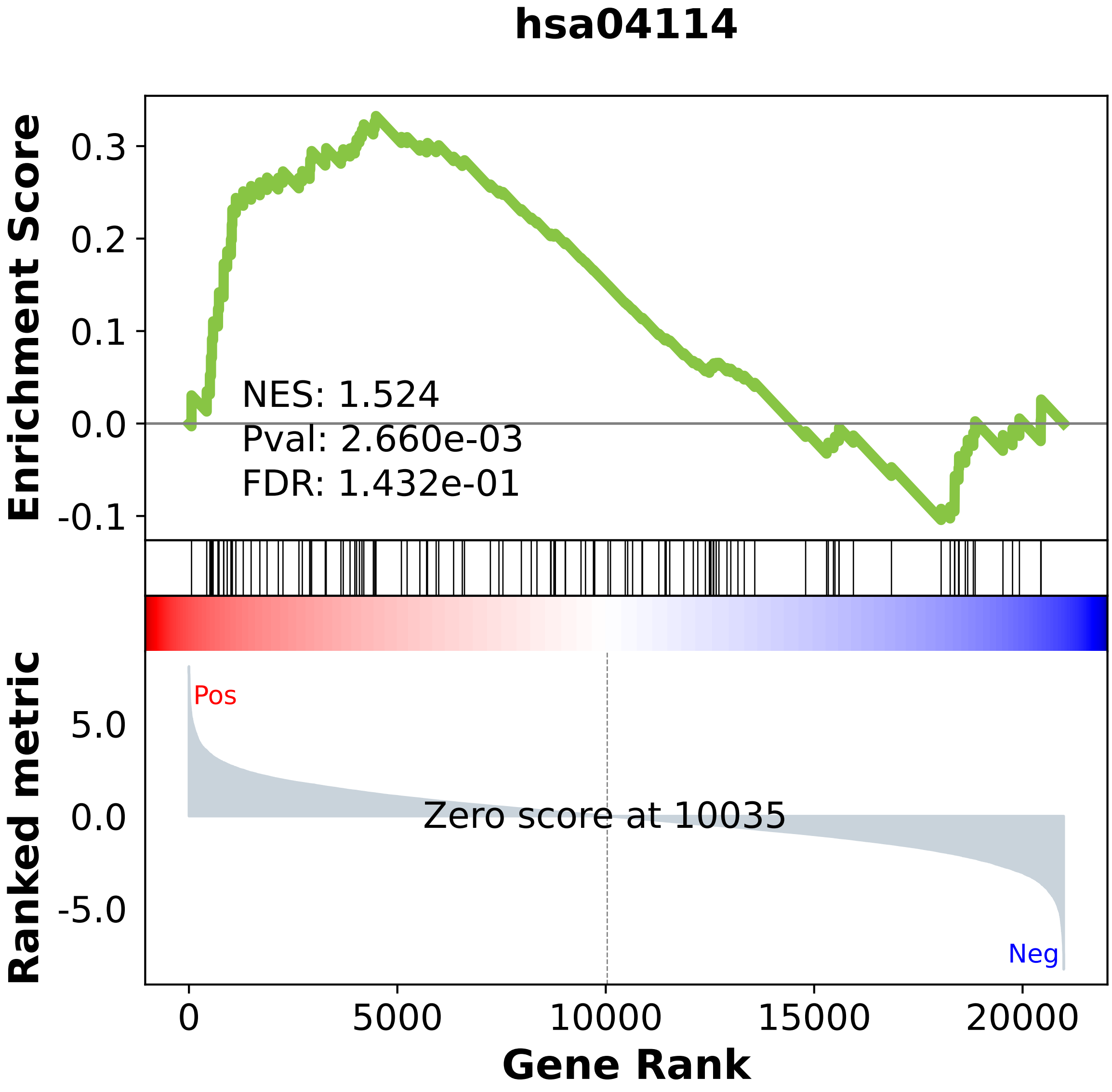

### hsa04115.pdf

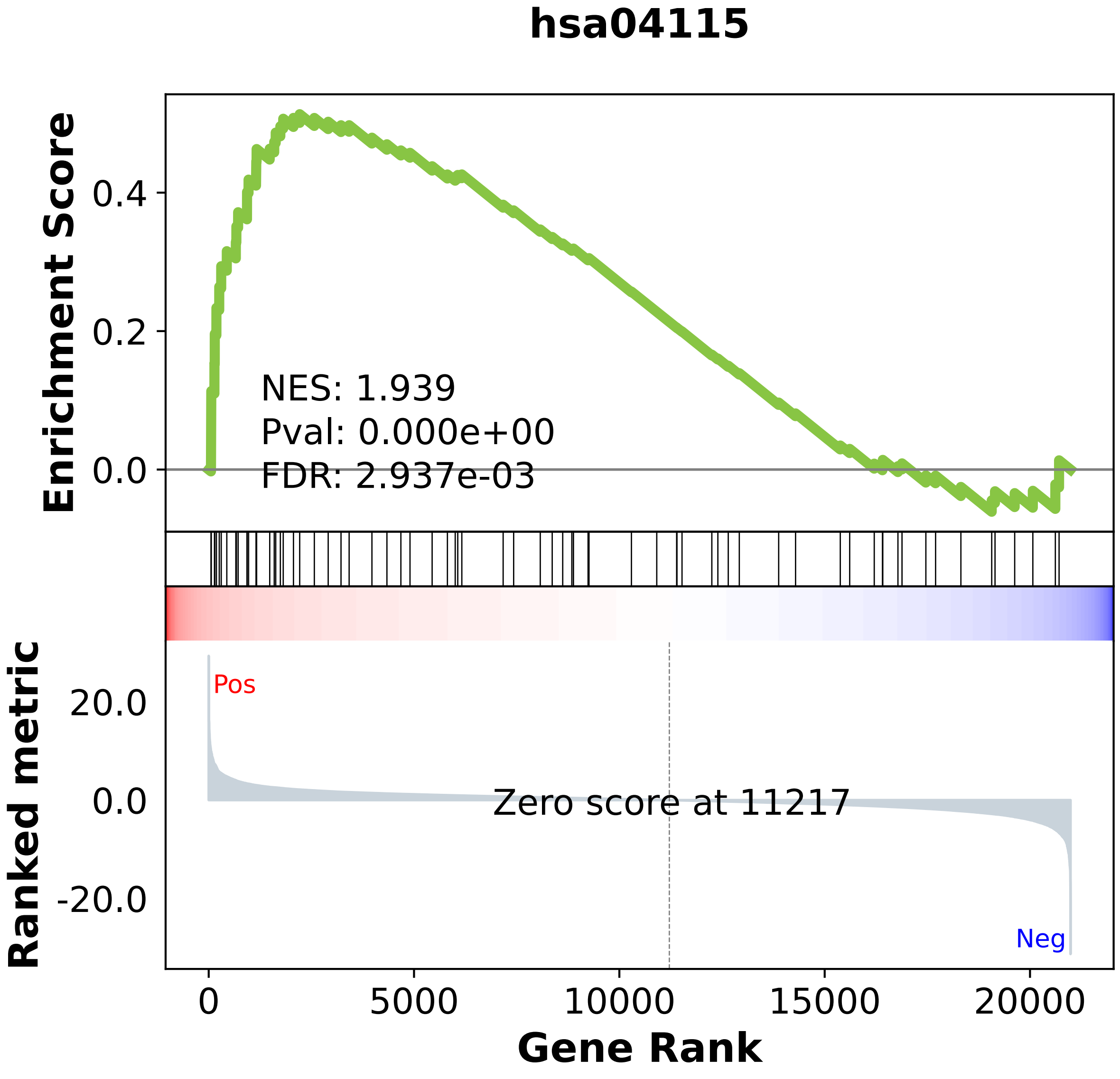

### hsa04120.pdf

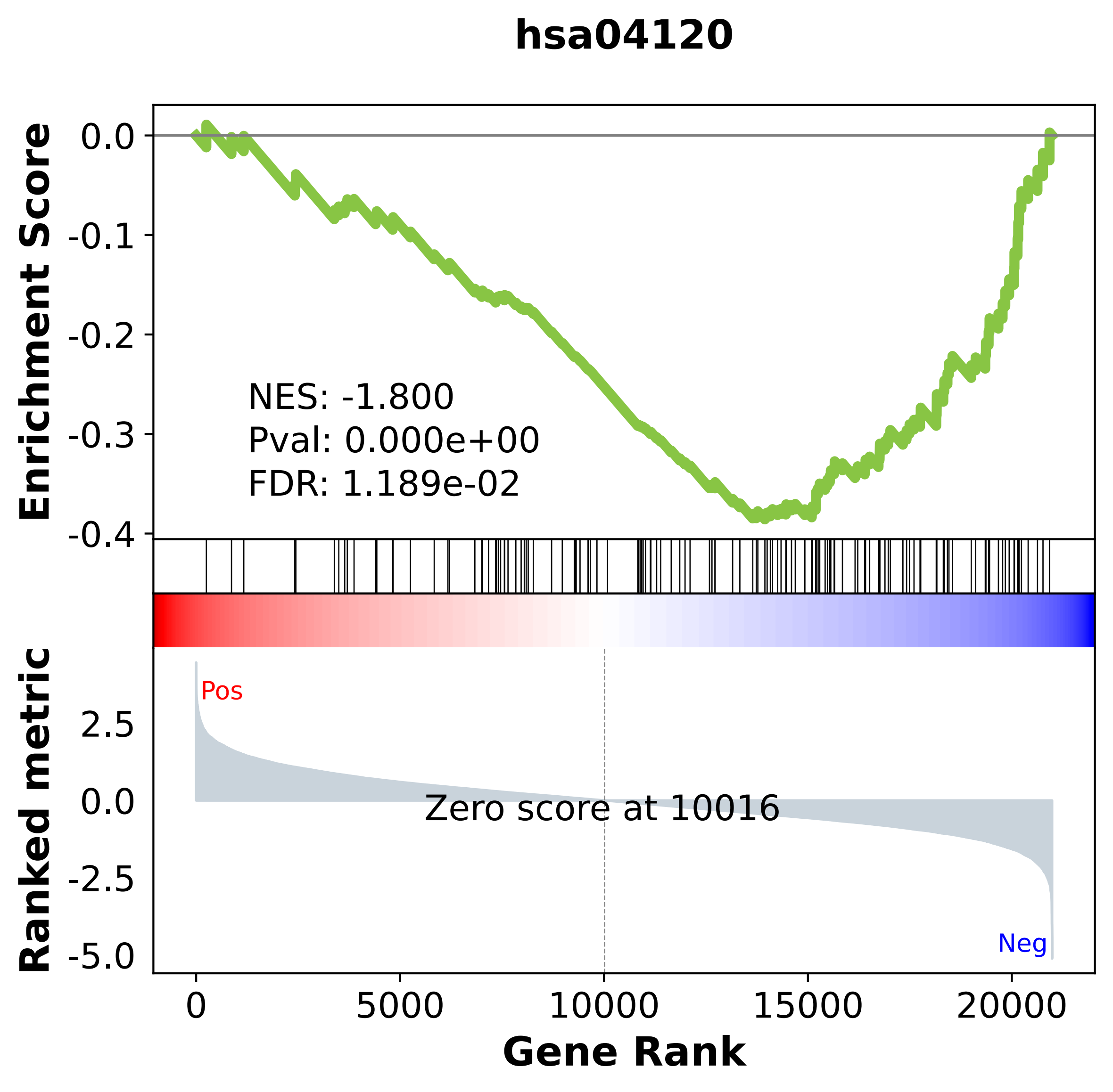

### hsa04130.pdf

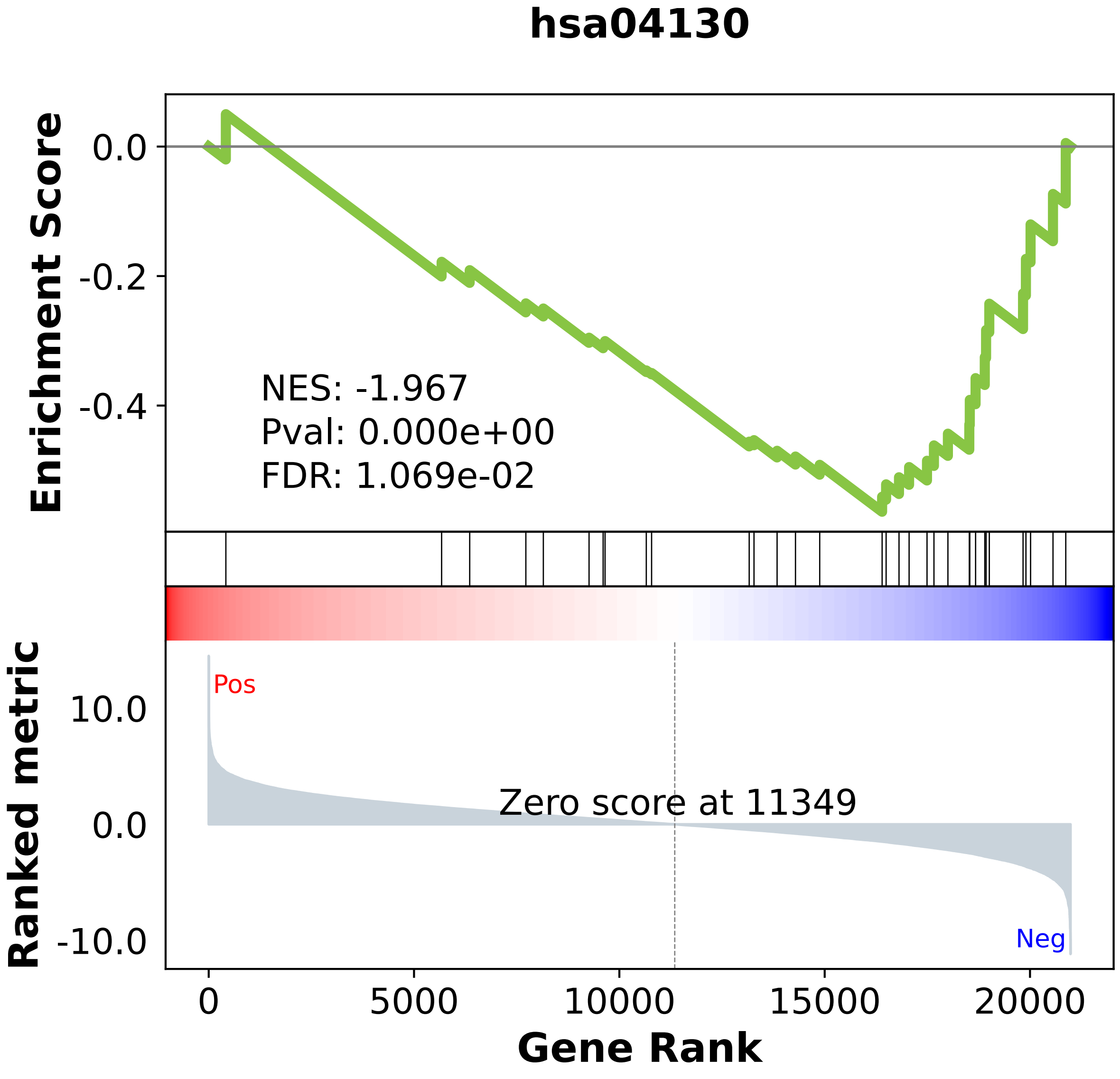

### hsa04140.pdf

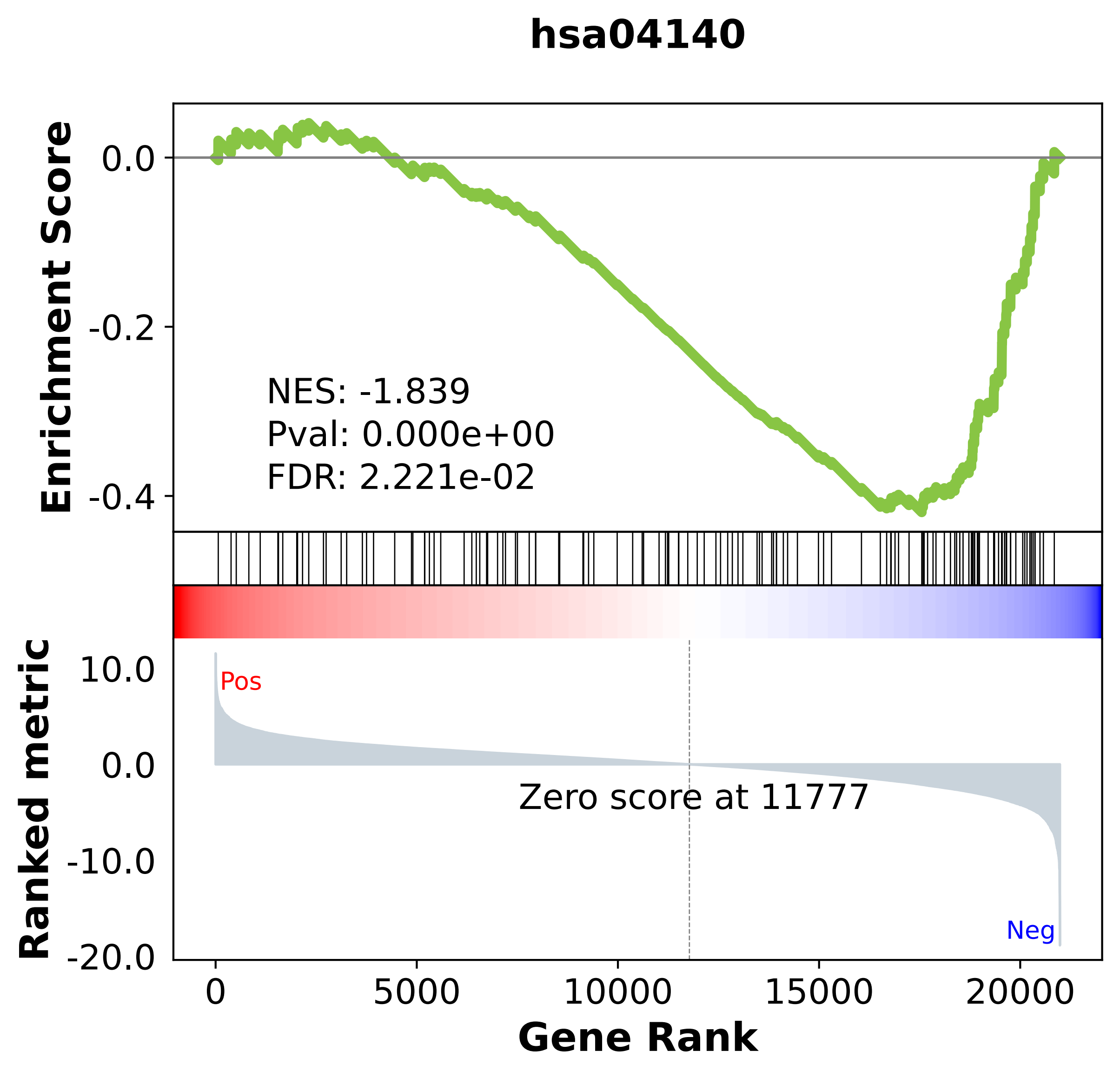

### hsa04141.pdf

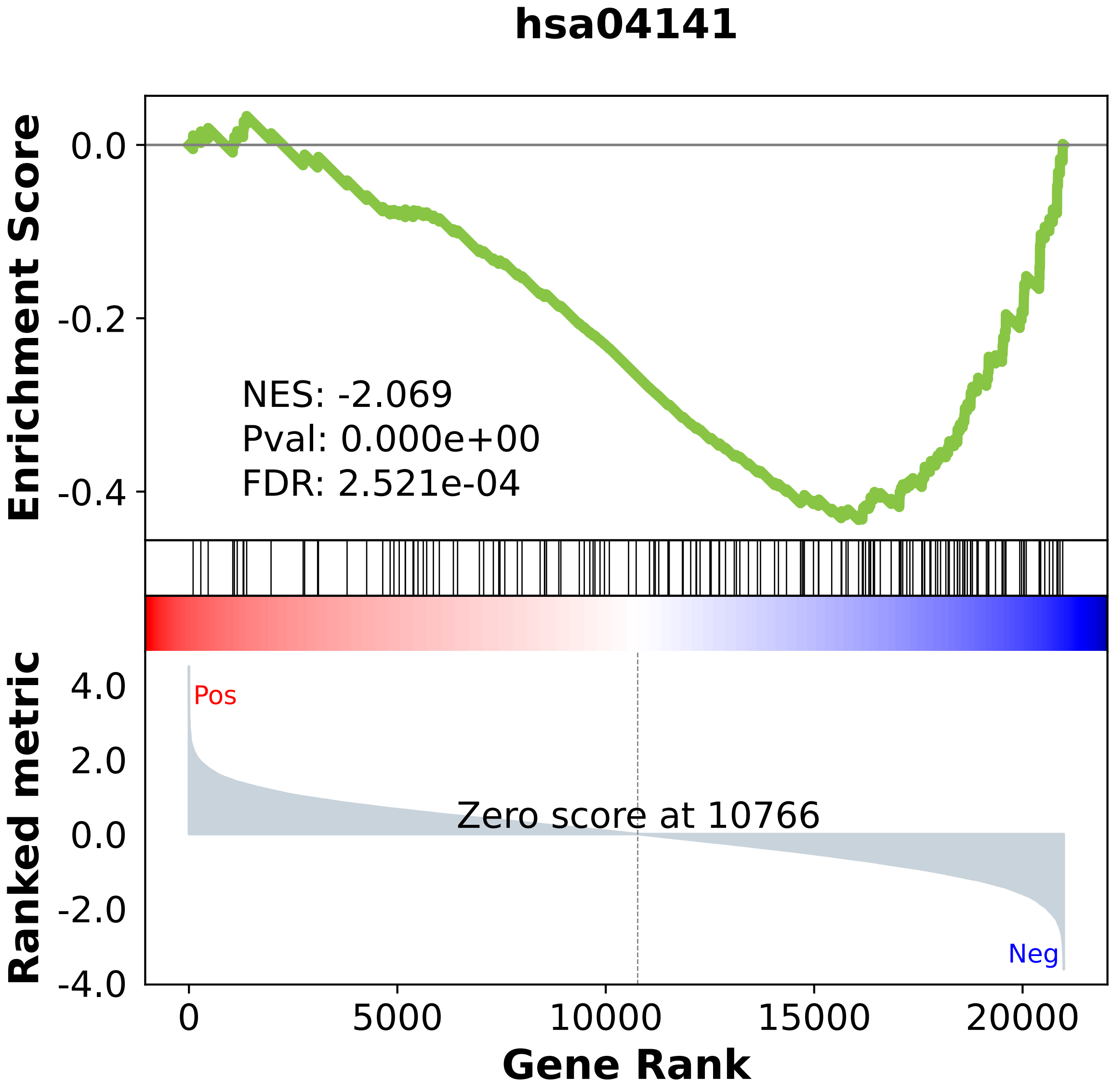

### hsa04142.pdf

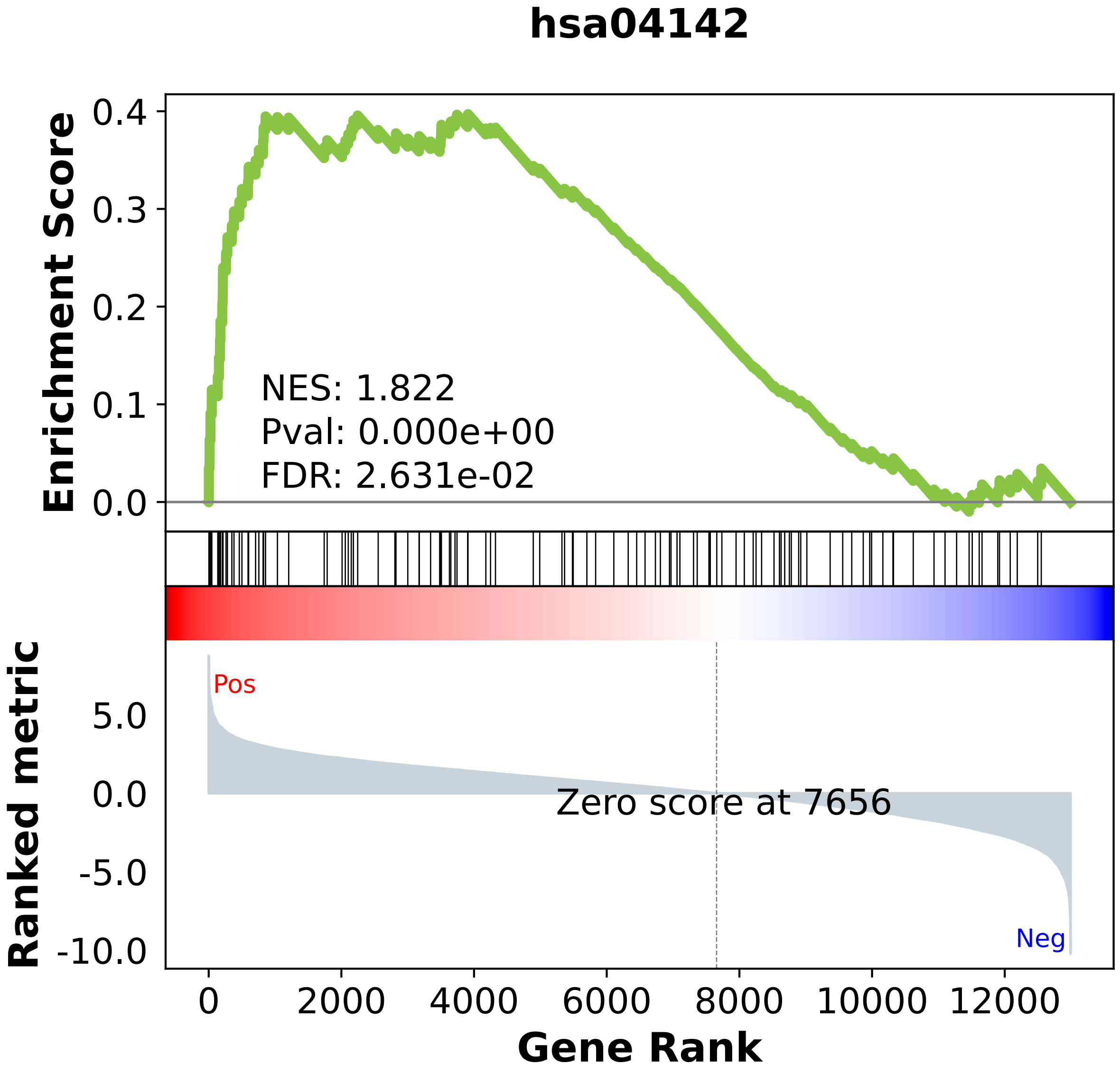

### hsa04144.pdf

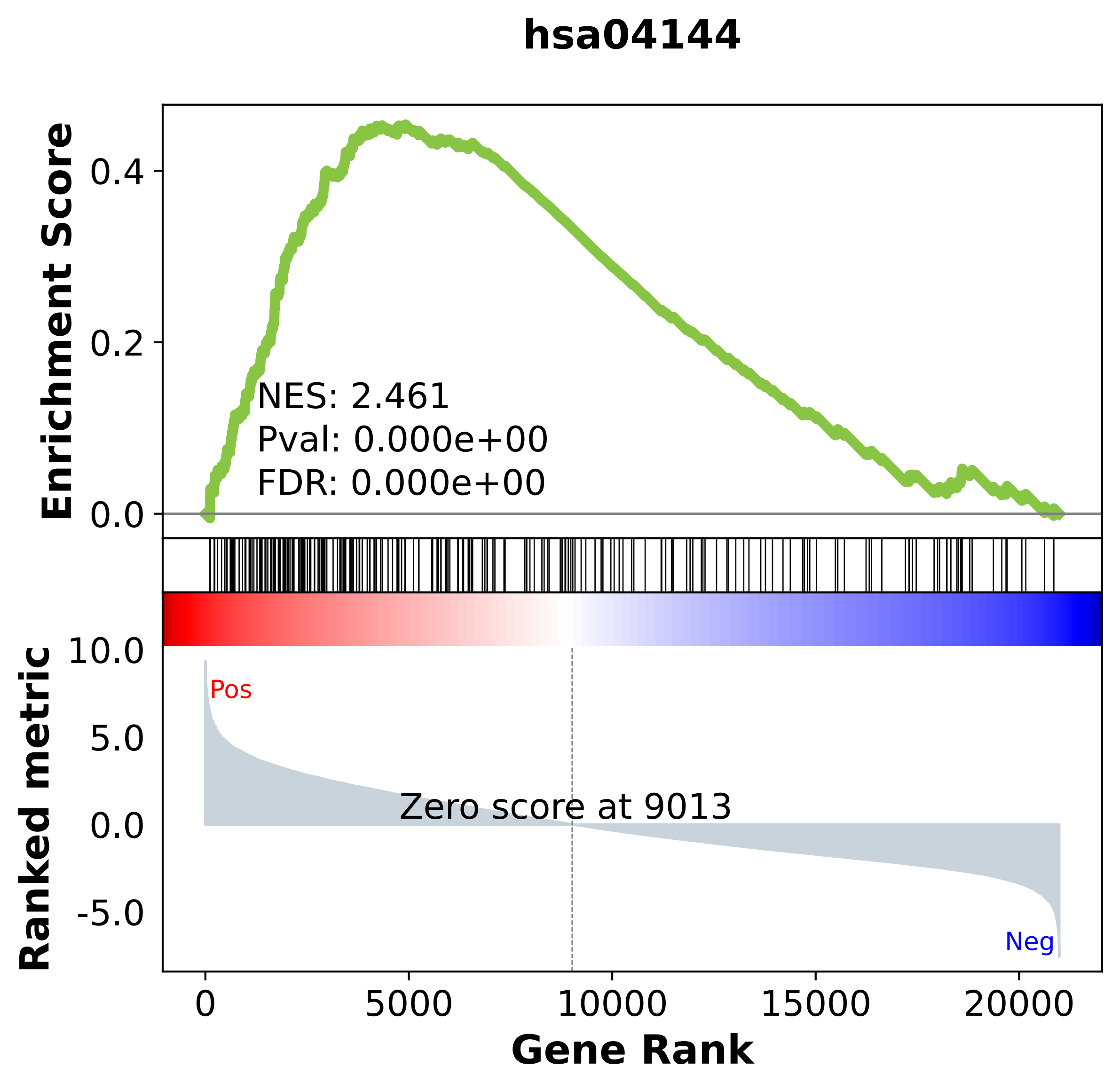

### hsa04145.pdf

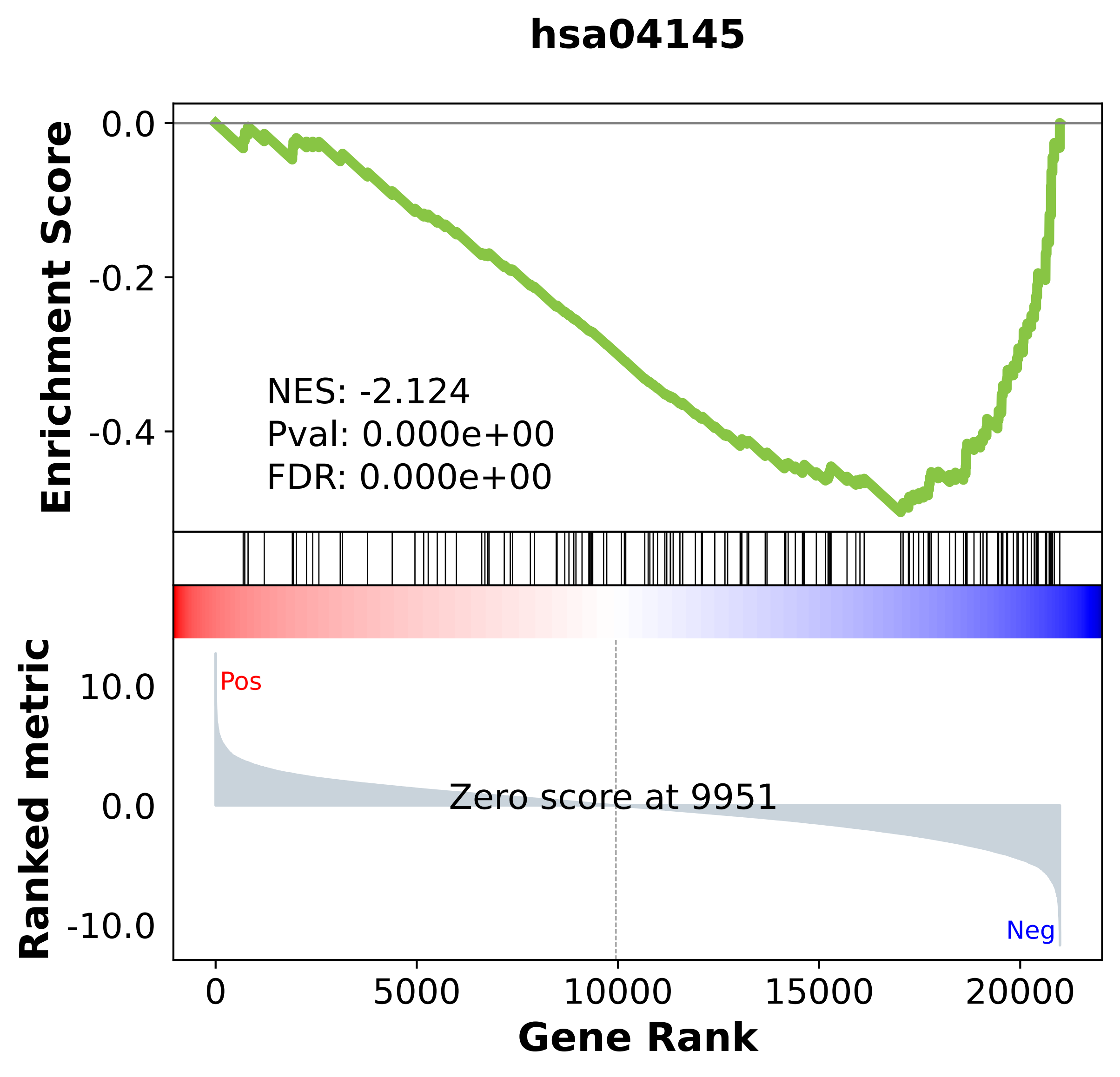

### hsa04146.pdf

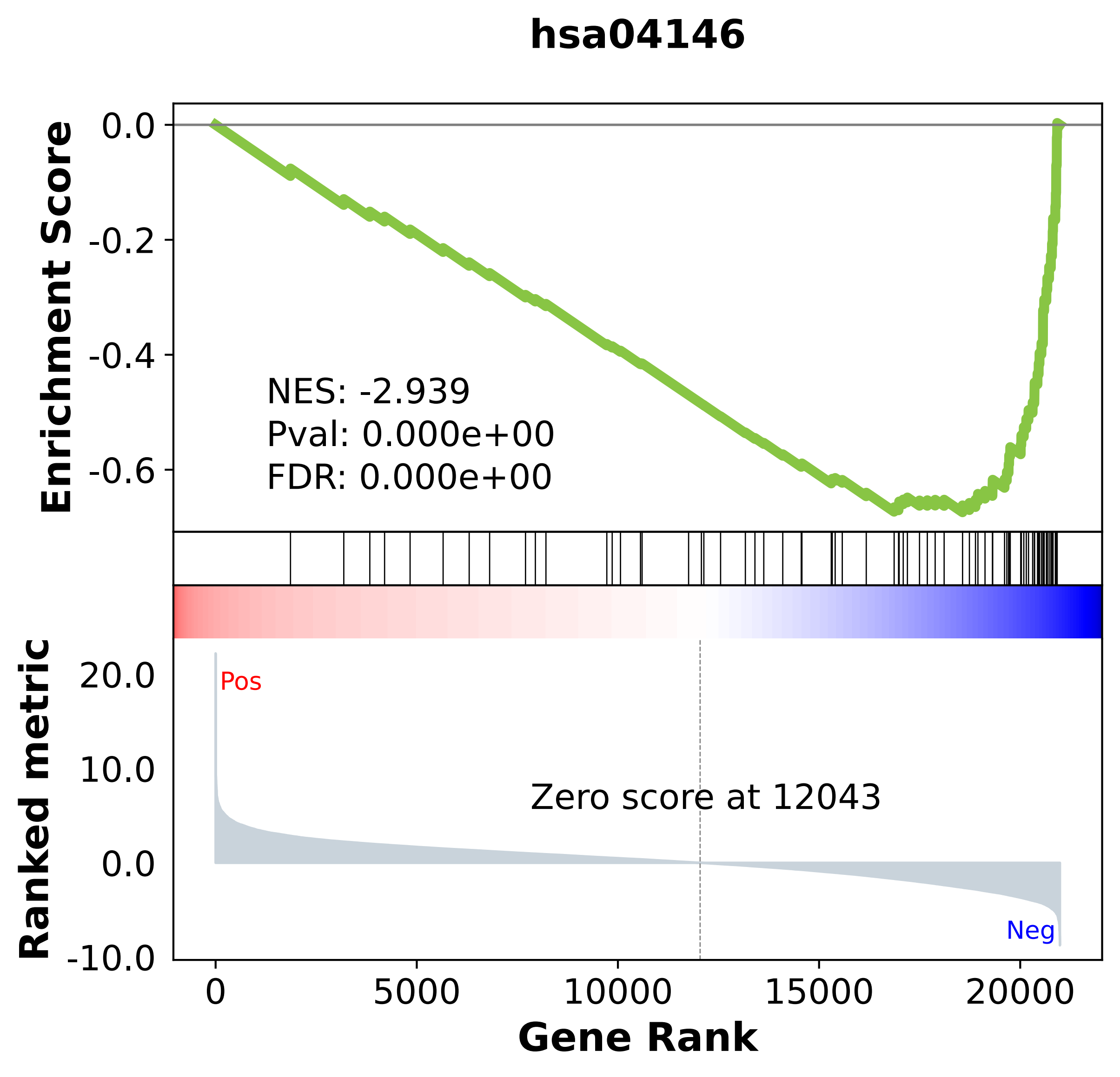

### hsa04210.pdf

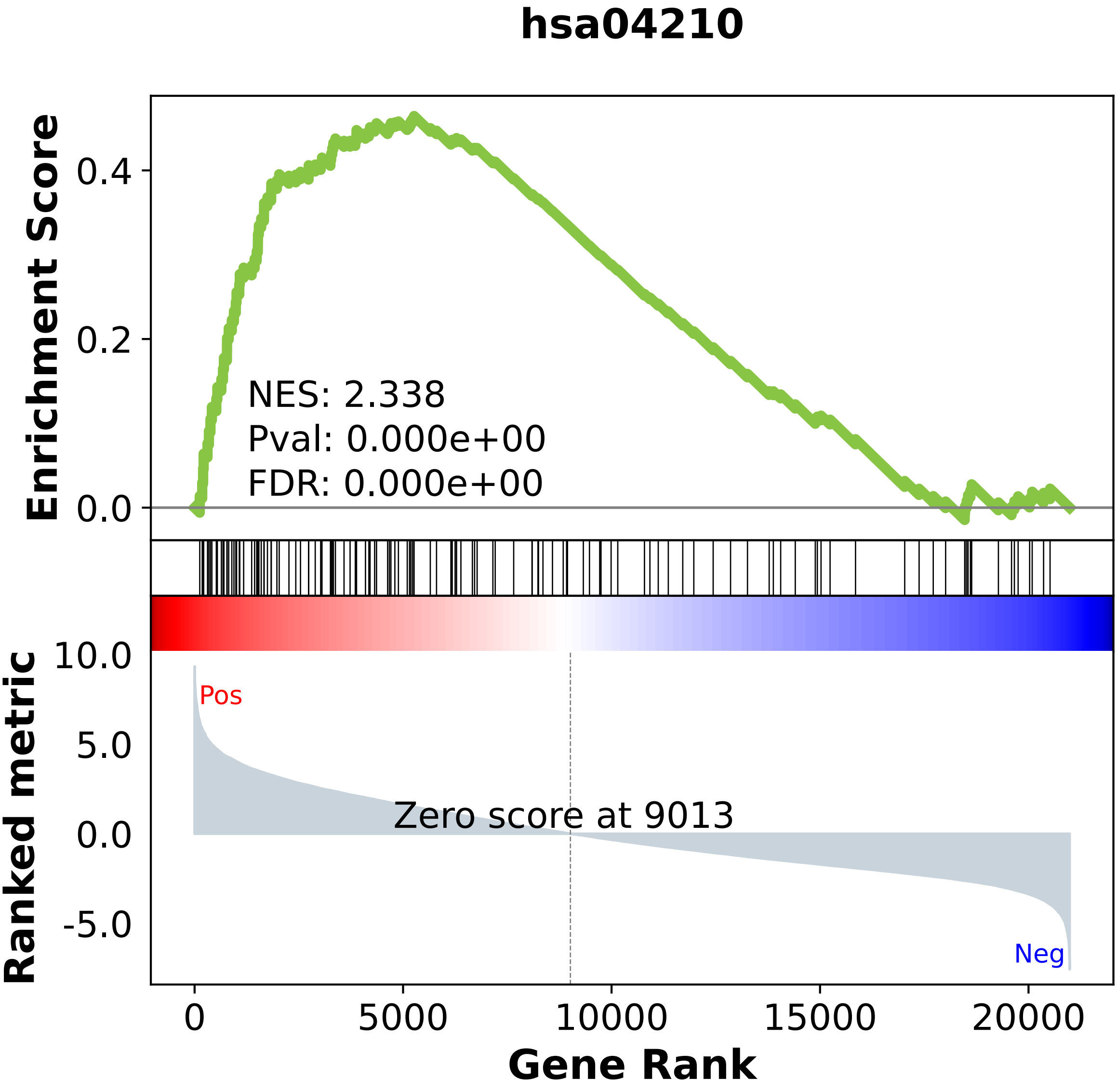

### hsa04216.pdf

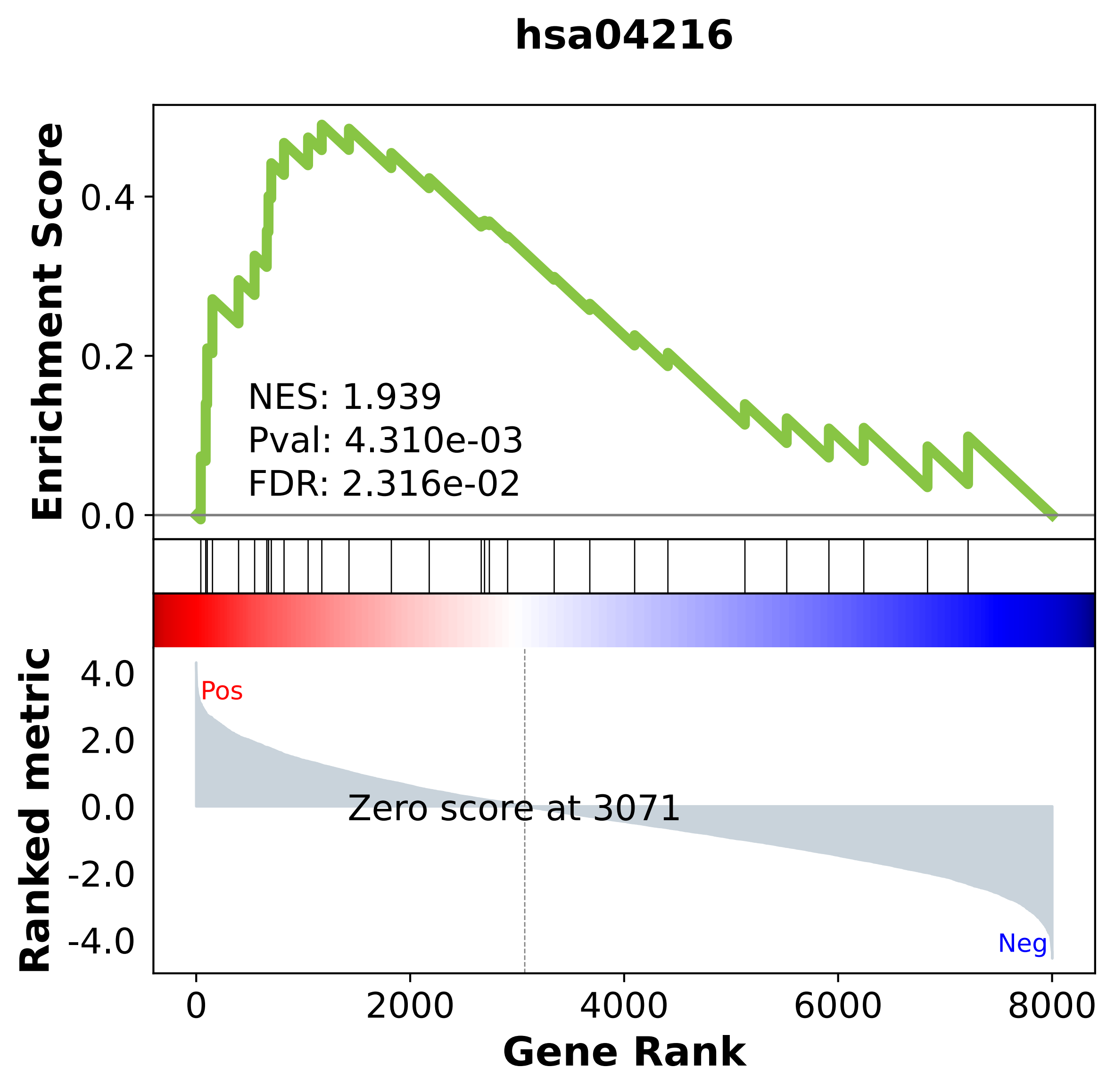

### hsa04217.pdf

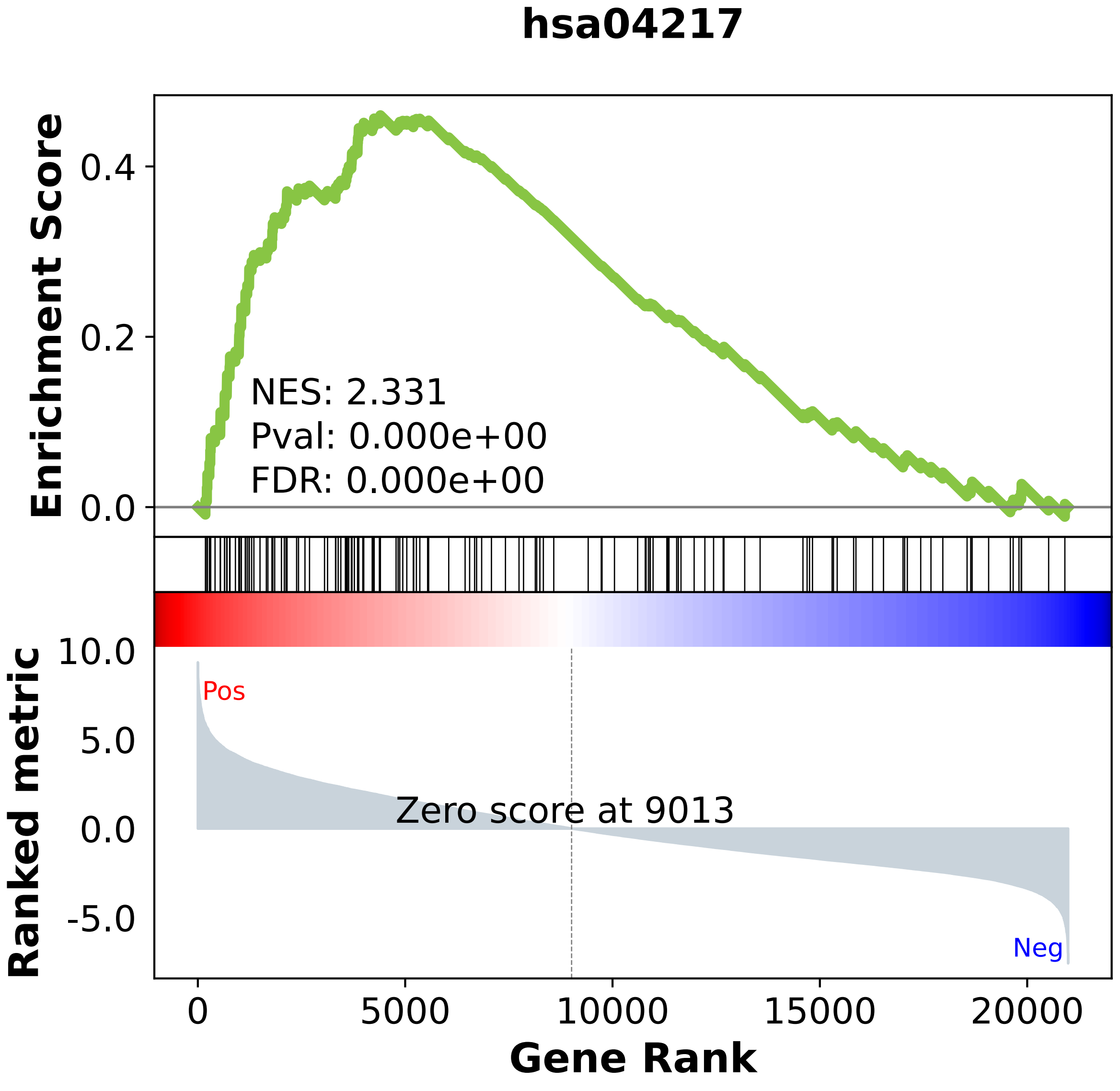

### hsa04218.pdf

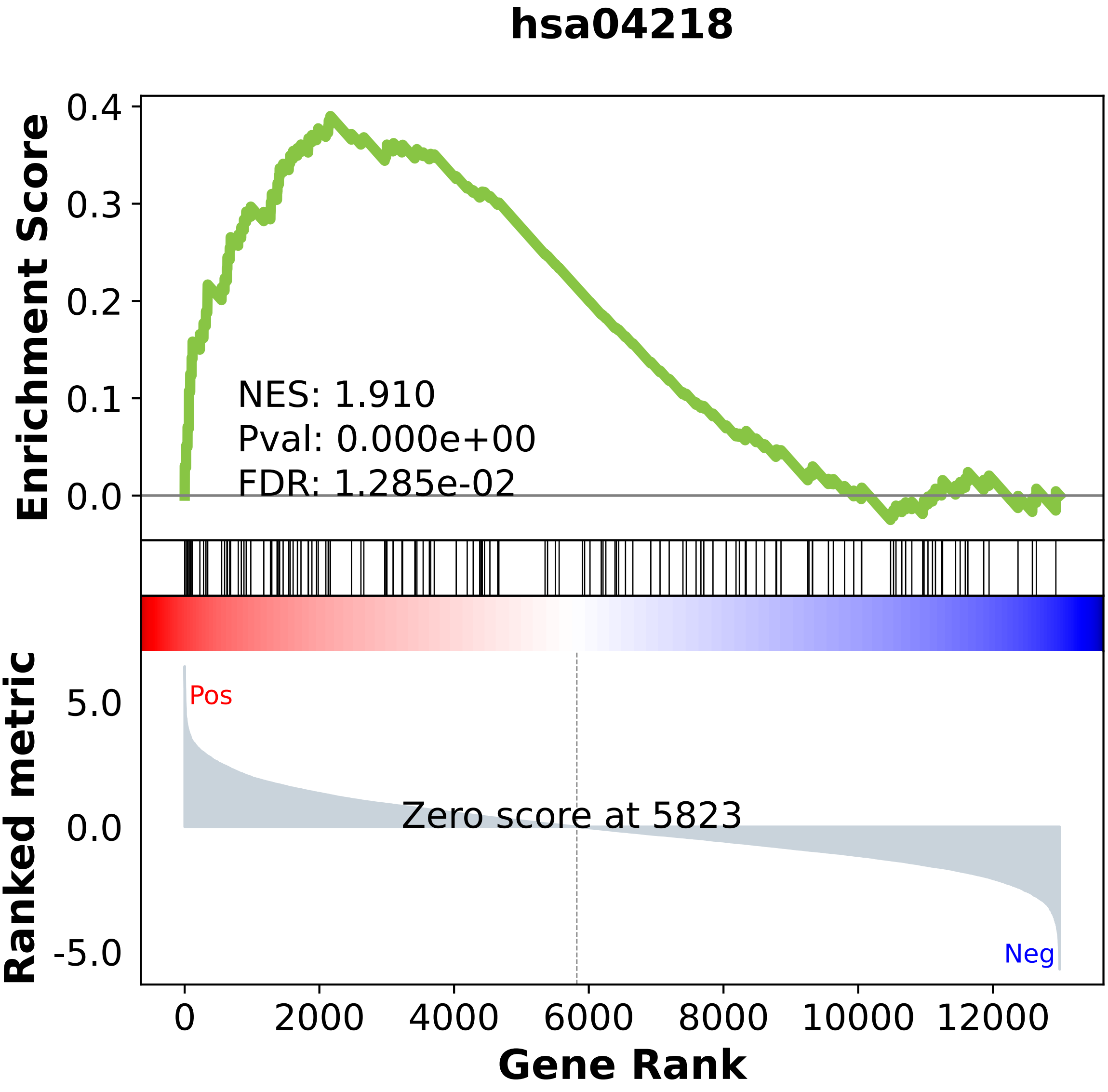

### hsa04260.pdf

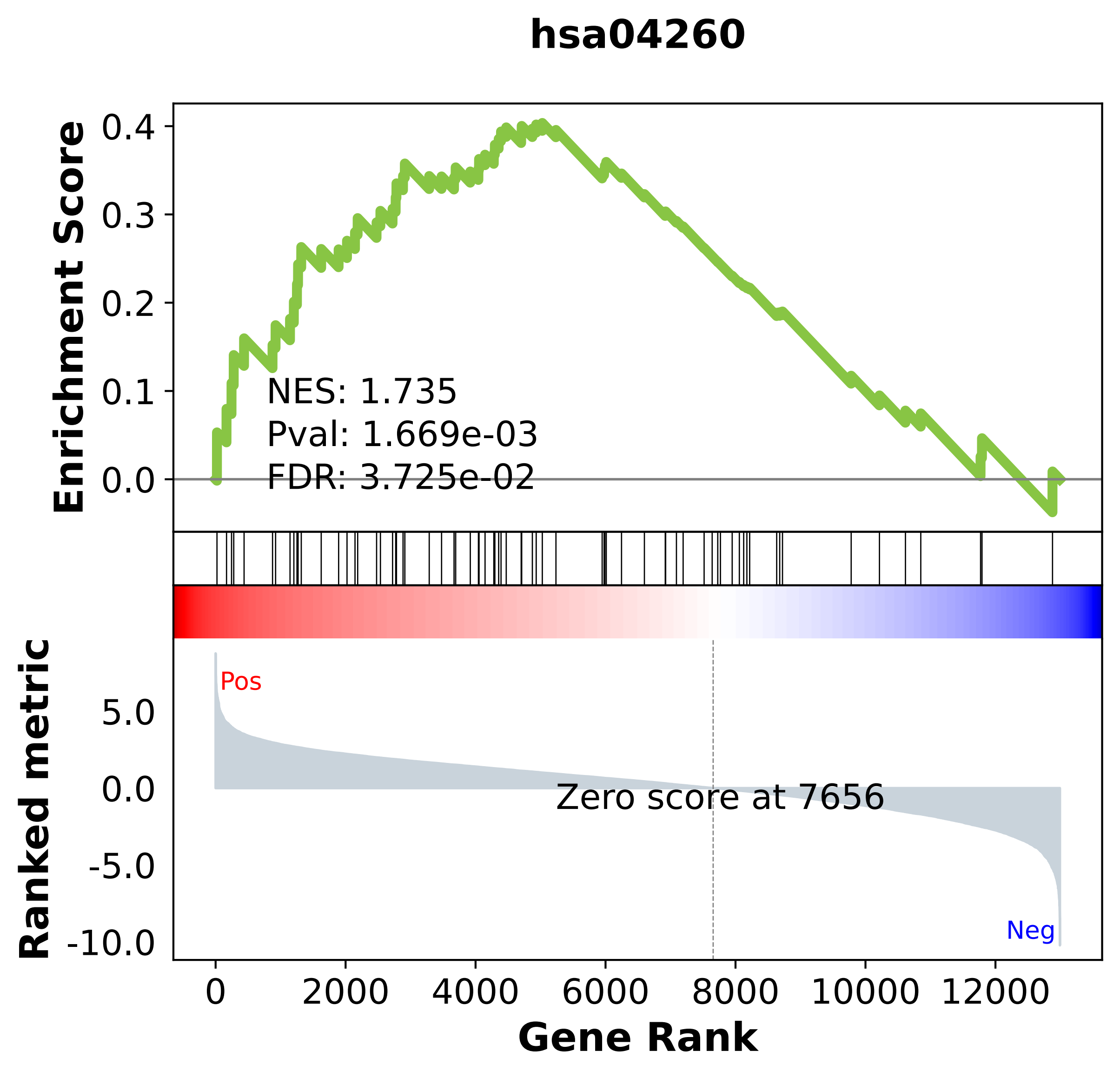

### hsa04270.pdf

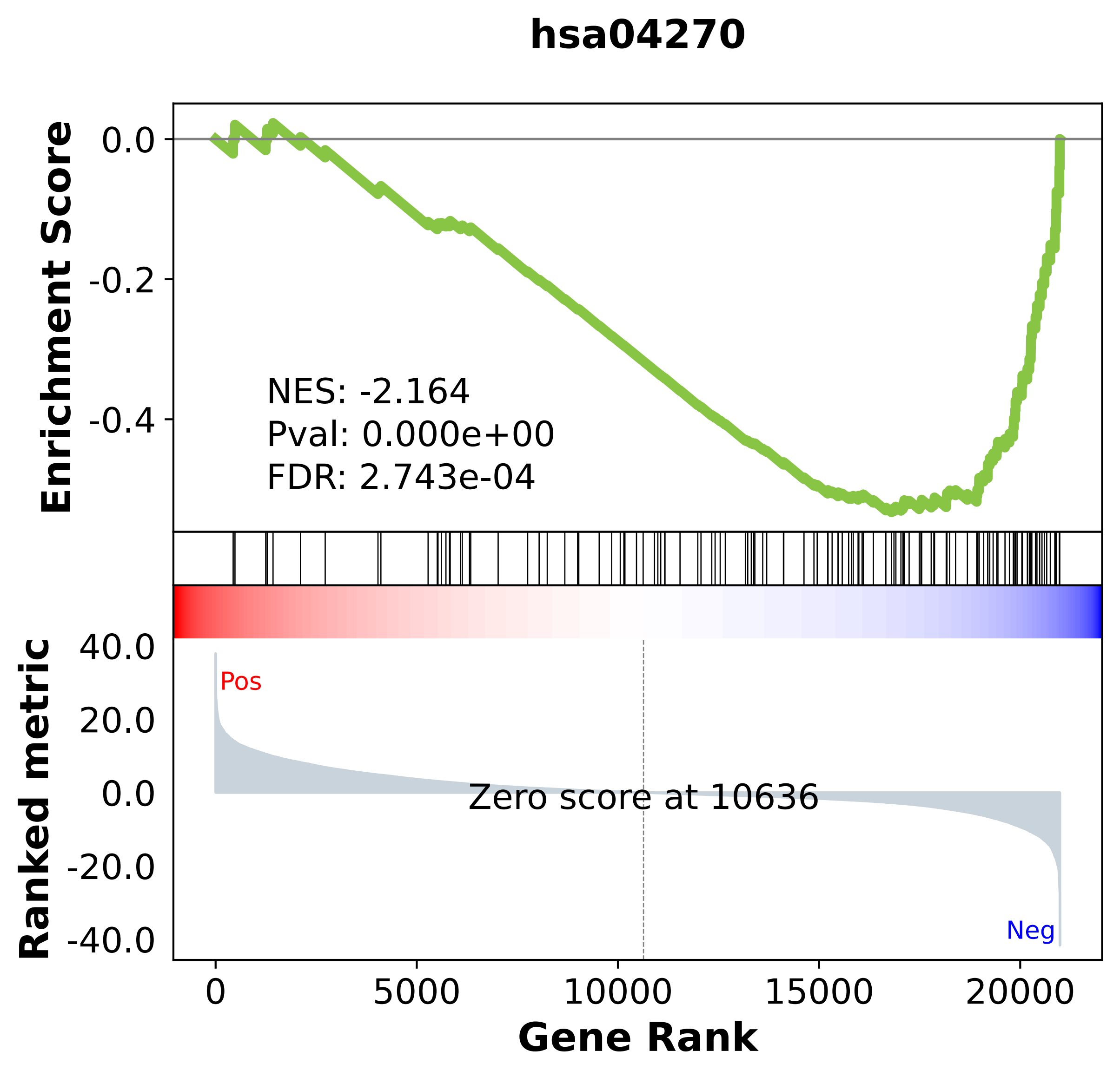
